# Changes in Ovary Transcriptome and Alternative Splicing at Estrus from Xiang Pigs with Large and Small Litter Size

**DOI:** 10.1101/547810

**Authors:** Fuping Zhang, Liangting Tang, Xueqin Ran, Ning Mao, Yiqi Ruan, Fanli Yi, Chang Liu, Xi Niu, Shihui Huang, Sheng Li, Jiafu Wang

## Abstract

**Background/Aims:** Litter size is one of the most important reproductive traits in pig breeding, which is affected by multiple genes and the environment. Ovaries are the most important reproductive organs and have a profound impact on the reproduction efficiency. Therefore, genetic differences in the ovaries may contribute to the observed differences in litter size. Although QTLs and candidate genes have been reported to affect the litter size in many pig breeds, however, the findings cannot elucidate the marked differences of the reproductive traits between breeds. The aim of present work is to elucidate the mechanisms of the differences for the reproductive traits and identify candidate genes associated with litter size in Xiang pig breed.

**Methods:** The changes in ovary transcriptome and alternative splicing were investigated at estrus between Xiang pigs with large and small litter size by RNA-seq technology. The RNA-seq results were confirmed by RT-qPCR method.

**Results:** We detected 16,219 - 16,285 expressed genes and 12 types of alternative splicing (AS) events in Xiang pig samples. A total of 762 differentially expressed genes were identified by XL (Xiang pig group with large litter size) vs XS (Xiang pig group with small litter size) sample comparisons. A total of 34 genes were upregulated and 728 genes were downregulated in XL ovary samples compared with the XS samples. Alternative splicing (AS) rates in XL samples were slightly lower than that observed in XS samples. Most of differentially expressed genes were differentially regulated on AS level. Eleven candidate genes were potentially identified to be related to Xiang pig fecundity and litter size, which may be closely related to the gonad development, oocyte maturation or embryo quality.

**Conclusion:** The significant changes in the expression of the protein-coding genes and the level of alternative splicing in estrus ovarian transcriptome between XL and XS groups probably are the molecular mechanisms of phenotypic variation in litter size.

## Introduction

In pig breeding, reproduction performance, particularly female reproductive performance, is a very important economic trait [1]. Reproductive traits are complex, and desirable reproductive phenotypes, such as litter size, are true polygenetic traits affected by interactions between multiple genes and the environment [2-5]. Each individual gene contributes to the overall variation in these traits [3, 6]. Litter size is one of the most important reproductive traits and has a great impact on the profit of pig producers [7]. As ovaries are the most important reproductive organs, they directly mediate ovulation and have a profound impact on the reproduction efficiency. Therefore, genetic differences in the ovaries may contribute to the observed differences in litter size [8-10].

To find the quantitative trait loci (QTL) and causal genes for these traits, several studies for linkages [11-13] and candidate genes ([14-17] have been conducted by using modern molecular information. Many of the major genes involved in the prolificacy of pig, such as *ESR*, *FSHB*, *RBP4*, *PRLR*, *MTNR1A*, *OPN*, *BMP* families, and *GDF9* genes have been characterized [18, 19]. Genetic studies have revealed an increasing number of associated QTLs and candidate genes involved in pig litter size [20-22]. Recently, several studies of the transcriptome for reproductive organs reveals differentially expressed genes affecting reproduction and candidate genes for litter size in European pig breeds [9, 15, 17] and Chinese indigenous pig breeds [14, 23]. Although QTLs and candidate genes have been reported to affect the litter size in many pig breeds, however, the studies were only focused on specific species or breeds and the findings cannot elucidate the marked differences of the reproductive traits between Asian and European pig breeds and the variations of litter size between pigs.

Chinese indigenous Xiang pig is one of the miniature pig breed originated from the mountain region in south east of Guizhou province. Small size, early sexual maturity, disease resistance, good meat quality and crude feed tolerance are the basic distinctively biological characteristics [24, 25]. Large phenotypic variation in litter size was observed within Xiang pig sows [26]. To analyze changes in gene expression across the entire transcriptome and to identify key genes relating to litter size, in the present study, we collected ovary samples of Xiang pigs having previously a large or small litter size. Then at estrus after weaning, a genome wide analysis of transcriptome and alternative splicing events of transcripts using RNA-seq technology was performed. Our results will give an insight of understanding on the molecular mechanisms about Xiang pig fecundity and litter size.

## Materials and Methods

### Ethics statement

Xiang pigs were obtained from Guizhou Dachang pig breeding Farm, in Congjiang county, Guizhou province, China. All animal procedures were approved by the Institutional Animal Care and Use Committee of Guizhou University (GZU-201709), and were conducted in accordance with the National Research Council Guide for Care and Use of Laboratory Animals.

### Animals and ovary collection

A total of forty Xiang pig gilts were used in this study from the sows that previously had a large litter size (XL: n = 20, TNB > 12), and a small litter size (XS: n = 20, TNB < 8), representing pigs with high and low fecundity, respectively. The gilts were selected after their first estrus cycle. The animals were reared in the same environment and fed the same diet under a standardized feeding regimen with free access to water during the experimental period. Information on the pedigree, breeding and performance of breeding animals was obtained from the farm records. After weaning, estrous detection was carefully performed twice to three times daily at proestrous and tested by a boar. Standing reflex was assessed by the back-pressure test. Reddening and swelling of the vulva were measured visually. The expression of estrus was scored according to a standard scoring system [14, 27]. In this study, we randomly chose six Xiang gilts from the XL (n = 3; TNB mean = 16.16) and XS (n = 3; TNB mean = 6.78) groups to perform the genome-wide analysis of transcriptomes by RNA-seq. The ovarian samples from 6 Xiang pig gilts were collected with surgery at the day of the third estrous when the gilts exhibited strong expression of reddening and swelling of the vulva, and standing reflex. Ovaries were dissected and collected with better ovulation points or preovulatory follicles on the surfaces of the ovaries. All samples were immediately frozen in liquid nitrogen and stored at −80 °C until total RNA was isolated.

### Library preparation and sequencing

Total RNA was extracted from the ovarian samples in the two groups with Trizol reagent (TIANGEN, China) following the manufacturer’s instructions, and then the DNase I was used to degrade any possible DNA contamination. The quantity and integrity of the total RNAs were assessed using the Agilent 2100 Bio-analyzer (Agilent Technologies, CA, USA). Total RNAs were stored at −80 °C until subsequent analysis. The same sample was used to both the sequencing and RT-qPCR analysis. The oligo (dT) magnetic beads were used to enrich mRNA. The purified mRNA was broke into short fragments by adding fragmentation buffer in the condition of appropriate temperature, and then cDNA was synthesized using random hexamer primers and mRNA fragments as templates. After adenylation of 3’ ends, DNA fragments were ligated to adapters. The cDNA fragments with 100–200 bp in length were selected for PCR amplification to generate cDNA libraries. The library preparations were sequenced on an Illumina HiSeqTM 2500 platform and generated 90 bp paired-end reads according to the manufacturer’s protocols at the Beijing Genomies Institute (BGI), Shenzhen, China.

### Data analysis

The raw sequence reads were generated by the Illumina platform and saved as a format of fastq. After removing low quality reads (more than 5% unknown nucleotides) and sequencing adapters and being assessed by the FastaQC software (Q20 < 20%), the clean reads were obtained. The pig reference genomes (Ssc 11.1) and the annotation files were downloaded from ENSEMBL database (http://www.ensembl.org/index.html). The genome index was built by the Bowtie2 (v2.2.3) software. TopHat2 (v2.0.12) software was used to align the clean reads onto the pig reference genome, which the maximum mismatch was not more than 2 bp. The Cufflinks software was used to assemble transcripts and to predict gene model. The HTSeq software was used to count the number of reads which aligned to each gene and exon. FPKMs (Fragments Per Kilobase of transcript per Million mapped reads) were then calculated to estimate the expression values of genes and alternative splicing variants. Asprofile (v1.0.4) software was used to classify and count the alternative splicing events in each sample. Venny online tool (http://bioinfogp.cnb.csic.es/tools/venny/index.html) was used for the statistics of the number of genes in each sample.

### Identification of differentially expressed genes and differential alternative splicing events

The Bio-conductor package limma [28] was used for analysis of differential gene expression and differential alternative splicing between two samples with biological replicates using a model based on the negative binomial distribution. The P-value had been assigned to each gene or AS and adjusted by the Benjamini and Hochberg’s approach for controlling the false discovery rate. P-value ≤ 0.01, adj P-value ≤ 0.05 and the absolute value of log2 Ratio ≥1 were employed as the threshold to determine the differential expression genes or differential alternative splicing events.

### GO enrichment analysis

Enrichment analysis of Gene ontology (GO) annotation was conducted on differentially expressed genes at gene or isoform level using Gene Ontology Consortium online (http://geneontology.org/).

### RT-qPCR validation

RNA samples from the 6 gilts in the RNA-seq experiment were used to validate the results by quantitative real-time RT-PCR (qPCR). After removing genomic DNA by RNase-free DNase (New England BioLabs, USA), the total cDNA was synthesized using the First Strand cDNA Synthesis Kit (Thermo Fisher Scientific, USA) according to the manufacturer’s protocol. Real-time PCR reactions were performed as described by the manufacturer in triplicate with Power SYBR Green PCR Master Mix on the BIO-RAD CFX98 Real-Time System (BIO-RAD, US). The reaction was performed in a 96-well plate. Each reaction contained 5 μL of SYBR Green PCR Master Mix, 0.4 μL of the forward and reverse primers (10 pM/μL), 1.0 μL of cDNA (5 ng/μL) and 3.6 μL of distilled water. Thermal cycling conditions consisted of an initial denaturation at 95 °C for 10 min, followed by 40 cycles of denaturation at 94 °C for 15 s, annealing at 60 °C for 30 s, and then a 55–95 °C melting curve detection. The *GAPDH* gene was used as a control in the experiments. All amplifications were followed by dissociation curve analysis of the amplified products. Relative expression levels of the selected target genes were calculated with the 2(-Delta Delta C[T]) method [29]. The statistical difference in gene expression between groups was analyzed by SPSS software (v21.0). The results were presented as mean ± standard deviation. The correlation between the results of RNA-seq and qPCR was calculated using correlation test (Pearson correlation) by SPSS Software (v21.0). Primer informations can be found in additional file: Table 5.

## Results

### Transcriptome sequencing and assemble

We performed an RNA-seq analysis on ovary transcriptome from Xiang pigs with large and small litter size. The cDNA libraries from 6 samples were sequenced using Illumina HiSeqTM 2500 platform. The total number of clean reads with maximum of 90 base-pair (bp) ranged from 49.9 to 50.39 million were produced after quality control and filtering. The reads were mapped by TopHat2 (version 2.0.12) to *Sus scrofa* reference genome (Ssc11.1). 81.5% to 84.6% clean reads were mapped onto the *Sus scrofa* reference genome respectively for each sample (Table 1). The unique mapped reads were used for subsequent analysis. FPKM was used to estimate the expression at gene and isoform levels.

**Table 1.**
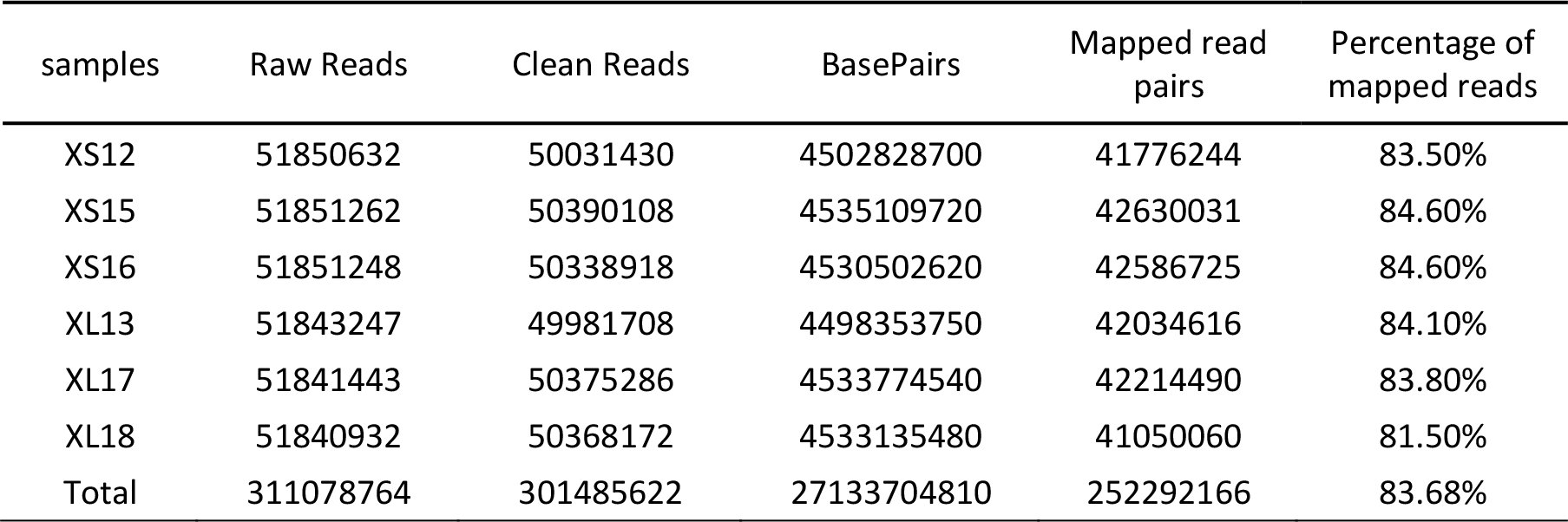
Summary of read mapping.

| samples | Raw Reads | Clean Reads | BasePairs | Mapped read pairs | Percentage of mapped reads |
| --- | --- | --- | --- | --- | --- |
| XS12 | 51850632 | 50031430 | 4502828700 | 41776244 | 83.50% |
| XS15 | 51851262 | 50390108 | 4535109720 | 42630031 | 84.60% |
| XS16 | 51851248 | 50338918 | 4530502620 | 42586725 | 84.60% |
| XL13 | 51843247 | 49981708 | 4498353750 | 42034616 | 84.10% |
| XL17 | 51841443 | 50375286 | 4533774540 | 42214490 | 83.80% |
| XL18 | 51840932 | 50368172 | 4533135480 | 41050060 | 81.50% |
| Total | 311078764 | 301485622 | 27133704810 | 252292166 | 83.68% |

### Changes of ovary transcriptome in XL and XS

After mapping onto *Sus scrofa* reference genome, in total, 16,454 genes were detected in all transcriptomes, including 13,899 known genes and 2,555 un-annotated genes (Supplementary Table S1, Fig 1). Among of those, 16,219 and 16,285 genes were detected from the XL and XS libraries, respectively. 16,050 genes were co-expressed in both XL and XS libraries, One hundred and sixty nine genes were exclusively expressed in XL libraries, and two hundred and thirty five genes were preferentially expressed in XS libraries (Fig 1). The number of un-annotated gene in each library from XL to XS was 2,400, and 2,441, respectively. The expression levels of those genes varied greatly, ranging from less than 0.1 FPKM to more than hundreds of thousands of FPKM (Supplementary Table S1). There were 27 top expressed genes in XL and/or XS samples respectively (FPKM > 1,000) (Table 2). 17 of 27 genes including *TPT1*, *EEF1A1*, *TMSB10*, *FTH1* and 11 members of ribosomal protein genes, were overlapped between XL and XS samples. Five genes (*GPX3, ACTG1, TIMP1*, *COL1A2*, and *VIM)* were dominantly expressed in XL libraries. Two genes (*CLU*, and *OVGP1*) were the top expressed genes in XS libraries. The sequencing frequency of the top expressed genes constituted 11.7 ∼ 15.5% of the total expressed values in XL and XS samples, respectively. Most of the genes specific for XL or XS were showed low or very low expression levels.

**Figure 1.**
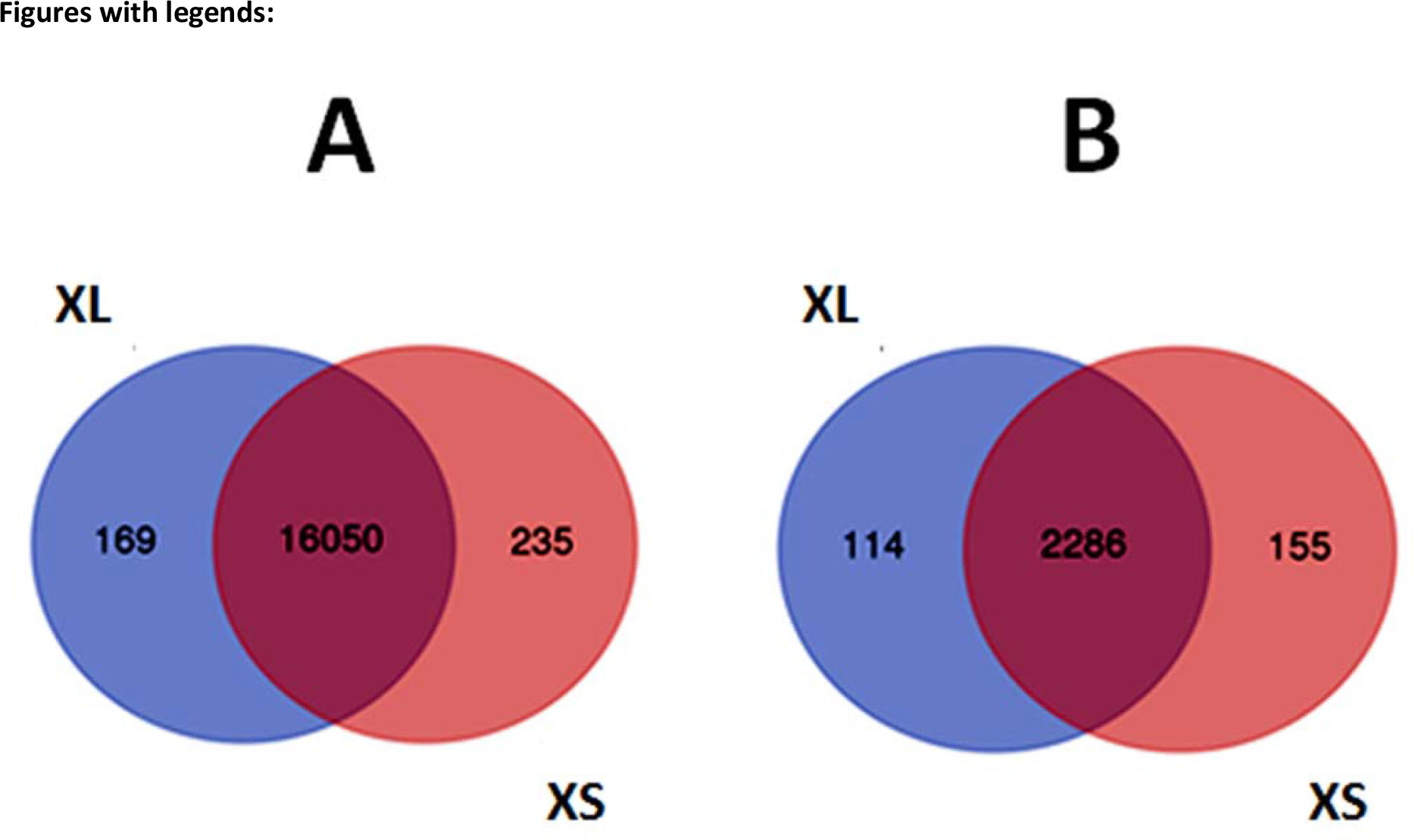
Profile of the gene expression between the ovaries of Xiang pigs with large (XL) and small (XS) litter size. Blue color represent genes only expressed in the XL group, red color show genes only expressed in the XS group, and common to both groups (intersection). A: Total of the expressed genes. B: Uncharacterized protein genes.

**Table 2.** The top highly expressed genes (FPKM > 1,000)

| Gene ID | Gene symbol | XL FPKM | XS FPKM | LogFC | P.Value | adj.P.Val | Main Function |
| --- | --- | --- | --- | --- | --- | --- | --- |
| ENSSSCG00000036438 | GPX3 | 1396.32 | 326.93 | 2.0945 | 0.0039 | 0.0597 | oocyte maturation, embryo quality |
| ENSSSCG00000028355 | ACTG1 | 1412.21 | 817.74 | 0.7882 | 0.0077 | 0.0826 | cellular response to interferon-gamma |
| ENSSSCG00000009668 | CLU | 253.88 | 1125.66 | -2.1485 | 0.0238 | 0.1477 | extracellular chaperone |
| ENSSSCG00000035997 | RPL10 | 1092.14 | 1392.33 | -0.3503 | 0.0399 | 0.1978 | translation, embryonic development |
| ENSSSCG00000004489 | EEF1A1 | 3534.45 | 4202.53 | -0.2497 | 0.0471 | 0.2134 | protein biosynthesis |
| ENSSSCG00000013597 | RPS28 | 2076.56 | 2321.82 | -0.1611 | 0.0498 | 0.2195 | translation |
| ENSSSCG00000037674 |  | 1544.21 | 1855.19 | -0.2646 | 0.0577 | 0.2356 | unknown |
| ENSSSCG00000014540 | FTH1 | 2907.67 | 1863.75 | 0.6416 | 0.0606 | 0.2418 | iron ion transport |
| ENSSSCG00000006791 | OVGP1 | 128.61 | 6752.92 | -5.7145 | 0.0684 | 0.2565 | carbohydrate metabolic process |
| ENSSSCG00000011033 | VIM | 1430.62 | 759.72 | 0.9131 | 0.0731 | 0.2653 | astrocyte development |
| ENSSSCG00000015326 | COL1A2 | 1192.61 | 634.94 | 0.9094 | 0.0801 | 0.2773 | blood vessel development |
| ENSSSCG00000024260 | RPL11 | 961.26 | 1061.56 | -0.1431 | 0.0877 | 0.2911 | translation |
| ENSSSCG00000012277 | TIMP1 | 1498.91 | 235.32 | 2.6711 | 0.0917 | 0.2972 | positive regulation of cell proliferation |
| ENSSSCG00000040929 | TPT1 | 2546.01 | 3172.08 | -0.3171 | 0.1172 | 0.3364 | cell differentiation |
| ENSSSCG00000035904 | RPL7A | 1121.11 | 1273.71 | -0.1841 | 0.1337 | 0.3607 | ribosome biogenesis |
| ENSSSCG00000038507 | RPS29 | 2071.41 | 2258.68 | -0.1248 | 0.1569 | 0.3919 | translation |
| ENSSSCG00000008245 | TMSB10 | 1434.32 | 1046.39 | 0.4549 | 0.1745 | 0.4142 | actin filament organization |
| ENSSSCG00000003042 | RPS19 | 957.37 | 1023.71 | -0.0966 | 0.2905 | 0.5415 | translation |
| ENSSSCG00000036014 | RPLP0 | 1191.31 | 1365.74 | -0.1971 | 0.2993 | 0.5503 | ribosome biogenesis |
| ENSSSCG00000003930 | RPS8 | 924.33 | 1012.12 | -0.131 | 0.3136 | 0.5641 | translation |
| ENSSSCG00000001502 | RPS18 | 1205.32 | 1150.56 | 0.0671 | 0.4464 | 0.6793 | translation |
| ENSSSCG00000039544 | RPS24 | 1170.68 | 1222.17 | -0.0621 | 0.4762 | 0.7047 | translation |
| ENSSSCG00000035768 | RPS15A | 1440.37 | 1395.94 | 0.0451 | 0.5956 | 0.7892 | regulation of cell cycle, translation |
| ENSSSCG00000012119 | TMSB4 | 1138.57 | 1164.31 | -0.0322 | 0.6229 | 0.8079 | organization of the cytoskeleton |
| ENSSSCG000000022176 |  | 1262.61 | 1418.36 | -0.1678 | 0.6416 | 0.8203 | unknown |
| ENSSSCG000000027358 | RPS17 | 1417.91 | 1448.65 | -0.0309 | 0.6752 | 0.8411 | translation |
| ENSSSCG000000020817 | RPS16 | 1294.51 | 1315.45 | -0.0231 | 0.7698 | 0.8973 | translation |

Differential gene expression from ovaries of Xiang pig gilts with large and small litter size were calculated by applying the cutoff of expression values at FPKM + 0.001, padj-value ≤ 0.05, and log2 fold-change ≥1 or ≤-1. The significance scores were corrected for multiple testing using Benjamini-Hochberg correction. A total of 762 genes differentially expressed between XL and XS samples. Compared with XS samples, 34 genes were up-regulated and 728 genes were down-regulated in XL samples (Supplemental Table S2). Most of the genes with greatest changes in expression were down-regulated genes. The range of log2 fold change values for DEGs was from 5.94 to −5.96. The scatter plots used to demonstrate the up- and down-regulated genes identified in the ovary tissues from XL and XS gilts (Figure 2). Of these genes, 47 were highly upregulated or down-regulated genes with a fold change of more than 4 (log2 > 2) or 30 (log2 < -5) in expression between XL and XS, including *STAR, NR5A1*, *MORN5, TSPAN1, FRK*, *TEKT1, LHCGR, MTHFD2*, *ZSWIM2* (Table 3)*. STAR* (P-adj = 9.5e-03) was the most up-regulated gene in XL group, whereas *ZSWIM2* showed the highest down-regulation (P-adj = 0.01 and logFC = −5.96).

**Figure 2.**
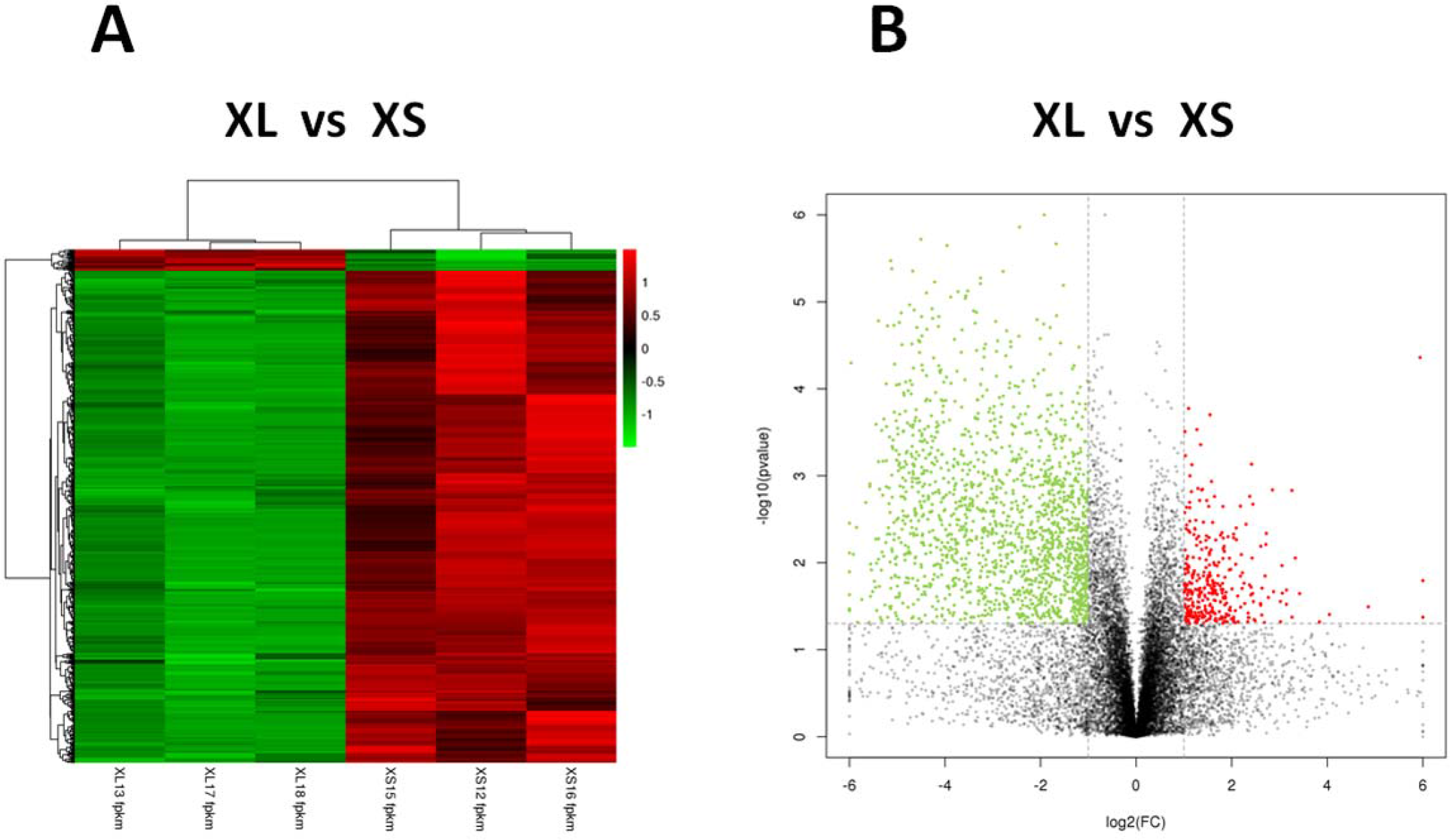
Differential expression of the genes between XL and XS groups. A: Scatter plot of differentially expressed genes (XL vs. XS). Each point in the figures represented one gene. Red points represent upregulated genes; Green points denote downregulated genes. Black points represent genes without significant difference. B: Heatmaps illustrate the most significantly upregulated and downregulated genes between two groups.

**Table 3.** The most significantly upregulated and downregulated genes in ovaries between XL and XS

| Gene ID | Gene | XL FPKM | XS FPKM | logFC | P.Value | adj.P.Val | Function |
| --- | --- | --- | --- | --- | --- | --- | --- |
| ENSSSCG00000004147 | ECT2L | 0.4911 | 17.2465 | -5.1341 | 3.36E-06 | 0.007373 | regulation of Rho protein signal transduction |
| ENSSSCG000000040973 | HYDIN | 0.7709 | 26.672 | -5.1125 | 4.16E-06 | 0.007373 | cilium movement,epithelial cell development |
| ENSSSCG000000002817 | DRC7 | 0.5791 | 24.2991 | -5.3911 | 1.65E-05 | 0.007981 | flagellar motility |
| ENSSSCG000000033394 | VWA3B | 0.1207 | 4.4461 | -5.2026 | 1.89E-05 | 0.007981 | apoptosis pathway |
| ENSSSCG000000002504 | AK7 | 1.5712 | 52.6944 | -5.0676 | 1.86E-05 | 0.007981 | nucleoside triphosphate biosynthetic process |
| ENSSSCG000000034942 | STAR | 101.3265 | 1.6502 | 5.9401 | 4.35797E-05 | 0.009582 | steroid biosynthetic process |
| ENSSSCG000000016030 | ZSWIM2 | 0.1006 | 6.2815 | -5.9638 | 5.04E-05 | 0.010794 | regulation of extrinsic apoptotic signaling |
| ENSSSCG000000005588 | NR5A1 | 25.0387 | 5.3585 | 2.2242 | 0.003041 | 0.01082 | female gonad development |
| ENSSSCG000000010100 | LRRC74B | 0.1608 | 5.3494 | -5.0559 | 5.21E-05 | 0.010921 | unknown |
| ENSSSCG000000017884 | TEKT1 | 1.2328 | 46.2493 | -5.2293 | 8.77E-05 | 0.012834 | flagellated sperm motility |
| ENSSSCG000000009428 | FAM216B | 1.5967 | 51.5777 | -5.0135 | 9.03E-05 | 0.012848 | unknown |
| ENSSSCG000000005529 | MORN5 | 1.4651 | 48.9129 | -5.0611 | 0.000181 | 0.016821 | follicle development |
| ENSSSCG000000003823 | C1orf87 | 0.5735 | 22.2651 | -5.2786 | 0.000222 | 0.017979 | unknown |
| ENSSSCG000000017344 |  | 0.1175 | 3.8177 | -5.0214 | 0.000233 | 0.018189 | unknown |
| ENSSSCG000000030800 |  | 0.0978 | 4.2525 | -5.4412 | 0.000255 | 0.018992 | unknown |
| ENSSSCG000000028471 | DTHD1 | 0.7056 | 26.3466 | -5.2224 | 0.000271 | 0.019146 | signal transduction |
| ENSSSCG000000001594 |  | 0.0673 | 2.6978 | -5.3241 | 0.000301 | 0.019961 | microtubule motor activity |
| ENSSSCG000000008908 | PDCL2 | 0.1407 | 5.0321 | -5.1595 | 0.000356 | 0.020796 | queuosine biosynthetic process |
| ENSSSCG000000015696 | MAP3K19 | 0.2522 | 8.5697 | -5.0864 | 0.000387 | 0.021837 | regulation of mitotic cell cycle |
| ENSSSCG000000020990 | DNAH12 | 0.3121 | 10.7355 | -5.1044 | 0.000458 | 0.023934 | cilium movement |
| ENSSSCG000000007236 | TTLL9 | 0.3516 | 11.9891 | -5.0916 | 0.000487 | 0.024699 | cellular protein modification process |
| ENSSSCG000000039903 | SPATS1 | 0.2478 | 8.0353 | -5.0187 | 0.000511 | 0.025256 | testis development |
| ENSSSCG000000000983 | IL17REL | 0.0978 | 3.6634 | -5.2264 | 0.000669 | 0.028073 | inflammatory responses |
| ENSSSCG000000015741 | CFAP221 | 0.2738 | 8.8253 | -5.0103 | 0.000657 | 0.028073 | cilium movement |
| ENSSSCG000000034242 | HNF4G | 0.0508 | 1.6365 | -5.0079 | 0.000666 | 0.028073 | positive regulation of transcription |
| ENSSSCG000000009809 | MORN3 | 0.4031 | 15.8584 | -5.2982 | 0.000682 | 0.028118 | unknown |
| ENSSSCG000000011078 | ARMC3 | 0.5553 | 23.2969 | -5.3905 | 0.000687 | 0.028154 | signal transduction, development |
| ENSSSCG000000012638 | GRIA3 | 15.8909 | 2.9806 | 2.4145 | 0.000735 | 0.028781 | glutamate-gated ion channel activity |
| ENSSSCG000000010478 | FFAR4 | 0.2932 | 10.1441 | -5.1123 | 0.000982 | 0.031997 | hormone secretion |
| ENSSSCG000000003908 | TSPAN1 | 5.3101 | 252.0776 | -5.5689 | 0.001237 | 0.035091 | cell proliferation |
| ENSSSCG000000003819 | ANGPTL3 | 0.0813 | 3.8734 | -5.5734 | 0.001327 | 0.035857 | angiogenesis,cholesterol metabolic process |
| ENSSSCG000000033462 |  | 0.7444 | 27.9106 | -5.2284 | 0.001334 | 0.035866 | unknown |
| ENSSSCG000000017495 | GRB7 | 0.1653 | 5.9526 | -5.1701 | 0.001377 | 0.036561 | negative regulation of translation |
| ENSSSCG000000032691 | ANKRD66 | 0.5455 | 22.5371 | -5.3683 | 0.001424 | 0.037318 | unknown |
| ENSSSCG000000008421 | LHCGR | 20.6097 | 2.8525 | 2.8529 | 0.001452 | 0.037422 | folliculogenesis and steroidogenesis |
| ENSSSCG000000031023 |  | 143.2986 | 14.9674 | 3.2591 | 0.001471 | 0.037631 | cell adhesion |
| ENSSSCG000000004434 | FRK | 0.0985 | 3.8885 | -5.3015 | 0.001674 | 0.039844 | cell differentiation |
| ENSSSCG000000008289 | MTHFD2 | 38.5153 | 7.4155 | 2.3768 | 0.001724 | 0.040391 | tetrahydrofolate metabolic process |
| ENSSSCG000000003048 | cxcl17 | 1.3079 | 44.9787 | -5.1038 | 0.001762 | 0.040612 | unknown |
| ENSSSCG000000011850 | mucin 4 | 0.4179 | 21.0885 | -5.6571 | 0.002031 | 0.042575 | cell-matrix adhesion |
| ENSSSCG000000007510 | C20orf85 | 4.4515 | 160.3676 | -5.1709 | 0.002089 | 0.043155 | unknown |
| ENSSSCG000000004195 | ARG1 | 1.6843 | 0.3098 | 2.4425 | 0.002122 | 0.043722 | urea cycle and Nitrogen metabolism. |
| ENSSSCG000000010437 | PAPSS2 | 88.1918 | 19.4349 | 2.1819 | 0.002234 | 0.044691 | folliculogenesis, bioavailability of androgen |
| ENSSSCG00000002734 | MARVELD3 | 0.2771 | 9.8266 | -5.1483 | 0.002641 | 0.047701 | regulation of epithelial cell proliferation |
| ENSSSCG00000003733 | KLHL14 | 0.2122 | 6.8489 | -5.0121 | 0.003203 | 0.053298 | protein ubiquitination |
| ENSSSCG000000036438 | GPX3 | 1396.32 | 326.93 | 2.0945 | 0.0039 | 0.0597 | oocyte maturation or embryo quality |
| ENSSSCG000000035945 | FDX1 | 99.9512 | 18.5901 | 2.4266 | 0.008821 | 0.088508 | steroid biosynthetic process |

### Alternative splicing events and differentially spliced genes in XL and XS

We used ASprofile program to detect 12 types of alternative splicing (AS) events in XL and XS data sets. The results were showed in Table S3. In total, 65,238 AS events from 15,047 genes and 66,086 AS events from 15,282 genes were detected in XL and XS data sets respectively. Approximately, 92.8∼93.8% of the total expressed genes underwent alternative splicing in XL and XS samples. The ovary transcriptome between XL and XS showed similar AS patterns (Table 4). Three types of AS events (TSS, TTS and SKIP) were predominant in pig ovary transcriptome, which accounted for more than 80% of AS events in each library.

**Table 4.**
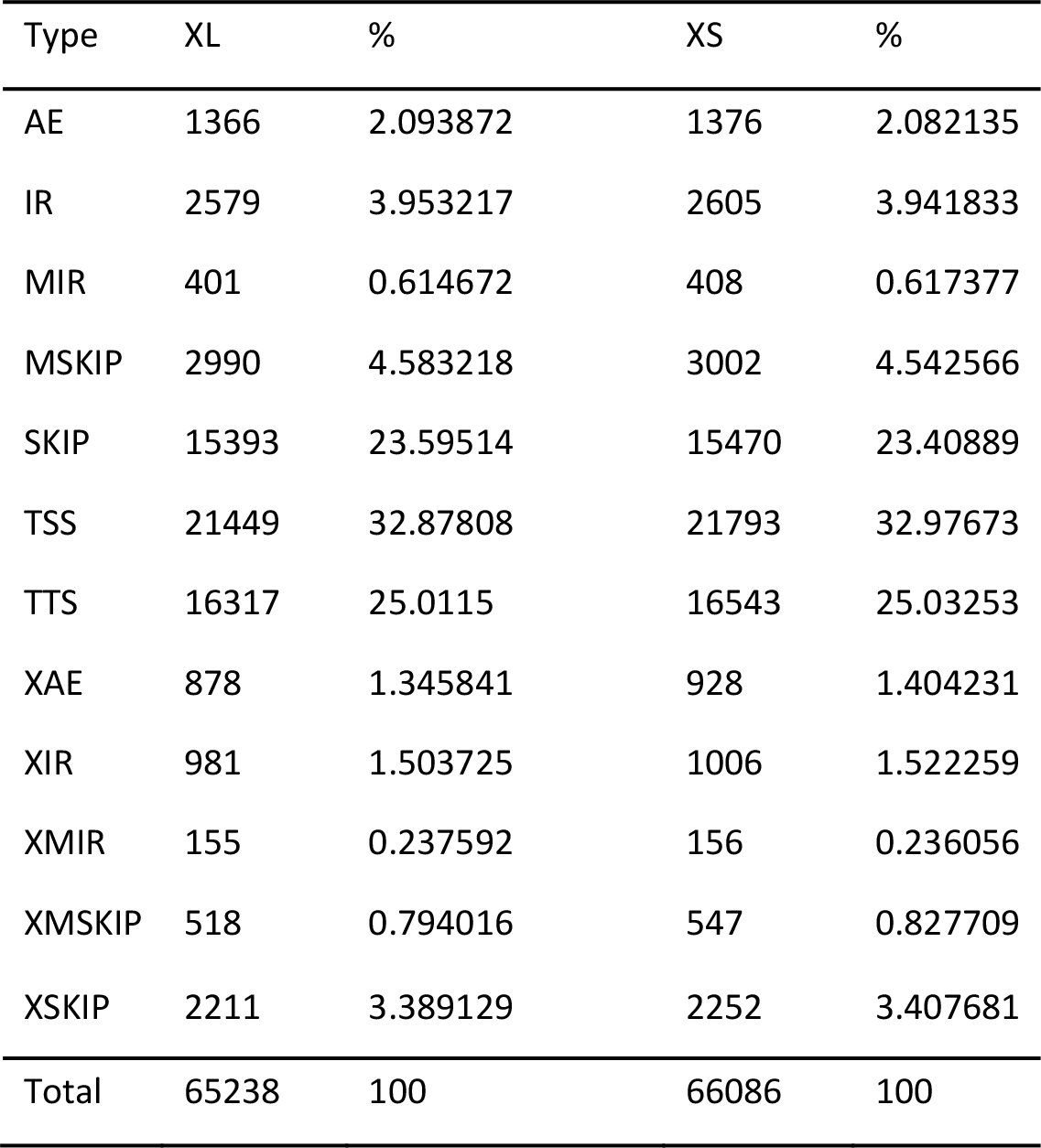
The pattern of alternative splicing events in XL and XS

| Type | XL | % | XS | % |
| --- | --- | --- | --- | --- |
| AE | 1366 | 2.093872 | 1376 | 2.082135 |
| IR | 2579 | 3.953217 | 2605 | 3.941833 |
| MIR | 401 | 0.614672 | 408 | 0.617377 |
| MSKIP | 2990 | 4.583218 | 3002 | 4.542566 |
| SKIP | 15393 | 23.59514 | 15470 | 23.40889 |
| TSS | 21449 | 32.87808 | 21793 | 32.97673 |
| TTS | 16317 | 25.0115 | 16543 | 25.03253 |
| XAE | 878 | 1.345841 | 928 | 1.404231 |
| XIR | 981 | 1.503725 | 1006 | 1.522259 |
| XMIR | 155 | 0.237592 | 156 | 0.236056 |
| XMSKIP | 518 | 0.794016 | 547 | 0.827709 |
| XSKIP | 2211 | 3.389129 | 2252 | 3.407681 |
| Total | 65238 | 100 | 66086 | 100 |

The differentially spliced genes (DSGs) were investigated by using the same cutoff with DEGs. In total, 808 DSGs, which represented 2,039 differentially splicing events, were identified between XL and XS samples (Table S4). Among these DSGs, 602 genes were both differentially regulated on the expression and AS level. Thirty seven of the differentially splicing events from 26 genes (e.g. *TTLL8*, *STAP2*, *ZUFSP*, *LNX1*, *ARNTL*, *SPAG9* etc.) showed large (log2 fold-change >10-fold) differences in AS level between XL and XS (Table S5). Furthermore, we found the DSGs were enriched in several gene families (Table S6), for example, cilia and flagella associated protein family (CFAP), intraflagellar transport family (IFT), kinesin family member (KIF), MORN repeat-containing protein family (MORN), NIMA related kinase family (NEK), solute carrier family member (SLC), spermatogenesis associated protein (SPATA), tektin family (TEKT), transmembrane protein family (TMEM), tubulin tyrosine ligase like protein (TTLL), ubiquitin specific peptidase (USP), zinc finger protein (ZNF) and so on.

### Gene ontology

Enrichment analysis of annotation was performed on the DEGs and DSGs using Gene Ontology (GO) Consortium online (http://geneontology.org/). The DEGs and DSGs were enriched in 120 and 110 GO terms (p < 0.05) (Table S7) based on biological process, molecular function, and cellular component ontologies. Most of the GO terms were in common between DEGs and DSGs. For the biological processes GO terms, “process”, “assembly” and “organization” were the predominant categories. For the GO terms of molecular function, most of DEGs and DSGs were annotated with the term “binding” and “activity”. For the cellular component category, the DEGs and DSGs were mainly enriched in GO terms of the “cilium” and “organelle”.

### Validation

To validate the RNA-seq results, nine genes (*SCARB1*, *LDLR*, *CYP11A1*, *HSD3B1*, *STAR*, *AKR1C1*, *SERPINE2*, *RARRES1*, *LRP8*) were randomly selected for quantitative PCR (qPCR). In the same RNA samples of the RNA-seq experiment, the expression fold changes (2^−ΔΔCt^) of these nine genes were tested by using qRT-PCR. The expression patterns of these nine genes were in agreement with the RNA-seq results (Table 5). This suggested that the results from the RNA-seq experiments were accurate and efficient.

**Table 5.**
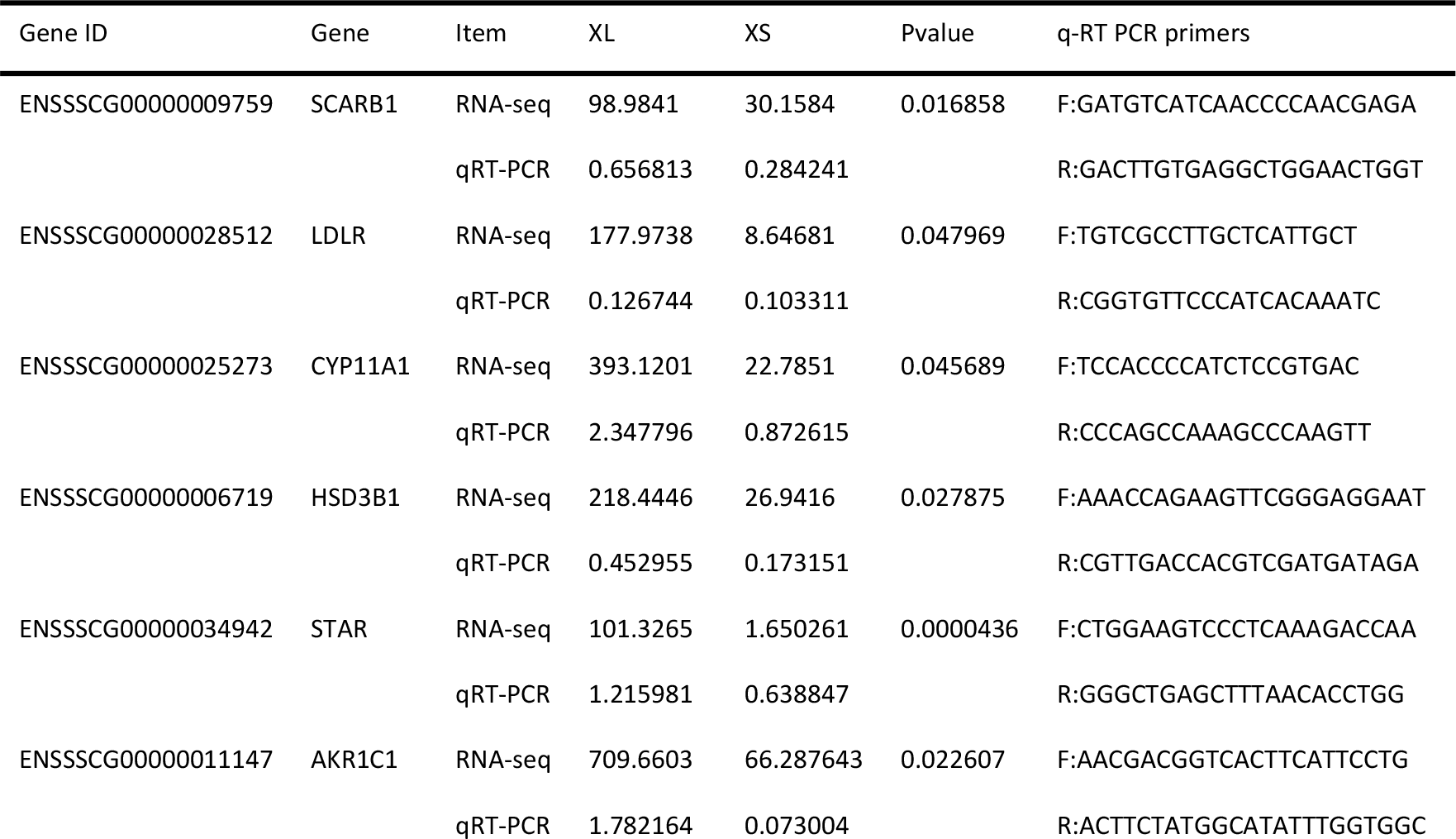
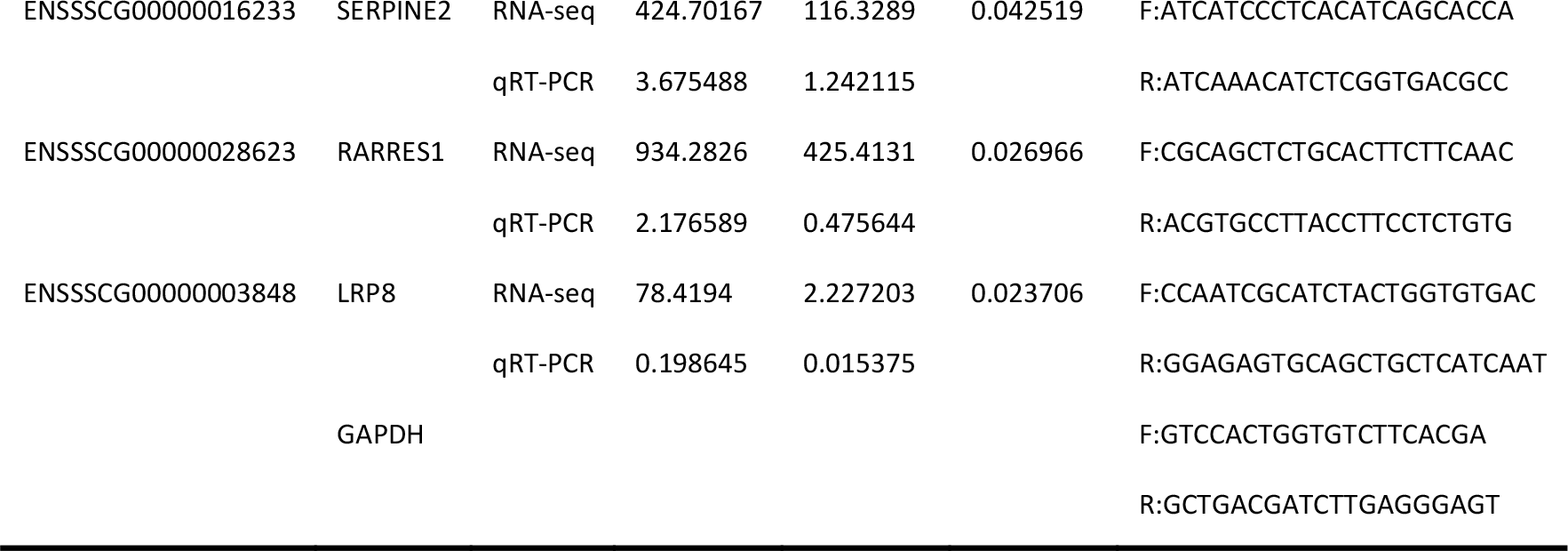
Validation of selected RNA-seq based gene expression by real-time RT-PCR analysis

## Discussion

In this study, we constructed two libraries prepared from Xiang pig gilt samples with extremely large and small litter size. We compared the ovarian transcriptome and alternative splicing of Xiang pig gilts at estrus with large (XL) and small (XS) litter sizes using Solexa sequencing technology. A total of 16,219 and 16,285 expressed genes were detected in ovaries of large and small litter size pigs, respectively. Among of those, 16,050 genes were co-expressed in both libraries. The expression levels of those genes varied greatly, ranging from less than 0.1 FPKM to more than hundreds of thousands of FPKM. The 27 top highly expressed genes (FPKM > 1,000) were identified in the ovaries of large and/or small litter size pig samples (Table 2). Of these top highly expressed genes, *GPX3* and *TIMP1* were upregulated and *OVGP1* and *CLU* genes were downregulated in the high litter size samples compared with small litter size ovary samples. Most of the genes specific for XL or XS showed low or very low expression values. The sequencing frequency of the top expressed genes constituted 11.7∼15.5% of the total expressed values in XL and XS samples. The strong expression of these genes in the ovaries suggests that they are important for ovaries development or specific reproductive processes. For example, more than half of those genes were ribosomal protein genes. Ribosomal proteins (RPs) are ubiquitous RNA binding proteins, the functions of which are not only to participate jointly with ribosome RNA (rRNA) in the synthesis of protein, but also to take part in the regulation of gene transcription and the regulation of cell proliferation and apoptosis through its extra-ribosomal functions [30]. The abnormal expression of RPs often results in serious diseases such as anemia and tumor [31]. *GPX3* gene (glutathione peroxidase 3) and *TIMP1* gene (metalloproteinase inhibitor 1) were proved to be related with oocyte maturation or embryo quality in previous reports [32, 33]. *TPT1* gene is thought to promote cell survival by enhancing the anti-apoptotic response and suppressing proapoptotic activities [34]. The expression of *CLU* gene appears to correlate with cell remodeling or differentiation that occurs during estrus [35].

Previous studies of goat ovaries suggest that some specific differentially expressed genes identified by RNA-Seq are likely to be important for improving litter size [8, 36]. Similar studies on litter size in Yorkshire pigs indicate that some of the top 10 most differentially expressed genes are suggested to be candidate genes involved in specific reproductive processes [19]. In this study, 762 genes were differentially regulated in the ovaries of large and small litter size pigs (Table S2). A number of highly upregulated genes with a fold change of more than 4 (log2 > 2) involved in steroid hormone metabolism and biosynthesis, cell adhesion, organic substance metabolism, Ion transport and signaling transduction [Table 3]. Even though the functions of a lot of genes are still unclear, some of the highly upregulated genes found in our study have been reported to be candidate genes involved in reproduction. For example, the *STAR* gene is required for cholesterol transport into mitochondria to initiate steroidogenesis in the adrenal and gonads [37]. The NR5A1 protein regulates the transcription of key genes involved in sexual development and reproduction, including *STAR*, *CYP17A1*, *CYP11A1*, *LHB*, *AMH*, *CYP19A1*, and *INHA* [38]. NR5A1 is expressed in multiple cell types in the fetal, postnatal, prepubertal, and mature ovary [39]. The inactivation of Nr5a1 specifically in mouse granulosa cells causes infertility associated with hypoplastic ovaries. Nr5a1-/- ovaries have follicles but lack corpora lutea, a finding that indicates impaired ovulation [38]. Follicle-stimulating hormone (FSH) and luteinizing hormone (LH) are gonadotropins (GTHs) that signal through their cognate receptors, FSH receptor (FSHR) and LH/choriogonadotropin receptor (LHCGR), to control major gonadal events in vertebrates, including folliculogenesis and steroidogenesis in the ovary [40]. MTHFD2 is likely responsible for mitochondrial production of both NADH and NADPH in rapidly proliferating cells [41]. *PAPSS2* gene plays an important role in modulating ovarian function and female fertility by control of the bioavailability of ovarian androgen [42]. Adrenodoxin, a versatile ferredoxin, is involved in steroid hormone biosynthesis and vitamin D and bile acid metabolism [43].

The most significantly downregulated genes with a fold change of more than 30 (log2 < -5), is presented in Table 3. A lot of downregulated genes are directly involved in cilium movement, receptor signaling pathway, hormone secretion, protein modification and regulation of apoptotic process. For example, the fyn-related kinase (FRK) is a non-receptor tyrosine kinas, which represses cell proliferation, migration and invasiveness by suppressing epithelial to mesenchymal transition [44]. *TEKT1* is a spermatid specific gene and plays an important role in spermatogenesis [45], but its function in ovary is unclear. TGFβ family control follicle development. MORN5 is involved in TGFβ signaling at all levels. BMP signaling is required for MORN5 expression and reduction of MORN5 derepresses several genes in the BMP and TGFβ signaling pathways [46]. TSPAN1, a new member of the tetraspanin family, could inhibit cell proliferation and migration. TSPAN1 is positive related to PTEN in both clinical specimen and mouse models [47].

In this study, a great number of alternative splicing (AS) events were predicted from the expressed genes in XL and XS data sets. Approximate 92.8% and 93.8% of the total expressed genes underwent alternative splicing in XL and XS samples. This result was similar to the AS rates in human [48]. We found that AS rates in XL samples were lower than that observed in XS samples. Alternative splicing is an important mechanism for regulating gene expression and generating proteome diversity [49]. An increasing number of evidence has revealed that AS influences gene functions in animals and plays a decisive role in the generation of receptor diversity and regulation of growth and development [50]. Many genetic diseases have been closely linked to higher than normal rates of AS [51]. Therefore, we speculated the higher AS rates in Xiang pigs with small litter size might be involved in the progression of lower reproductive capacity. In addition, we identified 808 differentially spliced genes (DSGs)(P-adj < 0.05) and found that 602 genes were differentially regulated on both the expression and AS level. The DEGs exhibited diverse and specific splicing patterns and events between the XL and XS samples. Most of top differentially regulated genes, including *STAR*, *TSPAN1, FRK, PAPSS2, MORN5, MTHFD2, LHCGR* and *TEKT1*, also exhibited highly differentially splicing on AS levels. These results indicate that changes in ovary transcriptome at estrus from Xiang pigs with large and small litter size were differentially regulated on the expression and AS level.

Finally, by analyzing the most highly expressed genes, the top DEGs and top DSGs in the large and small litter size samples, we identified a total of 11 candidate genes (*STAR*, *TSPAN1, FRK, PAPSS2, MORN5, MTHFD2, LHCGR*, *GPX3*, *FDX*1*, NR5A1* and *TEKT1*) relating to porcine fecundity and litter size (Table 6). The functions of *MTHFD2, GPX3, STAR, FDX1, NR5A1, LHCGR* and *PAPSS2* involve in cell proliferation, response to oxidative stress, ovarian steroid hormone synthesis and secretion which may be closely related to the gonad development, oocyte maturation or embryo quality. These genes increasingly expressed in ovary of Xiang pigs with large litter size and promoted ovarian steroidogenesis and oocytes quality. They may be associated with high fecundity in Xiang pigs. The functions of *TSPAN1*, *FRK* and *MORN5* may be related to repress cell proliferation, migration and invasiveness. Reduction of TSPAN1, FRK and MORN5 promotes cell proliferation. They may also be associated with high fecundity in Xiang pigs. *TEKT1* involves cell motility. It is a spermatid specific gene which plays an important role in spermatogenesis, but its function in pig ovary is unknown. In ovary of Xiang pigs with small litter size, the expression of *TEKT1* was found to be higher at estrus stage, which might have specific roles in gonadal physiology. Previous study identified 11 candidate genes for high litter size in Yorkshire pigs using the RNA-Seq method [19]. Interestedly, in our study, two of them (*GPX3, STAR*) were also identified as candidate genes that may be associated with high litter size in Xiang pigs. However, other candidate genes in Yorkshire pigs were completely different from that in Xiang pigs. The results suggest that litter size traits were not only affected by common genes, but regulated by different genes due to genetic differences between pig breeds.

**Table 6.** the candidate genes relating to porcine fecundity and litter size

| Gene ID | Gene symbol | XL FPKM | XS- FPKM | logFC | P.Value | adj.P.Val | function |
| --- | --- | --- | --- | --- | --- | --- | --- |
| ENSSSCG00000003908 | TSPAN1 | 5.3101 | 252.0777 | -5.5689 | 0.001237 | 0.035091 | cell proliferation |
| ENSSSCG00000004434 | FRK | 0.0986 | 3.8886 | -5.3015 | 0.001674 | 0.039844 | cell differentiation |
| ENSSSCG00000005529 | MORN5 | 1.4651 | 48.9129 | -5.0611 | 0.000185 | 0.016821 | follicle development |
| ENSSSCG00000005588 | NR5A1 | 25.0387 | 5.3585 | 2.2242 | 0.003044 | 0.010822 | sexual development and reproduction |
| ENSSSCG00000008289 | MTHFD2 | 38.5153 | 7.4155 | 2.3768 | 0.001724 | 0.040391 | cell proliferation |
| ENSSSCG00000008421 | LHCGR | 20.6097 | 2.8525 | 2.8529 | 0.001452 | 0.037422 | folliculogenesis and steroidogenesis |
| ENSSSCG00000017884 | TEKT1 | 1.2328 | 46.2493 | -5.2293 | 0.000087 | 0.012834 | cell motility |
| ENSSSCG000000034942 | STAR | 101.3265 | 1.6503 | 5.9401 | 0.000043 | 0.009582 | steroidogenesis |
| ENSSSCG00000010437 | PAPSS2 | 88.1918 | 19.4349 | 2.1819 | 0.002234 | 0.044691 | folliculogenesis, bioavailability of androgen |
| ENSSSCG000000035945 | FDX1 | 99.9513 | 18.5901 | 2.4266 | 0.008821 | 0.088508 | synthesis of various steroid hormones |
| ENSSSCG000000036438 | GPX3 | 1396.3267 | 326.9341 | 2.0945 | 0.003981 | 0.059721 | oocyte maturation or embryo quality |

## Conclusion

This work presented a genome-wide view of gene expression and alternative splicing in estrus ovarian between Xiang pig gilts with large and small litter size. The significant changes in the expression of at least 4.7% of all expressed genes were found in estrus ovarian between XL and XS groups. AS rates in XL samples were lower than that observed in XS samples. Most of differentially expressed genes were differentially regulated on both expression and AS level. After analyzing the function of these genes, 11 candidate genes were potentially identified to be related to Xiang pig fecundity and litter size. The significant changes in the expression of the protein-coding genes and the level of alternative splicing in estrus ovarian transcriptome between XL and XS groups probably are the molecular mechanisms of phenotypic variation in litter size.

## Supporting information

new simple supple Tables

## Acknowledgments

This work are funded by the National High Technology Research and Development Program of China (863 Program) [2013AA102503], the National Natural Science Foundation of China (31672390, 31401091), the Guizhou Province “Hundred” Innovative Talents Project [2016-4012], and the Guizhou Agriculture Research program (QKHZC[2017]2585, QKHZC[2017]2587).

## Disclosure Statement

The authors have no Disclosure Statement to declare.

## Cover Letter

The present manuscript focused on the ovary transcriptome and alternative splicing of Xiang pig between groups with large and small litter size. Here are instruction for the asked questions by the honored journal eLIFE editor:

### * How will your work make others in the field think differently and move the field forward?

Litter size is one of the most important reproductive traits in pig breeding, which is affected by multiple genes and the environment. Ovaries are the most important reproductive organs and have a profound impact on the reproduction efficiency. Therefore, genetic differences in the ovaries may contribute to the observed differences in litter size. Although QTLs and candidate genes have been reported to affect the litter size in many pig breeds, however, the studies were only focused on specific species or breeds and the findings cannot elucidate the marked differences of the reproductive traits between Asian and European pig breeds and the variations of litter size between pigs.

The distinctively biological characteristics of Xiang pig are small in body size together with early sexual maturity. A large phenotypic variation in litter size was observed within Xiang pig sows. To analyze changes in gene expression profiles, in the present study, we collected ovary of Xiang pigs having previously a large or small litter size at estrus after weaning and performed a genome wide analysis of transcriptome and alternative splicing events of transcripts using RNA-seq technology. RNA-seq technology produces vastly more sequence data in a cost-effective way and in a much shorter amount of time, allowing a deep characterization of the transcriptome and genome-wide analysis of alternative splicing in a variety of cells and conditions. Our results will give an insight of understanding on the molecular mechanisms of the marked differences of the reproductive traits between Asian and European pig breeds and the variations of litter size between pig breeds.

### * How does your work relate to the current literature on the topic?

On the topic for pig transcriptome and different expression genes, there are several research teams working on by using transcriptionomic techniques. For example, Zhang et al (2015) assessed gene expression differences between the ovaries of Yorkshire pigs with extremely high and low litter sizes using the RNA-Seq method. Chu et al (2017) investigated the mRNA expression profiles of follicular tissue from Large White gilts and Chinese indigenous Mi gilts in estrus using RNA sequencing technology. Both of them focused on the different expression patterns comparison between pig breeds of different pig population with high or low litter size. They got large amount of different expressed genes enriched in gonad development, ovarian steroidogenesis and oocyte maturation processes.

For the topic of alternative splicing events in transcripts, many work used RNA-seq technology to detect AS events in transcripts. For example, Wang et al (2008) analyzed AS event from the human transcriptomes of ten tissues based on deep sequencing of cDNA fragments [48] and the proteome diversity by trypsin [49]. They found that many genetic diseases was closely linked to higher than normal rates of AS [50, 51]. However, there are little reports on the roles of AS in regulation of gene expression and function in pig ovary.

Using the same design to take the ovaries with extremely high and low litter size and RNA-Seq method, we analyzed the expression patterns combined with AS events in transcriptome of Xiang pig ovary. Firstly, we found a large amount different expressed genes (DEG) between ovaries of Xiang pig with large and small litter size. Most of different expressed genes were enriched in the same process in gonad development, ovarian steroidogenesis and oocyte maturation processes, which were similar in Yorkshire, Large White and Mi pig breeds (19, 37). Interestedly, in our study, two of the top DEG (GPX3, STAR) were identified as candidate genes in both of Yorkshire and Xiang pigs. However, other candidate genes were completely different from that in Xiang pigs. The results suggest that litter size traits were not only affected by common genes, but regulated by different genes due to great genetic differences between pig breeds.

Secondly, We found that approximate 92.8% and 93.8% of the total expressed genes underwent alternative splicing in two Xiang pig groups. This result was similar to the AS rates in human [48]. Therefore, we speculated the higher AS rates in Xiang pigs with small litter size might be involved in the progression of lower reproductive capacity. The outstanding finding was that we identified 602 genes were differentially regulated on both the expression and AS level.

Most previous works take the version of Sscrofa10.2 as reference, which released on Aug 2011 and contained too much repeat sequences and gaps. Different version of pig genome reference might affect the results. It is worth to denote that the current version of pig genome sequence, Sscrofa11.1, was used as refer in the present manuscript. The reference sequence in version Sscrofa11.1 released on Jul 2017 was much more accurate and shorter (with 2.4 G base pairs) than that in version Sscrofa10.2 (3.0 G).

### * Who do you consider to be the most relevant audience for this work?

Students, teachers in University and institute; Farmers and researcher working in pig breeding might be interested in reading and referring to our work.

### * Have you made clear in the letter what the work has and has not achieved?

The outstanding findings in this work are following:

1. we detected a large amount of expressed genes and twelve types of alternative splicing (AS) events from Xiang pig ovaries. A total of 762 differentially expressed genes were identified based comparison between XL (Xiang pig group with large litter size) vs XS (Xiang pig group with small litter size) groups.
2. Alternative splicing (AS) rates in XL samples were slightly lower than that observed in XS samples. Most of differentially expressed genes were also significantly regulated on AS level.
3. Eleven candidate genes were identified to be related to Xiang pig fecundity and litter size, which may be closely related to the gonad development, oocyte maturation or embryo quality.

In this study, 762 genes were differentially regulated in the ovaries of large and small litter size pigs. A number of highly upregulated genes with a fold change of more than 4 (log2 > 2) involved in steroid hormone metabolism and biosynthesis, cell adhesion, organic substance metabolism, Ion transport and signaling transduction [Table 3]. The functions of some genes have been demonstrated in human or animals but lots of them are still unclear. For example, the STAR gene is required for cholesterol transport into mitochondria to initiate steroidogenesis in the adrenal and gonads of mouse [37]. The NR5A1 protein regulates the transcription of key genes involved in sexual development and reproduction, including STAR, CYP17A1, CYP11A1, LHB, AMH, CYP19A1, and INHA [38]. The inactivation of Nr5a1 specifically in mouse granulosa cells causes infertility associated with hypoplastic ovaries. These gene may be associated with high fecundity in Xiang pigs. TEKT1 involves in cell motility. It is a spermatid specific gene which plays an important role in spermatogenesis, but its function in pig ovary is unknown.

All of authors agreed to submit the manuscript to the journal, eLIFE, by the platform bioRxiv. And the simplest supplement files has been submitted, in which seven supplement tables contracted to 63 pages from the original 1837 pages.

Prof. Ran

Guizhou University

## References

1 Li PH, Ma X, Zhang YQ, Zhang Q, Huang RH: Progress in the physiological and genetic mechanisms underlying the high prolificacy of the Erhualian pig. Yi Chuan 2017; 39(11):1016–1024.

2 Andersson E, Frössling J, Engblom L, Algers B, Gunnarsson S: Impact of litter size on sow stayability in Swedish commercial piglet producing herds. Acta Vet Scand 2016; 58(1):31. DOI: 10.1186/s13028-016-0213-8.

3 Zak LJ, Gaustad AH, Bolarin A, Broekhuijse MLWJ, Walling GA, Knol EF: Genetic control of complex traits, with a focus on reproduction in pigs. Mol Reprod Dev 2017; 84:1004–1011.

4 Kwon WS, Rahman MS, Ryu DY, Khatun A, Pang MG: Comparison of markers predicting litter size in different pig breeds. Andrology 2017; 5(3):568–577.

5 Lawlor PG, Lynch PB: A review of factors influencing litter size in Irish sows. Ir Vet J 2007; 60(6): 359–366.

6 Varona L, Legarra A, Herring W, Vitezica ZG: Genomic selection models for directional dominance: an example for litter size in pigs. Genet Sel Evol 2018; 50:1 DOI: 10.1186/s12711-018-0374-1.

7 Guo X, Su G, Christensen OF, Janss L, Lund MS: Genome-wide association analyses using a Bayesian approach for litter size and piglet mortality in Danish Landrace and Yorkshire pigs. BMC Genomics 2016; 17:468. DOI: 10.1186/s12864-016-2806-z

8 Ling YH, Xiang H, Li YS, Liu Y, Zhang YH, Zhang ZJ, Ding JP, Zhang XR: Exploring differentially expressed genes in the ovaries of uniparous and multiparous goats using the RNA-Seq (Quantification) method. Gene 2014; 550:148–153.

9 Córdoba S, Balcells I, Castelló A, Ovilo C, Noguera JL, Timoneda O, Sánchez A: Endometrial gene expression profile of pregnant sows with extreme phenotypes for reproductive efficiency. Sci Rep 2015; 5:14416. DOI: 10.1038/srep14416

10 Ferrari M, Lindholm AK, König B: A genetic tool to manipulate litter size. Front Zool 2014; 11:18 DOI: 10.1186/1742-9994-11-18.

11 Lai FN, Zhai HL, Cheng M, Ma JY, Cheng SF, Ge W, Zhang GL, Wang JJ, Zhang RQ, Wang X, Min LJ, Song JZ, Shen W: Whole-genome scanning for the litter size trait associated genes and SNPs under selection in dairy goat (*Capra hircus*). Sci Rep 2016; 6:38096. DOI: 10.1038/srep38096.

12 Sell-Kubiak E, Duijvesteijn N, Lopes MS, Janss LL, Knol EF5, Bijma P, Mulder HA: Genome-wide association study reveals novel loci for litter size and its variability in a Large White pig population. BMC Genomics 2015; 16:1049 DOI: 10.1186/s12864-015-2273-y.

13 Sun X, Mei S, Tao H, Wang G, Su L, Jiang S, Deng C, Xiong Y, Li F: Microarray profiling for differential gene expression in PMSG-hCG stimulated preovulatory ovarian follicles of Chinese Taihu and Large White sows. BMC Genomics 2011; 12:111. DOI: 10.1186/1471-2164-12-111.

14 Chu Q, Zhou B, Xu F, Chen R, Shen C, Liang T, Li Y, Schinckel AP: Genome-wide differential mRNA expression profiles in follicles of two breeds and at two stages of estrus cycle of gilts. Sci Rep 2017; 7: 5052. DOI: 10.1038/s41598-017-04336-x.

15 Fischer D, Laiho A, Gyenesei A, Sironen A: Identification of reproduction-related gene polymorphisms using whole transcriptome sequencing in the Large White pig population. G3 2015; 5:1351–1360.

16 Chen X, Fu J, Wang A: Expression of genes involved in progesterone receptor paracrine signaling and their effect on litter size in pigs. J Anim Sci Biotechnol 2016; 7:31. DOI: 10.1186/s40104-016-0090-z.

17 Kwon SG, Hwang JH, Park DH, Kim TW, Kang DG, Kang KH, Kim IS, Park HC, Na CS, Ha J, Kim CW: Identification of differentially expressed genes associated with litter size in Berkshire pig placenta. PLoS ONE 2016; 11(4): e0153311. DOI: 10.1371/journal.pone.0153311.

18 Muñoz M, Fernández AI, Ovilo C, Muñoz G, Rodriguez C, Fernández A, Alves E, Silió L: Non-additive effects of RBP4, ESR1 and IGF2 polymorphisms on litter size at different parities in a Chinese-European porcine line. Genet Sel Evol 2010; 42:23. DOI: 10.1186/1297-9686-42-23.

19 Zhang X, Huang L, Wu T, Feng Y, Ding Y, Ye P, Yin Z: Transcriptomic analysis of ovaries from pigs with high and low litter Size. PLoS ONE 2015; 10(10): e0139514. DOI:10.1371/journal.pone.0139514.

20 Coster A, Madsen O, Heuven HCM, Dibbits B, Groenen MA, van Arendonk JA, Bovenhuis H: The imprinted gene DIO3 Is a candidate gene for litter size in pigs. PLoS ONE 2012; 7(2): e31825. DOI:10.1371/journal.pone.003182.

21 Wang Y, Ding X, Tan Z, Xing K, Yang T, Wang Y2, Sun D, Wang C: Genome-wide association study for reproductive traits in a Large White pig population. Anim Genet 2018; 49:127–131.

22 Ma X, Li PH, Zhu MX, He LC, Sui SP, Gao S, Su GS, Ding NS, Huang Y, Lu ZQ, Huang XG, Huang RH: Genome-wide association analysis reveals genomic regions on chromosome 13 affecting litter size and candidate genes for uterine horn length in Erhualian pigs. Animal 2018; 14:1–9.

23 Huang J, Liu R, Su L, Xiao Q, Yu M: Transcriptome analysis revealed the embryo-induced gene expression patterns in the endometrium from Meishan and Yorkshire pigs. Int J Mol Sci 2015; 16:22692–22710.

24 Luo ZY, Dai XL, Ran XQ, Cen YX, Niu X, Li S, Huang SH, Wang JF: Identification and profile of microRNAs in Xiang pig testes in four different ages detected by Solexa sequencing. Theriogenology 2018; 117:61–71.

25 Liu C, Ran X, Wang J, Li S, Liu J: Detection of genomic structural variations in Guizhou indigenous pigs and the comparison with other breeds. PLoS ONE 2018; 13(3): e0194282. DOI: 10.1371/journal.pone.0194282.

26 Liu C, Ran X, Yu C, Xu Q, Niu X, Zhao P, Wang J: Whole-genome analysis of structural variations between Xiang pigs with larger litter sizes and those with smaller litter sizes. Genomics 2018; pii: S0888-7543(18)30094-30096.

27 Knauer M, Cassady J, Newcom D, See M: Estimates of variance components for genetic correlations among swine estrus traits. J Anim Sci 2010; 88:2913–2919.

28 Smyth GK: Linear models and empirical bayes methods for assessing differential expression in microarray experiments. Stat Appl Genet Mol Biol 2004; 3:Article3.

29 Livak KJ, Schmittgen TD: Analysis of relative gene expression data using real-time quantitative PCR and the 2(-Delta Delta C(T)) Method. Methods 2001; 25: 402–408.

30 Wang W, Nag S, Zhang X, Wang MH, Wang H, Zhou J, Zhang R: Ribosomal proteins and human diseases: pathogenesis, molecular mechanisms, and therapeutic implications. Med Res Rev 2015; 35(2):225–285.

31 Warner JR, McIntosh KB: How common are extraribosomal functions of ribosomal proteins? Mol Cell 2009; 34(1):3–11.

32 Huang X, Hao C, Shen X, Zhang Y, Liu X: RUNX2, GPX3 and PTX3 gene expression profiling in cumulus cells are reflective oocyte/embryo competence and potentially reliable predictors of embryo developmental competence in PCOS patients. Reprod Biol Endocrinol 2013; 11:109. DOI: 10.1186/1477-7827-11-109.

33 Julie A.W. Stilley JA, Sharpe-Timms KL: TIMP1 contributes to ovarian anomalies in both an MMP-dependent and -independent manner in a rat model. Biol Reprod 2012; 86(2):47. DOI: 10.1095/biolreprod.111.094680.

34 Chen W, Wang H, Tao S, Zheng Y, Wu W, Lian F, Jaramillo M, Fang D, Zhang DD: Tumor protein translationally controlled 1 is a p53 target gene that promotes cell survival. Cell Cycle 2013; 12:2321–2328.

35 Leeb C, Eresheim C, Nimpf J: Clusterin is a ligand for apolipoprotein E receptor 2 (ApoER2) and very low density lipoprotein receptor (VLDLR) and signals via the Reelin-signaling pathway. J Biol Chem 2014; 289(7): 4161–4172.

36 Zhao ZQ, Wang LJ, Sun XW, Zhang JJ, Zhao YJ, Na RS, Zhang JH: Transcriptome analysis of the *Capra hircus* ovary. PLoS One 2015; 10(3): e0121586. DOI:10.1371/journal.pone.0121586.

37 Clark BJ, Hudson EA: StAR protein stability in Y1 and Kin-8 mouse adrenocortical cells. Biology 2015; 4(1):200–215.

38 Lourenço D, Brauner R, Lin L, De Perdigo A, Weryha G, Muresan M, Boudjenah R, Guerra-Junior G, Maciel-Guerra AT, Achermann JC, McElreavey K, Bashamboo A: Mutations in NR5A1 associated with ovarian insufficiency. N Engl J Med 2009; 360(12): 1200–1210.

39 Jeyasuria P, Ikeda Y, Jamin SP, Zhao L, De Rooij DG, Themmen AP, Behringer RR, Parker KL: Cell-specific knockout of steroidogenic factor 1 reveals its essential roles in gonadal function. Mol Endocrinol 2004; 18(7):1610–1619.

40 Liu KC, Ge W: Evidence for gating roles of protein kinase A and protein kinase C in estradiol-induced luteinizing hormone receptor (lhcgr) expression in zebrafish ovarian follicle cells. PLoS One 2013; 8(5): e62524. DOI: 10.1371/journal.pone.0062524

41 Shin M, Momb J, Appling DR: Human mitochondrial MTHFD2 is a dual redox cofactor-specific methylenetetrahydrofolate dehydrogenase/methenyltetrahydrofolate cyclohydrolase. Cancer Metab 2017; 5:11. DOI: 10.1186/s40170-017-0173-0.

42 Oostdijk W, Idkowiak J, Mueller JW, House PJ, Taylor AE, O’Reilly MW, Hughes BA, de Vries MC, Kant SG, Santen GW, Verkerk AJ, Uitterlinden AG, Wit JM, Losekoot M, Arlt W: PAPSS2 Deficiency Causes Androgen Excess via Impaired DHEA Sulfation—In Vitro and in Vivo Studies in a Family Harboring Two Novel PAPSS2 Mutations. J Clin Endocrinol Metab 2015; 100(4):E672–680.

43 Estrada DF: The cytochrome P450 24A1 interaction with adrenodoxin relies on multiple recognition sites that vary among species. J Biol Chem 2018; 293(11):4167–4179.

44 Ogunbolude Y, Dai C, Bagu ET, Goel RK, Miah S, MacAusland-Berg J, Ng CY, Chibbar R, Napper S, Raptis L, Vizeacoumar F, Vizeacoumar F, Bonham K, Lukong KE: FRK inhibits breast cancer cell migration and invasion by suppressing epithelial-mesenchymal transition. Oncotarget 2017; 8 (68): 113034–113065.

45 Larsson M, Norrander J, Gräslund S, Brundell E, Linck R, Ståhl S, Höög C: The spatial and temporal expression of Tekt1, a mouse tektin C homologue, during spermatogenesis suggest that it is involved in the development of the sperm tail basal body and axoneme. Eur J Cell Biol 2000; 79(10):718–725.

46 Cela P, Hampl M, FuK K, Kunova Bosakova M, Krejci P, Richman JM, Buchtova M: MORN5 expression during craniofacial development and its interaction with the BMP and TGFβ pathways. Front Physiol 2016; 7:378. DOI: 10.3389/fphys.2016.00378.

47 Wang HX, Kolesnikova TV, Denison C, Gygi SP, Hemler ME: The C-terminal tail of tetraspanin protein CD9 contributes to its function and molecular organization. J Cell Sci 2011; 124(16): 2702–2710.

48 Wang ET, Sandberg R, Luo S, Khrebtukova I, Zhang L, Mayr C, Kingsmore SF, Schroth GP, Burge CB: Alternative isoform regulation in human tissue transcriptomes. Nature 2008; 456(7221):470–476.

49 Wang X, Codreanu SG, Wen B, Li K, Chambers MC, Liebler DC, Zhang B: Detection of proteome diversity resulted from alternative splicing is limited by trypsin cleavage specificity. Mol Cell Proteomics 2018; 17(3):422–430.

50 Lee Y, Rio DC: Mechanisms and Regulation of Alternative Pre-mRNA Splicing. Annu Rev Biochem 2015; 84:291–323.

51 Chacko E, Ranganathan S: Genome-wide analysis of alternative splicing in cow: implications in bovine as a model for human diseases. BMC Genomics 2009; 10(Suppl 3):S11. DOI: 10.1186/1471-2164-10-S3-S11.

