## Supplementary material for "Changes in Ovary Transcriptome and Alternative Splicing at Estrus from Xiang Pigs with Large and Small Litter Size": new simple supple Tables

#### Table S1 The expression data from the estrus ovaries of Xiang pigs with large and small liter size

| gene id | chr | XL13 | XL17 | XL18 | mean fpkm | XS12 | XS15 | XS16 | mean fpkm |
|---|---|---|---|---|---|---|---|---|---|
| | | gene fpkm | gene fpkm | gene fpkm | | gene fpkm | gene fpkm | gene fpkm | |
| ENSSSCG00000004489 | 1 | 3409.80 | 3670.41 | 3523.14 | 3534.45 | 4162.42 | 3776.40 | 4668.77 | 4202.53 |
| ENSSSCG00000014540 | 2 | 3739.19 | 2812.97 | 2170.87 | 2907.68 | 1798.29 | 1841.20 | 1951.77 | 1863.75 |
| ENSSSCG00000040929 | 11 | 2580.82 | 2672.38 | 2384.82 | 2546.01 | 2658.37 | 3038.57 | 3819.32 | 3172.09 |
| ENSSSCG00000013597 | 2 | 2174.58 | 2078.48 | 1976.63 | 2076.56 | 2157.35 | 2387.24 | 2420.89 | 2321.83 |
| ENSSSCG00000038507 | 1 | 2049.98 | 2035.29 | 2128.93 | 2071.40 | 2030.75 | 2332.14 | 2413.17 | 2258.69 |
| ENSSSCG00000037674 | 1 | 1526.13 | 1585.29 | 1521.23 | 1544.22 | 1788.34 | 1666.58 | 2110.65 | 1855.19 |
| ENSSSCG00000012277 | X | 462.37 | 2635.12 | 1399.24 | 1498.91 | 84.34 | 114.06 | 507.60 | 235.33 |
| ENSSSCG00000035768 | 3 | 1448.41 | 1428.00 | 1444.70 | 1440.37 | 1237.93 | 1524.57 | 1425.33 | 1395.94 |
| ENSSSCG00000008245 | 3 | 1488.40 | 1250.64 | 1563.92 | 1434.32 | 615.60 | 1063.48 | 1460.11 | 1046.40 |
| ENSSSCG00000011033 | 10 | 1521.15 | 1122.31 | 1648.42 | 1430.63 | 218.41 | 1063.61 | 997.17 | 759.73 |
| ENSSSCG00000027358 | 7 | 1538.37 | 1328.93 | 1386.43 | 1417.91 | 1460.56 | 1511.68 | 1373.73 | 1448.66 |
| ENSSSCG00000028355 | 12 | 1620.77 | 1406.77 | 1209.07 | 1412.20 | 771.41 | 707.29 | 974.54 | 817.75 |
| ENSSSCG00000036438 | 16 | 1478.32 | 1110.57 | 1600.09 | 1396.33 | 104.47 | 472.37 | 403.96 | 326.93 |
| ENSSSCG00000020817 | 6 | 1289.52 | 1282.72 | 1311.31 | 1294.52 | 1177.48 | 1346.57 | 1422.32 | 1315.46 |
| ENSSSCG00000022176 | 5 | 1317.73 | 1104.95 | 1365.15 | 1262.61 | 2056.54 | 963.10 | 1235.47 | 1418.37 |
| ENSSSCG00000001502 | 7 | 1254.52 | 1177.03 | 1184.43 | 1205.33 | 1030.80 | 1159.49 | 1261.39 | 1150.56 |
| ENSSSCG00000015326 | 9 | 934.27 | 1244.69 | 1398.88 | 1192.61 | 230.02 | 637.98 | 1036.83 | 634.94 |
| ENSSSCG00000036014 | 14 | 1264.67 | 1143.84 | 1165.44 | 1191.32 | 1135.32 | 1296.31 | 1665.61 | 1365.75 |
| ENSSSCG00000039544 | 14 | 1133.11 | 1101.28 | 1277.65 | 1170.68 | 1149.24 | 1207.25 | 1310.04 | 1222.18 |
| ENSSSCG00000012119 | X | 1169.50 | 1133.37 | 1112.84 | 1138.57 | 1112.07 | 1264.54 | 1116.29 | 1164.30 |
| ENSSSCG00000035904 | 1 | 1173.41 | 1107.58 | 1082.36 | 1121.12 | 1168.00 | 1207.61 | 1445.50 | 1273.70 |
| ENSSSCG00000035997 | X | 1077.58 | 1113.11 | 1085.75 | 1092.15 | 1235.98 | 1326.30 | 1614.73 | 1392.34 |
| ENSSSCG00000027573 | 13 | 1071.38 | 943.70 | 944.94 | 986.67 | 830.78 | 911.59 | 1011.49 | 917.95 |
| ENSSSCG00000014855 | 9 | 1038.04 | 902.64 | 953.93 | 964.87 | 962.09 | 920.98 | 997.18 | 960.08 |
| ENSSSCG00000024260 | 6 | 985.02 | 948.54 | 950.25 | 961.27 | 1030.09 | 1157.37 | 997.25 | 1061.57 |
| ENSSSCG00000003042 | 6 | 936.21 | 977.68 | 958.22 | 957.37 | 948.80 | 982.45 | 1139.90 | 1023.72 |
| ENSSSCG00000029830 | 4 | 1005.17 | 916.50 | 923.66 | 948.45 | 902.00 | 871.48 | 889.01 | 887.50 |
| ENSSSCG00000021825 | 17 | 908.65 | 1005.80 | 916.35 | 943.60 | 707.08 | 1096.84 | 1163.09 | 989.00 |
| ENSSSCG00000036135 | 12 | 822.25 | 931.29 | 1060.18 | 937.91 | 148.36 | 540.05 | 938.28 | 542.23 |
| ENSSSCG00000028623 | 13 | 1648.66 | 394.88 | 759.31 | 934.28 | 2.68 | 801.77 | 471.79 | 425.41 |
| ENSSSCG00000017082 | 16 | 832.90 | 975.82 | 985.40 | 931.37 | 224.65 | 647.36 | 915.66 | 595.89 |
| ENSSSCG00000003930 | 6 | 939.47 | 885.00 | 948.56 | 924.34 | 876.86 | 1000.86 | 1158.65 | 1012.12 |
| ENSSSCG00000033169 | 2 | 913.01 | 909.26 | 915.28 | 912.52 | 708.63 | 833.20 | 960.21 | 834.01 |
| ENSSSCG00000013381 | 2 | 1010.33 | 866.67 | 857.02 | 911.34 | 810.69 | 895.48 | 858.13 | 854.77 |
| ENSSSCG00000016034 | 15 | 513.46 | 1066.35 | 1043.63 | 874.48 | 330.45 | 821.45 | 1373.46 | 841.79 |
| ENSSSCG00000011952 | 13 | 984.17 | 840.15 | 794.61 | 872.98 | 912.27 | 902.76 | 838.28 | 884.43 |
| ENSSSCG00000007585 | 3 | 869.40 | 780.45 | 839.68 | 829.84 | 795.02 | 519.04 | 628.04 | 647.37 |
| ENSSSCG00000004578 | 1 | 334.02 | 898.68 | 1241.40 | 824.70 | 359.10 | 296.97 | 1011.98 | 556.02 |
| ENSSSCG00000003153 | 6 | 596.39 | 1142.85 | 722.85 | 820.70 | 488.19 | 1013.89 | 534.68 | 678.92 |
| ENSSSCG00000008913 | 8 | 901.74 | 758.42 | 766.34 | 808.83 | 317.06 | 500.54 | 592.97 | 470.19 |
| ENSSSCG00000009019 | 8 | 821.40 | 780.52 | 796.12 | 799.35 | 948.33 | 1024.18 | 1009.06 | 993.86 |
| ENSSSCG00000040273 | 13 | 803.59 | 772.55 | 784.38 | 786.84 | 800.28 | 927.41 | 976.97 | 901.55 |
| ENSSSCG00000001543 | 7 | 818.01 | 737.57 | 780.69 | 778.76 | 727.16 | 785.58 | 840.08 | 784.27 |
| ENSSSCG00000008170 | 3 | 830.62 | 757.87 | 719.06 | 769.18 | 791.33 | 812.60 | 759.50 | 787.81 |
| ENSSSCG00000030849 | X | 785.85 | 749.04 | 744.33 | 759.74 | 672.05 | 759.63 | 782.54 | 738.07 |
| ENSSSCG00000017509 | 12 | 815.22 | 721.99 | 723.55 | 753.58 | 800.04 | 785.88 | 918.23 | 834.72 |
| ENSSSCG00000014133 | 2 | 746.51 | 754.66 | 755.83 | 752.33 | 527.97 | 713.25 | 770.79 | 670.67 |
| ENSSSCG00000012405 | X | 764.94 | 768.41 | 701.57 | 744.97 | 708.92 | 691.52 | 736.66 | 712.37 |
| ENSSSCG00000004177 | 1 | 664.04 | 762.00 | 790.93 | 738.99 | 670.21 | 802.24 | 849.13 | 773.86 |
| ENSSSCG00000011147 | 10 | 1111.77 | 578.11 | 439.10 | 709.66 | 6.03 | 99.31 | 93.52 | 66.29 |
| ENSSSCG00000029724 | 2 | 754.86 | 654.36 | 667.27 | 692.16 | 543.04 | 541.50 | 639.75 | 574.76 |
| ENSSSCG00000035007 | 1 | 730.49 | 642.50 | 676.33 | 683.11 | 673.13 | 778.70 | 822.86 | 758.23 |

| | | | | | | | | | |
|---|---|---|---|---|---|---|---|---|---|
| ENSSSCG00000013064 | 2 | 717.92 | 639.45 | 623.07 | 660.15 | 643.66 | 645.75 | 805.55 | 698.32 |
| ENSSSCG00000017239 | 12 | 673.64 | 655.79 | 622.03 | 650.49 | 603.05 | 677.45 | 662.84 | 647.78 |
| ENSSSCG00000039460 | 6 | 668.91 | 642.84 | 630.97 | 647.57 | 509.37 | 553.91 | 699.67 | 587.65 |
| ENSSSCG00000029785 | 6 | 684.56 | 523.26 | 709.27 | 639.03 | 752.02 | 489.05 | 592.09 | 611.05 |
| ENSSSCG00000004945 | 1 | 681.33 | 576.01 | 602.11 | 619.81 | 627.94 | 624.84 | 635.02 | 629.27 |
| ENSSSCG00000015140 | 9 | 685.01 | 840.61 | 318.98 | 614.87 | 375.58 | 341.55 | 386.03 | 367.72 |
| ENSSSCG00000014924 | 9 | 160.30 | 1096.73 | 558.49 | 605.17 | 70.45 | 115.34 | 176.03 | 120.61 |
| ENSSSCG00000033697 | 3 | 596.76 | 589.25 | 608.16 | 598.05 | 606.90 | 586.99 | 725.50 | 639.79 |
| ENSSSCG00000025928 | 6 | 636.54 | 573.85 | 575.97 | 595.45 | 507.39 | 542.21 | 601.45 | 550.35 |
| ENSSSCG00000025675 | 2 | 559.59 | 588.28 | 628.25 | 592.04 | 598.40 | 583.38 | 748.88 | 643.55 |
| ENSSSCG00000024974 | 6 | 581.21 | 553.78 | 584.41 | 573.13 | 453.71 | 490.49 | 552.93 | 499.04 |
| ENSSSCG00000000915 | 5 | 421.48 | 600.25 | 661.12 | 560.95 | 379.66 | 647.81 | 576.86 | 534.78 |
| ENSSSCG00000032111 | 4 | 607.04 | 500.59 | 563.92 | 557.18 | 570.13 | 654.15 | 462.55 | 562.28 |
| ENSSSCG00000005169 | 1 | 574.45 | 542.70 | 530.75 | 549.30 | 449.89 | 549.06 | 480.69 | 493.21 |
| ENSSSCG00000034617 | 3 | 543.65 | 536.13 | 508.93 | 529.57 | 624.63 | 665.63 | 708.14 | 666.14 |
| ENSSSCG00000037274 | 15 | 718.32 | 513.94 | 344.76 | 525.67 | 752.06 | 600.76 | 667.05 | 673.29 |
| ENSSSCG00000031370 | 2 | 623.37 | 485.71 | 458.84 | 522.64 | 374.60 | 464.46 | 454.36 | 431.14 |
| ENSSSCG00000000694 | 5 | 450.40 | 470.16 | 626.09 | 515.55 | 669.82 | 418.48 | 607.65 | 565.32 |
| ENSSSCG00000023666 | 14 | 393.86 | 573.16 | 569.90 | 512.31 | 115.38 | 286.57 | 259.31 | 220.42 |
| ENSSSCG00000010281 | 14 | 403.61 | 606.70 | 525.72 | 512.01 | 272.27 | 390.75 | 388.66 | 350.56 |
| ENSSSCG00000031838 | 4 | 507.72 | 525.00 | 499.50 | 510.74 | 539.01 | 619.77 | 743.51 | 634.09 |
| ENSSSCG00000025698 | 3 | 352.35 | 1023.89 | 135.29 | 503.84 | 14.17 | 19.89 | 62.52 | 32.19 |
| ENSSSCG00000006081 | 4 | 503.31 | 500.91 | 503.58 | 502.60 | 522.31 | 524.72 | 560.89 | 535.97 |
| ENSSSCG00000035520 | X | 595.05 | 425.25 | 481.95 | 500.75 | 558.58 | 480.59 | 448.19 | 495.79 |
| ENSSSCG00000021208 | 16 | 372.35 | 661.70 | 460.62 | 498.22 | 177.12 | 550.94 | 309.46 | 345.84 |
| ENSSSCG00000003165 | 6 | 516.54 | 481.00 | 442.19 | 479.91 | 439.12 | 496.29 | 527.05 | 487.49 |
| ENSSSCG00000031088 | 3 | 515.05 | 449.41 | 453.53 | 472.66 | 416.71 | 484.27 | 512.13 | 471.04 |
| ENSSSCG00000011272 | 13 | 501.28 | 465.39 | 443.56 | 470.07 | 585.00 | 580.47 | 595.30 | 586.92 |
| ENSSSCG00000009403 | 11 | 376.16 | 553.82 | 480.00 | 470.00 | 327.47 | 524.02 | 339.69 | 397.06 |
| ENSSSCG00000012173 | X | 341.51 | 571.79 | 466.46 | 459.92 | 396.01 | 532.80 | 663.58 | 530.80 |
| ENSSSCG00000011831 | 13 | 400.38 | 586.83 | 384.43 | 457.21 | 17.23 | 269.95 | 197.77 | 161.65 |
| ENSSSCG00000035811 | 13 | 460.67 | 455.44 | 453.70 | 456.60 | 456.26 | 484.34 | 471.60 | 470.74 |
| ENSSSCG00000000361 | 5 | 405.79 | 521.74 | 437.61 | 455.05 | 228.74 | 276.70 | 355.64 | 287.03 |
| ENSSSCG00000004970 | 1 | 420.94 | 445.90 | 498.19 | 455.01 | 556.53 | 522.88 | 715.71 | 598.37 |
| ENSSSCG00000013887 | 2 | 702.67 | 128.85 | 529.23 | 453.58 | 0.19 | 38.83 | 56.25 | 31.76 |
| ENSSSCG00000000374 | 5 | 431.08 | 442.88 | 480.20 | 451.39 | 401.35 | 496.29 | 651.07 | 516.24 |
| ENSSSCG00000023971 | 10 | 522.21 | 393.66 | 411.55 | 442.48 | 430.46 | 536.26 | 566.04 | 510.92 |
| ENSSSCG00000024263 | 1 | 433.84 | 401.79 | 485.95 | 440.52 | 333.83 | 346.33 | 509.66 | 396.61 |
| ENSSSCG00000012974 | 2 | 425.49 | 428.67 | 460.70 | 438.29 | 295.61 | 252.84 | 358.62 | 302.35 |
| ENSSSCG00000002669 | 6 | 93.94 | 823.44 | 394.19 | 437.19 | 23.61 | 75.14 | 72.42 | 57.06 |
| ENSSSCG00000039795 | 5 | 382.50 | 369.29 | 536.30 | 429.36 | 677.40 | 275.76 | 413.64 | 455.60 |
| ENSSSCG00000016233 | 15 | 619.45 | 255.66 | 399.00 | 424.70 | 2.58 | 179.71 | 166.70 | 116.33 |
| ENSSSCG00000036296 | 9 | 439.02 | 384.07 | 444.37 | 422.49 | 427.08 | 442.92 | 472.06 | 447.35 |
| ENSSSCG00000030300 | 6 | 412.62 | 598.20 | 252.45 | 421.09 | 235.37 | 231.02 | 561.41 | 342.60 |
| ENSSSCG00000039046 | X | 444.11 | 378.03 | 409.84 | 410.66 | 366.20 | 416.11 | 353.67 | 378.66 |
| ENSSSCG00000020771 | 15 | 193.93 | 432.96 | 580.35 | 402.42 | 0.39 | 28.96 | 209.95 | 79.77 |
| ENSSSCG00000028850 | 12 | 401.54 | 366.17 | 433.19 | 400.30 | 431.69 | 449.99 | 389.55 | 423.74 |
| ENSSSCG00000025273 | 7 | 754.97 | 265.80 | 158.58 | 393.12 | 2.08 | 35.49 | 30.78 | 22.79 |
| ENSSSCG00000007524 | 17 | 394.06 | 363.88 | 421.24 | 393.06 | 382.56 | 384.64 | 428.26 | 398.48 |
| ENSSSCG00000032556 | 3 | 463.62 | 366.96 | 346.52 | 392.36 | 201.85 | 203.30 | 281.25 | 228.80 |
| ENSSSCG00000009303 | 11 | 436.21 | 364.65 | 369.82 | 390.23 | 291.40 | 368.37 | 360.14 | 339.97 |
| ENSSSCG00000039468 | 9 | 496.44 | 328.55 | 320.45 | 381.81 | 138.34 | 166.54 | 231.51 | 178.79 |
| ENSSSCG00000007520 | 17 | 389.56 | 421.96 | 326.76 | 379.43 | 267.69 | 288.43 | 263.05 | 273.06 |
| ENSSSCG00000037030 | 13 | 376.61 | 382.65 | 373.11 | 377.46 | 373.20 | 360.21 | 369.78 | 367.73 |
| ENSSSCG00000013460 | 2 | 382.08 | 380.47 | 363.84 | 375.47 | 301.50 | 319.14 | 422.92 | 347.85 |
| ENSSSCG00000025507 | 12 | 370.12 | 364.01 | 389.01 | 374.38 | 315.52 | 374.18 | 368.78 | 352.83 |

| | | | | | | | | | |
|---|---|---|---|---|---|---|---|---|---|
| ENSSSCG00000024669 | 6 | 355.99 | 369.19 | 396.03 | 373.74 | 183.74 | 266.31 | 181.76 | 210.60 |
| ENSSSCG00000006899 | 4 | 381.17 | 371.45 | 361.50 | 371.37 | 427.61 | 407.92 | 376.73 | 404.09 |
| ENSSSCG00000000886 | 5 | 333.43 | 453.52 | 325.60 | 370.85 | 176.46 | 237.10 | 283.67 | 232.41 |
| ENSSSCG00000036350 | 18 | 424.94 | 348.66 | 338.86 | 370.82 | 357.27 | 316.32 | 229.39 | 300.99 |
| ENSSSCG00000033326 | 16 | 357.17 | 375.76 | 369.10 | 367.34 | 399.00 | 418.13 | 410.53 | 409.22 |
| ENSSSCG00000037567 | 3 | 318.80 | 439.67 | 336.28 | 364.92 | 218.85 | 141.18 | 217.89 | 192.64 |
| ENSSSCG00000005453 | 1 | 519.28 | 309.68 | 264.10 | 364.35 | 142.81 | 324.79 | 221.46 | 229.69 |
| ENSSSCG00000003555 | 6 | 396.12 | 332.90 | 360.57 | 363.19 | 489.94 | 395.60 | 439.42 | 441.65 |
| ENSSSCG00000017936 | 12 | 336.92 | 388.56 | 363.23 | 362.90 | 384.37 | 442.88 | 485.28 | 437.51 |
| ENSSSCG00000011803 | 13 | 350.97 | 340.71 | 387.54 | 359.74 | 480.89 | 408.12 | 419.87 | 436.29 |
| ENSSSCG00000010447 | 14 | 447.70 | 260.06 | 356.28 | 354.68 | 811.19 | 164.65 | 186.64 | 387.49 |
| ENSSSCG00000013405 | 2 | 318.39 | 373.77 | 370.92 | 354.36 | 228.74 | 241.04 | 272.63 | 247.47 |
| ENSSSCG00000025192 | 8 | 364.05 | 346.84 | 345.21 | 352.04 | 389.59 | 414.43 | 341.22 | 381.75 |
| ENSSSCG00000039426 | 13 | 216.74 | 426.33 | 396.62 | 346.56 | 125.74 | 158.82 | 217.59 | 167.38 |
| ENSSSCG00000012842 | 2 | 354.73 | 336.67 | 327.62 | 339.67 | 298.71 | 344.36 | 410.33 | 351.13 |
| ENSSSCG00000017430 | 12 | 343.22 | 457.06 | 215.22 | 338.50 | 218.23 | 185.13 | 217.75 | 207.04 |
| ENSSSCG00000017472 | 12 | 277.56 | 578.76 | 155.86 | 337.39 | 49.29 | 65.59 | 82.86 | 65.91 |
| ENSSSCG00000022343 | 6 | 376.23 | 226.88 | 400.04 | 334.38 | 112.16 | 213.64 | 257.45 | 194.42 |
| ENSSSCG00000004002 | 6 | 361.86 | 325.80 | 313.01 | 333.55 | 340.03 | 334.53 | 350.15 | 341.57 |
| ENSSSCG00000032092 | 12 | 356.24 | 358.45 | 251.79 | 322.16 | 282.28 | 277.86 | 243.73 | 267.96 |
| ENSSSCG00000013889 | 2 | 368.55 | 298.58 | 297.10 | 321.41 | 277.15 | 291.45 | 347.55 | 305.38 |
| ENSSSCG00000020830 | 5 | 345.01 | 292.53 | 321.31 | 319.62 | 446.17 | 420.47 | 271.63 | 379.42 |
| ENSSSCG00000009097 | 8 | 387.21 | 280.68 | 288.68 | 318.86 | 150.96 | 220.79 | 287.69 | 219.81 |
| ENSSSCG00000028485 | 6 | 339.29 | 289.01 | 320.67 | 316.32 | 221.47 | 236.90 | 246.05 | 234.81 |
| ENSSSCG00000006083 | 4 | 400.27 | 271.88 | 264.06 | 312.07 | 144.23 | 259.09 | 224.25 | 209.19 |
| ENSSSCG00000001528 | 7 | 294.79 | 310.56 | 318.48 | 307.94 | 261.55 | 314.39 | 372.93 | 316.29 |
| ENSSSCG00000037236 | 13 | 300.12 | 295.64 | 315.34 | 303.70 | 328.80 | 337.09 | 399.81 | 355.23 |
| ENSSSCG00000006063 | 4 | 295.15 | 309.94 | 296.10 | 300.40 | 201.72 | 246.78 | 231.63 | 226.71 |
| ENSSSCG00000006501 | 4 | 299.98 | 312.75 | 286.75 | 299.83 | 284.95 | 270.31 | 398.77 | 318.01 |
| ENSSSCG00000000190 | 5 | 416.62 | 276.00 | 202.65 | 298.42 | 312.52 | 262.27 | 265.03 | 279.94 |
| ENSSSCG00000003088 | 6 | 186.45 | 322.27 | 381.56 | 296.76 | 121.98 | 241.82 | 152.34 | 172.04 |
| ENSSSCG00000004700 | 1 | 315.49 | 271.75 | 293.39 | 293.54 | 206.94 | 200.77 | 379.12 | 262.28 |
| ENSSSCG00000016078 | 15 | 360.91 | 219.82 | 264.59 | 281.77 | 248.81 | 181.86 | 238.31 | 222.99 |
| ENSSSCG00000008602 | 3 | 220.75 | 328.68 | 283.70 | 277.71 | 254.32 | 235.65 | 261.34 | 250.44 |
| ENSSSCG00000005688 | 1 | 68.44 | 363.10 | 400.97 | 277.50 | 1.49 | 24.92 | 90.15 | 38.85 |
| ENSSSCG00000039962 | 3 | 242.77 | 343.94 | 244.87 | 277.19 | 155.70 | 99.73 | 152.84 | 136.09 |
| ENSSSCG00000040470 | 4 | 269.90 | 282.14 | 278.10 | 276.71 | 334.08 | 311.94 | 401.15 | 349.06 |
| ENSSSCG00000017164 | 12 | 189.56 | 382.23 | 241.71 | 271.17 | 117.68 | 152.53 | 111.27 | 127.16 |
| ENSSSCG00000004875 | 1 | 98.38 | 229.99 | 477.02 | 268.46 | 69.33 | 64.71 | 134.56 | 89.53 |
| ENSSSCG00000010400 | 14 | 338.11 | 111.13 | 338.83 | 262.69 | 15.60 | 44.14 | 95.30 | 51.68 |
| ENSSSCG00000010627 | 14 | 135.73 | 338.09 | 302.88 | 258.90 | 70.27 | 106.63 | 94.55 | 90.48 |
| ENSSSCG00000008433 | 3 | 254.38 | 269.36 | 241.40 | 255.05 | 364.87 | 345.08 | 300.96 | 336.97 |
| ENSSSCG00000038880 | 12 | 263.91 | 234.85 | 266.32 | 255.02 | 271.15 | 258.99 | 267.95 | 266.03 |
| ENSSSCG00000009668 | 14 | 270.29 | 324.78 | 166.58 | 253.88 | 1500.03 | 574.72 | 1302.24 | 1125.66 |
| ENSSSCG00000013746 | 2 | 295.34 | 262.68 | 199.45 | 252.49 | 186.45 | 157.62 | 307.28 | 217.12 |
| ENSSSCG00000039130 | 14 | 248.17 | 234.66 | 241.65 | 241.50 | 267.93 | 264.69 | 220.01 | 250.88 |
| ENSSSCG00000009216 | 8 | 27.47 | 273.73 | 419.47 | 240.22 | 316.09 | 390.02 | 1935.68 | 880.59 |
| ENSSSCG00000013849 | 2 | 223.20 | 232.09 | 257.01 | 237.43 | 73.22 | 107.96 | 137.99 | 106.39 |
| ENSSSCG00000000274 | 5 | 230.59 | 239.52 | 235.71 | 235.27 | 242.36 | 247.41 | 296.55 | 262.11 |
| ENSSSCG00000012784 | X | 226.12 | 234.24 | 243.05 | 234.47 | 189.36 | 244.75 | 438.26 | 290.79 |
| ENSSSCG00000016859 | 16 | 143.13 | 210.34 | 341.67 | 231.72 | 6.80 | 104.53 | 80.50 | 63.94 |
| ENSSSCG00000001930 | 7 | 300.33 | 232.37 | 152.51 | 228.40 | 109.63 | 159.84 | 188.06 | 152.51 |
| ENSSSCG00000001701 | 7 | 255.04 | 216.33 | 208.62 | 226.66 | 258.99 | 182.57 | 215.69 | 219.08 |
| ENSSSCG00000004241 | 1 | 388.58 | 197.09 | 84.15 | 223.27 | 61.94 | 130.70 | 78.22 | 90.29 |
| ENSSSCG00000006719 | 4 | 163.84 | 345.33 | 146.16 | 218.44 | 0.43 | 25.50 | 54.90 | 26.94 |
| ENSSSCG00000039148 | 12 | 208.11 | 195.95 | 250.23 | 218.09 | 145.53 | 170.86 | 260.71 | 192.37 |

| | | | | | | | | | |
|---|---|---|---|---|---|---|---|---|---|
| ENSSSCG00000017202 | 12 | 240.21 | 238.23 | 167.70 | 215.38 | 282.97 | 221.46 | 241.21 | 248.55 |
| ENSSSCG00000014569 | 9 | 235.84 | 211.24 | 196.38 | 214.49 | 232.72 | 220.53 | 237.09 | 230.12 |
| ENSSSCG00000038549 | 3 | 160.52 | 367.26 | 109.46 | 212.41 | 50.29 | 68.89 | 50.14 | 56.44 |
| ENSSSCG00000016737 | 18 | 256.82 | 190.40 | 184.77 | 210.66 | 167.50 | 129.09 | 189.85 | 162.15 |
| ENSSSCG00000004789 | 1 | 306.04 | 161.63 | 162.61 | 210.09 | 33.30 | 91.08 | 61.30 | 61.89 |
| ENSSSCG00000030921 | 9 | 298.04 | 172.36 | 158.85 | 209.75 | 23.64 | 64.04 | 125.78 | 71.15 |
| ENSSSCG00000000849 | 5 | 276.06 | 190.72 | 161.07 | 209.28 | 189.68 | 135.78 | 202.24 | 175.90 |
| ENSSSCG00000013260 | 2 | 228.31 | 233.70 | 161.90 | 207.97 | 376.87 | 350.45 | 575.45 | 434.26 |
| ENSSSCG00000029652 | 5 | 170.93 | 240.20 | 208.01 | 206.38 | 262.57 | 194.22 | 272.75 | 243.18 |
| ENSSSCG00000010703 | 14 | 221.43 | 214.80 | 181.12 | 205.78 | 82.06 | 150.69 | 178.00 | 136.92 |
| ENSSSCG00000014336 | 2 | 185.30 | 313.49 | 116.18 | 204.99 | 32.14 | 21.88 | 19.81 | 24.61 |
| ENSSSCG00000012022 | 13 | 183.04 | 241.18 | 190.70 | 204.97 | 206.22 | 138.05 | 131.48 | 158.58 |
| ENSSSCG00000017591 | 12 | 224.07 | 204.85 | 177.91 | 202.27 | 159.97 | 181.30 | 222.10 | 187.79 |
| ENSSSCG00000006395 | 4 | 284.50 | 248.44 | 73.46 | 202.13 | 119.80 | 116.80 | 164.11 | 133.57 |
| ENSSSCG00000033262 | 2 | 168.11 | 238.41 | 197.03 | 201.18 | 576.63 | 346.96 | 246.56 | 390.05 |
| ENSSSCG00000008061 | 3 | 209.24 | 191.81 | 202.48 | 201.17 | 186.12 | 171.77 | 225.53 | 194.47 |
| ENSSSCG00000000576 | 5 | 209.07 | 198.08 | 195.21 | 200.79 | 241.94 | 289.88 | 360.91 | 297.58 |
| ENSSSCG00000016522 | 18 | 178.55 | 147.75 | 273.90 | 200.06 | 26.88 | 70.67 | 58.06 | 51.87 |
| ENSSSCG00000025281 | 3 | 231.06 | 185.55 | 180.93 | 199.18 | 109.84 | 125.92 | 143.56 | 126.44 |
| ENSSSCG00000012961 | 2 | 212.48 | 175.65 | 202.29 | 196.81 | 253.87 | 167.89 | 227.25 | 216.34 |
| ENSSSCG00000017957 | 12 | 235.00 | 235.10 | 120.26 | 196.79 | 90.93 | 90.01 | 96.84 | 92.59 |
| ENSSSCG00000003691 | 6 | 197.20 | 182.52 | 208.42 | 196.05 | 166.68 | 150.32 | 195.70 | 170.90 |
| ENSSSCG00000014022 | 2 | 249.87 | 181.42 | 154.05 | 195.11 | 168.88 | 163.73 | 95.24 | 142.62 |
| ENSSSCG00000017024 | 16 | 310.02 | 125.33 | 148.68 | 194.68 | 113.51 | 140.85 | 70.09 | 108.15 |
| ENSSSCG00000002829 | 6 | 141.28 | 282.83 | 157.56 | 193.89 | 60.64 | 239.04 | 131.53 | 143.74 |
| ENSSSCG00000005601 | 1 | 245.58 | 210.11 | 125.79 | 193.82 | 135.99 | 108.52 | 196.22 | 146.91 |
| ENSSSCG00000031321 | 5 | 125.59 | 434.46 | 21.33 | 193.79 | 23.43 | 5.26 | 6.46 | 11.72 |
| ENSSSCG00000013181 | 2 | 176.55 | 186.57 | 217.83 | 193.65 | 226.72 | 154.76 | 169.34 | 183.60 |
| ENSSSCG00000027746 | 8 | 210.49 | 181.46 | 187.67 | 193.21 | 199.45 | 175.04 | 206.25 | 193.58 |
| ENSSSCG00000034448 | 6 | 179.74 | 202.59 | 195.26 | 192.53 | 186.83 | 223.70 | 258.69 | 223.08 |
| ENSSSCG00000017212 | 12 | 201.54 | 179.63 | 195.89 | 192.35 | 164.05 | 202.61 | 193.29 | 186.65 |
| ENSSSCG00000035419 | 18 | 131.59 | 201.98 | 240.18 | 191.25 | 94.02 | 164.86 | 116.17 | 125.02 |
| ENSSSCG00000009146 | 8 | 199.92 | 185.79 | 188.02 | 191.24 | 218.83 | 237.17 | 187.13 | 214.38 |
| ENSSSCG00000037508 | 1 | 176.26 | 252.16 | 142.84 | 190.42 | 189.38 | 252.68 | 204.39 | 215.48 |
| ENSSSCG00000012625 | X | 173.41 | 158.42 | 237.35 | 189.73 | 104.53 | 84.71 | 137.27 | 108.84 |
| ENSSSCG00000000010 | 5 | 139.11 | 217.73 | 210.08 | 188.97 | 248.85 | 206.02 | 283.29 | 246.05 |
| ENSSSCG00000000104 | 5 | 181.98 | 174.74 | 210.09 | 188.93 | 202.55 | 198.21 | 155.27 | 185.35 |
| ENSSSCG00000000084 | 5 | 198.68 | 214.54 | 153.47 | 188.89 | 100.85 | 95.19 | 129.63 | 108.56 |
| ENSSSCG00000007085 | 17 | 154.53 | 170.06 | 232.83 | 185.81 | 505.27 | 163.01 | 165.59 | 277.95 |
| ENSSSCG00000000633 | 5 | 231.43 | 211.27 | 114.56 | 185.75 | 38.56 | 80.62 | 62.10 | 60.42 |
| ENSSSCG00000029160 | 7 | 150.13 | 199.10 | 202.72 | 183.98 | 181.90 | 112.70 | 149.88 | 148.16 |
| ENSSSCG00000000405 | 5 | 213.58 | 161.49 | 175.25 | 183.44 | 178.41 | 143.67 | 178.98 | 167.02 |
| ENSSSCG00000013046 | 2 | 172.91 | 176.06 | 198.75 | 182.57 | 186.59 | 156.22 | 213.43 | 185.41 |
| ENSSSCG00000006887 | 4 | 230.19 | 201.41 | 116.08 | 182.56 | 115.47 | 129.00 | 130.34 | 124.94 |
| ENSSSCG00000003046 | 6 | 155.50 | 171.87 | 218.91 | 182.09 | 141.31 | 150.58 | 218.05 | 169.98 |
| ENSSSCG00000030309 | 7 | 171.04 | 171.94 | 203.07 | 182.02 | 148.19 | 208.34 | 113.95 | 156.82 |
| ENSSSCG00000017259 | 12 | 224.25 | 211.21 | 108.89 | 181.45 | 155.68 | 126.22 | 133.83 | 138.58 |
| ENSSSCG00000009772 | 14 | 199.83 | 172.82 | 170.03 | 180.89 | 180.38 | 153.26 | 197.90 | 177.18 |
| ENSSSCG00000038359 | 12 | 211.87 | 187.92 | 142.53 | 180.77 | 135.76 | 114.42 | 144.85 | 131.68 |
| ENSSSCG00000040162 | 3 | 147.11 | 202.46 | 190.11 | 179.89 | 71.10 | 179.41 | 112.89 | 121.13 |
| ENSSSCG00000030241 | X | 134.78 | 260.80 | 142.44 | 179.34 | 37.33 | 77.15 | 79.26 | 64.58 |
| ENSSSCG00000007743 | 3 | 193.33 | 179.90 | 164.21 | 179.15 | 139.68 | 121.98 | 184.28 | 148.65 |
| ENSSSCG00000028512 | 2 | 201.68 | 311.71 | 20.54 | 177.97 | 11.17 | 5.23 | 9.54 | 8.65 |
| ENSSSCG00000036716 | 15 | 182.16 | 172.75 | 176.43 | 177.11 | 187.07 | 179.70 | 214.50 | 193.76 |
| ENSSSCG00000011274 | 13 | 195.67 | 160.87 | 170.08 | 175.54 | 165.17 | 174.85 | 193.21 | 177.74 |
| ENSSSCG00000013736 | 2 | 169.88 | 206.05 | 147.15 | 174.36 | 49.27 | 55.37 | 101.12 | 68.59 |

| | | | | | | | | | |
|---|---|---|---|---|---|---|---|---|---|
| ENSSSCG00000024437 | 13 | 167.25 | 165.87 | 189.04 | 174.05 | 91.71 | 105.13 | 139.29 | 112.04 |
| ENSSSCG00000011727 | 13 | 15.39 | 329.64 | 177.04 | 174.02 | 2.75 | 8.29 | 22.64 | 11.23 |
| ENSSSCG00000022760 | 10 | 213.02 | 166.60 | 141.76 | 173.79 | 182.21 | 163.31 | 171.20 | 172.24 |
| ENSSSCG00000011351 | 13 | 182.84 | 162.87 | 174.88 | 173.53 | 177.14 | 170.27 | 203.76 | 183.72 |
| ENSSSCG00000006062 | 4 | 191.28 | 192.55 | 136.29 | 173.37 | 151.87 | 135.68 | 164.07 | 150.54 |
| ENSSSCG00000015545 | 9 | 142.02 | 161.16 | 215.86 | 173.01 | 72.54 | 132.41 | 139.26 | 114.73 |
| ENSSSCG00000002627 | 7 | 195.50 | 127.89 | 193.11 | 172.17 | 16.04 | 73.93 | 87.83 | 59.27 |
| ENSSSCG00000031262 | 4 | 173.29 | 143.12 | 199.45 | 171.95 | 296.65 | 289.97 | 183.51 | 256.71 |
| ENSSSCG00000038594 | 17 | 188.23 | 293.90 | 30.06 | 170.73 | 22.46 | 12.91 | 28.14 | 21.17 |
| ENSSSCG00000011867 | 13 | 205.65 | 142.40 | 162.96 | 170.34 | 982.17 | 115.38 | 122.17 | 406.57 |
| ENSSSCG00000033368 | 5 | 182.86 | 158.33 | 169.42 | 170.20 | 172.77 | 170.41 | 205.88 | 183.02 |
| ENSSSCG00000012534 | X | 195.65 | 155.03 | 159.62 | 170.10 | 137.72 | 126.74 | 150.24 | 138.23 |
| ENSSSCG00000004192 | 1 | 301.27 | 70.65 | 137.20 | 169.71 | 68.47 | 76.37 | 60.18 | 68.34 |
| ENSSSCG00000009851 | 14 | 168.68 | 156.56 | 182.39 | 169.21 | 162.30 | 162.70 | 167.94 | 164.31 |
| ENSSSCG00000006624 | 4 | 207.08 | 125.15 | 175.08 | 169.11 | 22.90 | 143.15 | 62.87 | 76.31 |
| ENSSSCG00000021084 | 4 | 180.98 | 136.37 | 184.18 | 167.18 | 112.10 | 148.38 | 270.12 | 176.87 |
| ENSSSCG00000035392 | 15 | 290.64 | 116.81 | 88.36 | 165.27 | 58.41 | 75.65 | 131.39 | 88.48 |
| ENSSSCG00000014156 | 2 | 140.16 | 212.83 | 130.04 | 161.01 | 50.48 | 79.28 | 50.69 | 60.15 |
| ENSSSCG00000003914 | 6 | 203.84 | 127.50 | 148.44 | 159.93 | 202.82 | 122.23 | 184.66 | 169.91 |
| ENSSSCG00000006954 | 4 | 181.64 | 151.25 | 146.59 | 159.83 | 187.80 | 176.69 | 193.03 | 185.84 |
| ENSSSCG00000037697 | 5 | 120.50 | 106.75 | 251.31 | 159.52 | 168.34 | 193.95 | 193.51 | 185.27 |
| ENSSSCG00000011101 | 10 | 156.12 | 148.23 | 170.49 | 158.28 | 118.25 | 103.25 | 98.32 | 106.61 |
| ENSSSCG00000033733 | 13 | 192.02 | 146.47 | 133.94 | 157.48 | 168.89 | 148.77 | 162.41 | 160.02 |
| ENSSSCG00000034072 | 5 | 193.11 | 161.09 | 118.12 | 157.44 | 145.33 | 111.83 | 160.28 | 139.14 |
| ENSSSCG00000015324 | 9 | 127.62 | 141.15 | 198.70 | 155.82 | 249.70 | 203.27 | 155.36 | 202.78 |
| ENSSSCG00000015322 | 9 | 110.12 | 153.18 | 203.63 | 155.64 | 251.40 | 158.08 | 255.97 | 221.82 |
| ENSSSCG00000040879 | 13 | 173.92 | 150.80 | 141.47 | 155.40 | 153.61 | 132.36 | 102.12 | 129.36 |
| ENSSSCG00000032165 | 14 | 176.22 | 142.73 | 141.65 | 153.53 | 119.33 | 134.37 | 138.75 | 130.82 |
| ENSSSCG00000036956 | 12 | 91.44 | 356.32 | 11.84 | 153.20 | 6.16 | 7.64 | 8.76 | 7.52 |
| ENSSSCG00000022506 | 13 | 141.95 | 175.84 | 141.74 | 153.18 | 152.86 | 122.61 | 126.47 | 133.98 |
| ENSSSCG00000015973 | 15 | 168.89 | 138.29 | 150.21 | 152.46 | 122.26 | 108.05 | 139.67 | 123.33 |
| ENSSSCG00000035684 | 7 | 172.75 | 136.83 | 147.03 | 152.20 | 152.17 | 138.46 | 206.21 | 165.62 |
| ENSSSCG00000008638 | 3 | 139.62 | 160.45 | 155.13 | 151.74 | 98.15 | 97.19 | 113.86 | 103.07 |
| ENSSSCG00000009327 | 11 | 169.92 | 126.64 | 157.79 | 151.45 | 143.41 | 152.27 | 121.42 | 139.03 |
| ENSSSCG00000037241 | 10 | 77.45 | 261.09 | 115.73 | 151.42 | 59.21 | 27.26 | 24.41 | 36.96 |
| ENSSSCG00000033727 | 13 | 115.74 | 136.41 | 201.37 | 151.17 | 127.27 | 90.26 | 176.15 | 131.22 |
| ENSSSCG00000031888 | 14 | 173.21 | 221.85 | 54.26 | 149.77 | 48.15 | 77.29 | 49.91 | 58.45 |
| ENSSSCG00000000522 | 5 | 144.09 | 152.67 | 152.25 | 149.67 | 144.68 | 190.87 | 145.74 | 160.43 |
| ENSSSCG00000000038 | 5 | 139.14 | 147.27 | 162.53 | 149.65 | 50.47 | 60.85 | 74.59 | 61.97 |
| ENSSSCG00000016535 | 18 | 99.17 | 160.79 | 187.90 | 149.28 | 152.80 | 115.96 | 48.71 | 105.82 |
| ENSSSCG00000029034 | 6 | 156.43 | 152.39 | 138.08 | 148.97 | 129.17 | 137.60 | 144.98 | 137.25 |
| ENSSSCG00000016554 | 18 | 274.40 | 141.49 | 30.73 | 148.88 | 6.06 | 20.55 | 19.46 | 15.36 |
| ENSSSCG00000039314 | 4 | 136.73 | 210.37 | 97.80 | 148.30 | 114.11 | 104.64 | 96.12 | 104.96 |
| ENSSSCG00000012042 | 13 | 161.99 | 128.63 | 152.73 | 147.78 | 172.86 | 151.13 | 190.73 | 171.57 |
| ENSSSCG00000032367 | 4 | 125.67 | 241.84 | 72.78 | 146.76 | 28.62 | 55.98 | 35.54 | 40.05 |
| ENSSSCG00000008056 | 3 | 129.74 | 154.95 | 153.13 | 145.94 | 100.89 | 105.76 | 115.46 | 107.37 |
| ENSSSCG00000035622 | 8 | 161.51 | 156.21 | 119.73 | 145.82 | 109.01 | 102.95 | 107.85 | 106.61 |
| ENSSSCG00000014071 | 2 | 150.95 | 151.89 | 134.49 | 145.78 | 180.19 | 166.43 | 155.50 | 167.37 |
| ENSSSCG00000026257 | 6 | 234.39 | 98.50 | 102.58 | 145.16 | 42.47 | 131.75 | 111.55 | 95.25 |
| ENSSSCG00000034313 | 6 | 128.81 | 104.67 | 201.19 | 144.89 | 170.73 | 99.32 | 133.24 | 134.43 |
| ENSSSCG00000004414 | 1 | 137.25 | 156.61 | 138.75 | 144.20 | 119.05 | 111.10 | 119.37 | 116.51 |
| ENSSSCG00000000660 | 5 | 167.22 | 154.05 | 109.46 | 143.58 | 9.68 | 77.69 | 65.07 | 50.81 |
| ENSSSCG00000031023 | 1 | 117.53 | 130.48 | 181.88 | 143.30 | 2.70 | 17.72 | 24.49 | 14.97 |
| ENSSSCG00000008980 | 8 | 123.59 | 129.59 | 175.70 | 142.96 | 49.19 | 98.38 | 83.71 | 77.09 |
| ENSSSCG00000006359 | 4 | 63.73 | 358.80 | 6.23 | 142.92 | 0.53 | 1.02 | 2.82 | 1.46 |
| ENSSSCG00000025768 | 7 | 133.66 | 116.45 | 178.00 | 142.70 | 350.96 | 288.94 | 346.93 | 328.94 |

| | | | | | | | | | |
|---|---|---|---|---|---|---|---|---|---|
| ENSSSCG00000014016 | 2 | 126.89 | 140.12 | 161.09 | 142.70 | 93.39 | 133.69 | 119.05 | 115.38 |
| ENSSSCG00000006237 | 4 | 126.53 | 177.96 | 123.59 | 142.69 | 78.09 | 129.33 | 100.21 | 102.54 |
| ENSSSCG00000002963 | 6 | 151.21 | 137.94 | 135.82 | 141.66 | 128.56 | 138.62 | 151.35 | 139.51 |
| ENSSSCG00000016267 | 15 | 112.02 | 167.37 | 143.92 | 141.10 | 109.27 | 59.49 | 122.45 | 97.07 |
| ENSSSCG00000004058 | 1 | 93.75 | 161.38 | 167.29 | 140.81 | 162.22 | 114.41 | 207.40 | 161.35 |
| ENSSSCG00000001868 | 7 | 147.01 | 135.32 | 139.98 | 140.77 | 122.64 | 136.36 | 130.92 | 129.97 |
| ENSSSCG00000039894 | Y | 121.23 | 169.03 | 131.57 | 140.61 | 73.65 | 118.09 | 143.29 | 111.67 |
| ENSSSCG00000002366 | 7 | 166.09 | 121.86 | 132.62 | 140.19 | 47.49 | 102.14 | 75.99 | 75.21 |
| ENSSSCG00000008540 | 3 | 120.04 | 177.00 | 122.89 | 139.98 | 127.71 | 104.40 | 100.95 | 111.02 |
| ENSSSCG00000022080 | 13 | 131.76 | 150.19 | 137.97 | 139.97 | 82.62 | 82.80 | 113.68 | 93.03 |
| ENSSSCG00000021880 | 6 | 137.98 | 147.52 | 134.03 | 139.84 | 45.76 | 145.76 | 66.79 | 86.10 |
| ENSSSCG00000012583 | X | 123.40 | 255.01 | 40.30 | 139.57 | 11.24 | 17.79 | 13.92 | 14.32 |
| ENSSSCG00000001769 | 7 | 144.75 | 128.56 | 142.90 | 138.74 | 141.89 | 135.54 | 150.67 | 142.70 |
| ENSSSCG00000027907 | 2 | 150.33 | 135.07 | 128.53 | 137.98 | 107.15 | 108.09 | 137.67 | 117.64 |
| ENSSSCG00000016075 | 15 | 135.68 | 137.84 | 139.94 | 137.82 | 136.57 | 170.29 | 101.05 | 135.97 |
| ENSSSCG00000039854 | 4 | 61.66 | 277.66 | 72.62 | 137.31 | 1.55 | 44.05 | 13.66 | 19.75 |
| ENSSSCG00000005316 | 1 | 100.83 | 134.01 | 176.78 | 137.21 | 611.22 | 36.81 | 83.41 | 243.81 |
| ENSSSCG00000013906 | 2 | 140.48 | 136.67 | 134.06 | 137.07 | 116.68 | 126.86 | 155.42 | 132.98 |
| ENSSSCG00000003187 | 6 | 137.16 | 131.71 | 140.26 | 136.38 | 122.36 | 149.30 | 188.72 | 153.46 |
| ENSSSCG00000000645 | 5 | 144.85 | 157.60 | 106.57 | 136.34 | 55.37 | 83.62 | 73.99 | 70.99 |
| ENSSSCG00000015056 | 9 | 220.75 | 106.10 | 81.19 | 136.01 | 52.55 | 67.48 | 93.99 | 71.34 |
| ENSSSCG00000004499 | 1 | 143.58 | 124.21 | 138.93 | 135.57 | 124.73 | 105.08 | 144.66 | 124.82 |
| ENSSSCG00000011399 | 13 | 130.48 | 137.65 | 137.77 | 135.30 | 123.34 | 120.33 | 135.85 | 126.50 |
| ENSSSCG00000021890 | 6 | 143.61 | 133.06 | 128.40 | 135.02 | 97.86 | 103.21 | 115.37 | 105.48 |
| ENSSSCG00000015337 | 9 | 146.68 | 123.35 | 134.93 | 134.98 | 116.25 | 120.44 | 115.45 | 117.38 |
| ENSSSCG00000034993 | 2 | 83.02 | 165.93 | 154.72 | 134.56 | 51.74 | 84.35 | 73.09 | 69.72 |
| ENSSSCG00000038628 | 2 | 142.56 | 129.10 | 130.84 | 134.17 | 156.38 | 125.13 | 124.71 | 135.41 |
| ENSSSCG00000002449 | 7 | 91.82 | 94.72 | 214.98 | 133.84 | 66.16 | 78.70 | 106.99 | 83.95 |
| ENSSSCG00000024791 | 2 | 117.85 | 122.55 | 160.12 | 133.51 | 217.77 | 227.07 | 280.96 | 241.93 |
| ENSSSCG00000005029 | 1 | 114.56 | 129.95 | 153.75 | 132.75 | 125.71 | 144.12 | 135.23 | 135.02 |
| ENSSSCG00000003520 | 6 | 136.76 | 135.02 | 126.12 | 132.63 | 114.54 | 122.02 | 148.27 | 128.28 |
| ENSSSCG00000004570 | 1 | 144.80 | 105.36 | 147.15 | 132.44 | 442.46 | 73.35 | 117.45 | 211.08 |
| ENSSSCG00000039514 | 6 | 165.37 | 155.34 | 70.86 | 130.52 | 49.95 | 117.41 | 55.82 | 74.39 |
| ENSSSCG00000012772 | X | 122.65 | 123.40 | 145.28 | 130.44 | 98.21 | 78.45 | 108.11 | 94.93 |
| ENSSSCG00000027130 | 3 | 136.97 | 208.28 | 43.99 | 129.75 | 12.83 | 18.52 | 23.09 | 18.15 |
| ENSSSCG00000016998 | 16 | 139.69 | 121.90 | 127.10 | 129.56 | 127.16 | 143.66 | 118.73 | 129.85 |
| ENSSSCG00000009545 | 11 | 135.35 | 127.45 | 124.50 | 129.10 | 39.13 | 55.38 | 86.51 | 60.34 |
| ENSSSCG00000006791 | 4 | 337.13 | 16.03 | 32.64 | 128.60 | 3957.80 | 3498.06 | 12802.90 | 6752.92 |
| ENSSSCG00000027357 | 13 | 92.75 | 174.81 | 118.17 | 128.58 | 86.63 | 113.05 | 99.19 | 99.62 |
| ENSSSCG00000033844 | 3 | 78.78 | 187.54 | 119.28 | 128.53 | 3.81 | 39.12 | 19.20 | 20.71 |
| ENSSSCG00000016185 | 15 | 118.74 | 137.89 | 127.19 | 127.94 | 91.38 | 101.48 | 115.28 | 102.71 |
| ENSSSCG00000013992 | 2 | 137.85 | 128.04 | 117.29 | 127.73 | 143.36 | 117.84 | 169.49 | 143.56 |
| ENSSSCG00000012597 | X | 153.76 | 99.91 | 129.22 | 127.63 | 42.01 | 165.29 | 83.59 | 96.96 |
| ENSSSCG00000001910 | 7 | 105.46 | 150.67 | 124.36 | 126.83 | 18.89 | 72.40 | 63.50 | 51.59 |
| ENSSSCG00000034131 | 7 | 156.26 | 116.40 | 107.57 | 126.74 | 66.35 | 102.05 | 117.69 | 95.36 |
| ENSSSCG00000025966 | 3 | 136.38 | 130.73 | 113.02 | 126.71 | 116.72 | 111.44 | 126.59 | 118.25 |
| ENSSSCG00000010203 | 14 | 148.08 | 137.95 | 93.95 | 126.66 | 153.41 | 134.34 | 148.44 | 145.40 |
| ENSSSCG00000002704 | 6 | 134.55 | 130.46 | 114.81 | 126.61 | 123.38 | 135.93 | 139.42 | 132.91 |
| ENSSSCG00000030996 | 14 | 147.80 | 143.71 | 87.76 | 126.42 | 118.41 | 94.45 | 142.48 | 118.45 |
| ENSSSCG00000002039 | 7 | 97.63 | 173.08 | 106.50 | 125.74 | 60.49 | 172.68 | 101.68 | 111.62 |
| ENSSSCG00000008987 | 8 | 131.03 | 126.78 | 118.66 | 125.49 | 121.26 | 149.09 | 143.93 | 138.09 |
| ENSSSCG00000020790 | 15 | 111.94 | 123.32 | 140.58 | 125.28 | 103.73 | 107.77 | 157.64 | 123.05 |
| ENSSSCG00000000071 | 5 | 123.98 | 120.04 | 130.97 | 125.00 | 134.62 | 120.37 | 120.28 | 125.09 |
| ENSSSCG00000034937 | 6 | 117.30 | 127.21 | 128.52 | 124.34 | 84.76 | 108.93 | 110.33 | 101.34 |
| ENSSSCG00000038947 | 7 | 129.86 | 130.07 | 113.06 | 124.33 | 130.42 | 127.12 | 158.50 | 138.68 |
| ENSSSCG00000002296 | 7 | 69.92 | 133.78 | 167.01 | 123.57 | 10.62 | 25.31 | 43.78 | 26.57 |

| | | | | | | | | | |
|---|---|---|---|---|---|---|---|---|---|
| ENSSSCG00000010591 | 14 | 273.72 | 2.69 | 93.89 | 123.43 | 0.10 | 0.49 | 7.90 | 2.83 |
| ENSSSCG00000039243 | 2 | 119.92 | 133.91 | 115.15 | 122.99 | 190.09 | 119.75 | 138.81 | 149.55 |
| ENSSSCG00000016991 | 16 | 64.00 | 231.46 | 72.74 | 122.73 | 86.44 | 34.32 | 37.33 | 52.70 |
| ENSSSCG00000017752 | 12 | 117.58 | 126.99 | 122.77 | 122.44 | 176.14 | 188.25 | 128.62 | 164.33 |
| ENSSSCG00000000089 | 5 | 137.73 | 125.97 | 102.19 | 121.96 | 117.65 | 109.98 | 127.46 | 118.36 |
| ENSSSCG00000038989 | 12 | 144.27 | 118.86 | 102.75 | 121.96 | 136.76 | 108.47 | 140.69 | 128.64 |
| ENSSSCG00000002873 | 6 | 66.89 | 110.45 | 187.26 | 121.53 | 69.37 | 65.64 | 95.48 | 76.83 |
| ENSSSCG00000022300 | 3 | 140.97 | 164.71 | 58.77 | 121.48 | 89.89 | 85.39 | 57.77 | 77.68 |
| ENSSSCG00000032330 | 9 | 59.01 | 153.81 | 149.15 | 120.66 | 42.45 | 82.07 | 72.65 | 65.72 |
| ENSSSCG00000035446 | 12 | 136.22 | 118.49 | 106.71 | 120.47 | 97.40 | 75.30 | 104.56 | 92.42 |
| ENSSSCG00000034086 | 10 | 225.11 | 77.70 | 57.19 | 120.00 | 31.77 | 35.29 | 46.32 | 37.79 |
| ENSSSCG00000004237 | 1 | 93.83 | 124.28 | 139.34 | 119.15 | 108.39 | 130.11 | 91.46 | 109.98 |
| ENSSSCG00000025295 | 2 | 104.29 | 121.95 | 131.04 | 119.09 | 95.89 | 104.61 | 127.70 | 109.40 |
| ENSSSCG00000007522 | 17 | 50.22 | 147.58 | 159.20 | 119.00 | 51.76 | 98.93 | 63.22 | 71.30 |
| ENSSSCG00000033299 | X | 111.82 | 123.48 | 121.46 | 118.92 | 311.64 | 62.34 | 73.79 | 149.26 |
| ENSSSCG00000035080 | 12 | 125.76 | 113.68 | 116.34 | 118.59 | 134.65 | 122.00 | 114.97 | 123.87 |
| ENSSSCG00000009422 | 11 | 80.98 | 216.36 | 57.84 | 118.39 | 74.98 | 59.17 | 58.44 | 64.20 |
| ENSSSCG00000010682 | 14 | 92.65 | 103.83 | 157.65 | 118.05 | 70.96 | 70.81 | 93.35 | 78.37 |
| ENSSSCG00000002623 | 7 | 49.38 | 106.91 | 196.48 | 117.59 | 3.15 | 12.45 | 71.75 | 29.12 |
| ENSSSCG00000001400 | 7 | 147.17 | 100.69 | 104.77 | 117.54 | 113.44 | 103.98 | 99.80 | 105.74 |
| ENSSSCG00000035090 | 3 | 115.67 | 118.45 | 117.92 | 117.34 | 97.85 | 97.54 | 110.45 | 101.94 |
| ENSSSCG00000040777 | 6 | 126.20 | 109.65 | 115.64 | 117.16 | 143.51 | 143.11 | 145.11 | 143.91 |
| ENSSSCG00000024267 | 6 | 89.47 | 124.31 | 136.06 | 116.61 | 100.97 | 102.58 | 122.45 | 108.67 |
| ENSSSCG00000001453 | 7 | 76.61 | 159.20 | 112.14 | 115.98 | 404.42 | 280.94 | 186.02 | 290.46 |
| ENSSSCG00000003653 | 6 | 121.65 | 104.93 | 119.72 | 115.43 | 112.10 | 118.07 | 150.74 | 126.97 |
| ENSSSCG00000023903 | 8 | 121.67 | 99.73 | 123.56 | 114.99 | 122.85 | 86.47 | 132.12 | 113.81 |
| ENSSSCG00000010877 | 10 | 129.76 | 120.62 | 90.93 | 113.77 | 100.07 | 103.71 | 86.63 | 96.80 |
| ENSSSCG00000035297 | 7 | 97.64 | 41.49 | 200.89 | 113.34 | 375.72 | 68.26 | 167.66 | 203.88 |
| ENSSSCG00000010320 | 14 | 101.05 | 113.54 | 124.89 | 113.16 | 71.14 | 64.52 | 88.96 | 74.87 |
| ENSSSCG00000020976 | 10 | 115.07 | 112.99 | 111.42 | 113.16 | 315.25 | 83.69 | 86.43 | 161.79 |
| ENSSSCG00000039647 | 9 | 138.80 | 101.49 | 98.65 | 112.98 | 90.71 | 102.04 | 124.08 | 105.61 |
| ENSSSCG00000009555 | 11 | 99.93 | 121.74 | 117.14 | 112.93 | 87.31 | 100.86 | 100.81 | 96.33 |
| ENSSSCG00000036114 | 13 | 115.35 | 109.99 | 109.78 | 111.70 | 55.12 | 89.64 | 70.28 | 71.68 |
| ENSSSCG00000031249 | 13 | 124.37 | 121.89 | 88.36 | 111.54 | 64.95 | 105.16 | 79.15 | 83.09 |
| ENSSSCG00000036592 | 12 | 144.48 | 101.34 | 87.92 | 111.25 | 98.63 | 89.20 | 75.52 | 87.79 |
| ENSSSCG00000022725 | 7 | 145.48 | 96.01 | 91.80 | 111.10 | 64.45 | 83.13 | 82.77 | 76.78 |
| ENSSSCG00000013401 | 2 | 158.76 | 70.65 | 102.96 | 110.79 | 12.84 | 49.00 | 84.05 | 48.63 |
| ENSSSCG00000026606 | 2 | 116.91 | 111.44 | 103.43 | 110.59 | 117.03 | 130.27 | 107.34 | 118.21 |
| ENSSSCG00000014020 | 2 | 121.17 | 105.24 | 105.10 | 110.50 | 133.58 | 100.69 | 126.15 | 120.14 |
| ENSSSCG00000023304 | 3 | 116.92 | 108.76 | 105.62 | 110.43 | 68.42 | 105.38 | 64.26 | 79.35 |
| ENSSSCG00000006527 | 4 | 107.70 | 110.92 | 110.02 | 109.55 | 108.32 | 98.43 | 141.78 | 116.17 |
| ENSSSCG00000032527 | 3 | 98.83 | 200.63 | 26.67 | 108.71 | 13.98 | 33.60 | 24.73 | 24.10 |
| ENSSSCG00000005055 | 1 | 110.23 | 115.17 | 100.51 | 108.63 | 249.15 | 213.19 | 268.93 | 243.76 |
| ENSSSCG00000032265 | 2 | 137.16 | 100.73 | 87.19 | 108.36 | 61.54 | 86.98 | 82.22 | 76.91 |
| ENSSSCG00000011137 | 10 | 115.64 | 107.89 | 101.36 | 108.30 | 52.36 | 61.25 | 86.61 | 66.74 |
| ENSSSCG00000016792 | 16 | 240.91 | 50.41 | 33.48 | 108.27 | 15.53 | 28.96 | 30.26 | 24.92 |
| ENSSSCG00000011747 | 13 | 74.59 | 123.74 | 126.43 | 108.25 | 4.98 | 69.56 | 44.22 | 39.59 |
| ENSSSCG00000040494 | 5 | 117.94 | 104.88 | 100.46 | 107.76 | 107.16 | 97.32 | 116.31 | 106.93 |
| ENSSSCG00000016892 | 16 | 53.12 | 229.97 | 39.30 | 107.46 | 1.06 | 34.35 | 12.60 | 16.00 |
| ENSSSCG00000011171 | 10 | 92.18 | 111.36 | 118.65 | 107.40 | 109.63 | 80.69 | 113.95 | 101.42 |
| ENSSSCG00000006016 | 4 | 116.40 | 105.94 | 99.25 | 107.20 | 142.87 | 133.54 | 130.46 | 135.63 |
| ENSSSCG00000015560 | 9 | 84.81 | 86.35 | 150.38 | 107.18 | 58.48 | 73.39 | 87.58 | 73.15 |
| ENSSSCG00000023562 | 9 | 122.61 | 99.77 | 97.71 | 106.70 | 97.49 | 109.25 | 108.32 | 105.02 |
| ENSSSCG00000001549 | 7 | 53.68 | 126.91 | 139.02 | 106.53 | 16.56 | 29.29 | 27.42 | 24.42 |
| ENSSSCG00000000119 | 5 | 105.52 | 106.91 | 106.33 | 106.25 | 90.88 | 107.95 | 119.31 | 106.05 |
| ENSSSCG00000023245 | X | 127.76 | 87.93 | 102.45 | 106.04 | 68.89 | 95.11 | 98.54 | 87.52 |

| | | | | | | | | | |
|---|---|---|---|---|---|---|---|---|---|
| ENSSSCG00000000808 | 5 | 83.16 | 123.12 | 110.85 | 105.71 | 38.85 | 67.58 | 40.75 | 49.06 |
| ENSSSCG00000028322 | 9 | 86.49 | 197.82 | 32.46 | 105.59 | 26.74 | 20.19 | 22.16 | 23.03 |
| ENSSSCG00000011000 | 10 | 98.04 | 138.68 | 79.35 | 105.36 | 74.30 | 57.97 | 51.62 | 61.29 |
| ENSSSCG00000003505 | 6 | 105.08 | 102.42 | 107.20 | 104.90 | 80.14 | 80.16 | 129.68 | 96.66 |
| ENSSSCG00000037360 | 17 | 85.90 | 120.17 | 108.61 | 104.90 | 137.76 | 100.62 | 102.39 | 113.59 |
| ENSSSCG00000016581 | 18 | 122.64 | 94.11 | 97.32 | 104.69 | 60.37 | 63.98 | 79.31 | 67.89 |
| ENSSSCG00000028423 | 6 | 115.56 | 95.12 | 103.28 | 104.66 | 98.42 | 108.93 | 99.11 | 102.15 |
| ENSSSCG00000027121 | 3 | 133.21 | 127.20 | 53.51 | 104.64 | 121.47 | 66.00 | 155.11 | 114.20 |
| ENSSSCG00000028227 | 17 | 98.81 | 88.80 | 125.01 | 104.21 | 63.31 | 52.02 | 85.38 | 66.90 |
| ENSSSCG00000014921 | 9 | 102.38 | 155.01 | 54.40 | 103.93 | 22.67 | 36.94 | 53.30 | 37.64 |
| ENSSSCG00000012252 | X | 102.84 | 113.22 | 95.54 | 103.87 | 89.55 | 81.88 | 66.64 | 79.36 |
| ENSSSCG00000015384 | 9 | 94.99 | 110.40 | 106.21 | 103.87 | 91.23 | 106.89 | 110.96 | 103.03 |
| ENSSSCG00000035790 | 5 | 104.39 | 119.70 | 87.50 | 103.86 | 116.64 | 67.27 | 76.06 | 86.65 |
| ENSSSCG00000028536 | 2 | 81.05 | 150.20 | 79.45 | 103.57 | 23.37 | 50.29 | 66.29 | 46.65 |
| ENSSSCG00000037803 | 1 | 129.80 | 84.51 | 95.62 | 103.31 | 49.01 | 123.03 | 76.17 | 82.74 |
| ENSSSCG00000016128 | 15 | 102.00 | 111.20 | 96.64 | 103.28 | 112.87 | 118.92 | 134.53 | 122.11 |
| ENSSSCG00000003903 | 6 | 101.58 | 100.18 | 107.96 | 103.24 | 129.00 | 110.72 | 117.50 | 119.07 |
| ENSSSCG00000012403 | X | 65.92 | 108.06 | 134.96 | 102.98 | 97.99 | 114.61 | 75.00 | 95.87 |
| ENSSSCG00000036711 | 2 | 103.27 | 114.18 | 91.37 | 102.94 | 81.97 | 69.30 | 85.36 | 78.88 |
| ENSSSCG00000033019 | 1 | 103.65 | 102.79 | 101.37 | 102.60 | 92.70 | 99.21 | 114.35 | 102.09 |
| ENSSSCG00000013602 | 2 | 127.56 | 92.28 | 87.09 | 102.31 | 85.33 | 75.53 | 73.26 | 78.04 |
| ENSSSCG00000035690 | 6 | 126.72 | 109.62 | 70.25 | 102.20 | 78.90 | 70.46 | 56.23 | 68.53 |
| ENSSSCG00000032580 | 5 | 96.87 | 75.18 | 132.91 | 101.65 | 99.68 | 141.20 | 150.61 | 130.49 |
| ENSSSCG00000002452 | 7 | 71.61 | 134.42 | 98.71 | 101.58 | 56.95 | 115.77 | 113.69 | 95.47 |
| ENSSSCG00000006033 | 4 | 93.32 | 107.34 | 103.84 | 101.50 | 111.81 | 164.47 | 97.82 | 124.70 |
| ENSSSCG00000013421 | 2 | 127.71 | 101.14 | 75.54 | 101.46 | 94.61 | 78.09 | 87.16 | 86.62 |
| ENSSSCG00000034227 | 1 | 106.25 | 101.65 | 96.23 | 101.38 | 95.21 | 103.34 | 119.57 | 106.04 |
| ENSSSCG00000034942 | 15 | 111.90 | 104.71 | 87.37 | 101.33 | 0.50 | 2.11 | 2.33 | 1.65 |
| ENSSSCG00000006556 | 4 | 145.36 | 89.55 | 68.44 | 101.12 | 73.79 | 100.51 | 140.74 | 105.01 |
| ENSSSCG00000036307 | 9 | 107.65 | 138.56 | 54.87 | 100.36 | 16.69 | 46.39 | 39.76 | 34.28 |
| ENSSSCG00000008692 | 8 | 50.56 | 103.57 | 146.74 | 100.29 | 41.63 | 48.00 | 71.19 | 53.61 |
| ENSSSCG00000006752 | 4 | 97.13 | 94.38 | 108.81 | 100.10 | 102.95 | 106.87 | 77.10 | 95.64 |
| ENSSSCG00000009889 | 14 | 52.78 | 127.16 | 120.10 | 100.01 | 42.85 | 69.86 | 61.58 | 58.10 |
| ENSSSCG00000035945 | 9 | 107.55 | 110.25 | 82.05 | 99.95 | 13.03 | 22.27 | 20.47 | 18.59 |
| ENSSSCG00000026761 | 13 | 81.51 | 95.03 | 121.90 | 99.48 | 62.98 | 60.85 | 61.08 | 61.64 |
| ENSSSCG00000009222 | 8 | 66.22 | 160.35 | 71.25 | 99.28 | 278.06 | 167.07 | 68.30 | 171.14 |
| ENSSSCG00000005373 | 1 | 96.57 | 88.29 | 112.94 | 99.27 | 88.53 | 95.40 | 99.75 | 94.56 |
| ENSSSCG00000003493 | 6 | 113.92 | 78.24 | 104.89 | 99.01 | 90.25 | 87.58 | 102.42 | 93.42 |
| ENSSSCG00000015710 | 15 | 104.36 | 110.33 | 82.33 | 99.01 | 53.64 | 62.68 | 60.17 | 58.83 |
| ENSSSCG00000009759 | 14 | 188.68 | 46.63 | 61.64 | 98.98 | 19.11 | 27.91 | 43.45 | 30.16 |
| ENSSSCG00000031640 | 6 | 58.56 | 133.85 | 104.30 | 98.90 | 64.47 | 154.68 | 58.84 | 92.67 |
| ENSSSCG00000001834 | 7 | 110.33 | 88.99 | 96.71 | 98.68 | 64.20 | 52.48 | 46.90 | 54.53 |
| ENSSSCG00000002383 | 7 | 80.84 | 180.09 | 34.17 | 98.37 | 53.97 | 8.23 | 25.87 | 29.36 |
| ENSSSCG00000022398 | 9 | 107.46 | 90.05 | 96.95 | 98.15 | 106.91 | 100.89 | 88.36 | 98.72 |
| ENSSSCG00000001229 | 7 | 76.36 | 128.66 | 89.24 | 98.09 | 675.92 | 388.91 | 406.45 | 490.43 |
| ENSSSCG00000005096 | 1 | 161.08 | 72.98 | 59.11 | 97.72 | 41.07 | 54.96 | 46.65 | 47.56 |
| ENSSSCG00000031380 | 6 | 175.37 | 74.05 | 43.74 | 97.72 | 42.32 | 64.17 | 67.44 | 57.97 |
| ENSSSCG00000016714 | 18 | 65.51 | 197.38 | 30.14 | 97.68 | 31.02 | 25.33 | 27.61 | 27.99 |
| ENSSSCG00000006740 | 4 | 100.39 | 107.83 | 84.50 | 97.58 | 280.00 | 156.03 | 334.38 | 256.80 |
| ENSSSCG00000005656 | 1 | 117.34 | 88.90 | 85.34 | 97.20 | 98.31 | 85.37 | 79.48 | 87.72 |
| ENSSSCG00000015988 | 15 | 121.16 | 86.67 | 83.01 | 96.95 | 99.32 | 88.83 | 77.17 | 88.44 |
| ENSSSCG00000017929 | 12 | 86.96 | 96.41 | 105.24 | 96.20 | 61.48 | 71.07 | 88.15 | 73.57 |
| ENSSSCG00000004385 | 1 | 87.26 | 92.62 | 107.54 | 95.81 | 79.53 | 82.46 | 71.55 | 77.85 |
| ENSSSCG00000028414 | 6 | 116.33 | 79.27 | 91.70 | 95.77 | 90.60 | 88.74 | 118.65 | 99.33 |
| ENSSSCG00000009464 | 11 | 81.80 | 123.28 | 82.10 | 95.73 | 22.35 | 30.15 | 37.94 | 30.15 |
| ENSSSCG00000012699 | X | 103.44 | 79.34 | 103.94 | 95.58 | 129.14 | 84.08 | 59.68 | 90.96 |

| | | | | | | | | | |
|---|---|---|---|---|---|---|---|---|---|
| ENSSSCG00000003132 | 6 | 88.57 | 88.66 | 109.23 | 95.49 | 68.04 | 72.19 | 97.19 | 79.14 |
| ENSSSCG00000003332 | 6 | 89.58 | 95.80 | 100.03 | 95.14 | 77.64 | 82.78 | 122.17 | 94.19 |
| ENSSSCG00000031866 | 5 | 120.56 | 92.44 | 72.22 | 95.07 | 122.18 | 64.11 | 68.20 | 84.83 |
| ENSSSCG00000010517 | 14 | 93.44 | 90.70 | 100.85 | 95.00 | 86.44 | 82.30 | 94.43 | 87.72 |
| ENSSSCG00000003702 | 6 | 74.66 | 89.45 | 120.62 | 94.91 | 20.88 | 51.82 | 55.73 | 42.81 |
| ENSSSCG00000017525 | 12 | 90.03 | 99.95 | 94.68 | 94.89 | 109.77 | 124.52 | 145.40 | 126.56 |
| ENSSSCG00000007362 | 17 | 119.40 | 92.96 | 71.84 | 94.73 | 81.80 | 84.46 | 75.63 | 80.63 |
| ENSSSCG00000033854 | 5 | 74.30 | 99.00 | 110.46 | 94.58 | 53.44 | 84.68 | 94.34 | 77.48 |
| ENSSSCG00000004480 | 1 | 122.52 | 67.03 | 94.19 | 94.58 | 94.01 | 82.68 | 125.05 | 100.58 |
| ENSSSCG00000009150 | 8 | 53.00 | 59.08 | 171.64 | 94.58 | 68.08 | 60.84 | 153.26 | 94.06 |
| ENSSSCG00000032213 | 15 | 107.68 | 89.13 | 85.19 | 94.00 | 73.67 | 75.62 | 83.54 | 77.61 |
| ENSSSCG00000034570 | 6 | 86.73 | 54.08 | 141.19 | 94.00 | 247.44 | 123.39 | 191.59 | 187.47 |
| ENSSSCG00000013427 | 2 | 103.55 | 85.64 | 92.53 | 93.91 | 167.66 | 169.05 | 133.85 | 156.85 |
| ENSSSCG00000011110 | 10 | 166.89 | 60.88 | 53.49 | 93.76 | 64.23 | 50.22 | 49.81 | 54.75 |
| ENSSSCG00000024149 | 7 | 50.90 | 131.11 | 99.04 | 93.69 | 22.82 | 29.52 | 38.49 | 30.28 |
| ENSSSCG00000001038 | 7 | 90.17 | 82.16 | 108.11 | 93.48 | 136.38 | 104.58 | 151.77 | 130.91 |
| ENSSSCG00000014031 | 2 | 124.27 | 96.74 | 58.69 | 93.23 | 59.76 | 55.28 | 60.19 | 58.41 |
| ENSSSCG00000028304 | 7 | 74.10 | 82.79 | 120.30 | 92.40 | 92.37 | 61.52 | 61.51 | 71.80 |
| ENSSSCG00000006848 | 4 | 52.70 | 101.39 | 123.07 | 92.39 | 16.50 | 22.47 | 39.54 | 26.17 |
| ENSSSCG00000012604 | X | 78.25 | 100.82 | 97.57 | 92.21 | 76.60 | 105.51 | 104.31 | 95.47 |
| ENSSSCG00000025941 | 1 | 106.91 | 114.11 | 55.15 | 92.06 | 62.38 | 66.24 | 77.36 | 68.66 |
| ENSSSCG00000037102 | 2 | 83.49 | 132.23 | 60.00 | 91.91 | 19.89 | 20.10 | 28.68 | 22.89 |
| ENSSSCG00000016077 | 15 | 112.59 | 82.26 | 80.68 | 91.84 | 99.83 | 59.40 | 63.02 | 74.08 |
| ENSSSCG00000012840 | 2 | 78.28 | 77.89 | 118.65 | 91.61 | 54.13 | 37.75 | 77.70 | 56.53 |
| ENSSSCG00000004142 | 1 | 75.89 | 105.11 | 93.61 | 91.54 | 69.52 | 61.70 | 45.92 | 59.05 |
| ENSSSCG00000024550 | 14 | 89.34 | 77.86 | 106.80 | 91.33 | 86.03 | 69.18 | 93.00 | 82.74 |
| ENSSSCG00000029352 | 13 | 80.62 | 137.09 | 55.35 | 91.02 | 57.99 | 54.50 | 30.35 | 47.61 |
| ENSSSCG00000006335 | 4 | 79.46 | 182.43 | 11.12 | 91.00 | 6.33 | 14.99 | 9.74 | 10.35 |
| ENSSSCG00000001571 | 7 | 86.63 | 100.93 | 85.34 | 90.97 | 82.09 | 77.45 | 88.58 | 82.71 |
| ENSSSCG00000027331 | 15 | 62.07 | 128.18 | 82.60 | 90.95 | 28.02 | 62.60 | 79.36 | 56.66 |
| ENSSSCG00000028431 | 2 | 47.36 | 82.53 | 142.46 | 90.78 | 28.89 | 36.97 | 43.82 | 36.56 |
| ENSSSCG00000014435 | 2 | 100.89 | 107.53 | 63.78 | 90.73 | 71.30 | 62.58 | 69.36 | 67.74 |
| ENSSSCG00000016018 | 15 | 72.65 | 89.13 | 109.84 | 90.54 | 4.73 | 34.72 | 32.18 | 23.87 |
| ENSSSCG00000011565 | 13 | 79.69 | 75.50 | 116.42 | 90.54 | 73.03 | 77.49 | 109.63 | 86.72 |
| ENSSSCG00000027257 | 1 | 95.77 | 81.86 | 93.52 | 90.38 | 85.00 | 72.09 | 93.63 | 83.58 |
| ENSSSCG00000012026 | 13 | 30.27 | 199.96 | 40.66 | 90.30 | 24.90 | 19.62 | 32.46 | 25.66 |
| ENSSSCG00000030125 | 4 | 92.95 | 88.10 | 89.71 | 90.25 | 72.25 | 69.95 | 103.65 | 81.95 |
| ENSSSCG00000012440 | X | 78.38 | 77.04 | 114.97 | 90.13 | 64.98 | 69.04 | 82.25 | 72.09 |
| ENSSSCG00000030798 | 3 | 137.91 | 76.29 | 55.95 | 90.05 | 104.16 | 68.15 | 57.64 | 76.65 |
| ENSSSCG00000025483 | 3 | 96.42 | 73.89 | 97.84 | 89.38 | 12.62 | 32.88 | 42.90 | 29.47 |
| ENSSSCG00000016370 | 15 | 93.79 | 89.71 | 84.01 | 89.17 | 80.75 | 72.51 | 81.92 | 78.39 |
| ENSSSCG00000011676 | 13 | 68.45 | 118.58 | 80.41 | 89.15 | 77.34 | 59.64 | 92.85 | 76.61 |
| ENSSSCG00000002050 | 7 | 91.72 | 70.14 | 105.54 | 89.13 | 77.18 | 68.79 | 113.44 | 86.47 |
| ENSSSCG00000023522 | 17 | 55.16 | 81.88 | 130.28 | 89.11 | 19.23 | 31.38 | 55.75 | 35.45 |
| ENSSSCG00000002023 | 7 | 114.02 | 66.97 | 86.15 | 89.05 | 77.46 | 93.07 | 73.88 | 81.47 |
| ENSSSCG00000023105 | 10 | 78.07 | 157.54 | 31.39 | 89.00 | 39.16 | 52.68 | 32.63 | 41.49 |
| ENSSSCG00000012766 | X | 91.67 | 82.22 | 93.10 | 88.99 | 34.69 | 48.46 | 83.91 | 55.68 |
| ENSSSCG00000011893 | 13 | 114.70 | 77.11 | 74.29 | 88.70 | 78.10 | 63.13 | 104.02 | 81.75 |
| ENSSSCG00000008557 | 3 | 127.61 | 74.88 | 62.66 | 88.38 | 56.10 | 52.45 | 51.19 | 53.25 |
| ENSSSCG00000013318 | 2 | 94.50 | 87.39 | 83.09 | 88.33 | 43.27 | 62.94 | 97.18 | 67.80 |
| ENSSSCG00000010437 | 14 | 72.31 | 93.55 | 98.72 | 88.19 | 2.03 | 33.14 | 23.13 | 19.43 |
| ENSSSCG00000007435 | 17 | 55.58 | 82.30 | 126.68 | 88.19 | 23.44 | 54.48 | 29.58 | 35.83 |
| ENSSSCG00000027072 | 6 | 84.21 | 73.93 | 106.30 | 88.15 | 76.37 | 69.44 | 83.83 | 76.54 |
| ENSSSCG00000032166 | 12 | 96.71 | 79.48 | 87.39 | 87.86 | 82.91 | 79.97 | 103.65 | 88.84 |
| ENSSSCG00000014036 | 2 | 96.26 | 78.58 | 88.60 | 87.81 | 59.11 | 64.01 | 91.51 | 71.54 |
| ENSSSCG00000030857 | 6 | 92.98 | 93.29 | 76.47 | 87.58 | 57.84 | 82.18 | 45.23 | 61.75 |

| | | | | | | | | | |
|---|---|---|---|---|---|---|---|---|---|
| ENSSSCG00000008701 | 8 | 82.51 | 88.80 | 90.12 | 87.14 | 68.12 | 66.70 | 92.13 | 75.65 |
| ENSSSCG00000037821 | 4 | 124.61 | 68.96 | 67.79 | 87.12 | 28.27 | 78.11 | 36.38 | 47.59 |
| ENSSSCG00000030268 | 1 | 91.35 | 71.63 | 98.08 | 87.02 | 45.58 | 56.36 | 47.22 | 49.72 |
| ENSSSCG00000040710 | 1 | 95.26 | 82.67 | 80.53 | 86.15 | 59.44 | 53.45 | 89.64 | 67.51 |
| ENSSSCG00000000810 | 5 | 70.49 | 64.30 | 122.54 | 85.78 | 11.33 | 12.83 | 46.97 | 23.71 |
| ENSSSCG00000017065 | 16 | 40.42 | 131.26 | 83.98 | 85.22 | 29.34 | 38.55 | 54.19 | 40.69 |
| ENSSSCG00000006487 | 4 | 105.19 | 80.75 | 69.18 | 85.04 | 71.86 | 59.41 | 76.44 | 69.24 |
| ENSSSCG00000039740 | 2 | 86.05 | 88.15 | 80.82 | 85.00 | 85.26 | 82.89 | 91.23 | 86.46 |
| ENSSSCG00000022925 | 5 | 76.72 | 153.44 | 24.55 | 84.90 | 19.32 | 15.38 | 18.07 | 17.59 |
| ENSSSCG00000025109 | 6 | 70.62 | 95.75 | 87.45 | 84.61 | 46.67 | 75.26 | 69.65 | 63.86 |
| ENSSSCG00000000406 | 5 | 107.12 | 80.66 | 65.96 | 84.58 | 143.27 | 97.89 | 107.16 | 116.11 |
| ENSSSCG00000029066 | 10 | 158.86 | 22.40 | 72.17 | 84.47 | 23.62 | 24.81 | 46.28 | 31.57 |
| ENSSSCG00000026414 | 16 | 87.69 | 77.52 | 88.05 | 84.42 | 59.31 | 78.67 | 132.54 | 90.17 |
| ENSSSCG00000014136 | 2 | 87.54 | 66.18 | 98.70 | 84.14 | 8.00 | 31.53 | 34.70 | 24.74 |
| ENSSSCG00000013863 | 2 | 77.31 | 82.80 | 92.17 | 84.09 | 106.96 | 96.49 | 105.55 | 103.00 |
| ENSSSCG00000002444 | 7 | 21.05 | 149.61 | 81.44 | 84.04 | 15.78 | 28.15 | 33.04 | 25.66 |
| ENSSSCG00000008247 | 3 | 79.61 | 76.38 | 95.04 | 83.68 | 89.11 | 78.69 | 108.45 | 92.08 |
| ENSSSCG00000009249 | 8 | 111.03 | 66.69 | 73.21 | 83.64 | 83.17 | 78.70 | 77.81 | 79.90 |
| ENSSSCG00000014850 | 9 | 86.46 | 76.89 | 85.97 | 83.10 | 78.87 | 71.05 | 91.53 | 80.48 |
| ENSSSCG00000039847 | 5 | 52.16 | 90.46 | 106.54 | 83.06 | 80.09 | 81.42 | 68.08 | 76.53 |
| ENSSSCG00000040012 | 6 | 95.29 | 70.80 | 82.66 | 82.92 | 63.90 | 54.65 | 74.07 | 64.20 |
| ENSSSCG00000010238 | 14 | 82.80 | 86.06 | 79.72 | 82.86 | 81.83 | 77.13 | 59.53 | 72.83 |
| ENSSSCG00000000058 | 5 | 95.20 | 85.87 | 67.38 | 82.82 | 84.05 | 78.20 | 83.71 | 81.99 |
| ENSSSCG00000032028 | 5 | 84.14 | 80.37 | 83.86 | 82.79 | 92.54 | 104.70 | 97.82 | 98.36 |
| ENSSSCG00000000492 | 5 | 17.55 | 29.88 | 200.43 | 82.62 | 16.75 | 9.74 | 13.64 | 13.37 |
| ENSSSCG00000003354 | 6 | 88.73 | 67.60 | 91.49 | 82.61 | 71.47 | 73.15 | 53.59 | 66.07 |
| ENSSSCG00000004870 | 1 | 47.30 | 56.84 | 142.75 | 82.30 | 40.29 | 43.66 | 73.72 | 52.56 |
| ENSSSCG00000014577 | 9 | 75.14 | 86.47 | 84.96 | 82.19 | 42.95 | 49.82 | 69.12 | 53.96 |
| ENSSSCG00000013366 | 2 | 75.00 | 93.09 | 77.44 | 81.85 | 71.04 | 65.76 | 84.09 | 73.63 |
| ENSSSCG00000031808 | 11 | 92.90 | 80.73 | 71.85 | 81.83 | 92.53 | 80.75 | 102.91 | 92.06 |
| ENSSSCG00000022820 | 3 | 74.08 | 83.99 | 86.83 | 81.64 | 134.55 | 112.58 | 175.12 | 140.75 |
| ENSSSCG00000020962 | 12 | 75.37 | 97.94 | 71.39 | 81.57 | 105.38 | 104.79 | 126.37 | 112.18 |
| ENSSSCG00000009821 | 14 | 78.79 | 79.30 | 86.48 | 81.52 | 60.69 | 75.14 | 94.32 | 76.72 |
| ENSSSCG00000033500 | 6 | 81.47 | 83.85 | 79.07 | 81.46 | 50.70 | 74.79 | 83.50 | 69.66 |
| ENSSSCG00000022447 | 4 | 127.40 | 76.63 | 40.09 | 81.37 | 13.72 | 20.68 | 21.14 | 18.52 |
| ENSSSCG00000031577 | 1 | 78.92 | 81.78 | 83.10 | 81.27 | 72.56 | 96.95 | 96.83 | 88.78 |
| ENSSSCG00000003514 | 6 | 78.52 | 81.17 | 83.85 | 81.18 | 35.48 | 49.53 | 48.12 | 44.38 |
| ENSSSCG00000011621 | 13 | 80.60 | 81.71 | 81.09 | 81.14 | 76.55 | 55.08 | 87.14 | 72.92 |
| ENSSSCG00000022236 | 9 | 49.47 | 98.20 | 95.53 | 81.07 | 86.88 | 111.90 | 157.49 | 118.75 |
| ENSSSCG00000023277 | X | 88.83 | 68.05 | 86.08 | 80.99 | 89.01 | 73.17 | 94.05 | 85.41 |
| ENSSSCG00000013029 | 2 | 89.27 | 77.29 | 76.33 | 80.96 | 85.29 | 67.58 | 75.31 | 76.06 |
| ENSSSCG00000038991 | 4 | 110.91 | 56.82 | 74.47 | 80.73 | 63.57 | 111.64 | 80.27 | 85.16 |
| ENSSSCG00000003261 | 6 | 86.92 | 71.78 | 83.33 | 80.68 | 101.30 | 68.32 | 104.98 | 91.53 |
| ENSSSCG00000003150 | 6 | 77.96 | 79.88 | 83.67 | 80.50 | 60.37 | 64.38 | 95.19 | 73.31 |
| ENSSSCG00000023604 | 8 | 68.39 | 98.57 | 74.12 | 80.36 | 45.66 | 53.98 | 57.23 | 52.29 |
| ENSSSCG00000010450 | 14 | 66.05 | 86.42 | 88.08 | 80.18 | 120.94 | 83.52 | 93.80 | 99.42 |
| ENSSSCG00000024823 | 6 | 97.77 | 46.74 | 95.63 | 80.04 | 38.22 | 41.65 | 57.64 | 45.84 |
| ENSSSCG00000038514 | 7 | 85.29 | 73.03 | 81.23 | 79.85 | 81.44 | 97.85 | 105.58 | 94.96 |
| ENSSSCG00000023380 | 14 | 78.22 | 75.66 | 85.29 | 79.72 | 82.15 | 67.86 | 79.54 | 76.52 |
| ENSSSCG00000000444 | 5 | 75.97 | 76.03 | 87.16 | 79.72 | 71.79 | 85.89 | 98.53 | 85.40 |
| ENSSSCG00000013302 | 2 | 40.75 | 80.17 | 118.24 | 79.72 | 60.86 | 65.72 | 46.27 | 57.61 |
| ENSSSCG00000002307 | 7 | 96.20 | 69.16 | 73.47 | 79.61 | 73.73 | 66.10 | 78.04 | 72.62 |
| ENSSSCG00000014184 | 2 | 79.31 | 73.21 | 86.31 | 79.61 | 33.52 | 53.90 | 57.65 | 48.35 |
| ENSSSCG00000008633 | 3 | 92.69 | 63.53 | 82.21 | 79.48 | 45.27 | 48.79 | 85.07 | 59.71 |
| ENSSSCG00000016186 | 15 | 68.54 | 99.72 | 69.93 | 79.39 | 62.09 | 54.88 | 64.35 | 60.44 |
| ENSSSCG00000005300 | 1 | 81.46 | 71.00 | 85.40 | 79.29 | 75.09 | 62.32 | 95.10 | 77.51 |

| | | |
|---|---|---|
| A≧0.1 | 15609 | |
| Total | 16219 | |
| | | 16455 |
| B≧0.1 | 15906 | |
| Total | 16285 | |

| FPKM | A | B | Overlap |
|---|---|---|---|
| ≧1000 | 22 | 22 | 18 |
| 500-1000 | 54 | 50 | 39 |
| 100-500 | 345 | 300 | |
| 50-100 | 543 | 487 | |
| 10-50 | 4433 | 4882 | |
| 1--10 | 6621 | 6996 | |
| 0.1-1 | 3589 | 3165 | |
| | 15607 | 15902 | |

### Table S2 Differecially expressed genes in ovaries between XL and XS groups

| data...1. | XL13 fpkm | XL17 fpkm | XL18 fpkm | XS12 fpkm | XS15 fpkm | XS16 fpkm | XL AveFPKM | XS-AveFPKM | FC | logFC | AveExpr | t | P.Value | adj.P.Val | B |
|---|---|---|---|---|---|---|---|---|---|---|---|---|---|---|---|
| ENSSSCG00000017915 | 3.82 | 4.58 | 4.55 | 16.20 | 16.65 | 16.30 | 4.32 | 16.38 | 0.26 | -1.92 | 10.35 | 41.23 | 0.0000 | 0.0031 | 5.75 |
| ENSSSCG00000017994 | 4.41 | 0.86 | 0.75 | 46.28 | 43.37 | 46.83 | 2.01 | 45.49 | 0.04 | -4.50 | 23.75 | 29.03 | 0.0000 | 0.0062 | 5.13 |
| ENSSSCG00000010298 | 2.45 | 0.71 | 1.06 | 21.51 | 21.08 | 22.94 | 1.41 | 21.84 | 0.06 | -3.96 | 11.62 | 28.00 | 0.0000 | 0.0062 | 5.05 |
| ENSSSCG00000038825 | 3.00 | 4.28 | 4.19 | 21.31 | 20.00 | 20.97 | 3.82 | 20.76 | 0.18 | -2.44 | 12.29 | 31.11 | 0.0000 | 0.0062 | 5.28 |
| ENSSSCG00000006294 | 8.27 | 8.37 | 7.69 | 25.89 | 26.95 | 24.78 | 8.11 | 25.87 | 0.31 | -1.67 | 16.99 | 28.30 | 0.0000 | 0.0062 | 5.07 |
| ENSSSCG00000004147 | 1.27 | 0.09 | 0.12 | 18.35 | 16.93 | 16.46 | 0.49 | 17.25 | 0.03 | -5.13 | 8.87 | 25.69 | 0.0000 | 0.0074 | 4.84 |
| ENSSSCG00000040973 | 1.85 | 0.25 | 0.21 | 26.08 | 25.32 | 28.61 | 0.77 | 26.67 | 0.03 | -5.11 | 13.72 | 24.53 | 0.0000 | 0.0074 | 4.72 |
| ENSSSCG00000004413 | 3.66 | 0.15 | 0.63 | 36.66 | 36.60 | 40.17 | 1.48 | 37.81 | 0.04 | -4.67 | 19.64 | 24.20 | 0.0000 | 0.0074 | 4.68 |
| ENSSSCG00000004602 | 1.81 | 1.15 | 1.37 | 10.53 | 9.78 | 9.56 | 1.45 | 9.95 | 0.15 | -2.78 | 5.70 | 24.15 | 0.0000 | 0.0074 | 4.67 |
| ENSSSCG00000009388 | 1.72 | 0.33 | 0.81 | 20.31 | 18.38 | 21.32 | 0.96 | 20.00 | 0.05 | -4.39 | 10.48 | 21.36 | 0.0000 | 0.0075 | 4.32 |
| ENSSSCG00000016542 | 0.48 | 0.17 | 0.10 | 4.43 | 4.73 | 4.68 | 0.25 | 4.61 | 0.05 | -4.22 | 2.43 | 22.73 | 0.0000 | 0.0075 | 4.50 |
| ENSSSCG00000031086 | 3.22 | 0.53 | 1.19 | 25.65 | 24.40 | 22.77 | 1.65 | 24.27 | 0.07 | -3.88 | 12.96 | 20.87 | 0.0000 | 0.0075 | 4.25 |
| ENSSSCG00000037461 | 10.66 | 4.27 | 4.84 | 85.14 | 82.60 | 94.13 | 6.59 | 87.29 | 0.08 | -3.73 | 46.94 | 21.50 | 0.0000 | 0.0075 | 4.34 |
| ENSSSCG00000015031 | 2.21 | 0.77 | 0.95 | 16.49 | 15.72 | 14.52 | 1.31 | 15.57 | 0.08 | -3.57 | 8.44 | 20.71 | 0.0000 | 0.0075 | 4.22 |
| ENSSSCG00000026689 | 4.19 | 1.16 | 2.24 | 27.94 | 31.67 | 30.15 | 2.53 | 29.92 | 0.08 | -3.56 | 16.23 | 21.01 | 0.0000 | 0.0075 | 4.27 |
| ENSSSCG00000033546 | 6.84 | 1.26 | 2.64 | 40.28 | 40.41 | 42.84 | 3.58 | 41.17 | 0.09 | -3.52 | 22.38 | 21.60 | 0.0000 | 0.0075 | 4.35 |
| ENSSSCG00000032929 | 6.07 | 1.81 | 2.39 | 33.11 | 33.24 | 31.92 | 3.42 | 32.76 | 0.10 | -3.26 | 18.09 | 22.53 | 0.0000 | 0.0075 | 4.48 |
| ENSSSCG00000015794 | 2.50 | 1.11 | 1.85 | 17.00 | 16.62 | 18.53 | 1.82 | 17.38 | 0.10 | -3.26 | 9.60 | 23.25 | 0.0000 | 0.0075 | 4.57 |
| ENSSSCG00000034398 | 3.46 | 3.17 | 3.55 | 9.94 | 10.04 | 9.26 | 3.39 | 9.75 | 0.35 | -1.52 | 6.57 | 22.31 | 0.0000 | 0.0075 | 4.45 |
| ENSSSCG00000002817 | 1.44 | 0.11 | 0.18 | 24.21 | 22.04 | 26.64 | 0.58 | 24.30 | 0.02 | -5.39 | 12.44 | 18.21 | 0.0000 | 0.0080 | 3.80 |
| ENSSSCG00000033394 | 0.31 | 0.02 | 0.02 | 4.42 | 4.11 | 4.80 | 0.12 | 4.45 | 0.03 | -5.20 | 2.28 | 17.67 | 0.0000 | 0.0080 | 3.69 |
| ENSSSCG00000002504 | 3.62 | 0.31 | 0.78 | 54.86 | 46.89 | 56.34 | 1.57 | 52.69 | 0.03 | -5.07 | 27.13 | 17.73 | 0.0000 | 0.0080 | 3.71 |
| ENSSSCG00000000982 | 0.26 | 0.07 | 0.00 | 3.24 | 3.58 | 3.29 | 0.11 | 3.37 | 0.03 | -4.95 | 1.74 | 18.01 | 0.0000 | 0.0080 | 3.76 |
| ENSSSCG00000032827 | 1.06 | 0.07 | 0.13 | 13.15 | 11.51 | 13.43 | 0.42 | 12.70 | 0.03 | -4.92 | 6.56 | 19.04 | 0.0000 | 0.0080 | 3.95 |
| ENSSSCG00000040950 | 0.43 | 0.00 | 0.12 | 4.75 | 5.25 | 4.74 | 0.18 | 4.92 | 0.04 | -4.74 | 2.55 | 19.98 | 0.0000 | 0.0080 | 4.11 |
| ENSSSCG00000011855 | 0.84 | 0.02 | 0.20 | 8.34 | 8.52 | 9.54 | 0.35 | 8.80 | 0.04 | -4.65 | 4.57 | 19.38 | 0.0000 | 0.0080 | 4.01 |
| ENSSSCG00000011069 | 4.70 | 0.39 | 0.62 | 50.35 | 43.86 | 43.06 | 1.90 | 45.76 | 0.04 | -4.59 | 23.83 | 17.51 | 0.0000 | 0.0080 | 3.66 |
| ENSSSCG00000023393 | 1.11 | 0.13 | 0.16 | 11.23 | 10.61 | 9.64 | 0.47 | 10.49 | 0.04 | -4.49 | 5.48 | 18.63 | 0.0000 | 0.0080 | 3.88 |
| ENSSSCG00000011737 | 1.00 | 0.15 | 0.04 | 7.90 | 7.93 | 7.24 | 0.40 | 7.69 | 0.05 | -4.27 | 4.05 | 19.51 | 0.0000 | 0.0080 | 4.03 |

| | | | | | | | | | | | | | | | |
|---|---|---|---|---|---|---|---|---|---|---|---|---|---|---|---|
| ENSSSCG00000012558 | 2.10 | 0.06 | 0.19 | 14.43 | 14.55 | 15.94 | 0.78 | 14.97 | 0.05 | -4.26 | 7.88 | 18.45 | 0.0000 | 0.0080 | 3.84 |
| ENSSSCG00000011892 | 1.56 | 0.24 | 0.29 | 11.21 | 12.84 | 12.86 | 0.70 | 12.30 | 0.06 | -4.14 | 6.50 | 17.62 | 0.0000 | 0.0080 | 3.68 |
| ENSSSCG00000033381 | 0.75 | 0.42 | 0.26 | 7.97 | 7.90 | 9.16 | 0.48 | 8.34 | 0.06 | -4.13 | 4.41 | 18.62 | 0.0000 | 0.0080 | 3.88 |
| ENSSSCG00000006069 | 1.11 | 0.42 | 0.77 | 8.48 | 8.34 | 7.43 | 0.77 | 8.08 | 0.09 | -3.40 | 4.42 | 19.27 | 0.0000 | 0.0080 | 3.99 |
| ENSSSCG00000030513 | 1.31 | 0.47 | 0.75 | 9.42 | 8.18 | 8.47 | 0.84 | 8.69 | 0.10 | -3.37 | 4.77 | 17.95 | 0.0000 | 0.0080 | 3.75 |
| ENSSSCG00000028682 | 6.34 | 1.58 | 2.42 | 36.84 | 33.46 | 34.57 | 3.45 | 34.95 | 0.10 | -3.34 | 19.20 | 19.11 | 0.0000 | 0.0080 | 3.96 |
| ENSSSCG00000014196 | 0.80 | 0.32 | 0.52 | 4.08 | 4.36 | 4.16 | 0.55 | 4.20 | 0.13 | -2.94 | 2.37 | 18.12 | 0.0000 | 0.0080 | 3.78 |
| ENSSSCG00000007094 | 9.48 | 6.78 | 11.45 | 40.65 | 36.88 | 39.11 | 9.23 | 38.88 | 0.24 | -2.07 | 24.06 | 18.32 | 0.0000 | 0.0080 | 3.82 |
| ENSSSCG00000025417 | 7.89 | 6.05 | 8.83 | 28.04 | 27.97 | 30.99 | 7.59 | 29.00 | 0.26 | -1.93 | 18.29 | 17.86 | 0.0000 | 0.0080 | 3.73 |
| ENSSSCG00000010737 | 8.34 | 5.67 | 6.06 | 23.04 | 22.44 | 24.40 | 6.69 | 23.29 | 0.29 | -1.80 | 14.99 | 17.49 | 0.0000 | 0.0080 | 3.66 |
| ENSSSCG00000038080 | 9.03 | 8.29 | 9.17 | 26.75 | 27.16 | 30.11 | 8.83 | 28.01 | 0.32 | -1.67 | 18.42 | 18.76 | 0.0000 | 0.0080 | 3.90 |
| ENSSSCG00000024357 | 3.20 | 0.17 | 0.42 | 28.03 | 23.97 | 26.87 | 1.26 | 26.29 | 0.05 | -4.38 | 13.78 | 17.33 | 0.0000 | 0.0081 | 3.62 |
| ENSSSCG00000010768 | 3.24 | 0.79 | 1.00 | 22.16 | 19.17 | 21.45 | 1.67 | 20.92 | 0.08 | -3.64 | 11.30 | 17.27 | 0.0000 | 0.0081 | 3.61 |
| ENSSSCG00000033624 | 0.62 | 0.54 | 0.50 | 3.83 | 3.41 | 3.62 | 0.55 | 3.62 | 0.15 | -2.71 | 2.09 | 17.18 | 0.0000 | 0.0081 | 3.59 |
| ENSSSCG00000003336 | 2.10 | 0.07 | 0.22 | 16.35 | 19.35 | 17.64 | 0.80 | 17.78 | 0.04 | -4.48 | 9.29 | 16.70 | 0.0000 | 0.0085 | 3.49 |
| ENSSSCG00000027467 | 2.59 | 1.61 | 1.92 | 11.22 | 10.29 | 12.00 | 2.04 | 11.17 | 0.18 | -2.45 | 6.61 | 16.66 | 0.0000 | 0.0085 | 3.48 |
| ENSSSCG00000016133 | 0.94 | 0.03 | 0.16 | 11.83 | 9.94 | 12.04 | 0.38 | 11.27 | 0.03 | -4.90 | 5.82 | 15.86 | 0.0000 | 0.0085 | 3.30 |
| ENSSSCG00000017610 | 0.62 | 0.00 | 0.04 | 5.89 | 6.06 | 5.15 | 0.22 | 5.70 | 0.04 | -4.69 | 2.96 | 15.93 | 0.0000 | 0.0085 | 3.31 |
| ENSSSCG00000022305 | 2.40 | 0.29 | 0.78 | 25.29 | 30.27 | 25.72 | 1.16 | 27.10 | 0.04 | -4.55 | 14.13 | 16.26 | 0.0000 | 0.0085 | 3.39 |
| ENSSSCG00000040019 | 0.26 | 0.08 | 0.09 | 3.13 | 3.17 | 3.59 | 0.14 | 3.30 | 0.04 | -4.53 | 1.72 | 16.11 | 0.0000 | 0.0085 | 3.36 |
| ENSSSCG00000028996 | 9.32 | 2.72 | 3.61 | 66.03 | 72.56 | 79.70 | 5.22 | 72.76 | 0.07 | -3.80 | 38.99 | 16.35 | 0.0000 | 0.0085 | 3.41 |
| ENSSSCG00000008745 | 0.66 | 0.92 | 0.70 | 6.45 | 6.33 | 7.51 | 0.76 | 6.76 | 0.11 | -3.15 | 3.76 | 15.90 | 0.0000 | 0.0085 | 3.31 |
| ENSSSCG00000004608 | 0.52 | 0.34 | 0.52 | 3.43 | 3.63 | 3.21 | 0.46 | 3.42 | 0.13 | -2.90 | 1.94 | 16.21 | 0.0000 | 0.0085 | 3.38 |
| ENSSSCG00000026473 | 0.50 | 0.78 | 1.19 | 5.22 | 5.24 | 4.81 | 0.82 | 5.09 | 0.16 | -2.63 | 2.96 | 16.13 | 0.0000 | 0.0085 | 3.36 |
| ENSSSCG00000002412 | 1.26 | 1.11 | 1.30 | 4.95 | 5.24 | 4.59 | 1.23 | 4.93 | 0.25 | -2.01 | 3.08 | 16.42 | 0.0000 | 0.0085 | 3.43 |
| ENSSSCG00000011090 | 5.17 | 4.63 | 3.55 | 13.81 | 12.69 | 13.50 | 4.45 | 13.33 | 0.33 | -1.58 | 8.89 | 16.01 | 0.0000 | 0.0085 | 3.33 |
| ENSSSCG00000022512 | 0.54 | 0.57 | 0.91 | 4.22 | 4.32 | 3.84 | 0.67 | 4.13 | 0.16 | -2.62 | 2.40 | 15.67 | 0.0000 | 0.0087 | 3.25 |
| ENSSSCG00000004856 | 3.90 | 4.13 | 3.63 | 9.08 | 9.29 | 8.36 | 3.89 | 8.91 | 0.44 | -1.20 | 6.40 | 15.64 | 0.0000 | 0.0087 | 3.24 |
| ENSSSCG00000008072 | 0.69 | 1.33 | 1.19 | 10.80 | 9.36 | 9.00 | 1.07 | 9.72 | 0.11 | -3.18 | 5.40 | 15.51 | 0.0000 | 0.0089 | 3.21 |
| ENSSSCG00000004415 | 1.39 | 0.20 | 0.43 | 13.66 | 15.99 | 16.98 | 0.67 | 15.54 | 0.04 | -4.53 | 8.11 | 15.16 | 0.0000 | 0.0091 | 3.12 |
| ENSSSCG00000040714 | 0.36 | 0.21 | 0.14 | 3.18 | 2.78 | 3.07 | 0.24 | 3.01 | 0.08 | -3.66 | 1.62 | 15.18 | 0.0000 | 0.0091 | 3.13 |
| ENSSSCG00000016365 | 1.52 | 0.40 | 0.29 | 7.64 | 7.04 | 7.76 | 0.74 | 7.48 | 0.10 | -3.34 | 4.11 | 15.34 | 0.0000 | 0.0091 | 3.17 |
| ENSSSCG00000004469 | 2.51 | 0.65 | 1.12 | 11.05 | 11.23 | 10.17 | 1.43 | 10.82 | 0.13 | -2.92 | 6.12 | 15.26 | 0.0000 | 0.0091 | 3.15 |
| ENSSSCG00000033268 | 1.03 | 0.80 | 0.91 | 3.21 | 3.21 | 3.11 | 0.91 | 3.18 | 0.29 | -1.80 | 2.04 | 15.02 | 0.0000 | 0.0093 | 3.09 |
| ENSSSCG00000022739 | 6.47 | 2.42 | 2.51 | 26.66 | 24.53 | 24.23 | 3.80 | 25.14 | 0.15 | -2.73 | 14.47 | 14.88 | 0.0000 | 0.0094 | 3.05 |

| | | | | | | | | | | | | | | | |
|---|---|---|---|---|---|---|---|---|---|---|---|---|---|---|---|
| ENSSSCG00000036261 | 1.88 | 0.28 | 0.28 | 17.01 | 13.93 | 14.92 | 0.81 | 15.29 | 0.05 | -4.24 | 8.05 | 14.69 | 0.0000 | 0.0096 | 3.00 |
| ENSSSCG00000017098 | 8.49 | 3.28 | 3.30 | 42.92 | 43.55 | 50.43 | 5.03 | 45.63 | 0.11 | -3.18 | 25.33 | 14.77 | 0.0000 | 0.0096 | 3.02 |
| ENSSSCG00000034942 | 111.90 | 104.71 | 87.37 | 0.50 | 2.11 | 2.33 | 101.33 | 1.65 | 61.40 | 5.94 | 51.49 | -14.73 | 0.0000 | 0.0096 | 3.01 |
| ENSSSCG00000016030 | 0.23 | 0.00 | 0.07 | 5.40 | 6.73 | 6.71 | 0.10 | 6.28 | 0.02 | -5.96 | 3.19 | 14.27 | 0.0001 | 0.0108 | 2.89 |
| ENSSSCG00000005588 | 33.93 | 19.31 | 21.88 | 5.09 | 5.64 | 5.35 | 25.04 | 5.36 | 4.67 | 2.22 | 15.20 | -3.90 | 0.0030 | 0.0108 | -2.95 |
| ENSSSCG00000010100 | 0.36 | 0.10 | 0.03 | 5.93 | 4.70 | 5.42 | 0.16 | 5.35 | 0.03 | -5.06 | 2.76 | 14.17 | 0.0001 | 0.0109 | 2.86 |
| ENSSSCG00000035668 | 0.99 | 0.10 | 0.12 | 8.02 | 8.92 | 9.99 | 0.40 | 8.98 | 0.04 | -4.48 | 4.69 | 14.10 | 0.0001 | 0.0109 | 2.84 |
| ENSSSCG00000016215 | 5.10 | 3.46 | 0.61 | 59.63 | 47.01 | 53.90 | 3.06 | 53.51 | 0.06 | -4.13 | 28.29 | 14.04 | 0.0001 | 0.0109 | 2.82 |
| ENSSSCG00000005272 | 1.96 | 2.23 | 2.61 | 5.57 | 5.42 | 5.75 | 2.27 | 5.58 | 0.41 | -1.30 | 3.92 | 14.06 | 0.0001 | 0.0109 | 2.82 |
| ENSSSCG00000005425 | 20.67 | 19.37 | 22.65 | 50.18 | 48.37 | 55.47 | 20.90 | 51.34 | 0.41 | -1.30 | 36.12 | 14.04 | 0.0001 | 0.0109 | 2.82 |
| ENSSSCG00000016105 | 0.87 | 0.00 | 0.12 | 9.33 | 7.46 | 8.26 | 0.33 | 8.35 | 0.04 | -4.66 | 4.34 | 13.85 | 0.0001 | 0.0110 | 2.76 |
| ENSSSCG00000002356 | 0.39 | 0.00 | 0.02 | 2.84 | 3.27 | 2.99 | 0.14 | 3.03 | 0.05 | -4.47 | 1.58 | 13.68 | 0.0001 | 0.0110 | 2.71 |
| ENSSSCG00000017158 | 2.56 | 0.19 | 0.33 | 18.99 | 15.62 | 18.64 | 1.03 | 17.75 | 0.06 | -4.11 | 9.39 | 13.64 | 0.0001 | 0.0110 | 2.70 |
| ENSSSCG00000037723 | 2.41 | 0.13 | 0.00 | 15.52 | 13.09 | 14.40 | 0.85 | 14.34 | 0.06 | -4.08 | 7.59 | 13.71 | 0.0001 | 0.0110 | 2.72 |
| ENSSSCG00000033043 | 1.47 | 0.59 | 1.59 | 10.93 | 10.40 | 12.76 | 1.22 | 11.37 | 0.11 | -3.22 | 6.29 | 13.77 | 0.0001 | 0.0110 | 2.74 |
| ENSSSCG00000012970 | 1.86 | 1.96 | 2.01 | 13.22 | 12.19 | 10.59 | 1.94 | 12.00 | 0.16 | -2.63 | 6.97 | 13.94 | 0.0001 | 0.0110 | 2.79 |
| ENSSSCG00000036964 | 2.38 | 1.61 | 2.57 | 8.64 | 7.65 | 8.79 | 2.19 | 8.36 | 0.26 | -1.93 | 5.27 | 13.71 | 0.0001 | 0.0110 | 2.72 |
| ENSSSCG00000027169 | 2.84 | 3.24 | 3.50 | 7.58 | 7.96 | 8.59 | 3.19 | 8.04 | 0.40 | -1.33 | 5.62 | 13.79 | 0.0001 | 0.0110 | 2.75 |
| ENSSSCG00000013476 | 0.69 | 0.08 | 0.09 | 4.06 | 3.59 | 3.90 | 0.29 | 3.85 | 0.07 | -3.75 | 2.07 | 13.60 | 0.0001 | 0.0110 | 2.69 |
| ENSSSCG00000010384 | 1.95 | 0.00 | 0.00 | 10.55 | 10.69 | 12.15 | 0.65 | 11.13 | 0.06 | -4.10 | 5.89 | 13.49 | 0.0001 | 0.0113 | 2.66 |
| ENSSSCG00000003814 | 3.51 | 2.55 | 3.27 | 13.15 | 13.99 | 11.54 | 3.11 | 12.89 | 0.24 | -2.05 | 8.00 | 13.41 | 0.0001 | 0.0115 | 2.63 |
| ENSSSCG00000016255 | 2.07 | 0.03 | 0.23 | 17.64 | 14.15 | 17.00 | 0.78 | 16.26 | 0.05 | -4.39 | 8.52 | 13.26 | 0.0001 | 0.0119 | 2.58 |
| ENSSSCG00000008480 | 14.34 | 8.99 | 10.26 | 79.63 | 78.70 | 96.56 | 11.20 | 84.96 | 0.13 | -2.92 | 48.08 | 13.21 | 0.0001 | 0.0119 | 2.57 |
| ENSSSCG00000028122 | 0.55 | 0.10 | 0.12 | 4.26 | 3.54 | 4.20 | 0.26 | 4.00 | 0.06 | -3.97 | 2.13 | 13.15 | 0.0001 | 0.0120 | 2.55 |
| ENSSSCG00000025266 | 3.50 | 0.17 | 0.39 | 34.93 | 31.74 | 40.89 | 1.35 | 35.85 | 0.04 | -4.73 | 18.60 | 12.87 | 0.0001 | 0.0127 | 2.46 |
| ENSSSCG00000040418 | 0.44 | 0.16 | 0.18 | 2.93 | 3.49 | 3.04 | 0.26 | 3.15 | 0.08 | -3.58 | 1.71 | 12.94 | 0.0001 | 0.0127 | 2.48 |
| ENSSSCG00000039342 | 3.79 | 1.87 | 2.82 | 13.81 | 13.69 | 16.15 | 2.82 | 14.55 | 0.19 | -2.37 | 8.69 | 12.85 | 0.0001 | 0.0127 | 2.45 |
| ENSSSCG00000007774 | 0.79 | 0.52 | 1.10 | 3.51 | 3.50 | 3.72 | 0.80 | 3.58 | 0.22 | -2.15 | 2.19 | 12.90 | 0.0001 | 0.0127 | 2.47 |
| ENSSSCG00000011192 | 5.82 | 4.44 | 5.45 | 10.49 | 10.51 | 10.85 | 5.24 | 10.62 | 0.49 | -1.02 | 7.93 | 12.85 | 0.0001 | 0.0127 | 2.45 |
| ENSSSCG00000011738 | 0.54 | 0.00 | 0.02 | 5.85 | 4.62 | 5.53 | 0.19 | 5.33 | 0.04 | -4.82 | 2.76 | 12.78 | 0.0001 | 0.0127 | 2.43 |
| ENSSSCG00000037478 | 2.13 | 0.33 | 0.33 | 16.23 | 13.10 | 16.17 | 0.93 | 15.17 | 0.06 | -4.03 | 8.05 | 12.77 | 0.0001 | 0.0127 | 2.43 |
| ENSSSCG00000004302 | 1.95 | 0.08 | 0.15 | 10.86 | 9.55 | 11.45 | 0.72 | 10.62 | 0.07 | -3.87 | 5.67 | 12.68 | 0.0001 | 0.0128 | 2.40 |
| ENSSSCG00000011423 | 0.72 | 0.55 | 0.58 | 4.00 | 3.40 | 4.15 | 0.62 | 3.85 | 0.16 | -2.64 | 2.23 | 12.69 | 0.0001 | 0.0128 | 2.40 |
| ENSSSCG00000017884 | 3.18 | 0.19 | 0.33 | 48.05 | 39.11 | 51.60 | 1.23 | 46.25 | 0.03 | -5.23 | 23.74 | 12.64 | 0.0001 | 0.0128 | 2.38 |
| ENSSSCG00000038684 | 0.53 | 0.05 | 0.16 | 8.83 | 6.78 | 7.48 | 0.25 | 7.70 | 0.03 | -4.95 | 3.97 | 12.62 | 0.0001 | 0.0128 | 2.38 |

| | | | | | | | | | | | | | | |
|---|---|---|---|---|---|---|---|---|---|---|---|---|---|---|
| ENSSSCG00000035933 | 3.35 | 0.24 | 0.54 | 31.94 | 33.77 | 40.96 | 1.38 | 35.56 | 0.04 | -4.69 | 18.47 | 12.59 | 0.0001 | 0.0128 | 2.37 |
| ENSSSCG00000029778 | 1.41 | 0.30 | 0.44 | 6.45 | 6.49 | 7.56 | 0.72 | 6.83 | 0.10 | -3.25 | 3.77 | 12.60 | 0.0001 | 0.0128 | 2.37 |
| ENSSSCG00000009428 | 4.09 | 0.36 | 0.34 | 43.78 | 53.23 | 57.73 | 1.60 | 51.58 | 0.03 | -5.01 | 26.59 | 12.56 | 0.0001 | 0.0128 | 2.36 |
| ENSSSCG00000009824 | 0.66 | 0.41 | 0.60 | 2.96 | 3.51 | 3.17 | 0.56 | 3.21 | 0.17 | -2.53 | 1.88 | 12.49 | 0.0001 | 0.0130 | 2.33 |
| ENSSSCG00000011915 | 1.09 | 1.01 | 1.40 | 5.65 | 4.91 | 4.73 | 1.17 | 5.10 | 0.23 | -2.13 | 3.13 | 12.47 | 0.0001 | 0.0130 | 2.33 |
| ENSSSCG00000028501 | 4.11 | 2.67 | 3.87 | 18.63 | 18.09 | 22.29 | 3.55 | 19.67 | 0.18 | -2.47 | 11.61 | 12.43 | 0.0001 | 0.0131 | 2.31 |
| ENSSSCG00000008942 | 16.32 | 13.55 | 13.91 | 34.61 | 33.72 | 30.28 | 14.59 | 32.87 | 0.44 | -1.17 | 23.73 | 12.41 | 0.0001 | 0.0131 | 2.31 |
| ENSSSCG00000034843 | 1.49 | 1.16 | 0.96 | 7.03 | 7.05 | 8.58 | 1.20 | 7.56 | 0.16 | -2.65 | 4.38 | 12.35 | 0.0001 | 0.0132 | 2.28 |
| ENSSSCG00000011751 | 9.34 | 0.21 | 0.70 | 96.70 | 78.85 | 103.65 | 3.42 | 93.07 | 0.04 | -4.77 | 48.24 | 12.15 | 0.0001 | 0.0137 | 2.22 |
| ENSSSCG00000015281 | 1.03 | 0.42 | 0.45 | 6.12 | 4.88 | 5.59 | 0.64 | 5.53 | 0.11 | -3.12 | 3.08 | 12.20 | 0.0001 | 0.0137 | 2.23 |
| ENSSSCG00000033574 | 1.19 | 0.05 | 0.12 | 13.11 | 15.10 | 11.38 | 0.45 | 13.20 | 0.03 | -4.86 | 6.82 | 12.00 | 0.0001 | 0.0140 | 2.16 |
| ENSSSCG00000007857 | 9.07 | 0.43 | 0.79 | 99.75 | 93.42 | 75.48 | 3.43 | 89.55 | 0.04 | -4.71 | 46.49 | 11.92 | 0.0001 | 0.0140 | 2.13 |
| ENSSSCG00000001042 | 2.77 | 0.70 | 0.45 | 25.77 | 20.15 | 25.85 | 1.31 | 23.92 | 0.05 | -4.20 | 12.61 | 12.02 | 0.0001 | 0.0140 | 2.17 |
| ENSSSCG00000008378 | 2.52 | 1.01 | 1.58 | 9.87 | 9.68 | 8.28 | 1.70 | 9.28 | 0.18 | -2.44 | 5.49 | 11.98 | 0.0001 | 0.0140 | 2.15 |
| ENSSSCG00000011881 | 1.60 | 1.15 | 1.87 | 4.70 | 4.64 | 4.38 | 1.54 | 4.57 | 0.34 | -1.57 | 3.06 | 11.97 | 0.0001 | 0.0140 | 2.15 |
| ENSSSCG00000008004 | 4.91 | 4.01 | 4.92 | 9.89 | 11.28 | 11.14 | 4.61 | 10.77 | 0.43 | -1.22 | 7.69 | 11.98 | 0.0001 | 0.0140 | 2.15 |
| ENSSSCG00000036679 | 5.91 | 4.86 | 6.22 | 20.96 | 17.23 | 20.61 | 5.67 | 19.60 | 0.29 | -1.79 | 12.63 | 11.87 | 0.0001 | 0.0142 | 2.11 |
| ENSSSCG00000028184 | 0.91 | 0.12 | 0.21 | 10.65 | 11.29 | 13.96 | 0.41 | 11.97 | 0.03 | -4.86 | 6.19 | 11.82 | 0.0001 | 0.0142 | 2.10 |
| ENSSSCG00000038660 | 1.44 | 1.64 | 1.45 | 5.93 | 5.60 | 4.84 | 1.51 | 5.45 | 0.28 | -1.85 | 3.48 | 11.84 | 0.0001 | 0.0142 | 2.10 |
| ENSSSCG00000024800 | 3.13 | 0.91 | 1.16 | 15.64 | 14.38 | 18.13 | 1.73 | 16.05 | 0.11 | -3.21 | 8.89 | 11.76 | 0.0001 | 0.0143 | 2.08 |
| ENSSSCG00000002440 | 3.40 | 1.36 | 1.42 | 15.86 | 13.58 | 13.01 | 2.06 | 14.15 | 0.15 | -2.78 | 8.10 | 11.77 | 0.0001 | 0.0143 | 2.08 |
| ENSSSCG00000010764 | 1.59 | 0.63 | 0.23 | 5.98 | 5.56 | 6.27 | 0.82 | 5.93 | 0.14 | -2.86 | 3.38 | 11.67 | 0.0001 | 0.0147 | 2.04 |
| ENSSSCG00000031963 | 2.82 | 0.21 | 0.37 | 25.00 | 20.17 | 26.80 | 1.13 | 23.99 | 0.05 | -4.40 | 12.56 | 11.43 | 0.0001 | 0.0151 | 1.95 |
| ENSSSCG00000032421 | 0.75 | 0.08 | 0.06 | 6.60 | 6.70 | 5.22 | 0.30 | 6.17 | 0.05 | -4.38 | 3.23 | 11.57 | 0.0001 | 0.0151 | 2.00 |
| ENSSSCG00000002767 | 0.86 | 0.04 | 0.27 | 7.84 | 6.56 | 8.73 | 0.39 | 7.71 | 0.05 | -4.31 | 4.05 | 11.47 | 0.0001 | 0.0151 | 1.96 |
| ENSSSCG00000011205 | 0.38 | 0.01 | 0.01 | 2.40 | 2.16 | 2.39 | 0.14 | 2.31 | 0.06 | -4.08 | 1.23 | 11.51 | 0.0001 | 0.0151 | 1.98 |
| ENSSSCG00000021448 | 1.37 | 0.17 | 0.46 | 10.55 | 9.75 | 12.75 | 0.67 | 11.02 | 0.06 | -4.04 | 5.84 | 11.44 | 0.0001 | 0.0151 | 1.95 |
| ENSSSCG00000014233 | 1.26 | 0.46 | 0.28 | 10.80 | 8.91 | 11.87 | 0.67 | 10.53 | 0.06 | -3.98 | 5.60 | 11.47 | 0.0001 | 0.0151 | 1.97 |
| ENSSSCG00000013584 | 0.45 | 0.29 | 0.28 | 5.50 | 4.19 | 4.58 | 0.34 | 4.75 | 0.07 | -3.79 | 2.55 | 11.41 | 0.0001 | 0.0151 | 1.94 |
| ENSSSCG00000018015 | 2.95 | 0.94 | 1.70 | 21.19 | 16.12 | 18.88 | 1.87 | 18.73 | 0.10 | -3.33 | 10.30 | 11.50 | 0.0001 | 0.0151 | 1.98 |
| ENSSSCG00000015767 | 0.34 | 0.28 | 0.29 | 2.84 | 2.72 | 3.39 | 0.30 | 2.98 | 0.10 | -3.31 | 1.64 | 11.51 | 0.0001 | 0.0151 | 1.98 |
| ENSSSCG00000001095 | 8.55 | 8.40 | 7.30 | 18.22 | 20.48 | 21.89 | 8.09 | 20.20 | 0.40 | -1.32 | 14.14 | 11.37 | 0.0001 | 0.0152 | 1.93 |
| ENSSSCG00000001011 | 12.73 | 16.36 | 11.79 | 32.59 | 29.43 | 31.14 | 13.63 | 31.05 | 0.44 | -1.19 | 22.34 | 11.24 | 0.0001 | 0.0156 | 1.88 |
| ENSSSCG00000007958 | 3.18 | 2.58 | 3.22 | 5.87 | 6.46 | 6.35 | 2.99 | 6.23 | 0.48 | -1.06 | 4.61 | 11.23 | 0.0001 | 0.0156 | 1.87 |
| ENSSSCG00000034633 | 1.00 | 1.84 | 1.28 | 8.50 | 9.39 | 11.04 | 1.37 | 9.64 | 0.14 | -2.81 | 5.51 | 11.22 | 0.0002 | 0.0156 | 1.87 |

| | | | | | | | | | | | | | | |
|---|---|---|---|---|---|---|---|---|---|---|---|---|---|---|
| ENSSSCG00000038186 | 0.81 | 0.12 | 0.15 | 6.66 | 5.05 | 5.95 | 0.36 | 5.89 | 0.06 | -4.04 | 3.12 | 11.10 | 0.0002 | 0.0159 | 1.82 |
| ENSSSCG00000006950 | 3.84 | 1.63 | 1.95 | 34.90 | 25.84 | 32.47 | 2.47 | 31.07 | 0.08 | -3.65 | 16.77 | 11.04 | 0.0002 | 0.0159 | 1.79 |
| ENSSSCG00000011263 | 3.80 | 2.17 | 2.24 | 24.69 | 18.71 | 21.04 | 2.74 | 21.48 | 0.13 | -2.97 | 12.11 | 11.08 | 0.0002 | 0.0159 | 1.81 |
| ENSSSCG00000013018 | 1.68 | 1.07 | 1.53 | 9.39 | 7.21 | 8.68 | 1.43 | 8.43 | 0.17 | -2.56 | 4.93 | 11.05 | 0.0002 | 0.0159 | 1.80 |
| ENSSSCG00000016511 | 3.66 | 2.34 | 2.76 | 14.81 | 12.00 | 15.12 | 2.92 | 13.98 | 0.21 | -2.26 | 8.45 | 11.08 | 0.0002 | 0.0159 | 1.81 |
| ENSSSCG00000001459 | 1.86 | 1.21 | 1.38 | 5.47 | 4.73 | 4.78 | 1.48 | 4.99 | 0.30 | -1.75 | 3.24 | 11.13 | 0.0002 | 0.0159 | 1.83 |
| ENSSSCG00000036096 | 1.02 | 1.15 | 1.40 | 3.56 | 3.26 | 3.70 | 1.19 | 3.50 | 0.34 | -1.56 | 2.35 | 11.13 | 0.0002 | 0.0159 | 1.83 |
| ENSSSCG00000013765 | 0.54 | 0.54 | 0.30 | 4.15 | 3.28 | 4.16 | 0.46 | 3.86 | 0.12 | -3.06 | 2.16 | 10.94 | 0.0002 | 0.0162 | 1.76 |
| ENSSSCG00000015509 | 12.70 | 11.48 | 13.42 | 27.51 | 23.72 | 27.16 | 12.53 | 26.13 | 0.48 | -1.06 | 19.33 | 10.92 | 0.0002 | 0.0162 | 1.75 |
| ENSSSCG00000022159 | 42.31 | 45.92 | 44.72 | 17.67 | 24.64 | 19.75 | 44.32 | 20.69 | 2.14 | 1.10 | 32.50 | -10.95 | 0.0002 | 0.0162 | 1.76 |
| ENSSSCG00000024109 | 1.88 | 0.99 | 1.00 | 4.74 | 4.58 | 4.58 | 1.29 | 4.63 | 0.28 | -1.85 | 2.96 | 10.89 | 0.0002 | 0.0162 | 1.74 |
| ENSSSCG00000004410 | 1.41 | 0.00 | 0.25 | 15.05 | 14.07 | 11.09 | 0.55 | 13.40 | 0.04 | -4.60 | 6.98 | 10.87 | 0.0002 | 0.0163 | 1.73 |
| ENSSSCG00000002668 | 3.42 | 0.13 | 0.20 | 32.36 | 32.01 | 41.92 | 1.25 | 35.43 | 0.04 | -4.83 | 18.34 | 10.77 | 0.0002 | 0.0168 | 1.69 |
| ENSSSCG00000005529 | 3.92 | 0.25 | 0.22 | 47.19 | 57.64 | 41.91 | 1.47 | 48.91 | 0.03 | -5.06 | 25.19 | 10.71 | 0.0002 | 0.0168 | 1.66 |
| ENSSSCG00000002803 | 1.15 | 0.03 | 0.24 | 8.35 | 6.55 | 8.58 | 0.47 | 7.82 | 0.06 | -4.05 | 4.15 | 10.71 | 0.0002 | 0.0168 | 1.66 |
| ENSSSCG00000008402 | 2.38 | 1.24 | 1.78 | 17.23 | 15.73 | 12.78 | 1.80 | 15.25 | 0.12 | -3.08 | 8.52 | 10.72 | 0.0002 | 0.0168 | 1.66 |
| ENSSSCG00000038001 | 2.18 | 2.70 | 2.36 | 7.51 | 6.92 | 8.58 | 2.41 | 7.67 | 0.31 | -1.67 | 5.04 | 10.70 | 0.0002 | 0.0168 | 1.66 |
| ENSSSCG00000007839 | 6.80 | 6.74 | 8.03 | 16.49 | 16.05 | 18.95 | 7.19 | 17.16 | 0.42 | -1.26 | 12.18 | 10.70 | 0.0002 | 0.0168 | 1.66 |
| ENSSSCG00000003965 | 1.71 | 0.35 | 0.89 | 11.34 | 8.90 | 9.08 | 0.99 | 9.77 | 0.10 | -3.31 | 5.38 | 10.65 | 0.0002 | 0.0170 | 1.63 |
| ENSSSCG00000038993 | 2.57 | 0.76 | 1.78 | 13.15 | 11.91 | 15.55 | 1.71 | 13.54 | 0.13 | -2.99 | 7.62 | 10.66 | 0.0002 | 0.0170 | 1.64 |
| ENSSSCG00000023834 | 4.90 | 0.26 | 0.39 | 66.56 | 47.83 | 56.63 | 1.85 | 57.01 | 0.03 | -4.95 | 29.43 | 10.59 | 0.0002 | 0.0172 | 1.61 |
| ENSSSCG00000037854 | 1.84 | 1.39 | 2.64 | 11.89 | 10.40 | 9.20 | 1.96 | 10.50 | 0.19 | -2.42 | 6.23 | 10.58 | 0.0002 | 0.0172 | 1.61 |
| ENSSSCG00000001715 | 2.51 | 1.14 | 2.09 | 9.96 | 8.28 | 10.40 | 1.92 | 9.55 | 0.20 | -2.32 | 5.73 | 10.60 | 0.0002 | 0.0172 | 1.61 |
| ENSSSCG00000035596 | 1.11 | 0.67 | 0.89 | 3.09 | 3.27 | 2.87 | 0.89 | 3.08 | 0.29 | -1.79 | 1.98 | 10.56 | 0.0002 | 0.0173 | 1.60 |
| ENSSSCG00000027465 | 3.55 | 3.43 | 3.92 | 1.25 | 1.47 | 1.01 | 3.63 | 1.24 | 2.92 | 1.54 | 2.44 | -10.55 | 0.0002 | 0.0173 | 1.59 |
| ENSSSCG00000011217 | 1.22 | 0.29 | 0.14 | 12.27 | 8.91 | 11.09 | 0.55 | 10.75 | 0.05 | -4.28 | 5.65 | 10.49 | 0.0002 | 0.0176 | 1.57 |
| ENSSSCG00000040550 | 1.13 | 0.82 | 1.22 | 4.64 | 4.27 | 3.73 | 1.06 | 4.21 | 0.25 | -1.99 | 2.63 | 10.49 | 0.0002 | 0.0176 | 1.57 |
| ENSSSCG00000018028 | 0.55 | 0.11 | 0.03 | 6.37 | 4.65 | 6.00 | 0.23 | 5.67 | 0.04 | -4.61 | 2.95 | 10.38 | 0.0002 | 0.0176 | 1.52 |
| ENSSSCG00000034743 | 3.01 | 1.10 | 2.00 | 22.31 | 17.30 | 23.46 | 2.04 | 21.02 | 0.10 | -3.37 | 11.53 | 10.38 | 0.0002 | 0.0176 | 1.52 |
| ENSSSCG00000009765 | 2.56 | 1.05 | 1.42 | 13.46 | 10.20 | 12.00 | 1.68 | 11.89 | 0.14 | -2.83 | 6.78 | 10.41 | 0.0002 | 0.0176 | 1.54 |
| ENSSSCG00000010491 | 1.08 | 0.43 | 0.66 | 4.15 | 4.29 | 3.51 | 0.72 | 3.98 | 0.18 | -2.46 | 2.35 | 10.38 | 0.0002 | 0.0176 | 1.52 |
| ENSSSCG00000006495 | 2.05 | 2.93 | 2.44 | 11.61 | 9.33 | 9.45 | 2.47 | 10.13 | 0.24 | -2.04 | 6.30 | 10.39 | 0.0002 | 0.0176 | 1.52 |
| ENSSSCG00000029553 | 2.46 | 2.32 | 2.96 | 5.53 | 5.31 | 6.00 | 2.58 | 5.61 | 0.46 | -1.12 | 4.10 | 10.41 | 0.0002 | 0.0176 | 1.54 |
| ENSSSCG00000023423 | 18.61 | 14.57 | 16.37 | 39.83 | 38.40 | 45.95 | 16.51 | 41.39 | 0.40 | -1.33 | 28.95 | 10.34 | 0.0002 | 0.0179 | 1.50 |
| ENSSSCG00000017932 | 0.51 | 0.08 | 0.06 | 2.26 | 2.13 | 2.23 | 0.21 | 2.21 | 0.10 | -3.36 | 1.21 | 10.32 | 0.0002 | 0.0179 | 1.50 |

| | | | | | | | | | | | | | | |
|---|---|---|---|---|---|---|---|---|---|---|---|---|---|---|
| ENSSSCG00000003823 | 1.55 | 0.05 | 0.13 | 20.61 | 19.53 | 26.66 | 0.57 | 22.27 | 0.03 | -5.28 | 11.42 | 10.29 | 0.0002 | 0.0180 | 1.48 |
| ENSSSCG00000000117 | 2.94 | 1.51 | 1.83 | 21.52 | 17.22 | 23.73 | 2.09 | 20.82 | 0.10 | -3.32 | 11.46 | 10.29 | 0.0002 | 0.0180 | 1.48 |
| ENSSSCG00000003781 | 2.71 | 1.02 | 2.52 | 16.72 | 13.65 | 18.36 | 2.08 | 16.24 | 0.13 | -2.96 | 9.16 | 10.28 | 0.0002 | 0.0180 | 1.48 |
| ENSSSCG00000036033 | 1.34 | 1.83 | 2.35 | 5.85 | 5.20 | 5.33 | 1.84 | 5.46 | 0.34 | -1.57 | 3.65 | 10.26 | 0.0002 | 0.0181 | 1.47 |
| ENSSSCG00000017344 | 0.23 | 0.06 | 0.06 | 3.17 | 3.86 | 4.42 | 0.12 | 3.82 | 0.03 | -5.02 | 1.97 | 10.18 | 0.0002 | 0.0182 | 1.43 |
| ENSSSCG00000011391 | 2.79 | 0.55 | 0.69 | 28.11 | 32.16 | 39.38 | 1.34 | 33.22 | 0.04 | -4.63 | 17.28 | 10.18 | 0.0002 | 0.0182 | 1.43 |
| ENSSSCG00000032775 | 0.94 | 0.00 | 0.10 | 9.05 | 6.56 | 8.30 | 0.34 | 7.97 | 0.04 | -4.53 | 4.16 | 10.18 | 0.0002 | 0.0182 | 1.43 |
| ENSSSCG00000031352 | 0.50 | 0.11 | 0.03 | 3.61 | 2.78 | 3.24 | 0.21 | 3.21 | 0.07 | -3.90 | 1.71 | 10.23 | 0.0002 | 0.0182 | 1.45 |
| ENSSSCG00000023273 | 1.98 | 1.69 | 1.35 | 14.43 | 11.38 | 15.64 | 1.67 | 13.82 | 0.12 | -3.05 | 7.74 | 10.18 | 0.0002 | 0.0182 | 1.43 |
| ENSSSCG00000015549 | 1.36 | 1.36 | 1.93 | 4.63 | 4.03 | 4.37 | 1.55 | 4.34 | 0.36 | -1.49 | 2.95 | 10.21 | 0.0002 | 0.0182 | 1.44 |
| ENSSSCG00000017551 | 3.56 | 2.91 | 3.32 | 11.80 | 10.40 | 13.47 | 3.26 | 11.89 | 0.27 | -1.87 | 7.57 | 10.14 | 0.0002 | 0.0184 | 1.42 |
| ENSSSCG00000026564 | 0.79 | 0.45 | 0.19 | 7.46 | 8.10 | 10.23 | 0.48 | 8.59 | 0.06 | -4.17 | 4.54 | 10.10 | 0.0002 | 0.0186 | 1.40 |
| ENSSSCG00000017963 | 1.14 | 0.24 | 0.25 | 10.08 | 7.29 | 9.16 | 0.54 | 8.84 | 0.06 | -4.03 | 4.69 | 10.09 | 0.0002 | 0.0186 | 1.39 |
| ENSSSCG00000009237 | 1.28 | 2.18 | 1.11 | 10.08 | 9.84 | 12.76 | 1.52 | 10.89 | 0.14 | -2.84 | 6.21 | 10.09 | 0.0002 | 0.0186 | 1.39 |
| ENSSSCG00000010379 | 0.78 | 0.03 | 0.07 | 7.25 | 7.38 | 9.68 | 0.29 | 8.10 | 0.04 | -4.79 | 4.20 | 10.05 | 0.0002 | 0.0187 | 1.38 |
| ENSSSCG00000030800 | 0.24 | 0.00 | 0.05 | 4.70 | 4.64 | 3.41 | 0.10 | 4.25 | 0.02 | -5.44 | 2.18 | 9.97 | 0.0003 | 0.0190 | 1.34 |
| ENSSSCG00000037649 | 0.72 | 0.09 | 0.17 | 8.64 | 9.32 | 6.60 | 0.32 | 8.18 | 0.04 | -4.66 | 4.25 | 9.96 | 0.0003 | 0.0190 | 1.33 |
| ENSSSCG00000025788 | 3.03 | 2.86 | 4.04 | 22.56 | 16.64 | 21.04 | 3.31 | 20.08 | 0.16 | -2.60 | 11.69 | 9.96 | 0.0003 | 0.0190 | 1.33 |
| ENSSSCG00000008852 | 1.14 | 1.12 | 1.50 | 7.50 | 6.49 | 5.64 | 1.26 | 6.54 | 0.19 | -2.38 | 3.90 | 10.00 | 0.0003 | 0.0190 | 1.35 |
| ENSSSCG00000007454 | 7.89 | 5.62 | 6.88 | 20.78 | 16.88 | 19.44 | 6.79 | 19.03 | 0.36 | -1.49 | 12.91 | 9.96 | 0.0003 | 0.0190 | 1.33 |
| ENSSSCG00000015499 | 4.83 | 5.96 | 6.61 | 12.28 | 13.59 | 14.44 | 5.80 | 13.44 | 0.43 | -1.21 | 9.62 | 9.96 | 0.0003 | 0.0190 | 1.33 |
| ENSSSCG00000028471 | 1.78 | 0.11 | 0.23 | 31.79 | 22.86 | 24.39 | 0.71 | 26.35 | 0.03 | -5.22 | 13.53 | 9.84 | 0.0003 | 0.0191 | 1.28 |
| ENSSSCG00000023904 | 0.83 | 0.04 | 0.07 | 9.46 | 6.74 | 8.82 | 0.31 | 8.34 | 0.04 | -4.73 | 4.33 | 9.91 | 0.0003 | 0.0191 | 1.31 |
| ENSSSCG00000039815 | 0.22 | 0.02 | 0.06 | 1.83 | 2.04 | 2.34 | 0.10 | 2.07 | 0.05 | -4.35 | 1.08 | 9.84 | 0.0003 | 0.0191 | 1.28 |
| ENSSSCG00000032814 | 0.99 | 0.05 | 0.12 | 9.13 | 7.88 | 6.55 | 0.39 | 7.85 | 0.05 | -4.35 | 4.12 | 9.85 | 0.0003 | 0.0191 | 1.28 |
| ENSSSCG00000008970 | 1.24 | 0.25 | 0.26 | 11.88 | 8.47 | 10.43 | 0.58 | 10.26 | 0.06 | -4.14 | 5.42 | 9.92 | 0.0003 | 0.0191 | 1.31 |
| ENSSSCG00000003257 | 1.74 | 0.85 | 0.75 | 7.16 | 9.48 | 9.32 | 1.11 | 8.65 | 0.13 | -2.96 | 4.88 | 9.86 | 0.0003 | 0.0191 | 1.29 |
| ENSSSCG00000002353 | 2.22 | 1.06 | 1.64 | 10.69 | 8.08 | 10.18 | 1.64 | 9.65 | 0.17 | -2.56 | 5.65 | 9.85 | 0.0003 | 0.0191 | 1.28 |
| ENSSSCG00000008761 | 2.93 | 2.14 | 3.17 | 6.10 | 6.79 | 6.29 | 2.75 | 6.39 | 0.43 | -1.22 | 4.57 | 9.85 | 0.0003 | 0.0191 | 1.28 |
| ENSSSCG00000013482 | 1.88 | 1.17 | 1.15 | 6.69 | 8.67 | 8.75 | 1.40 | 8.04 | 0.17 | -2.52 | 4.72 | 9.82 | 0.0003 | 0.0193 | 1.27 |
| ENSSSCG00000010664 | 2.90 | 0.16 | 0.50 | 26.18 | 28.52 | 36.41 | 1.19 | 30.37 | 0.04 | -4.68 | 15.78 | 9.80 | 0.0003 | 0.0193 | 1.26 |
| ENSSSCG00000015109 | 2.28 | 0.22 | 0.14 | 14.36 | 14.54 | 18.94 | 0.88 | 15.95 | 0.06 | -4.18 | 8.41 | 9.80 | 0.0003 | 0.0193 | 1.26 |
| ENSSSCG00000017609 | 0.52 | 0.03 | 0.02 | 7.18 | 5.15 | 5.70 | 0.19 | 6.01 | 0.03 | -4.99 | 3.10 | 9.74 | 0.0003 | 0.0197 | 1.23 |
| ENSSSCG00000008410 | 0.30 | 0.04 | 0.16 | 4.16 | 3.08 | 3.26 | 0.16 | 3.50 | 0.05 | -4.42 | 1.83 | 9.69 | 0.0003 | 0.0198 | 1.21 |
| ENSSSCG00000014004 | 1.75 | 1.38 | 1.76 | 20.13 | 15.65 | 22.34 | 1.63 | 19.38 | 0.08 | -3.57 | 10.50 | 9.69 | 0.0003 | 0.0198 | 1.21 |

| | | | | | | | | | | | | | | |
|---|---|---|---|---|---|---|---|---|---|---|---|---|---|---|
| ENSSSCG00000031112 | 0.60 | 0.72 | 0.76 | 2.52 | 2.34 | 2.83 | 0.70 | 2.57 | 0.27 | -1.88 | 1.63 | 9.71 | 0.0003 | 0.0198 | 1.22 |
| ENSSSCG00000031102 | 3.33 | 2.75 | 3.61 | 6.77 | 6.87 | 7.79 | 3.23 | 7.14 | 0.45 | -1.14 | 5.19 | 9.69 | 0.0003 | 0.0198 | 1.21 |
| ENSSSCG00000000038 | 139.14 | 147.27 | 162.53 | 50.47 | 60.85 | 74.59 | 149.65 | 61.97 | 2.41 | 1.27 | 105.81 | -9.67 | 0.0003 | 0.0198 | 1.20 |
| ENSSSCG00000001594 | 0.13 | 0.03 | 0.03 | 2.70 | 2.25 | 3.14 | 0.07 | 2.70 | 0.02 | -5.32 | 1.38 | 9.62 | 0.0003 | 0.0200 | 1.18 |
| ENSSSCG00000015100 | 0.26 | 0.04 | 0.04 | 2.74 | 3.27 | 3.85 | 0.11 | 3.29 | 0.03 | -4.87 | 1.70 | 9.60 | 0.0003 | 0.0200 | 1.17 |
| ENSSSCG00000037044 | 5.13 | 0.39 | 0.57 | 38.51 | 47.15 | 55.13 | 2.03 | 46.93 | 0.04 | -4.53 | 24.48 | 9.60 | 0.0003 | 0.0200 | 1.17 |
| ENSSSCG00000039802 | 2.59 | 1.12 | 1.47 | 8.21 | 6.98 | 6.88 | 1.72 | 7.36 | 0.23 | -2.09 | 4.54 | 9.61 | 0.0003 | 0.0200 | 1.17 |
| ENSSSCG00000015048 | 0.51 | 0.20 | 0.03 | 8.54 | 5.90 | 7.22 | 0.25 | 7.22 | 0.03 | -4.88 | 3.73 | 9.55 | 0.0003 | 0.0200 | 1.14 |
| ENSSSCG00000021764 | 0.33 | 0.12 | 0.14 | 4.14 | 3.30 | 3.02 | 0.19 | 3.49 | 0.06 | -4.17 | 1.84 | 9.54 | 0.0003 | 0.0200 | 1.13 |
| ENSSSCG00000003455 | 2.39 | 0.53 | 0.66 | 24.40 | 17.06 | 22.37 | 1.19 | 21.28 | 0.06 | -4.16 | 11.23 | 9.53 | 0.0003 | 0.0200 | 1.13 |
| ENSSSCG00000016414 | 0.50 | 0.15 | 0.15 | 2.89 | 2.64 | 3.47 | 0.26 | 3.00 | 0.09 | -3.50 | 1.63 | 9.55 | 0.0003 | 0.0200 | 1.14 |
| ENSSSCG00000007935 | 2.03 | 1.17 | 1.52 | 12.24 | 10.52 | 14.65 | 1.57 | 12.47 | 0.13 | -2.99 | 7.02 | 9.54 | 0.0003 | 0.0200 | 1.14 |
| ENSSSCG00000031727 | 2.58 | 1.80 | 2.34 | 7.76 | 7.04 | 9.05 | 2.24 | 7.95 | 0.28 | -1.83 | 5.09 | 9.54 | 0.0003 | 0.0200 | 1.14 |
| ENSSSCG00000012126 | 34.09 | 36.75 | 37.33 | 15.95 | 21.38 | 15.98 | 36.05 | 17.77 | 2.03 | 1.02 | 26.91 | -9.55 | 0.0003 | 0.0200 | 1.14 |
| ENSSSCG00000007488 | 0.12 | 0.02 | 0.05 | 1.47 | 1.34 | 1.36 | 0.06 | 1.39 | 0.04 | -4.48 | 0.73 | 9.44 | 0.0003 | 0.0202 | 1.09 |
| ENSSSCG00000036621 | 0.36 | 0.17 | 0.16 | 2.20 | 2.10 | 2.68 | 0.23 | 2.33 | 0.10 | -3.32 | 1.28 | 9.46 | 0.0003 | 0.0202 | 1.10 |
| ENSSSCG00000005221 | 4.25 | 1.69 | 4.16 | 25.75 | 21.75 | 30.13 | 3.37 | 25.88 | 0.13 | -2.94 | 14.62 | 9.48 | 0.0003 | 0.0202 | 1.11 |
| ENSSSCG00000013313 | 1.81 | 3.76 | 2.18 | 15.11 | 13.39 | 17.99 | 2.58 | 15.50 | 0.17 | -2.59 | 9.04 | 9.45 | 0.0003 | 0.0202 | 1.09 |
| ENSSSCG00000025041 | 1.44 | 0.96 | 0.91 | 5.65 | 4.60 | 6.11 | 1.10 | 5.46 | 0.20 | -2.31 | 3.28 | 9.44 | 0.0003 | 0.0202 | 1.09 |
| ENSSSCG00000004270 | 0.93 | 0.38 | 0.46 | 2.99 | 2.59 | 2.81 | 0.59 | 2.80 | 0.21 | -2.25 | 1.69 | 9.49 | 0.0003 | 0.0202 | 1.11 |
| ENSSSCG00000004172 | 0.66 | 0.74 | 1.13 | 4.00 | 4.29 | 3.35 | 0.84 | 3.88 | 0.22 | -2.21 | 2.36 | 9.50 | 0.0003 | 0.0202 | 1.12 |
| ENSSSCG00000026367 | 7.13 | 7.77 | 6.80 | 26.01 | 21.71 | 28.81 | 7.24 | 25.51 | 0.28 | -1.82 | 16.37 | 9.44 | 0.0003 | 0.0202 | 1.09 |
| ENSSSCG00000001786 | 3.04 | 0.34 | 0.16 | 41.11 | 29.35 | 41.88 | 1.18 | 37.45 | 0.03 | -4.99 | 19.31 | 9.41 | 0.0003 | 0.0204 | 1.07 |
| ENSSSCG00000012235 | 3.71 | 1.70 | 2.15 | 17.69 | 13.93 | 13.67 | 2.52 | 15.10 | 0.17 | -2.58 | 8.81 | 9.40 | 0.0003 | 0.0204 | 1.07 |
| ENSSSCG00000033000 | 1.55 | 2.31 | 1.80 | 7.42 | 6.07 | 6.02 | 1.89 | 6.51 | 0.29 | -1.79 | 4.20 | 9.41 | 0.0003 | 0.0204 | 1.07 |
| ENSSSCG00000003508 | 0.40 | 0.11 | 0.16 | 3.18 | 2.34 | 2.85 | 0.22 | 2.79 | 0.08 | -3.64 | 1.51 | 9.37 | 0.0003 | 0.0204 | 1.06 |
| ENSSSCG00000023957 | 17.51 | 6.15 | 8.77 | 131.26 | 106.41 | 152.39 | 10.81 | 130.02 | 0.08 | -3.59 | 70.41 | 9.37 | 0.0003 | 0.0204 | 1.06 |
| ENSSSCG00000004388 | 1.56 | 0.50 | 2.16 | 6.03 | 5.73 | 5.85 | 1.41 | 5.87 | 0.24 | -2.06 | 3.64 | 9.39 | 0.0003 | 0.0204 | 1.06 |
| ENSSSCG00000006309 | 1.86 | 1.13 | 1.86 | 4.87 | 5.79 | 4.92 | 1.62 | 5.19 | 0.31 | -1.68 | 3.41 | 9.36 | 0.0003 | 0.0204 | 1.05 |
| ENSSSCG00000033739 | 0.72 | 0.04 | 0.09 | 8.35 | 7.54 | 5.78 | 0.28 | 7.22 | 0.04 | -4.67 | 3.75 | 9.35 | 0.0003 | 0.0205 | 1.04 |
| ENSSSCG00000017187 | 20.93 | 0.44 | 0.88 | 161.12 | 138.81 | 197.38 | 7.42 | 165.77 | 0.04 | -4.48 | 86.59 | 9.31 | 0.0003 | 0.0207 | 1.03 |
| ENSSSCG00000010164 | 3.18 | 2.67 | 3.75 | 9.44 | 7.85 | 9.69 | 3.20 | 8.99 | 0.36 | -1.49 | 6.10 | 9.32 | 0.0003 | 0.0207 | 1.03 |
| ENSSSCG00000023057 | 0.66 | 0.23 | 0.23 | 3.45 | 2.75 | 3.65 | 0.37 | 3.28 | 0.11 | -3.13 | 1.83 | 9.30 | 0.0003 | 0.0207 | 1.02 |
| ENSSSCG00000009796 | 1.25 | 0.09 | 0.13 | 10.86 | 10.19 | 7.66 | 0.49 | 9.57 | 0.05 | -4.28 | 5.03 | 9.29 | 0.0004 | 0.0207 | 1.01 |
| ENSSSCG00000028060 | 0.82 | 0.21 | 0.36 | 5.60 | 4.11 | 5.54 | 0.46 | 5.08 | 0.09 | -3.46 | 2.77 | 9.27 | 0.0004 | 0.0207 | 1.01 |

| | | | | | | | | | | | | | | |
|---|---|---|---|---|---|---|---|---|---|---|---|---|---|---|
| ENSSSCG00000013546 | 1.05 | 1.64 | 1.79 | 4.67 | 4.01 | 4.59 | 1.49 | 4.42 | 0.34 | -1.57 | 2.96 | 9.27 | 0.0004 | 0.0207 | 1.01 |
| ENSSSCG00000008908 | 0.42 | 0.00 | 0.00 | 4.25 | 4.79 | 6.05 | 0.14 | 5.03 | 0.03 | -5.16 | 2.59 | 9.26 | 0.0004 | 0.0208 | 1.00 |
| ENSSSCG00000000679 | 3.83 | 0.28 | 0.56 | 27.78 | 24.28 | 34.43 | 1.56 | 28.83 | 0.05 | -4.21 | 15.19 | 9.23 | 0.0004 | 0.0210 | 0.98 |
| ENSSSCG00000008397 | 3.07 | 4.60 | 5.39 | 29.93 | 28.88 | 39.05 | 4.35 | 32.62 | 0.13 | -2.91 | 18.49 | 9.23 | 0.0004 | 0.0210 | 0.99 |
| ENSSSCG00000001203 | 1.63 | 0.77 | 1.23 | 3.75 | 3.69 | 4.06 | 1.21 | 3.83 | 0.32 | -1.66 | 2.52 | 9.20 | 0.0004 | 0.0212 | 0.97 |
| ENSSSCG00000027144 | 0.90 | 0.19 | 0.53 | 5.20 | 4.36 | 3.86 | 0.54 | 4.47 | 0.12 | -3.05 | 2.50 | 9.12 | 0.0004 | 0.0217 | 0.93 |
| ENSSSCG00000009956 | 6.82 | 7.32 | 6.50 | 22.39 | 19.21 | 25.49 | 6.88 | 22.36 | 0.31 | -1.70 | 14.62 | 9.12 | 0.0004 | 0.0217 | 0.93 |
| ENSSSCG00000031876 | 4.86 | 5.23 | 7.67 | 15.49 | 18.83 | 16.86 | 5.92 | 17.06 | 0.35 | -1.53 | 11.49 | 9.13 | 0.0004 | 0.0217 | 0.93 |
| ENSSSCG00000028126 | 2.37 | 1.50 | 2.82 | 5.61 | 5.94 | 5.90 | 2.23 | 5.82 | 0.38 | -1.38 | 4.03 | 9.10 | 0.0004 | 0.0218 | 0.92 |
| ENSSSCG00000015197 | 3.82 | 0.60 | 0.59 | 29.32 | 35.47 | 42.85 | 1.67 | 35.88 | 0.05 | -4.43 | 18.78 | 9.10 | 0.0004 | 0.0218 | 0.92 |
| ENSSSCG00000015696 | 0.70 | 0.01 | 0.05 | 10.47 | 7.47 | 7.77 | 0.25 | 8.57 | 0.03 | -5.09 | 4.41 | 9.09 | 0.0004 | 0.0218 | 0.91 |
| ENSSSCG00000031652 | 2.64 | 1.92 | 2.45 | 8.52 | 7.97 | 6.57 | 2.33 | 7.69 | 0.30 | -1.72 | 5.01 | 9.08 | 0.0004 | 0.0219 | 0.91 |
| ENSSSCG00000006726 | 0.62 | 0.03 | 0.06 | 6.45 | 5.12 | 4.55 | 0.23 | 5.37 | 0.04 | -4.52 | 2.80 | 9.03 | 0.0004 | 0.0220 | 0.89 |
| ENSSSCG00000010614 | 0.73 | 0.05 | 0.19 | 8.36 | 7.43 | 5.70 | 0.32 | 7.16 | 0.05 | -4.47 | 3.74 | 9.00 | 0.0004 | 0.0220 | 0.87 |
| ENSSSCG00000034547 | 0.59 | 0.04 | 0.05 | 4.85 | 3.47 | 4.66 | 0.23 | 4.33 | 0.05 | -4.26 | 2.28 | 9.04 | 0.0004 | 0.0220 | 0.89 |
| ENSSSCG00000031764 | 1.09 | 0.85 | 1.99 | 5.11 | 4.78 | 5.71 | 1.31 | 5.20 | 0.25 | -1.99 | 3.25 | 9.05 | 0.0004 | 0.0220 | 0.89 |
| ENSSSCG00000002805 | 7.56 | 5.70 | 5.74 | 20.55 | 16.07 | 19.08 | 6.33 | 18.57 | 0.34 | -1.55 | 12.45 | 9.03 | 0.0004 | 0.0220 | 0.88 |
| ENSSSCG00000005655 | 22.57 | 14.22 | 14.69 | 50.95 | 44.98 | 54.94 | 17.16 | 50.29 | 0.34 | -1.55 | 33.73 | 9.02 | 0.0004 | 0.0220 | 0.88 |
| ENSSSCG00000011849 | 7.83 | 6.08 | 8.15 | 16.00 | 14.10 | 16.46 | 7.35 | 15.52 | 0.47 | -1.08 | 11.44 | 9.00 | 0.0004 | 0.0220 | 0.87 |
| ENSSSCG00000033351 | 2.34 | 2.30 | 3.15 | 5.61 | 5.19 | 5.40 | 2.60 | 5.40 | 0.48 | -1.05 | 4.00 | 9.00 | 0.0004 | 0.0220 | 0.87 |
| ENSSSCG00000016203 | 1.97 | 0.04 | 0.16 | 18.04 | 12.60 | 14.38 | 0.72 | 15.01 | 0.05 | -4.37 | 7.87 | 8.95 | 0.0004 | 0.0225 | 0.84 |
| ENSSSCG00000011249 | 1.14 | 0.31 | 0.28 | 7.40 | 5.43 | 5.78 | 0.58 | 6.20 | 0.09 | -3.43 | 3.39 | 8.87 | 0.0004 | 0.0230 | 0.80 |
| ENSSSCG00000012295 | 2.15 | 1.59 | 2.14 | 6.93 | 5.46 | 6.95 | 1.96 | 6.45 | 0.30 | -1.72 | 4.20 | 8.85 | 0.0004 | 0.0232 | 0.79 |
| ENSSSCG00000016444 | 43.38 | 41.94 | 43.24 | 10.61 | 20.82 | 19.00 | 42.85 | 16.81 | 2.55 | 1.35 | 29.83 | -8.84 | 0.0004 | 0.0232 | 0.78 |
| ENSSSCG00000007991 | 2.71 | 1.07 | 1.36 | 12.53 | 9.17 | 10.47 | 1.71 | 10.73 | 0.16 | -2.65 | 6.22 | 8.77 | 0.0005 | 0.0239 | 0.75 |
| ENSSSCG00000020990 | 0.79 | 0.03 | 0.11 | 11.27 | 12.59 | 8.34 | 0.31 | 10.74 | 0.03 | -5.10 | 5.52 | 8.75 | 0.0005 | 0.0239 | 0.74 |
| ENSSSCG00000032076 | 1.34 | 0.07 | 0.09 | 13.37 | 10.06 | 14.99 | 0.50 | 12.81 | 0.04 | -4.69 | 6.65 | 8.75 | 0.0005 | 0.0239 | 0.74 |
| ENSSSCG00000015823 | 3.62 | 3.11 | 2.94 | 13.95 | 10.74 | 14.61 | 3.22 | 13.10 | 0.25 | -2.02 | 8.16 | 8.72 | 0.0005 | 0.0242 | 0.72 |
| ENSSSCG00000038153 | 1.28 | 0.28 | 0.31 | 13.12 | 9.35 | 13.76 | 0.62 | 12.08 | 0.05 | -4.28 | 6.35 | 8.69 | 0.0005 | 0.0243 | 0.70 |
| ENSSSCG00000017446 | 3.92 | 3.32 | 3.28 | 22.19 | 21.77 | 29.79 | 3.51 | 24.58 | 0.14 | -2.81 | 14.04 | 8.69 | 0.0005 | 0.0243 | 0.71 |
| ENSSSCG00000039469 | 8.51 | 9.85 | 11.44 | 33.39 | 25.67 | 28.67 | 9.93 | 29.24 | 0.34 | -1.56 | 19.59 | 8.67 | 0.0005 | 0.0245 | 0.69 |
| ENSSSCG00000004008 | 1.05 | 0.53 | 0.96 | 2.81 | 2.83 | 2.55 | 0.85 | 2.73 | 0.31 | -1.69 | 1.79 | 8.64 | 0.0005 | 0.0246 | 0.68 |
| ENSSSCG00000005055 | 110.23 | 115.17 | 100.51 | 249.15 | 213.19 | 268.93 | 108.63 | 243.76 | 0.45 | -1.17 | 176.20 | 8.64 | 0.0005 | 0.0246 | 0.68 |
| ENSSSCG00000007236 | 0.91 | 0.03 | 0.12 | 13.24 | 9.15 | 13.57 | 0.35 | 11.99 | 0.03 | -5.09 | 6.17 | 8.63 | 0.0005 | 0.0247 | 0.67 |
| ENSSSCG00000008186 | 3.70 | 2.53 | 3.49 | 18.39 | 19.08 | 13.70 | 3.24 | 17.06 | 0.19 | -2.40 | 10.15 | 8.61 | 0.0005 | 0.0247 | 0.66 |

| | | | | | | | | | | | | | | |
|---|---|---|---|---|---|---|---|---|---|---|---|---|---|---|
| ENSSSCG00000033259 | 1.36 | 0.31 | 0.48 | 9.68 | 8.23 | 12.03 | 0.71 | 9.98 | 0.07 | -3.80 | 5.35 | 8.60 | 0.0005 | 0.0248 | 0.66 |
| ENSSSCG00000039903 | 0.08 | 0.24 | 0.42 | 9.54 | 6.22 | 8.35 | 0.25 | 8.04 | 0.03 | -5.02 | 4.14 | 8.54 | 0.0005 | 0.0253 | 0.62 |
| ENSSSCG00000006462 | 2.88 | 0.22 | 0.41 | 25.90 | 20.54 | 30.85 | 1.17 | 25.77 | 0.05 | -4.46 | 13.47 | 8.56 | 0.0005 | 0.0253 | 0.63 |
| ENSSSCG00000016166 | 0.76 | 0.20 | 0.25 | 8.56 | 5.70 | 7.32 | 0.41 | 7.20 | 0.06 | -4.15 | 3.80 | 8.54 | 0.0005 | 0.0253 | 0.63 |
| ENSSSCG00000007149 | 6.61 | 2.74 | 2.06 | 29.74 | 25.89 | 36.44 | 3.80 | 30.69 | 0.12 | -3.01 | 17.25 | 8.54 | 0.0005 | 0.0253 | 0.63 |
| ENSSSCG00000040989 | 3.19 | 1.79 | 1.35 | 27.50 | 20.86 | 31.28 | 2.11 | 26.55 | 0.08 | -3.65 | 14.33 | 8.51 | 0.0005 | 0.0254 | 0.61 |
| ENSSSCG00000026175 | 1.17 | 0.03 | 0.13 | 9.37 | 13.50 | 13.91 | 0.44 | 12.26 | 0.04 | -4.79 | 6.35 | 8.49 | 0.0005 | 0.0256 | 0.60 |
| ENSSSCG00000025685 | 1.97 | 0.11 | 0.25 | 22.35 | 14.73 | 19.35 | 0.78 | 18.81 | 0.04 | -4.60 | 9.79 | 8.46 | 0.0005 | 0.0257 | 0.58 |
| ENSSSCG00000003825 | 1.36 | 1.89 | 0.68 | 10.43 | 14.18 | 15.39 | 1.31 | 13.33 | 0.10 | -3.35 | 7.32 | 8.43 | 0.0005 | 0.0257 | 0.56 |
| ENSSSCG00000010770 | 1.30 | 0.72 | 0.60 | 9.77 | 6.88 | 7.49 | 0.88 | 8.05 | 0.11 | -3.20 | 4.46 | 8.44 | 0.0005 | 0.0257 | 0.57 |
| ENSSSCG00000011744 | 0.57 | 0.70 | 0.56 | 5.37 | 3.71 | 4.95 | 0.61 | 4.67 | 0.13 | -2.94 | 2.64 | 8.44 | 0.0005 | 0.0257 | 0.57 |
| ENSSSCG00000010402 | 0.81 | 0.31 | 0.61 | 4.03 | 2.95 | 3.49 | 0.57 | 3.49 | 0.16 | -2.60 | 2.03 | 8.47 | 0.0005 | 0.0257 | 0.58 |
| ENSSSCG00000027587 | 4.58 | 2.28 | 2.16 | 17.28 | 12.96 | 14.27 | 3.00 | 14.84 | 0.20 | -2.30 | 8.92 | 8.46 | 0.0005 | 0.0257 | 0.58 |
| ENSSSCG00000014818 | 2.51 | 2.98 | 3.87 | 8.53 | 7.05 | 7.95 | 3.12 | 7.84 | 0.40 | -1.33 | 5.48 | 8.43 | 0.0005 | 0.0257 | 0.56 |
| ENSSSCG00000009783 | 5.81 | 4.02 | 6.22 | 12.08 | 11.06 | 12.72 | 5.35 | 11.95 | 0.45 | -1.16 | 8.65 | 8.46 | 0.0005 | 0.0257 | 0.58 |
| ENSSSCG00000008834 | 6.36 | 0.65 | 0.68 | 77.31 | 55.14 | 84.05 | 2.56 | 72.17 | 0.04 | -4.82 | 37.37 | 8.41 | 0.0005 | 0.0259 | 0.55 |
| ENSSSCG00000032203 | 0.36 | 0.08 | 0.07 | 2.87 | 4.11 | 3.15 | 0.17 | 3.38 | 0.05 | -4.30 | 1.77 | 8.40 | 0.0006 | 0.0259 | 0.55 |
| ENSSSCG00000005967 | 2.48 | 1.17 | 2.60 | 15.87 | 13.02 | 18.97 | 2.09 | 15.95 | 0.13 | -2.93 | 9.02 | 8.40 | 0.0006 | 0.0259 | 0.55 |
| ENSSSCG00000031441 | 4.73 | 3.17 | 3.38 | 8.67 | 8.13 | 9.49 | 3.76 | 8.76 | 0.43 | -1.22 | 6.26 | 8.38 | 0.0006 | 0.0260 | 0.54 |
| ENSSSCG00000040224 | 1.46 | 0.08 | 0.13 | 13.38 | 12.31 | 17.99 | 0.56 | 14.56 | 0.04 | -4.71 | 7.56 | 8.37 | 0.0006 | 0.0261 | 0.53 |
| ENSSSCG00000035577 | 0.62 | 0.01 | 0.03 | 8.46 | 5.77 | 6.31 | 0.22 | 6.84 | 0.03 | -4.96 | 3.53 | 8.34 | 0.0006 | 0.0262 | 0.51 |
| ENSSSCG00000039395 | 1.36 | 0.15 | 0.42 | 14.73 | 9.89 | 14.36 | 0.64 | 12.99 | 0.05 | -4.34 | 6.82 | 8.32 | 0.0006 | 0.0262 | 0.50 |
| ENSSSCG00000028587 | 0.64 | 0.03 | 0.10 | 4.52 | 3.38 | 4.94 | 0.26 | 4.28 | 0.06 | -4.07 | 2.27 | 8.32 | 0.0006 | 0.0262 | 0.51 |
| ENSSSCG00000008491 | 10.18 | 3.27 | 3.98 | 93.17 | 78.82 | 118.34 | 5.81 | 96.78 | 0.06 | -4.06 | 51.29 | 8.35 | 0.0006 | 0.0262 | 0.52 |
| ENSSSCG00000011181 | 0.99 | 0.38 | 0.61 | 8.97 | 6.48 | 9.69 | 0.66 | 8.38 | 0.08 | -3.67 | 4.52 | 8.35 | 0.0006 | 0.0262 | 0.52 |
| ENSSSCG00000017601 | 1.55 | 0.82 | 0.92 | 6.38 | 4.86 | 6.74 | 1.10 | 5.99 | 0.18 | -2.45 | 3.55 | 8.32 | 0.0006 | 0.0262 | 0.50 |
| ENSSSCG00000028144 | 1.91 | 1.61 | 2.59 | 5.18 | 4.57 | 5.18 | 2.04 | 4.98 | 0.41 | -1.29 | 3.51 | 8.33 | 0.0006 | 0.0262 | 0.51 |
| ENSSSCG00000003755 | 0.78 | 0.32 | 0.39 | 4.22 | 3.71 | 5.30 | 0.50 | 4.41 | 0.11 | -3.14 | 2.46 | 8.31 | 0.0006 | 0.0263 | 0.50 |
| ENSSSCG00000015021 | 3.50 | 0.08 | 0.35 | 23.42 | 32.96 | 35.38 | 1.31 | 30.58 | 0.04 | -4.55 | 15.95 | 8.29 | 0.0006 | 0.0265 | 0.48 |
| ENSSSCG00000022643 | 0.41 | 0.19 | 0.15 | 3.31 | 2.44 | 3.53 | 0.25 | 3.09 | 0.08 | -3.62 | 1.67 | 8.28 | 0.0006 | 0.0265 | 0.48 |
| ENSSSCG00000039793 | 5.30 | 5.77 | 5.28 | 12.36 | 10.15 | 10.46 | 5.45 | 10.99 | 0.50 | -1.01 | 8.22 | 8.28 | 0.0006 | 0.0265 | 0.48 |
| ENSSSCG00000030167 | 44.79 | 48.30 | 44.40 | 18.51 | 20.97 | 27.82 | 45.83 | 22.43 | 2.04 | 1.03 | 34.13 | -8.27 | 0.0006 | 0.0265 | 0.48 |
| ENSSSCG00000040698 | 3.34 | 1.78 | 2.37 | 9.49 | 7.49 | 9.56 | 2.50 | 8.84 | 0.28 | -1.82 | 5.67 | 8.25 | 0.0006 | 0.0266 | 0.46 |
| ENSSSCG00000031781 | 9.90 | 9.28 | 7.44 | 25.05 | 19.95 | 21.06 | 8.87 | 22.02 | 0.40 | -1.31 | 15.45 | 8.25 | 0.0006 | 0.0266 | 0.46 |
| ENSSSCG00000009332 | 1.52 | 0.57 | 0.21 | 24.52 | 16.52 | 18.07 | 0.76 | 19.70 | 0.04 | -4.69 | 10.23 | 8.22 | 0.0006 | 0.0270 | 0.45 |

### Table S3 Alternative splicing analysis

| event_type | gene_id | chr | event start | event end | XL13 | | XL17 | | XL8 | | XS12 | | XS15 | | XS16 | |
|---|---|---|---|---|---|---|---|---|---|---|---|---|---|---|---|---|
| | | | | | gene fpkm | AS fpkm | gene fpkm | AS fpkm | gene fpkm | AS fpkm | gene fpkm | AS fpkm | gene fpkm | AS fpkm | gene fpkm | AS fpkm |
| IR_OFF | ENSSSCG00000000002 | 5 | 3230944 | 3230975 | 2.87 | 0.27 | 1.91 | 0.16 | 2.41 | 0.06 | 1.31 | 0.04 | 3.61 | 0.13 | 3.20 | 0.17 |
| XSKIP_ON | ENSSSCG00000000002 | 5 | 3229095 | 3229139 | 2.87 | 0.27 | 1.91 | 0.16 | 2.41 | 0.06 | 1.31 | 0.04 | 3.61 | 0.13 | 3.20 | 0.17 |
| TSS | ENSSSCG00000000002 | 5 | 3246191 | 3246479 | 2.87 | 0.25 | 1.91 | 0.16 | 2.41 | 0.20 | 1.31 | 0.11 | 3.61 | 0.48 | 3.20 | 0.22 |
| XAE | ENSSSCG00000000002 | 5 | 3245642 | 3245737 | 2.87 | 0.25 | 1.91 | 0.16 | 2.41 | 0.20 | 1.31 | 0.11 | 3.61 | 0.48 | 3.20 | 0.22 |
| IR_ON | ENSSSCG00000000002 | 5 | 3230944 | 3230975 | 2.87 | 2.35 | 1.91 | 1.59 | 2.41 | 2.15 | 1.31 | 1.16 | 3.61 | 3.00 | 3.20 | 2.81 |
| TSS | ENSSSCG00000000002 | 5 | 3246207 | 3246243 | 2.87 | 2.35 | 1.91 | 1.59 | 2.41 | 2.15 | 1.31 | 1.16 | 3.61 | 3.00 | 3.20 | 2.81 |
| XAE | ENSSSCG00000000002 | 5 | 3245642 | 3245750 | 2.87 | 2.35 | 1.91 | 1.59 | 2.41 | 2.15 | 1.31 | 1.16 | 3.61 | 3.00 | 3.20 | 2.81 |
| XSKIP_OFF | ENSSSCG00000000002 | 5 | 3229095 | 3229139 | 2.87 | 2.35 | 1.91 | 1.59 | 2.41 | 2.15 | 1.31 | 1.16 | 3.61 | 3.00 | 3.20 | 2.81 |
| TTS | ENSSSCG00000000002 | 5 | 3228196 | 3228279 | 2.87 | 2.62 | 1.91 | 1.75 | 2.41 | 2.21 | 1.31 | 1.21 | 3.61 | 3.13 | 3.20 | 2.98 |
| SKIP_OFF | ENSSSCG00000000003 | 5 | 3256861 | 3256899 | 2.72 | 0.00 | 2.90 | 0.08 | 3.34 | 0.06 | 8.51 | 0.07 | 7.50 | 0.06 | 13.87 | 0.06 |
| TSS | ENSSSCG00000000003 | 5 | 3270722 | 3270814 | 2.72 | 2.72 | 2.90 | 2.82 | 3.34 | 3.28 | 8.51 | 8.44 | 7.50 | 7.44 | 13.87 | 13.81 |
| SKIP_ON | ENSSSCG00000000003 | 5 | 3256861 | 3256899 | 2.72 | 2.72 | 2.90 | 2.82 | 3.34 | 3.28 | 8.51 | 8.44 | 7.50 | 7.44 | 13.87 | 13.81 |
| TTS | ENSSSCG00000000003 | 5 | 3245883 | 3249295 | 2.72 | 2.72 | 2.90 | 2.90 | 3.34 | 3.34 | 8.51 | 8.51 | 7.50 | 7.50 | 13.87 | 13.87 |
| TSS | ENSSSCG00000000005 | 5 | 3287462 | 3287506 | 7.35 | 7.35 | 5.39 | 5.39 | 12.67 | 12.67 | 9.60 | 9.60 | 8.93 | 8.93 | 10.06 | 10.06 |
| TTS | ENSSSCG00000000005 | 5 | 3294047 | 3294638 | 7.35 | 7.35 | 5.39 | 5.39 | 12.67 | 12.67 | 9.60 | 9.60 | 8.93 | 8.93 | 10.06 | 10.06 |
| TSS | ENSSSCG00000000006 | 5 | 3320884 | 3321091 | 2.24 | 2.24 | 2.70 | 2.70 | 3.07 | 3.07 | 1.50 | 1.50 | 2.69 | 2.69 | 1.54 | 1.54 |
| TTS | ENSSSCG00000000006 | 5 | 3302755 | 3303002 | 2.24 | 2.24 | 2.70 | 2.70 | 3.07 | 3.07 | 1.50 | 1.50 | 2.69 | 2.69 | 1.54 | 1.54 |
| SKIP_OFF | ENSSSCG00000000007 | 5 | 3213140 | 3213312 | 6.48 | 1.26 | 4.50 | 0.06 | 5.79 | 0.58 | 7.05 | 1.63 | 5.03 | 0.84 | 5.44 | 1.13 |
| SKIP_ON | ENSSSCG00000000007 | 5 | 3213140 | 3213312 | 6.48 | 5.22 | 4.50 | 4.44 | 5.79 | 5.21 | 7.05 | 5.43 | 5.03 | 4.19 | 5.44 | 4.32 |
| TSS | ENSSSCG00000000007 | 5 | 3224232 | 3224387 | 6.48 | 6.48 | 4.50 | 4.50 | 5.79 | 5.79 | 7.05 | 7.05 | 5.03 | 5.03 | 5.44 | 5.44 |
| TTS | ENSSSCG00000000007 | 5 | 3208118 | 3208591 | 6.48 | 6.48 | 4.50 | 4.50 | 5.79 | 5.79 | 7.05 | 7.05 | 5.03 | 5.03 | 5.44 | 5.44 |
| TTS | ENSSSCG00000000010 | 5 | 3873380 | 3874892 | 139.11 | 41.92 | 217.73 | 62.71 | 210.08 | 60.73 | 248.85 | 75.06 | 206.02 | 51.23 | 283.29 | 84.57 |
| TSS | ENSSSCG00000000010 | 5 | 3926979 | 3927217 | 139.11 | 97.19 | 217.73 | 155.03 | 210.08 | 149.35 | 248.85 | 173.79 | 206.02 | 154.78 | 283.29 | 198.72 |
| TTS | ENSSSCG00000000010 | 5 | 3838211 | 3840291 | 139.11 | 97.19 | 217.73 | 155.03 | 210.08 | 149.35 | 248.85 | 173.79 | 206.02 | 154.78 | 283.29 | 198.72 |
| SKIP_OFF | ENSSSCG00000000014 | 5 | 4083099 | 4083131 | 9.67 | 1.00 | 8.83 | 1.96 | 7.82 | 0.36 | 20.51 | 2.15 | 13.39 | 1.17 | 17.52 | 1.42 |
| SKIP_ON | ENSSSCG00000000014 | 5 | 4083099 | 4083131 | 9.67 | 8.67 | 8.83 | 6.87 | 7.82 | 7.46 | 20.51 | 18.37 | 13.39 | 12.22 | 17.52 | 16.10 |
| TSS | ENSSSCG00000000014 | 5 | 4101016 | 4101269 | 9.67 | 9.67 | 8.83 | 8.83 | 7.82 | 7.82 | 20.51 | 20.51 | 13.39 | 13.39 | 17.52 | 17.52 |
| TTS | ENSSSCG00000000014 | 5 | 4070703 | 4071801 | 9.67 | 9.67 | 8.83 | 8.83 | 7.82 | 7.82 | 20.51 | 20.51 | 13.39 | 13.39 | 17.52 | 17.52 |
| TTS | ENSSSCG00000000018 | 5 | 4191887 | 4195613 | 14.72 | 0.00 | 16.73 | 0.30 | 19.00 | 0.38 | 9.78 | 0.16 | 13.77 | 0.00 | 15.01 | 0.00 |
| TSS | ENSSSCG00000000018 | 5 | 4179663 | 4179825 | 14.72 | 4.90 | 16.73 | 3.70 | 19.00 | 6.46 | 9.78 | 2.12 | 13.77 | 4.55 | 15.01 | 5.09 |
| TSS | ENSSSCG00000000018 | 5 | 4179562 | 4179602 | 14.72 | 5.13 | 16.73 | 6.44 | 19.00 | 5.09 | 9.78 | 2.32 | 13.77 | 4.04 | 15.01 | 4.03 |
| TSS | ENSSSCG00000000018 | 5 | 4152072 | 4152540 | 14.72 | 4.69 | 16.73 | 6.59 | 19.00 | 7.45 | 9.78 | 5.33 | 13.77 | 5.18 | 15.01 | 5.90 |
| SKIP_OFF | ENSSSCG00000000019 | 5 | 4212719 | 4212802 | 20.21 | 1.19 | 23.40 | 2.38 | 15.22 | 1.83 | 15.91 | 0.36 | 15.96 | 1.35 | 16.78 | 1.56 |
| SKIP_ON | ENSSSCG00000000019 | 5 | 4212719 | 4212802 | 20.21 | 19.02 | 23.40 | 21.01 | 15.22 | 13.39 | 15.91 | 15.55 | 15.96 | 14.62 | 16.78 | 15.21 |
| TSS | ENSSSCG00000000019 | 5 | 4218932 | 4219415 | 20.21 | 20.21 | 23.40 | 23.40 | 15.22 | 15.22 | 15.91 | 15.91 | 15.96 | 15.96 | 16.78 | 16.78 |
| TTS | ENSSSCG00000000019 | 5 | 4197544 | 4201319 | 20.21 | 20.21 | 23.40 | 23.40 | 15.22 | 15.22 | 15.91 | 15.91 | 15.96 | 15.96 | 16.78 | 16.78 |
| TSS | ENSSSCG00000000020 | 5 | 4291399 | 4291452 | 0.21 | 0.21 | 0.10 | 0.10 | 0.39 | 0.39 | 1.20 | 1.20 | 0.25 | 0.25 | 0.17 | 0.17 |
| TTS | ENSSSCG00000000020 | 5 | 4385028 | 4385246 | 0.21 | 0.21 | 0.10 | 0.10 | 0.39 | 0.39 | 1.20 | 1.20 | 0.25 | 0.25 | 0.17 | 0.17 |
| SKIP_OFF | ENSSSCG00000000021 | 5 | 4412191 | 4412301 | 1.07 | 0.17 | 0.71 | 0.48 | 0.86 | 0.44 | 10.32 | 0.55 | 5.59 | 1.12 | 8.31 | 3.19 |
| SKIP_ON | ENSSSCG00000000021 | 5 | 4412191 | 4412301 | 1.07 | 0.89 | 0.71 | 0.23 | 0.86 | 0.42 | 10.32 | 9.77 | 5.59 | 4.47 | 8.31 | 5.12 |
| TSS | ENSSSCG00000000021 | 5 | 4435218 | 4435341 | 1.07 | 1.07 | 0.71 | 0.71 | 0.86 | 0.86 | 10.32 | 10.32 | 5.59 | 5.59 | 8.31 | 8.31 |
| TTS | ENSSSCG00000000021 | 5 | 4399744 | 4400271 | 1.07 | 1.07 | 0.71 | 0.71 | 0.86 | 0.86 | 10.32 | 10.32 | 5.59 | 5.59 | 8.31 | 8.31 |
| TSS | ENSSSCG00000000024 | 5 | 5015225 | 5015339 | 2.99 | 2.99 | 5.61 | 5.61 | 4.72 | 4.72 | 1.56 | 1.56 | 2.85 | 2.85 | 3.36 | 3.36 |

| | | | | | | | | | | | | | | | | |
|---|---|---|---|---|---|---|---|---|---|---|---|---|---|---|---|---|
| TTS | ENSSSCG00000000024 | 5 | 4909649 | 4909725 | 2.99 | 2.99 | 5.61 | 5.61 | 4.72 | 4.72 | 1.56 | 1.56 | 2.85 | 2.85 | 3.36 | 3.36 |
| TSS | ENSSSCG00000000025 | 5 | 4898421 | 4898706 | 1.20 | 1.15 | 2.39 | 2.31 | 2.25 | 2.25 | 2.75 | 2.64 | 3.42 | 3.33 | 2.04 | 1.95 |
| TTS | ENSSSCG00000000025 | 5 | 4870350 | 4874668 | 1.20 | 1.20 | 2.39 | 2.39 | 2.25 | 2.25 | 2.75 | 2.75 | 3.42 | 3.42 | 2.04 | 2.04 |
| TSS | ENSSSCG00000000026 | 5 | 5131162 | 5131504 | 0.16 | 0.16 | 0.04 | 0.04 | 0.05 | 0.05 | 0.17 | 0.17 | 0.14 | 0.14 | 0.02 | 0.02 |
| TTS | ENSSSCG00000000026 | 5 | 5155495 | 5157857 | 0.16 | 0.16 | 0.04 | 0.04 | 0.05 | 0.05 | 0.17 | 0.17 | 0.14 | 0.14 | 0.02 | 0.02 |
| TSS | ENSSSCG00000000028 | 5 | 5375714 | 5375861 | 0.64 | 0.64 | 0.32 | 0.32 | 0.32 | 0.32 | 5.96 | 5.96 | 3.71 | 3.71 | 4.13 | 4.13 |
| TTS | ENSSSCG00000000028 | 5 | 5407724 | 5408124 | 0.64 | 0.64 | 0.32 | 0.32 | 0.32 | 0.32 | 5.96 | 5.96 | 3.71 | 3.71 | 4.13 | 4.13 |
| SKIP_ON | ENSSSCG00000000029 | 5 | 5635705 | 5635794 | 11.30 | 2.73 | 3.39 | 0.76 | 5.19 | 0.74 | 5.38 | 1.03 | 6.74 | 1.63 | 4.60 | 1.44 |
| XAE | ENSSSCG00000000029 | 5 | 5661874 | 5662010 | 11.30 | 2.73 | 3.39 | 0.76 | 5.19 | 0.74 | 5.38 | 1.03 | 6.74 | 1.63 | 4.60 | 1.44 |
| SKIP_OFF | ENSSSCG00000000029 | 5 | 5635705 | 5635794 | 11.30 | 8.57 | 3.39 | 2.62 | 5.19 | 4.45 | 5.38 | 4.35 | 6.74 | 5.11 | 4.60 | 3.16 |
| XAE | ENSSSCG00000000029 | 5 | 5661874 | 5661996 | 11.30 | 8.57 | 3.39 | 2.62 | 5.19 | 4.45 | 5.38 | 4.35 | 6.74 | 5.11 | 4.60 | 3.16 |
| TSS | ENSSSCG00000000029 | 5 | 5561606 | 5561693 | 11.30 | 11.30 | 3.39 | 3.39 | 5.19 | 5.19 | 5.38 | 5.38 | 6.74 | 6.74 | 4.60 | 4.60 |
| TTS | ENSSSCG00000000029 | 5 | 5686689 | 5686841 | 11.30 | 11.30 | 3.39 | 3.39 | 5.19 | 5.19 | 5.38 | 5.38 | 6.74 | 6.74 | 4.60 | 4.60 |
| TTS | ENSSSCG00000000031 | 5 | 5751451 | 5751692 | 4.58 | 4.00 | 3.06 | 2.72 | 4.85 | 4.28 | 7.54 | 6.64 | 5.44 | 4.81 | 7.81 | 6.71 |
| TSS | ENSSSCG00000000031 | 5 | 5742162 | 5742681 | 4.58 | 4.58 | 3.06 | 3.06 | 4.85 | 4.85 | 7.54 | 7.54 | 5.44 | 5.44 | 7.81 | 7.81 |
| SKIP_OFF | ENSSSCG00000000033 | 5 | 5726251 | 5726461 | 23.05 | 0.00 | 37.66 | 0.00 | 40.57 | 0.00 | 62.79 | 0.00 | 49.57 | 0.00 | 61.56 | 0.00 |
| SKIP_ON | ENSSSCG00000000033 | 5 | 5726251 | 5726461 | 23.05 | 23.05 | 37.66 | 37.66 | 40.57 | 40.57 | 62.79 | 62.79 | 49.57 | 49.57 | 61.56 | 61.56 |
| TSS | ENSSSCG00000000033 | 5 | 5735059 | 5735131 | 23.05 | 23.05 | 37.66 | 37.66 | 40.57 | 40.57 | 62.79 | 62.79 | 49.57 | 49.57 | 61.56 | 61.56 |
| TTS | ENSSSCG00000000033 | 5 | 5723190 | 5723599 | 23.05 | 23.05 | 37.66 | 37.66 | 40.57 | 40.57 | 62.79 | 62.79 | 49.57 | 49.57 | 61.56 | 61.56 |
| TSS | ENSSSCG00000000034 | 5 | 5701532 | 5701799 | 46.67 | 46.67 | 32.17 | 32.17 | 34.27 | 34.27 | 34.43 | 34.43 | 33.22 | 33.22 | 39.13 | 39.13 |
| TTS | ENSSSCG00000000034 | 5 | 5717664 | 5718999 | 46.67 | 46.67 | 32.17 | 32.17 | 34.27 | 34.27 | 34.43 | 34.43 | 33.22 | 33.22 | 39.13 | 39.13 |
| SKIP_OFF | ENSSSCG00000000035 | 5 | 5800250 | 5800322 | 14.07 | 0.00 | 9.73 | 0.00 | 12.01 | 1.19 | 20.08 | 0.18 | 17.23 | 0.60 | 27.30 | 1.18 |
| SKIP_ON | ENSSSCG00000000035 | 5 | 5800250 | 5800322 | 14.07 | 13.98 | 9.73 | 9.59 | 12.01 | 10.73 | 20.08 | 19.76 | 17.23 | 16.46 | 27.30 | 26.03 |
| TSS | ENSSSCG00000000035 | 5 | 5793781 | 5793921 | 14.07 | 13.98 | 9.73 | 9.59 | 12.01 | 11.93 | 20.08 | 19.94 | 17.23 | 17.05 | 27.30 | 27.20 |
| TTS | ENSSSCG00000000035 | 5 | 5827344 | 5827632 | 14.07 | 14.07 | 9.73 | 9.73 | 12.01 | 12.01 | 20.08 | 20.08 | 17.23 | 17.23 | 27.30 | 27.30 |
| TSS | ENSSSCG00000000036 | 5 | 5823811 | 5824711 | 20.25 | 0.33 | 18.21 | 0.07 | 17.29 | 0.27 | 23.95 | 0.52 | 21.38 | 0.51 | 21.02 | 0.43 |
| TSS | ENSSSCG00000000036 | 5 | 5899217 | 5899310 | 20.25 | 2.63 | 18.21 | 0.59 | 17.29 | 1.45 | 23.95 | 4.75 | 21.38 | 2.24 | 21.02 | 0.00 |
| TTS | ENSSSCG00000000036 | 5 | 5984704 | 5985167 | 20.25 | 2.21 | 18.21 | 2.72 | 17.29 | 1.76 | 23.95 | 3.11 | 21.38 | 2.04 | 21.02 | 2.82 |
| SKIP_OFF | ENSSSCG00000000036 | 5 | 5979833 | 5979955 | 20.25 | 4.06 | 18.21 | 1.50 | 17.29 | 1.29 | 23.95 | 5.28 | 21.38 | 5.11 | 21.02 | 3.12 |
| SKIP_ON | ENSSSCG00000000036 | 5 | 5890224 | 5890285 | 20.25 | 6.28 | 18.21 | 4.23 | 17.29 | 3.05 | 23.95 | 8.38 | 21.38 | 7.15 | 21.02 | 5.95 |
| SKIP_OFF | ENSSSCG00000000036 | 5 | 5890224 | 5890285 | 20.25 | 11.01 | 18.21 | 13.32 | 17.29 | 12.52 | 23.95 | 10.30 | 21.38 | 11.48 | 21.02 | 14.64 |
| SKIP_ON | ENSSSCG00000000036 | 5 | 5979833 | 5979955 | 20.25 | 16.19 | 18.21 | 16.71 | 17.29 | 16.00 | 23.95 | 18.67 | 21.38 | 16.26 | 21.02 | 17.90 |
| TSS | ENSSSCG00000000036 | 5 | 5853382 | 5853549 | 20.25 | 17.29 | 18.21 | 17.55 | 17.29 | 15.57 | 23.95 | 18.68 | 21.38 | 18.63 | 21.02 | 20.59 |
| SKIP_OFF | ENSSSCG00000000037 | 5 | 6047749 | 6047870 | 53.00 | 0.12 | 45.96 | 0.34 | 26.79 | 0.12 | 24.78 | 0.00 | 28.97 | 0.23 | 38.26 | 0.11 |
| TSS | ENSSSCG00000000037 | 5 | 5993808 | 5993944 | 53.00 | 0.55 | 45.96 | 1.32 | 26.79 | 0.24 | 24.78 | 0.31 | 28.97 | 0.46 | 38.26 | 0.80 |
| SKIP_OFF | ENSSSCG00000000037 | 5 | 6033636 | 6033767 | 53.00 | 1.80 | 45.96 | 0.81 | 26.79 | 0.96 | 24.78 | 0.59 | 28.97 | 1.41 | 38.26 | 0.26 |
| SKIP_ON | ENSSSCG00000000037 | 5 | 6033636 | 6033767 | 53.00 | 51.20 | 45.96 | 45.15 | 26.79 | 25.83 | 24.78 | 24.19 | 28.97 | 27.56 | 38.26 | 38.00 |
| TSS | ENSSSCG00000000037 | 5 | 5994094 | 5994345 | 53.00 | 52.46 | 45.96 | 44.65 | 26.79 | 26.55 | 24.78 | 24.47 | 28.97 | 28.51 | 38.26 | 37.46 |
| SKIP_ON | ENSSSCG00000000037 | 5 | 6047749 | 6047870 | 53.00 | 52.89 | 45.96 | 45.62 | 26.79 | 26.67 | 24.78 | 24.78 | 28.97 | 28.74 | 38.26 | 38.14 |
| TTS | ENSSSCG00000000037 | 5 | 6049462 | 6049479 | 53.00 | 53.00 | 45.96 | 45.96 | 26.79 | 26.79 | 24.78 | 24.78 | 28.97 | 28.97 | 38.26 | 38.26 |
| TSS | ENSSSCG00000000038 | 5 | 6152922 | 6152942 | 139.14 | 139.14 | 147.27 | 147.27 | 162.53 | 162.53 | 50.47 | 50.47 | 60.85 | 60.85 | 74.59 | 74.59 |
| TTS | ENSSSCG00000000038 | 5 | 6169090 | 6169262 | 139.14 | 139.14 | 147.27 | 147.27 | 162.53 | 162.53 | 50.47 | 50.47 | 60.85 | 60.85 | 74.59 | 74.59 |
| TSS | ENSSSCG00000000039 | 5 | 6179625 | 6182078 | 39.79 | 0.23 | 32.11 | 0.56 | 37.97 | 1.00 | 39.07 | 1.12 | 34.67 | 1.18 | 39.59 | 0.96 |
| SKIP_OFF | ENSSSCG00000000039 | 5 | 6189355 | 6189447 | 39.79 | 1.37 | 32.11 | 2.62 | 37.97 | 0.51 | 39.07 | 1.75 | 34.67 | 0.57 | 39.59 | 0.31 |
| SKIP_ON | ENSSSCG00000000039 | 5 | 6189355 | 6189447 | 39.79 | 38.42 | 32.11 | 29.49 | 37.97 | 37.46 | 39.07 | 37.32 | 34.67 | 34.10 | 39.59 | 39.29 |
| TSS | ENSSSCG00000000039 | 5 | 6174559 | 6174617 | 39.79 | 39.56 | 32.11 | 31.55 | 37.97 | 36.97 | 39.07 | 37.95 | 34.67 | 33.49 | 39.59 | 38.64 |
| TTS | ENSSSCG00000000039 | 5 | 6199634 | 6202081 | 39.79 | 39.79 | 32.11 | 32.11 | 37.97 | 37.97 | 39.07 | 39.07 | 34.67 | 34.67 | 39.59 | 39.59 |
| TTS | ENSSSCG00000000040 | 5 | 6623314 | 6623373 | 0.67 | 0.00 | 0.63 | 0.00 | 0.42 | 0.00 | 0.54 | 0.02 | 0.48 | 0.00 | 0.14 | 0.00 |
| TTS | ENSSSCG00000000040 | 5 | 6634209 | 6635113 | 0.67 | 0.12 | 0.63 | 0.24 | 0.42 | 0.14 | 0.54 | 0.16 | 0.48 | 0.04 | 0.14 | 0.07 |
| TTS | ENSSSCG00000000040 | 5 | 6630062 | 6633091 | 0.67 | 0.56 | 0.63 | 0.27 | 0.42 | 0.24 | 0.54 | 0.28 | 0.48 | 0.40 | 0.14 | 0.07 |

| | | | | | | | | | | | | | | | | |
|---|---|---|---|---|---|---|---|---|---|---|---|---|---|---|---|---|
| TSS | ENSSSCG00000000040 | 5 | 6653892 | 6654438 | 0.67 | 0.67 | 0.63 | 0.51 | 0.42 | 0.38 | 0.54 | 0.46 | 0.48 | 0.44 | 0.14 | 0.14 |
| TSS | ENSSSCG00000000045 | 5 | 6562051 | 6562229 | 17.47 | 17.47 | 24.38 | 24.38 | 22.02 | 22.02 | 16.04 | 16.04 | 14.96 | 14.96 | 16.97 | 16.97 |
| TTS | ENSSSCG00000000045 | 5 | 6566714 | 6567647 | 17.47 | 17.47 | 24.38 | 24.38 | 22.02 | 22.02 | 16.04 | 16.04 | 14.96 | 14.96 | 16.97 | 16.97 |
| TSS | ENSSSCG00000000046 | 5 | 6538907 | 6539274 | 1.29 | 1.29 | 0.26 | 0.26 | 0.78 | 0.78 | 1.99 | 1.99 | 0.22 | 0.22 | 0.48 | 0.48 |
| TTS | ENSSSCG00000000046 | 5 | 6543392 | 6543746 | 1.29 | 1.29 | 0.26 | 0.26 | 0.78 | 0.78 | 1.99 | 1.99 | 0.22 | 0.22 | 0.48 | 0.48 |
| TSS | ENSSSCG00000000050 | 5 | 6579988 | 6581440 | 23.37 | 2.33 | 24.73 | 1.57 | 34.59 | 4.33 | 33.56 | 2.80 | 30.41 | 4.30 | 40.19 | 3.30 |
| SKIP_ON | ENSSSCG00000000050 | 5 | 6579322 | 6579652 | 23.37 | 4.96 | 24.73 | 4.73 | 34.59 | 7.63 | 33.56 | 6.40 | 30.41 | 6.20 | 40.19 | 6.11 |
| TSS | ENSSSCG00000000050 | 5 | 6580141 | 6580156 | 23.37 | 4.73 | 24.73 | 7.50 | 34.59 | 7.93 | 33.56 | 12.08 | 30.41 | 8.93 | 40.19 | 11.92 |
| SKIP_OFF | ENSSSCG00000000050 | 5 | 6579322 | 6579652 | 23.37 | 11.35 | 24.73 | 10.92 | 34.59 | 14.71 | 33.56 | 12.29 | 30.41 | 10.98 | 40.19 | 18.84 |
| TSS | ENSSSCG00000000050 | 5 | 6571005 | 6571411 | 23.37 | 16.31 | 24.73 | 15.66 | 34.59 | 22.34 | 33.56 | 18.69 | 30.41 | 17.18 | 40.19 | 24.96 |
| TTS | ENSSSCG00000000050 | 5 | 6587416 | 6588054 | 23.37 | 23.37 | 24.73 | 24.73 | 34.59 | 34.59 | 33.56 | 33.56 | 30.41 | 30.41 | 40.19 | 40.19 |
| XSKIP_ON | ENSSSCG00000000058 | 5 | 6891302 | 6891452 | 95.20 | 0.10 | 85.87 | 0.00 | 67.38 | 0.10 | 84.05 | 0.00 | 78.20 | 0.00 | 83.71 | 0.32 |
| TTS | ENSSSCG00000000058 | 5 | 6899232 | 6900015 | 95.20 | 4.99 | 85.87 | 3.11 | 67.38 | 1.34 | 84.05 | 4.35 | 78.20 | 4.46 | 83.71 | 4.58 |
| TSS | ENSSSCG00000000058 | 5 | 6896168 | 6896305 | 95.20 | 15.61 | 85.87 | 14.01 | 67.38 | 7.38 | 84.05 | 5.59 | 78.20 | 10.98 | 83.71 | 14.12 |
| XSKIP_OFF | ENSSSCG00000000058 | 5 | 6891302 | 6891452 | 95.20 | 79.49 | 85.87 | 71.86 | 67.38 | 59.90 | 84.05 | 78.46 | 78.20 | 67.22 | 83.71 | 69.27 |
| TSS | ENSSSCG00000000058 | 5 | 6891002 | 6891135 | 95.20 | 79.59 | 85.87 | 71.86 | 67.38 | 60.00 | 84.05 | 78.46 | 78.20 | 67.22 | 83.71 | 69.59 |
| TSS | ENSSSCG00000000059 | 5 | 6928856 | 6929220 | 9.51 | 0.73 | 7.82 | 0.09 | 8.61 | 0.16 | 10.44 | 1.22 | 10.73 | 1.78 | 14.42 | 3.12 |
| TTS | ENSSSCG00000000059 | 5 | 6899725 | 6903905 | 9.51 | 9.51 | 7.82 | 7.82 | 8.61 | 8.61 | 10.44 | 10.44 | 10.73 | 10.73 | 14.42 | 14.42 |
| TSS | ENSSSCG00000000061 | 5 | 6964999 | 6965417 | 12.66 | 12.66 | 12.75 | 12.75 | 11.91 | 11.91 | 6.67 | 6.67 | 13.21 | 13.21 | 10.99 | 10.99 |
| TTS | ENSSSCG00000000061 | 5 | 6975371 | 6975904 | 12.66 | 12.66 | 12.75 | 12.75 | 11.91 | 11.91 | 6.67 | 6.67 | 13.21 | 13.21 | 10.99 | 10.99 |
| TSS | ENSSSCG00000000062 | 5 | 6988230 | 6988352 | 69.29 | 69.29 | 77.23 | 77.23 | 35.14 | 35.14 | 13.00 | 13.00 | 48.53 | 48.53 | 20.19 | 20.19 |
| TTS | ENSSSCG00000000062 | 5 | 6976277 | 6977934 | 69.29 | 69.29 | 77.23 | 77.23 | 35.14 | 35.14 | 13.00 | 13.00 | 48.53 | 48.53 | 20.19 | 20.19 |
| SKIP_OFF | ENSSSCG00000000063 | 5 | 7010279 | 7010365 | 6.21 | 0.00 | 5.15 | 0.78 | 6.85 | 3.44 | 5.91 | 1.44 | 6.07 | 0.63 | 5.46 | 0.61 |
| SKIP_ON | ENSSSCG00000000063 | 5 | 7010279 | 7010365 | 6.21 | 6.21 | 5.15 | 4.37 | 6.85 | 3.41 | 5.91 | 4.47 | 6.07 | 5.44 | 5.46 | 4.85 |
| TSS | ENSSSCG00000000063 | 5 | 6999981 | 7000890 | 6.21 | 6.21 | 5.15 | 5.15 | 6.85 | 6.85 | 5.91 | 5.91 | 6.07 | 6.07 | 5.46 | 5.46 |
| TTS | ENSSSCG00000000063 | 5 | 7013122 | 7014430 | 6.21 | 6.21 | 5.15 | 5.15 | 6.85 | 6.85 | 5.91 | 5.91 | 6.07 | 6.07 | 5.46 | 5.46 |
| TSS | ENSSSCG00000000064 | 5 | 7070959 | 7071023 | 46.45 | 46.45 | 43.76 | 43.76 | 46.98 | 46.98 | 47.44 | 47.44 | 41.86 | 41.86 | 57.68 | 57.68 |
| TTS | ENSSSCG00000000064 | 5 | 7014383 | 7014812 | 46.45 | 46.45 | 43.76 | 43.76 | 46.98 | 46.98 | 47.44 | 47.44 | 41.86 | 41.86 | 57.68 | 57.68 |
| TSS | ENSSSCG00000000066 | 5 | 7277488 | 7277678 | 3.69 | 3.69 | 2.93 | 2.93 | 3.39 | 3.39 | 3.32 | 3.32 | 2.97 | 2.97 | 4.06 | 4.06 |
| TTS | ENSSSCG00000000066 | 5 | 7253385 | 7258495 | 3.69 | 3.69 | 2.93 | 2.93 | 3.39 | 3.39 | 3.32 | 3.32 | 2.97 | 2.97 | 4.06 | 4.06 |
| MIR_ON | ENSSSCG00000000067 | 5 | 7250059 | 7250263 | 0.65 | 0.00 | 0.42 | 0.00 | 1.20 | 0.00 | 0.95 | 0.00 | 1.36 | 0.00 | 1.06 | 0.00 |
| TSS | ENSSSCG00000000067 | 5 | 7246700 | 7246728 | 0.65 | 0.00 | 0.42 | 0.00 | 1.20 | 0.00 | 0.95 | 0.00 | 1.36 | 0.00 | 1.06 | 0.00 |
| MIR_OFF | ENSSSCG00000000067 | 5 | 7250059 | 7250263 | 0.65 | 0.65 | 0.42 | 0.42 | 1.20 | 1.20 | 0.95 | 0.95 | 1.36 | 1.36 | 1.06 | 1.06 |
| TTS | ENSSSCG00000000067 | 5 | 7252705 | 7252936 | 0.65 | 0.65 | 0.42 | 0.42 | 1.20 | 1.20 | 0.95 | 0.95 | 1.36 | 1.36 | 1.06 | 1.06 |
| TTS | ENSSSCG00000000068 | 5 | 7312262 | 7312843 | 13.03 | 0.00 | 15.91 | 0.26 | 14.21 | 0.12 | 15.36 | 0.00 | 14.40 | 0.00 | 12.65 | 0.00 |
| TTS | ENSSSCG00000000068 | 5 | 7312262 | 7314457 | 13.03 | 13.03 | 15.91 | 15.66 | 14.21 | 14.09 | 15.36 | 15.36 | 14.40 | 14.40 | 12.65 | 12.65 |
| TSS | ENSSSCG00000000068 | 5 | 7373368 | 7373440 | 13.03 | 13.03 | 15.91 | 15.91 | 14.21 | 14.21 | 15.36 | 15.36 | 14.40 | 14.40 | 12.65 | 12.65 |
| AE | ENSSSCG00000000071 | 5 | 7614532 | 7614665 | 123.98 | 54.26 | 120.04 | 56.49 | 130.97 | 60.53 | 134.62 | 54.89 | 120.37 | 52.26 | 120.28 | 54.15 |
| TSS | ENSSSCG00000000071 | 5 | 7583048 | 7583427 | 123.98 | 54.26 | 120.04 | 56.49 | 130.97 | 60.53 | 134.62 | 54.89 | 120.37 | 52.26 | 120.28 | 54.15 |
| TTS | ENSSSCG00000000071 | 5 | 7616178 | 7616382 | 123.98 | 54.26 | 120.04 | 56.49 | 130.97 | 60.53 | 134.62 | 54.89 | 120.37 | 52.26 | 120.28 | 54.15 |
| AE | ENSSSCG00000000071 | 5 | 7614556 | 7614665 | 123.98 | 69.72 | 120.04 | 63.55 | 130.97 | 70.45 | 134.62 | 79.74 | 120.37 | 68.11 | 120.28 | 66.13 |
| TSS | ENSSSCG00000000071 | 5 | 7583095 | 7583605 | 123.98 | 69.72 | 120.04 | 63.55 | 130.97 | 70.45 | 134.62 | 79.74 | 120.37 | 68.11 | 120.28 | 66.13 |
| MSKIP_OFF | ENSSSCG00000000072 | 5 | 7647344 | 7649331 | 33.55 | 7.08 | 39.33 | 9.02 | 38.61 | 3.29 | 12.00 | 3.00 | 19.71 | 2.69 | 27.88 | 5.22 |
| TTS | ENSSSCG00000000072 | 5 | 7671247 | 7671931 | 33.55 | 9.86 | 39.33 | 15.76 | 38.61 | 17.22 | 12.00 | 4.85 | 19.71 | 8.03 | 27.88 | 11.57 |
| MSKIP_ON | ENSSSCG00000000072 | 5 | 7647344 | 7649331 | 33.55 | 9.97 | 39.33 | 15.82 | 38.61 | 17.50 | 12.00 | 5.02 | 19.71 | 8.08 | 27.88 | 11.69 |
| MSKIP_ON | ENSSSCG00000000072 | 5 | 7643801 | 7647410 | 33.55 | 9.97 | 39.33 | 15.82 | 38.61 | 17.50 | 12.00 | 5.02 | 19.71 | 8.08 | 27.88 | 11.69 |
| MSKIP_OFF | ENSSSCG00000000072 | 5 | 7643801 | 7647410 | 33.55 | 16.50 | 39.33 | 14.50 | 38.61 | 17.81 | 12.00 | 3.98 | 19.71 | 8.93 | 27.88 | 10.97 |
| TSS | ENSSSCG00000000072 | 5 | 7622247 | 7622833 | 33.55 | 33.55 | 39.33 | 39.33 | 38.61 | 38.61 | 12.00 | 12.00 | 19.71 | 19.71 | 27.88 | 27.88 |
| TSS | ENSSSCG00000000073 | 5 | 7673983 | 7673997 | 0.65 | 0.65 | 0.79 | 0.79 | 0.16 | 0.16 | 0.00 | 0.00 | 0.00 | 0.00 | 0.00 | 0.00 |
| TTS | ENSSSCG00000000073 | 5 | 7674834 | 7675196 | 0.65 | 0.65 | 0.79 | 0.79 | 0.16 | 0.16 | 0.00 | 0.00 | 0.00 | 0.00 | 0.00 | 0.00 |

| | | | | | | | | | | | | | | | | |
|---|---|---|---|---|---|---|---|---|---|---|---|---|---|---|---|---|
| TSS | ENSSSCG00000000074 | 5 | 7738489 | 7738847 | 0.22 | 0.00 | 0.56 | 0.00 | 0.23 | 0.00 | 0.03 | 0.00 | 0.00 | 0.00 | 0.09 | 0.00 |
| TSS | ENSSSCG00000000074 | 5 | 7737483 | 7737597 | 0.22 | 0.00 | 0.56 | 0.00 | 0.23 | 0.00 | 0.03 | 0.00 | 0.00 | 0.00 | 0.09 | 0.00 |
| TTS | ENSSSCG00000000074 | 5 | 7734250 | 7737291 | 0.22 | 0.00 | 0.56 | 0.00 | 0.23 | 0.00 | 0.03 | 0.00 | 0.00 | 0.00 | 0.09 | 0.00 |
| TSS | ENSSSCG00000000075 | 5 | 7933873 | 7935363 | 8.26 | 0.07 | 7.99 | 0.00 | 14.93 | 0.17 | 8.09 | 0.18 | 7.19 | 0.29 | 10.60 | 0.07 |
| TSS | ENSSSCG00000000075 | 5 | 7874347 | 7874622 | 8.26 | 1.08 | 7.99 | 0.52 | 14.93 | 6.76 | 8.09 | 0.49 | 7.19 | 0.42 | 10.60 | 1.79 |
| SKIP_OFF | ENSSSCG00000000075 | 5 | 7813518 | 7813579 | 8.26 | 2.12 | 7.99 | 3.63 | 14.93 | 2.76 | 8.09 | 2.11 | 7.19 | 1.34 | 10.60 | 3.01 |
| SKIP_ON | ENSSSCG00000000075 | 5 | 7813518 | 7813579 | 8.26 | 5.00 | 7.99 | 3.84 | 14.93 | 5.23 | 8.09 | 5.30 | 7.19 | 5.14 | 10.60 | 5.74 |
| TSS | ENSSSCG00000000075 | 5 | 7777044 | 7777303 | 8.26 | 7.11 | 7.99 | 7.47 | 14.93 | 7.99 | 8.09 | 7.41 | 7.19 | 6.48 | 10.60 | 8.75 |
| TTS | ENSSSCG00000000075 | 5 | 7974429 | 7975923 | 8.26 | 8.26 | 7.99 | 7.99 | 14.93 | 14.93 | 8.09 | 8.09 | 7.19 | 7.19 | 10.60 | 10.60 |
| SKIP_OFF | ENSSSCG00000000076 | 5 | 7976847 | 7976907 | 6.87 | 0.20 | 5.74 | 0.00 | 7.59 | 0.00 | 8.06 | 0.00 | 7.12 | 0.00 | 10.59 | 0.00 |
| SKIP_ON | ENSSSCG00000000076 | 5 | 7976847 | 7976907 | 6.87 | 6.67 | 5.74 | 5.74 | 7.59 | 7.59 | 8.06 | 8.06 | 7.12 | 7.12 | 10.59 | 10.59 |
| TSS | ENSSSCG00000000076 | 5 | 8017664 | 8017703 | 6.87 | 6.87 | 5.74 | 5.74 | 7.59 | 7.59 | 8.06 | 8.06 | 7.12 | 7.12 | 10.59 | 10.59 |
| TTS | ENSSSCG00000000076 | 5 | 7976099 | 7976603 | 6.87 | 6.87 | 5.74 | 5.74 | 7.59 | 7.59 | 8.06 | 8.06 | 7.12 | 7.12 | 10.59 | 10.59 |
| TSS | ENSSSCG00000000077 | 5 | 8045310 | 8045528 | 12.22 | 12.22 | 9.97 | 9.97 | 13.44 | 13.44 | 14.19 | 14.19 | 14.02 | 14.02 | 14.44 | 14.44 |
| TTS | ENSSSCG00000000077 | 5 | 8027588 | 8027885 | 12.22 | 12.22 | 9.97 | 9.97 | 13.44 | 13.44 | 14.19 | 14.19 | 14.02 | 14.02 | 14.44 | 14.44 |
| MSKIP_ON | ENSSSCG00000000078 | 5 | 8098068 | 8102846 | 2.45 | 0.00 | 3.29 | 0.00 | 3.40 | 0.00 | 3.94 | 0.00 | 4.60 | 0.01 | 3.88 | 0.00 |
| SKIP_ON | ENSSSCG00000000078 | 5 | 8102676 | 8102846 | 2.45 | 0.00 | 3.29 | 0.00 | 3.40 | 0.00 | 3.94 | 0.00 | 4.60 | 0.01 | 3.88 | 0.00 |
| TSS | ENSSSCG00000000078 | 5 | 8242528 | 8242601 | 2.45 | 0.00 | 3.29 | 0.00 | 3.40 | 0.00 | 3.94 | 0.00 | 4.60 | 0.01 | 3.88 | 0.00 |
| TSS | ENSSSCG00000000078 | 5 | 8175608 | 8175663 | 2.45 | 0.00 | 3.29 | 0.00 | 3.40 | 0.00 | 3.94 | 0.00 | 4.60 | 0.00 | 3.88 | 0.00 |
| MSKIP_OFF | ENSSSCG00000000078 | 5 | 8098068 | 8102846 | 2.45 | 0.16 | 3.29 | 0.27 | 3.40 | 0.00 | 3.94 | 0.45 | 4.60 | 0.18 | 3.88 | 0.49 |
| SKIP_OFF | ENSSSCG00000000078 | 5 | 8098068 | 8098193 | 2.45 | 0.16 | 3.29 | 0.27 | 3.40 | 0.00 | 3.94 | 0.45 | 4.60 | 0.18 | 3.88 | 0.49 |
| TSS | ENSSSCG00000000078 | 5 | 8202378 | 8202390 | 2.45 | 0.59 | 3.29 | 0.65 | 3.40 | 0.44 | 3.94 | 0.37 | 4.60 | 0.71 | 3.88 | 0.77 |
| TTS | ENSSSCG00000000078 | 5 | 8064075 | 8064129 | 2.45 | 0.59 | 3.29 | 0.65 | 3.40 | 0.44 | 3.94 | 0.37 | 4.60 | 0.71 | 3.88 | 0.77 |
| TSS | ENSSSCG00000000078 | 5 | 8311038 | 8311105 | 2.45 | 0.54 | 3.29 | 0.86 | 3.40 | 1.05 | 3.94 | 0.50 | 4.60 | 1.02 | 3.88 | 0.59 |
| TSS | ENSSSCG00000000078 | 5 | 8200252 | 8200256 | 2.45 | 1.32 | 3.29 | 1.79 | 3.40 | 1.92 | 3.94 | 3.08 | 4.60 | 2.86 | 3.88 | 2.52 |
| SKIP_OFF | ENSSSCG00000000078 | 5 | 8102676 | 8102846 | 2.45 | 2.29 | 3.29 | 3.03 | 3.40 | 3.40 | 3.94 | 3.50 | 4.60 | 4.41 | 3.88 | 3.39 |
| SKIP_ON | ENSSSCG00000000078 | 5 | 8098068 | 8098193 | 2.45 | 2.29 | 3.29 | 3.03 | 3.40 | 3.40 | 3.94 | 3.50 | 4.60 | 4.41 | 3.88 | 3.39 |
| TSS | ENSSSCG00000000079 | 5 | 8354168 | 8354723 | 0.31 | 0.31 | 0.07 | 0.07 | 0.07 | 0.07 | 1.70 | 1.70 | 1.56 | 1.56 | 2.81 | 2.81 |
| TTS | ENSSSCG00000000079 | 5 | 8320978 | 8325458 | 0.31 | 0.31 | 0.07 | 0.07 | 0.07 | 0.07 | 1.70 | 1.70 | 1.56 | 1.56 | 2.81 | 2.81 |
| TSS | ENSSSCG00000000080 | 5 | 8444825 | 8445071 | 0.31 | 0.04 | 0.28 | 0.02 | 0.10 | 0.01 | 0.92 | 0.10 | 0.99 | 0.15 | 0.57 | 0.13 |
| TTS | ENSSSCG00000000080 | 5 | 8356286 | 8358681 | 0.31 | 0.04 | 0.28 | 0.02 | 0.10 | 0.01 | 0.92 | 0.10 | 0.99 | 0.15 | 0.57 | 0.13 |
| TSS | ENSSSCG00000000080 | 5 | 8417023 | 8417248 | 0.31 | 0.27 | 0.28 | 0.26 | 0.10 | 0.09 | 0.92 | 0.82 | 0.99 | 0.83 | 0.57 | 0.43 |
| TSS | ENSSSCG00000000082 | 5 | 8700978 | 8701228 | 0.33 | 0.33 | 0.20 | 0.20 | 0.92 | 0.92 | 0.47 | 0.47 | 0.29 | 0.29 | 0.22 | 0.22 |
| TTS | ENSSSCG00000000082 | 5 | 8597053 | 8597560 | 0.33 | 0.33 | 0.20 | 0.20 | 0.92 | 0.92 | 0.47 | 0.47 | 0.29 | 0.29 | 0.22 | 0.22 |
| TSS | ENSSSCG00000000083 | 5 | 8735382 | 8735515 | 27.22 | 27.22 | 20.69 | 20.69 | 22.37 | 22.37 | 20.60 | 20.60 | 20.55 | 20.55 | 28.30 | 28.30 |
| TTS | ENSSSCG00000000083 | 5 | 8738352 | 8738817 | 27.22 | 27.22 | 20.69 | 20.69 | 22.37 | 22.37 | 20.60 | 20.60 | 20.55 | 20.55 | 28.30 | 28.30 |
| TTS | ENSSSCG00000000084 | 5 | 8738880 | 8743669 | 198.68 | 1.22 | 214.54 | 1.39 | 153.47 | 1.02 | 100.85 | 0.72 | 95.19 | 0.60 | 129.63 | 0.61 |
| TSS | ENSSSCG00000000084 | 5 | 8744854 | 8745368 | 198.68 | 197.46 | 214.54 | 213.14 | 153.47 | 152.45 | 100.85 | 100.13 | 95.19 | 94.58 | 129.63 | 129.02 |
| TTS | ENSSSCG00000000084 | 5 | 8738880 | 8743867 | 198.68 | 197.46 | 214.54 | 213.14 | 153.47 | 152.45 | 100.85 | 100.13 | 95.19 | 94.58 | 129.63 | 129.02 |
| TSS | ENSSSCG00000000087 | 5 | 8852688 | 8852850 | 5.47 | 5.47 | 4.72 | 4.61 | 6.46 | 6.28 | 6.21 | 6.21 | 5.22 | 5.13 | 6.43 | 6.43 |
| TTS | ENSSSCG00000000087 | 5 | 8824675 | 8829684 | 5.47 | 5.47 | 4.72 | 4.72 | 6.46 | 6.46 | 6.21 | 6.21 | 5.22 | 5.22 | 6.43 | 6.43 |
| TSS | ENSSSCG00000000089 | 5 | 8927875 | 8927953 | 137.73 | 137.73 | 125.97 | 125.97 | 102.19 | 102.19 | 117.65 | 117.65 | 109.98 | 109.98 | 127.46 | 127.46 |
| TTS | ENSSSCG00000000089 | 5 | 8923674 | 8923910 | 137.73 | 137.73 | 125.97 | 125.97 | 102.19 | 102.19 | 117.65 | 117.65 | 109.98 | 109.98 | 127.46 | 127.46 |
| TSS | ENSSSCG00000000090 | 5 | 9070353 | 9070603 | 2.17 | 2.17 | 4.09 | 4.09 | 4.01 | 4.01 | 6.81 | 6.81 | 5.46 | 5.46 | 3.24 | 3.24 |
| TTS | ENSSSCG00000000090 | 5 | 9087316 | 9087634 | 2.17 | 2.17 | 4.09 | 4.09 | 4.01 | 4.01 | 6.81 | 6.81 | 5.46 | 5.46 | 3.24 | 3.24 |
| MSKIP_OFF | ENSSSCG00000000091 | 5 | 9100881 | 9110036 | 5.63 | 0.00 | 3.95 | 0.00 | 5.48 | 0.00 | 6.90 | 0.00 | 10.55 | 0.49 | 9.36 | 0.42 |
| TSS | ENSSSCG00000000091 | 5 | 9105621 | 9105778 | 5.63 | 0.00 | 3.95 | 0.13 | 5.48 | 0.00 | 6.90 | 0.68 | 10.55 | 1.36 | 9.36 | 0.85 |
| TTS | ENSSSCG00000000091 | 5 | 9108471 | 9108533 | 5.63 | 0.15 | 3.95 | 0.00 | 5.48 | 0.00 | 6.90 | 0.02 | 10.55 | 0.00 | 9.36 | 0.62 |
| TTS | ENSSSCG00000000091 | 5 | 9108164 | 9108746 | 5.63 | 1.10 | 3.95 | 0.67 | 5.48 | 1.00 | 6.90 | 0.87 | 10.55 | 1.31 | 9.36 | 0.95 |
| MSKIP_ON | ENSSSCG00000000091 | 5 | 9100881 | 9110036 | 5.63 | 0.86 | 3.95 | 0.45 | 5.48 | 1.63 | 6.90 | 2.37 | 10.55 | 2.12 | 9.36 | 2.44 |

| | | | | | | | | | | | | | | | | |
|---|---|---|---|---|---|---|---|---|---|---|---|---|---|---|---|---|
| XSKIP_ON | ENSSSCG00000000091 | 5 | 9105854 | 9105917 | 5.63 | 0.86 | 3.95 | 0.45 | 5.48 | 1.63 | 6.90 | 2.37 | 10.55 | 2.12 | 9.36 | 2.44 |
| TSS | ENSSSCG00000000091 | 5 | 9107252 | 9107471 | 5.63 | 1.17 | 3.95 | 0.64 | 5.48 | 1.60 | 6.90 | 1.41 | 10.55 | 3.43 | 9.36 | 2.53 |
| MSKIP_ON | ENSSSCG00000000091 | 5 | 9100881 | 9110036 | 5.63 | 2.35 | 3.95 | 2.06 | 5.48 | 1.25 | 6.90 | 1.55 | 10.55 | 1.84 | 9.36 | 1.54 |
| XSKIP_OFF | ENSSSCG00000000091 | 5 | 9105854 | 9105917 | 5.63 | 2.35 | 3.95 | 2.06 | 5.48 | 1.25 | 6.90 | 1.55 | 10.55 | 1.84 | 9.36 | 1.54 |
| TSS | ENSSSCG00000000091 | 5 | 9115817 | 9116450 | 5.63 | 4.46 | 3.95 | 3.18 | 5.48 | 3.88 | 6.90 | 4.82 | 10.55 | 5.76 | 9.36 | 5.98 |
| TSS | ENSSSCG00000000092 | 5 | 9207671 | 9208255 | 0.53 | 0.53 | 0.26 | 0.26 | 0.28 | 0.28 | 2.90 | 2.90 | 0.45 | 0.45 | 1.14 | 1.14 |
| TTS | ENSSSCG00000000092 | 5 | 9226160 | 9230351 | 0.53 | 0.53 | 0.26 | 0.26 | 0.28 | 0.28 | 2.90 | 2.90 | 0.45 | 0.45 | 1.14 | 1.14 |
| TSS | ENSSSCG00000000093 | 5 | 9239960 | 9244886 | 14.27 | 0.81 | 7.79 | 0.48 | 11.44 | 0.91 | 40.66 | 2.18 | 32.50 | 3.10 | 46.27 | 3.38 |
| TSS | ENSSSCG00000000093 | 5 | 9236625 | 9236733 | 14.27 | 13.46 | 7.79 | 7.32 | 11.44 | 10.54 | 40.66 | 38.48 | 32.50 | 29.40 | 46.27 | 42.89 |
| TTS | ENSSSCG00000000093 | 5 | 9247707 | 9248726 | 14.27 | 14.27 | 7.79 | 7.79 | 11.44 | 11.44 | 40.66 | 40.66 | 32.50 | 32.50 | 46.27 | 46.27 |
| SKIP_OFF | ENSSSCG00000000094 | 5 | 9279042 | 9279137 | 50.52 | 0.00 | 42.51 | 0.32 | 42.86 | 0.00 | 41.05 | 0.00 | 59.27 | 0.81 | 41.16 | 0.39 |
| SKIP_ON | ENSSSCG00000000094 | 5 | 9275767 | 9275881 | 50.52 | 0.20 | 42.51 | 0.53 | 42.86 | 0.17 | 41.05 | 1.54 | 59.27 | 0.39 | 41.16 | 0.00 |
| TSS | ENSSSCG00000000094 | 5 | 9273228 | 9273407 | 50.52 | 8.71 | 42.51 | 6.52 | 42.86 | 7.26 | 41.05 | 13.68 | 59.27 | 12.88 | 41.16 | 11.38 |
| TSS | ENSSSCG00000000094 | 5 | 9273131 | 9273493 | 50.52 | 41.81 | 42.51 | 35.98 | 42.86 | 35.60 | 41.05 | 27.37 | 59.27 | 46.39 | 41.16 | 29.78 |
| SKIP_OFF | ENSSSCG00000000094 | 5 | 9275767 | 9275881 | 50.52 | 50.32 | 42.51 | 41.98 | 42.86 | 42.69 | 41.05 | 39.51 | 59.27 | 58.88 | 41.16 | 41.16 |
| SKIP_ON | ENSSSCG00000000094 | 5 | 9279042 | 9279137 | 50.52 | 50.52 | 42.51 | 42.19 | 42.86 | 42.86 | 41.05 | 41.05 | 59.27 | 58.46 | 41.16 | 40.78 |
| TTS | ENSSSCG00000000094 | 5 | 9291646 | 9293255 | 50.52 | 50.52 | 42.51 | 42.51 | 42.86 | 42.86 | 41.05 | 41.05 | 59.27 | 59.27 | 41.16 | 41.16 |
| XAE | ENSSSCG00000000095 | 5 | 9316163 | 9318717 | 22.71 | 1.75 | 20.81 | 1.97 | 18.42 | 1.98 | 20.11 | 1.80 | 19.17 | 1.96 | 20.67 | 2.03 |
| XSKIP_ON | ENSSSCG00000000095 | 5 | 9316163 | 9318717 | 22.71 | 1.75 | 20.81 | 1.97 | 18.42 | 1.98 | 20.11 | 1.80 | 19.17 | 1.96 | 20.67 | 2.03 |
| XSKIP_ON | ENSSSCG00000000095 | 5 | 9319847 | 9319874 | 22.71 | 1.75 | 20.81 | 1.97 | 18.42 | 1.98 | 20.11 | 1.80 | 19.17 | 1.96 | 20.67 | 2.03 |
| AE | ENSSSCG00000000095 | 5 | 9319707 | 9319874 | 22.71 | 2.95 | 20.81 | 2.11 | 18.42 | 1.65 | 20.11 | 2.00 | 19.17 | 2.31 | 20.67 | 1.24 |
| TSS | ENSSSCG00000000095 | 5 | 9321646 | 9321899 | 22.71 | 2.95 | 20.81 | 2.11 | 18.42 | 1.65 | 20.11 | 2.00 | 19.17 | 2.31 | 20.67 | 1.24 |
| XAE | ENSSSCG00000000095 | 5 | 9319707 | 9319874 | 22.71 | 2.95 | 20.81 | 2.11 | 18.42 | 1.65 | 20.11 | 2.00 | 19.17 | 2.31 | 20.67 | 1.24 |
| SKIP_OFF | ENSSSCG00000000095 | 5 | 9319847 | 9319874 | 22.71 | 3.08 | 20.81 | 2.63 | 18.42 | 1.10 | 20.11 | 2.39 | 19.17 | 1.62 | 20.67 | 3.01 |
| XSKIP_OFF | ENSSSCG00000000095 | 5 | 9319847 | 9319874 | 22.71 | 3.08 | 20.81 | 2.63 | 18.42 | 1.10 | 20.11 | 2.39 | 19.17 | 1.62 | 20.67 | 3.01 |
| TSS | ENSSSCG00000000095 | 5 | 9318940 | 9319874 | 22.71 | 2.85 | 20.81 | 3.13 | 18.42 | 3.68 | 20.11 | 3.41 | 19.17 | 3.15 | 20.67 | 3.03 |
| TTS | ENSSSCG00000000095 | 5 | 9247203 | 9247289 | 22.71 | 2.85 | 20.81 | 3.13 | 18.42 | 3.68 | 20.11 | 3.41 | 19.17 | 3.15 | 20.67 | 3.03 |
| SKIP_ON | ENSSSCG00000000095 | 5 | 9316163 | 9316274 | 22.71 | 8.89 | 20.81 | 7.86 | 18.42 | 6.43 | 20.11 | 7.80 | 19.17 | 7.08 | 20.67 | 7.29 |
| XAE | ENSSSCG00000000095 | 5 | 9316163 | 9316274 | 22.71 | 8.89 | 20.81 | 7.86 | 18.42 | 6.43 | 20.11 | 7.80 | 19.17 | 7.08 | 20.67 | 7.29 |
| SKIP_OFF | ENSSSCG00000000095 | 5 | 9316163 | 9316274 | 22.71 | 12.08 | 20.81 | 10.98 | 18.42 | 10.01 | 20.11 | 10.51 | 19.17 | 10.12 | 20.67 | 11.34 |
| SKIP_ON | ENSSSCG00000000095 | 5 | 9319847 | 9319874 | 22.71 | 12.08 | 20.81 | 10.98 | 18.42 | 10.01 | 20.11 | 10.51 | 19.17 | 10.12 | 20.67 | 11.34 |
| XSKIP_OFF | ENSSSCG00000000095 | 5 | 9316163 | 9318717 | 22.71 | 12.08 | 20.81 | 10.98 | 18.42 | 10.01 | 20.11 | 10.51 | 19.17 | 10.12 | 20.67 | 11.34 |
| AE | ENSSSCG00000000095 | 5 | 9319847 | 9319874 | 22.71 | 13.83 | 20.81 | 12.95 | 18.42 | 11.99 | 20.11 | 12.31 | 19.17 | 12.09 | 20.67 | 13.38 |
| XAE | ENSSSCG00000000095 | 5 | 9319847 | 9319874 | 22.71 | 13.83 | 20.81 | 12.95 | 18.42 | 11.99 | 20.11 | 12.31 | 19.17 | 12.09 | 20.67 | 13.38 |
| TSS | ENSSSCG00000000095 | 5 | 9322033 | 9322127 | 22.71 | 16.91 | 20.81 | 15.57 | 18.42 | 13.09 | 20.11 | 14.70 | 19.17 | 13.71 | 20.67 | 16.39 |
| TSS | ENSSSCG00000000097 | 5 | 9357847 | 9358142 | 45.08 | 45.08 | 37.06 | 37.06 | 47.38 | 47.38 | 48.48 | 48.48 | 46.03 | 46.03 | 62.40 | 62.40 |
| TTS | ENSSSCG00000000097 | 5 | 9336129 | 9336848 | 45.08 | 45.08 | 37.06 | 37.06 | 47.38 | 47.38 | 48.48 | 48.48 | 46.03 | 46.03 | 62.40 | 62.40 |
| TSS | ENSSSCG00000000103 | 5 | 9433058 | 9433109 | 0.04 | 0.04 | 0.00 | 0.00 | 0.04 | 0.04 | 0.28 | 0.28 | 0.22 | 0.22 | 0.16 | 0.16 |
| TTS | ENSSSCG00000000103 | 5 | 9469247 | 9470549 | 0.04 | 0.04 | 0.00 | 0.00 | 0.04 | 0.04 | 0.28 | 0.28 | 0.22 | 0.22 | 0.16 | 0.16 |
| IR_OFF | ENSSSCG00000000104 | 5 | 9494087 | 9494136 | 181.98 | 2.20 | 174.74 | 2.17 | 210.09 | 2.75 | 202.55 | 3.17 | 198.21 | 2.93 | 155.27 | 1.17 |
| SKIP_ON | ENSSSCG00000000104 | 5 | 9499278 | 9499513 | 181.98 | 37.49 | 174.74 | 24.64 | 210.09 | 70.46 | 202.55 | 45.57 | 198.21 | 61.78 | 155.27 | 43.26 |
| SKIP_OFF | ENSSSCG00000000104 | 5 | 9499278 | 9499513 | 181.98 | 144.48 | 174.74 | 150.10 | 210.09 | 139.62 | 202.55 | 156.98 | 198.21 | 136.43 | 155.27 | 112.01 |
| IR_ON | ENSSSCG00000000104 | 5 | 9494087 | 9494136 | 181.98 | 179.77 | 174.74 | 172.57 | 210.09 | 207.34 | 202.55 | 199.38 | 198.21 | 195.29 | 155.27 | 154.10 |
| TSS | ENSSSCG00000000104 | 5 | 9485487 | 9486078 | 181.98 | 181.98 | 174.74 | 174.74 | 210.09 | 210.09 | 202.55 | 202.55 | 198.21 | 198.21 | 155.27 | 155.27 |
| TTS | ENSSSCG00000000104 | 5 | 9502133 | 9505152 | 181.98 | 181.98 | 174.74 | 174.74 | 210.09 | 210.09 | 202.55 | 202.55 | 198.21 | 198.21 | 155.27 | 155.27 |
| TSS | ENSSSCG00000000105 | 5 | 9515461 | 9515734 | 31.56 | 31.56 | 24.56 | 24.56 | 31.97 | 31.97 | 11.38 | 11.38 | 21.96 | 21.96 | 31.64 | 31.64 |
| TTS | ENSSSCG00000000105 | 5 | 9505157 | 9506113 | 31.56 | 31.56 | 24.56 | 24.56 | 31.97 | 31.97 | 11.38 | 11.38 | 21.96 | 21.96 | 31.64 | 31.64 |
| TSS | ENSSSCG00000000107 | 5 | 9581997 | 9582104 | 22.62 | 1.62 | 26.46 | 2.27 | 15.77 | 0.47 | 14.98 | 0.58 | 17.09 | 1.63 | 14.09 | 0.72 |
| SKIP_ON | ENSSSCG00000000107 | 5 | 9616485 | 9616545 | 22.62 | 1.63 | 26.46 | 2.27 | 15.77 | 0.47 | 14.98 | 0.58 | 17.09 | 1.63 | 14.09 | 0.72 |
| TSS | ENSSSCG00000000107 | 5 | 9594464 | 9595355 | 22.62 | 1.79 | 26.46 | 2.17 | 15.77 | 1.84 | 14.98 | 2.68 | 17.09 | 1.47 | 14.09 | 1.94 |

| | | | | | | | | | | | | | | | | |
|---|---|---|---|---|---|---|---|---|---|---|---|---|---|---|---|---|
| SKIP_OFF | ENSSSCG00000000107 | 5 | 9607844 | 9607954 | 22.62 | 5.80 | 26.46 | 6.44 | 15.77 | 4.16 | 14.98 | 5.13 | 17.09 | 5.09 | 14.09 | 4.75 |
| IR_ON | ENSSSCG00000000107 | 5 | 9613265 | 9613846 | 22.62 | 5.99 | 26.46 | 7.52 | 15.77 | 3.86 | 14.98 | 2.15 | 17.09 | 2.91 | 14.09 | 2.04 |
| IR_OFF | ENSSSCG00000000107 | 5 | 9613265 | 9613846 | 22.62 | 7.42 | 26.46 | 8.05 | 15.77 | 5.43 | 14.98 | 4.43 | 17.09 | 5.99 | 14.09 | 4.64 |
| TSS | ENSSSCG00000000107 | 5 | 9594987 | 9595143 | 22.62 | 7.42 | 26.46 | 8.05 | 15.77 | 5.43 | 14.98 | 4.43 | 17.09 | 5.99 | 14.09 | 4.64 |
| SKIP_OFF | ENSSSCG00000000107 | 5 | 9616485 | 9616545 | 22.62 | 7.58 | 26.46 | 8.62 | 15.77 | 6.00 | 14.98 | 7.81 | 17.09 | 6.56 | 14.09 | 6.69 |
| TTS | ENSSSCG00000000107 | 5 | 9619246 | 9623906 | 22.62 | 9.21 | 26.46 | 10.89 | 15.77 | 6.47 | 14.98 | 8.40 | 17.09 | 8.19 | 14.09 | 7.41 |
| TTS | ENSSSCG00000000107 | 5 | 9614047 | 9615759 | 22.62 | 13.41 | 26.46 | 15.58 | 15.77 | 9.29 | 14.98 | 6.58 | 17.09 | 8.90 | 14.09 | 6.68 |
| SKIP_ON | ENSSSCG00000000107 | 5 | 9607844 | 9607954 | 22.62 | 16.82 | 26.46 | 20.02 | 15.77 | 11.61 | 14.98 | 9.85 | 17.09 | 11.99 | 14.09 | 9.34 |
| SKIP_OFF | ENSSSCG00000000108 | 5 | 9678805 | 9678825 | 12.32 | 3.53 | 14.96 | 4.15 | 12.50 | 3.45 | 9.87 | 3.05 | 9.41 | 2.60 | 11.73 | 4.16 |
| TSS | ENSSSCG00000000108 | 5 | 9632919 | 9633382 | 12.32 | 3.53 | 14.96 | 4.15 | 12.50 | 3.45 | 9.87 | 3.05 | 9.41 | 2.60 | 11.73 | 4.16 |
| SKIP_ON | ENSSSCG00000000108 | 5 | 9678805 | 9678825 | 12.32 | 5.17 | 14.96 | 7.13 | 12.50 | 6.02 | 9.87 | 4.58 | 9.41 | 3.45 | 11.73 | 5.08 |
| TSS | ENSSSCG00000000108 | 5 | 9633555 | 9633704 | 12.32 | 8.78 | 14.96 | 10.81 | 12.50 | 9.05 | 9.87 | 6.82 | 9.41 | 6.81 | 11.73 | 7.57 |
| TTS | ENSSSCG00000000108 | 5 | 9679734 | 9682473 | 12.32 | 8.70 | 14.96 | 11.28 | 12.50 | 9.46 | 9.87 | 7.63 | 9.41 | 6.05 | 11.73 | 9.24 |
| SKIP_ON | ENSSSCG00000000110 | 5 | 9754176 | 9754337 | 2.82 | 0.22 | 3.81 | 0.52 | 4.89 | 0.53 | 4.11 | 0.92 | 5.32 | 0.94 | 4.51 | 1.21 |
| SKIP_ON | ENSSSCG00000000110 | 5 | 9723857 | 9723891 | 2.82 | 0.28 | 3.81 | 0.35 | 4.89 | 0.90 | 4.11 | 0.63 | 5.32 | 0.00 | 4.51 | 0.97 |
| XIR_OFF | ENSSSCG00000000110 | 5 | 9723891 | 9723991 | 2.82 | 0.28 | 3.81 | 0.35 | 4.89 | 0.90 | 4.11 | 0.63 | 5.32 | 0.00 | 4.51 | 0.97 |
| TSS | ENSSSCG00000000110 | 5 | 9713968 | 9714050 | 2.82 | 1.18 | 3.81 | 0.75 | 4.89 | 1.45 | 4.11 | 1.33 | 5.32 | 0.24 | 4.51 | 0.40 |
| XIR_ON | ENSSSCG00000000110 | 5 | 9723891 | 9723991 | 2.82 | 1.18 | 3.81 | 0.75 | 4.89 | 1.45 | 4.11 | 1.33 | 5.32 | 0.24 | 4.51 | 0.40 |
| SKIP_OFF | ENSSSCG00000000110 | 5 | 9723857 | 9723891 | 2.82 | 1.14 | 3.81 | 2.20 | 4.89 | 2.01 | 4.11 | 1.23 | 5.32 | 4.14 | 4.51 | 1.94 |
| TSS | ENSSSCG00000000110 | 5 | 9713765 | 9713927 | 2.82 | 1.42 | 3.81 | 2.55 | 4.89 | 2.91 | 4.11 | 1.86 | 5.32 | 4.14 | 4.51 | 2.91 |
| SKIP_OFF | ENSSSCG00000000110 | 5 | 9754176 | 9754337 | 2.82 | 2.60 | 3.81 | 3.29 | 4.89 | 4.36 | 4.11 | 3.19 | 5.32 | 4.38 | 4.51 | 3.30 |
| TTS | ENSSSCG00000000110 | 5 | 9777116 | 9778191 | 2.82 | 2.82 | 3.81 | 3.81 | 4.89 | 4.89 | 4.11 | 4.11 | 5.32 | 5.32 | 4.51 | 4.51 |
| TSS | ENSSSCG00000000111 | 5 | 9778727 | 9778877 | 0.10 | 0.10 | 0.26 | 0.26 | 0.07 | 0.07 | 0.38 | 0.38 | 0.13 | 0.13 | 0.33 | 0.33 |
| TTS | ENSSSCG00000000111 | 5 | 9799215 | 9799703 | 0.10 | 0.10 | 0.26 | 0.26 | 0.07 | 0.07 | 0.38 | 0.38 | 0.13 | 0.13 | 0.33 | 0.33 |
| SKIP_OFF | ENSSSCG00000000114 | 5 | 9810675 | 9810725 | 4.39 | 0.00 | 4.00 | 0.00 | 5.32 | 0.00 | 9.51 | 0.00 | 7.35 | 0.00 | 9.64 | 0.00 |
| TSS | ENSSSCG00000000114 | 5 | 9825363 | 9826322 | 4.39 | 1.90 | 4.00 | 1.44 | 5.32 | 1.85 | 9.51 | 3.33 | 7.35 | 2.13 | 9.64 | 3.06 |
| TSS | ENSSSCG00000000114 | 5 | 9889814 | 9889930 | 4.39 | 2.47 | 4.00 | 2.52 | 5.32 | 3.46 | 9.51 | 6.11 | 7.35 | 5.11 | 9.64 | 6.47 |
| SKIP_ON | ENSSSCG00000000114 | 5 | 9810675 | 9810725 | 4.39 | 4.39 | 4.00 | 4.00 | 5.32 | 5.32 | 9.51 | 9.51 | 7.35 | 7.35 | 9.64 | 9.64 |
| TTS | ENSSSCG00000000114 | 5 | 9804946 | 9809287 | 4.39 | 4.39 | 4.00 | 4.00 | 5.32 | 5.32 | 9.51 | 9.51 | 7.35 | 7.35 | 9.64 | 9.64 |
| TSS | ENSSSCG00000000116 | 5 | 9917590 | 9917744 | 35.56 | 35.56 | 30.33 | 30.33 | 29.36 | 29.36 | 38.25 | 38.25 | 29.83 | 29.83 | 32.14 | 32.14 |
| TTS | ENSSSCG00000000116 | 5 | 9904948 | 9905540 | 35.56 | 35.56 | 30.33 | 30.33 | 29.36 | 29.36 | 38.25 | 38.25 | 29.83 | 29.83 | 32.14 | 32.14 |
| TSS | ENSSSCG00000000117 | 5 | 9917751 | 9918044 | 2.94 | 2.94 | 1.51 | 1.51 | 1.83 | 1.83 | 21.52 | 21.52 | 17.22 | 17.22 | 23.73 | 23.73 |
| TTS | ENSSSCG00000000117 | 5 | 9924014 | 9925634 | 2.94 | 2.94 | 1.51 | 1.51 | 1.83 | 1.83 | 21.52 | 21.52 | 17.22 | 17.22 | 23.73 | 23.73 |
| IR_OFF | ENSSSCG00000000118 | 5 | 9939885 | 9939901 | 5.83 | 0.49 | 4.62 | 0.28 | 5.61 | 0.00 | 10.69 | 0.63 | 6.68 | 0.79 | 8.51 | 0.33 |
| IR_ON | ENSSSCG00000000118 | 5 | 9939885 | 9939901 | 5.83 | 5.34 | 4.62 | 4.34 | 5.61 | 5.61 | 10.69 | 10.06 | 6.68 | 5.89 | 8.51 | 8.19 |
| TSS | ENSSSCG00000000118 | 5 | 9956682 | 9956827 | 5.83 | 5.83 | 4.62 | 4.62 | 5.61 | 5.61 | 10.69 | 10.69 | 6.68 | 6.68 | 8.51 | 8.51 |
| TTS | ENSSSCG00000000118 | 5 | 9927320 | 9927441 | 5.83 | 5.83 | 4.62 | 4.62 | 5.61 | 5.61 | 10.69 | 10.69 | 6.68 | 6.68 | 8.51 | 8.51 |
| TSS | ENSSSCG00000000119 | 5 | 9993804 | 9994259 | 105.52 | 3.27 | 106.91 | 3.29 | 106.33 | 3.64 | 90.88 | 3.73 | 107.95 | 4.90 | 119.31 | 3.32 |
| XIR_OFF | ENSSSCG00000000119 | 5 | 9993752 | 9993804 | 105.52 | 3.27 | 106.91 | 3.29 | 106.33 | 3.64 | 90.88 | 3.73 | 107.95 | 4.90 | 119.31 | 3.32 |
| IR_ON | ENSSSCG00000000119 | 5 | 9976896 | 9976993 | 105.52 | 9.39 | 106.91 | 9.35 | 106.33 | 10.08 | 90.88 | 10.61 | 107.95 | 9.81 | 119.31 | 10.03 |
| IR_OFF | ENSSSCG00000000119 | 5 | 9976896 | 9976993 | 105.52 | 96.13 | 106.91 | 97.56 | 106.33 | 96.24 | 90.88 | 80.27 | 107.95 | 98.14 | 119.31 | 109.28 |
| TSS | ENSSSCG00000000119 | 5 | 9995961 | 9996260 | 105.52 | 102.25 | 106.91 | 103.62 | 106.33 | 102.69 | 90.88 | 87.16 | 107.95 | 103.05 | 119.31 | 115.99 |
| XIR_ON | ENSSSCG00000000119 | 5 | 9993752 | 9993804 | 105.52 | 102.25 | 106.91 | 103.62 | 106.33 | 102.69 | 90.88 | 87.16 | 107.95 | 103.05 | 119.31 | 115.99 |
| TTS | ENSSSCG00000000119 | 5 | 9966122 | 9970228 | 105.52 | 105.52 | 106.91 | 106.91 | 106.33 | 106.33 | 90.88 | 90.88 | 107.95 | 107.95 | 119.31 | 119.31 |
| TSS | ENSSSCG00000000120 | 5 | 10003836 | 10004343 | 4.49 | 4.49 | 3.28 | 3.28 | 4.26 | 4.26 | 6.13 | 6.13 | 5.01 | 5.01 | 6.59 | 6.59 |
| TTS | ENSSSCG00000000120 | 5 | 10016647 | 10018944 | 4.49 | 4.49 | 3.28 | 3.28 | 4.26 | 4.26 | 6.13 | 6.13 | 5.01 | 5.01 | 6.59 | 6.59 |
| TSS | ENSSSCG00000000121 | 5 | 10025743 | 10026101 | 0.13 | 0.00 | 0.19 | 0.00 | 0.41 | 0.00 | 0.53 | 0.00 | 0.37 | 0.00 | 0.43 | 0.00 |
| TTS | ENSSSCG00000000121 | 5 | 10024348 | 10025119 | 0.13 | 0.00 | 0.19 | 0.00 | 0.41 | 0.00 | 0.53 | 0.00 | 0.37 | 0.00 | 0.43 | 0.00 |
| TSS | ENSSSCG00000000130 | 5 | 10521565 | 10521735 | 2.03 | 2.03 | 4.18 | 4.18 | 3.32 | 3.32 | 4.22 | 4.22 | 4.37 | 4.37 | 2.78 | 2.78 |
| TTS | ENSSSCG00000000130 | 5 | 10489289 | 10491274 | 2.03 | 2.03 | 4.18 | 4.18 | 3.32 | 3.32 | 4.22 | 4.22 | 4.37 | 4.37 | 2.78 | 2.78 |

| | | | | | | | | | | | | | | | | |
|---|---|---|---|---|---|---|---|---|---|---|---|---|---|---|---|---|
| TSS | ENSSSCG00000000133 | 5 | 10750018 | 10750113 | 8.52 | 0.17 | 5.68 | 0.25 | 9.80 | 0.25 | 17.02 | 0.39 | 7.75 | 0.31 | 13.43 | 0.30 |
| TSS | ENSSSCG00000000133 | 5 | 10746890 | 10747021 | 8.52 | 1.04 | 5.68 | 1.06 | 9.80 | 2.82 | 17.02 | 4.48 | 7.75 | 0.00 | 13.43 | 1.47 |
| TSS | ENSSSCG00000000133 | 5 | 10749467 | 10749613 | 8.52 | 7.31 | 5.68 | 4.37 | 9.80 | 6.73 | 17.02 | 12.16 | 7.75 | 7.44 | 13.43 | 11.66 |
| TTS | ENSSSCG00000000133 | 5 | 10759154 | 10759698 | 8.52 | 8.52 | 5.68 | 5.68 | 9.80 | 9.80 | 17.02 | 17.02 | 7.75 | 7.75 | 13.43 | 13.43 |
| TSS | ENSSSCG00000000135 | 5 | 10718741 | 10718983 | 4.65 | 4.65 | 2.36 | 2.36 | 4.29 | 4.29 | 3.56 | 3.56 | 4.52 | 4.52 | 6.17 | 6.17 |
| TTS | ENSSSCG00000000135 | 5 | 10707354 | 10708212 | 4.65 | 4.65 | 2.36 | 2.36 | 4.29 | 4.29 | 3.56 | 3.56 | 4.52 | 4.52 | 6.17 | 6.17 |
| TSS | ENSSSCG00000000136 | 5 | 10840309 | 10840808 | 1.34 | 1.34 | 4.03 | 4.03 | 2.74 | 2.74 | 1.50 | 1.50 | 2.37 | 2.37 | 1.15 | 1.15 |
| TTS | ENSSSCG00000000136 | 5 | 10814008 | 10819547 | 1.34 | 1.34 | 4.03 | 4.03 | 2.74 | 2.74 | 1.50 | 1.50 | 2.37 | 2.37 | 1.15 | 1.15 |
| TSS | ENSSSCG00000000137 | 5 | 10992748 | 10992803 | 3.06 | 0.00 | 7.76 | 0.00 | 6.73 | 0.36 | 4.04 | 0.14 | 7.50 | 0.48 | 4.06 | 0.51 |
| TSS | ENSSSCG00000000137 | 5 | 10897750 | 10898001 | 3.06 | 3.06 | 7.76 | 7.76 | 6.73 | 6.37 | 4.04 | 3.90 | 7.50 | 7.02 | 4.06 | 3.55 |
| TTS | ENSSSCG00000000137 | 5 | 10878435 | 10878818 | 3.06 | 3.06 | 7.76 | 7.76 | 6.73 | 6.73 | 4.04 | 4.04 | 7.50 | 7.50 | 4.06 | 4.06 |
| TSS | ENSSSCG00000000138 | 5 | 10954978 | 10955691 | 1.03 | 0.00 | 0.00 | 0.00 | 0.35 | 0.04 | 0.95 | 0.06 | 0.19 | 0.06 | 0.74 | 0.06 |
| TSS | ENSSSCG00000000138 | 5 | 10953580 | 10953664 | 1.03 | 1.03 | 0.00 | 0.00 | 0.35 | 0.31 | 0.95 | 0.89 | 0.19 | 0.13 | 0.74 | 0.68 |
| TTS | ENSSSCG00000000138 | 5 | 10973077 | 10974006 | 1.03 | 1.03 | 0.00 | 0.00 | 0.35 | 0.35 | 0.95 | 0.95 | 0.19 | 0.19 | 0.74 | 0.74 |
| TSS | ENSSSCG00000000139 | 5 | 11000412 | 11000460 | 7.97 | 7.97 | 5.82 | 5.82 | 6.81 | 6.81 | 23.62 | 23.62 | 13.00 | 13.00 | 20.24 | 20.24 |
| TTS | ENSSSCG00000000139 | 5 | 11015229 | 11015327 | 7.97 | 7.97 | 5.82 | 5.82 | 6.81 | 6.81 | 23.62 | 23.62 | 13.00 | 13.00 | 20.24 | 20.24 |
| TSS | ENSSSCG00000000141 | 5 | 11213281 | 11213786 | 21.62 | 21.62 | 17.88 | 17.88 | 22.79 | 22.79 | 20.31 | 20.31 | 18.59 | 18.59 | 27.23 | 27.23 |
| TTS | ENSSSCG00000000141 | 5 | 11229011 | 11233149 | 21.62 | 21.62 | 17.88 | 17.88 | 22.79 | 22.79 | 20.31 | 20.31 | 18.59 | 18.59 | 27.23 | 27.23 |
| TSS | ENSSSCG00000000142 | 5 | 11233845 | 11235100 | 3.70 | 1.49 | 11.52 | 5.53 | 9.92 | 3.87 | 2.58 | 1.12 | 3.77 | 1.65 | 2.83 | 1.66 |
| TSS | ENSSSCG00000000142 | 5 | 11233921 | 11233932 | 3.70 | 2.21 | 11.52 | 5.99 | 9.92 | 6.05 | 2.58 | 1.46 | 3.77 | 2.13 | 2.83 | 1.18 |
| TTS | ENSSSCG00000000142 | 5 | 11250815 | 11254712 | 3.70 | 3.70 | 11.52 | 11.52 | 9.92 | 9.92 | 2.58 | 2.58 | 3.77 | 3.77 | 2.83 | 2.83 |
| TSS | ENSSSCG00000000144 | 5 | 11264602 | 11265705 | 37.77 | 3.23 | 46.40 | 6.24 | 49.24 | 4.76 | 41.74 | 2.62 | 35.35 | 2.76 | 44.76 | 2.82 |
| TSS | ENSSSCG00000000144 | 5 | 11264465 | 11264658 | 37.77 | 34.54 | 46.40 | 40.16 | 49.24 | 44.48 | 41.74 | 39.12 | 35.35 | 32.59 | 44.76 | 41.94 |
| TTS | ENSSSCG00000000144 | 5 | 11276793 | 11277631 | 37.77 | 37.77 | 46.40 | 46.40 | 49.24 | 49.24 | 41.74 | 41.74 | 35.35 | 35.35 | 44.76 | 44.76 |
| TTS | ENSSSCG00000000145 | 5 | 11417917 | 11417931 | 53.57 | 0.00 | 74.64 | 0.00 | 52.73 | 0.00 | 42.41 | 0.44 | 47.35 | 0.83 | 50.75 | 0.00 |
| TTS | ENSSSCG00000000145 | 5 | 11454937 | 11455054 | 53.57 | 53.57 | 74.64 | 74.64 | 52.73 | 52.73 | 42.41 | 41.97 | 47.35 | 46.51 | 50.75 | 50.75 |
| TSS | ENSSSCG00000000145 | 5 | 11398166 | 11398498 | 53.57 | 53.57 | 74.64 | 74.64 | 52.73 | 52.73 | 42.41 | 42.41 | 47.35 | 47.35 | 50.75 | 50.75 |
| TSS | ENSSSCG00000000146 | 3 | 7019423 | 7019484 | 35.97 | 0.00 | 39.36 | 0.00 | 41.14 | 0.00 | 419.62 | 0.33 | 30.73 | 0.00 | 18.77 | 0.00 |
| IR_OFF | ENSSSCG00000000146 | 3 | 7113077 | 7113213 | 35.97 | 0.57 | 39.36 | 2.35 | 41.14 | 0.92 | 419.62 | 15.96 | 30.73 | 1.68 | 18.77 | 1.17 |
| SKIP_OFF | ENSSSCG00000000146 | 3 | 7138810 | 7138904 | 35.97 | 1.62 | 39.36 | 1.61 | 41.14 | 2.11 | 419.62 | 97.53 | 30.73 | 1.99 | 18.77 | 1.39 |
| SKIP_ON | ENSSSCG00000000146 | 3 | 7139020 | 7139058 | 35.97 | 1.62 | 39.36 | 1.61 | 41.14 | 2.11 | 419.62 | 97.53 | 30.73 | 1.99 | 18.77 | 1.39 |
| MSKIP_ON | ENSSSCG00000000146 | 3 | 7138810 | 7139058 | 35.97 | 6.48 | 39.36 | 11.33 | 41.14 | 10.35 | 419.62 | 15.68 | 30.73 | 7.17 | 18.77 | 2.90 |
| SKIP_ON | ENSSSCG00000000146 | 3 | 7138810 | 7138904 | 35.97 | 6.48 | 39.36 | 11.33 | 41.14 | 10.35 | 419.62 | 15.68 | 30.73 | 7.17 | 18.77 | 2.90 |
| SKIP_OFF | ENSSSCG00000000146 | 3 | 7075419 | 7075439 | 35.97 | 7.35 | 39.36 | 14.14 | 41.14 | 11.56 | 419.62 | 32.51 | 30.73 | 10.36 | 18.77 | 5.41 |
| MSKIP_OFF | ENSSSCG00000000146 | 3 | 7138810 | 7139058 | 35.97 | 26.99 | 39.36 | 23.61 | 41.14 | 27.47 | 419.62 | 289.58 | 30.73 | 18.38 | 18.77 | 11.97 |
| SKIP_OFF | ENSSSCG00000000146 | 3 | 7139020 | 7139058 | 35.97 | 26.99 | 39.36 | 23.61 | 41.14 | 27.47 | 419.62 | 289.58 | 30.73 | 18.38 | 18.77 | 11.97 |
| SKIP_ON | ENSSSCG00000000146 | 3 | 7075419 | 7075439 | 35.97 | 28.61 | 39.36 | 25.22 | 41.14 | 29.58 | 419.62 | 387.10 | 30.73 | 20.38 | 18.77 | 13.36 |
| TTS | ENSSSCG00000000146 | 3 | 7142366 | 7143088 | 35.97 | 35.10 | 39.36 | 36.55 | 41.14 | 39.93 | 419.62 | 402.78 | 30.73 | 27.54 | 18.77 | 16.26 |
| IR_ON | ENSSSCG00000000146 | 3 | 7113077 | 7113213 | 35.97 | 35.39 | 39.36 | 37.01 | 41.14 | 40.22 | 419.62 | 403.65 | 30.73 | 29.05 | 18.77 | 17.60 |
| TSS | ENSSSCG00000000146 | 3 | 7002706 | 7002821 | 35.97 | 35.97 | 39.36 | 39.36 | 41.14 | 41.14 | 419.62 | 419.28 | 30.73 | 30.73 | 18.77 | 18.77 |
| TSS | ENSSSCG00000000148 | 5 | 11542534 | 11542607 | 0.94 | 0.09 | 1.48 | 0.10 | 0.64 | 0.24 | 2.57 | 0.53 | 1.50 | 0.29 | 1.57 | 0.39 |
| XIR_ON | ENSSSCG00000000148 | 5 | 11516454 | 11516636 | 0.94 | 0.09 | 1.48 | 0.10 | 0.64 | 0.24 | 2.57 | 0.53 | 1.50 | 0.29 | 1.57 | 0.39 |
| AE | ENSSSCG00000000148 | 5 | 11541362 | 11542628 | 0.94 | 0.35 | 1.48 | 0.70 | 0.64 | 0.05 | 2.57 | 0.17 | 1.50 | 0.31 | 1.57 | 0.08 |
| SKIP_ON | ENSSSCG00000000148 | 5 | 11516636 | 11516677 | 0.94 | 0.35 | 1.48 | 0.70 | 0.64 | 0.05 | 2.57 | 0.17 | 1.50 | 0.31 | 1.57 | 0.08 |
| XIR_OFF | ENSSSCG00000000148 | 5 | 11516454 | 11516636 | 0.94 | 0.35 | 1.48 | 0.70 | 0.64 | 0.05 | 2.57 | 0.17 | 1.50 | 0.31 | 1.57 | 0.08 |
| AE | ENSSSCG00000000148 | 5 | 11541404 | 11542628 | 0.94 | 0.50 | 1.48 | 0.67 | 0.64 | 0.35 | 2.57 | 1.86 | 1.50 | 0.90 | 1.57 | 1.10 |
| SKIP_OFF | ENSSSCG00000000148 | 5 | 11516636 | 11516677 | 0.94 | 0.50 | 1.48 | 0.67 | 0.64 | 0.35 | 2.57 | 1.86 | 1.50 | 0.90 | 1.57 | 1.10 |
| TSS | ENSSSCG00000000148 | 5 | 11542768 | 11543023 | 0.94 | 0.85 | 1.48 | 1.37 | 0.64 | 0.40 | 2.57 | 2.03 | 1.50 | 1.21 | 1.57 | 1.18 |
| TTS | ENSSSCG00000000148 | 5 | 11509336 | 11511176 | 0.94 | 0.94 | 1.48 | 1.48 | 0.64 | 0.64 | 2.57 | 2.57 | 1.50 | 1.50 | 1.57 | 1.57 |
| TSS | ENSSSCG00000000152 | 5 | 11799218 | 11799676 | 20.19 | 0.00 | 21.61 | 0.00 | 29.15 | 0.19 | 22.90 | 0.25 | 19.59 | 0.00 | 20.68 | 0.00 |

| | | | | | | | | | | | | | | | | |
|---|---|---|---|---|---|---|---|---|---|---|---|---|---|---|---|---|
| MSKIP_OFF | ENSSSCG00000000152 | 5 | 11887743 | 11897527 | 20.19 | 0.12 | 21.61 | 0.00 | 29.15 | 0.13 | 22.90 | 0.00 | 19.59 | 0.00 | 20.68 | 0.06 |
| SKIP_OFF | ENSSSCG00000000152 | 5 | 11897439 | 11897527 | 20.19 | 0.12 | 21.61 | 0.00 | 29.15 | 0.13 | 22.90 | 0.00 | 19.59 | 0.00 | 20.68 | 0.06 |
| SKIP_ON | ENSSSCG00000000152 | 5 | 11887743 | 11887782 | 20.19 | 0.00 | 21.61 | 0.00 | 29.15 | 0.77 | 22.90 | 0.00 | 19.59 | 0.00 | 20.68 | 0.00 |
| TSS | ENSSSCG00000000152 | 5 | 11752664 | 11752875 | 20.19 | 0.14 | 21.61 | 0.28 | 29.15 | 0.81 | 22.90 | 4.18 | 19.59 | 0.15 | 20.68 | 1.03 |
| MSKIP_ON | ENSSSCG00000000152 | 5 | 11890802 | 11897527 | 20.19 | 0.14 | 21.61 | 0.28 | 29.15 | 0.99 | 22.90 | 4.42 | 19.59 | 0.15 | 20.68 | 1.03 |
| SKIP_OFF | ENSSSCG00000000152 | 5 | 11887743 | 11887782 | 20.19 | 0.14 | 21.61 | 0.28 | 29.15 | 0.99 | 22.90 | 4.42 | 19.59 | 0.15 | 20.68 | 1.03 |
| SKIP_ON | ENSSSCG00000000152 | 5 | 11890802 | 11890844 | 20.19 | 0.14 | 21.61 | 0.28 | 29.15 | 0.99 | 22.90 | 4.42 | 19.59 | 0.15 | 20.68 | 1.03 |
| SKIP_OFF | ENSSSCG00000000152 | 5 | 11875949 | 11876041 | 20.19 | 0.89 | 21.61 | 0.97 | 29.15 | 0.60 | 22.90 | 0.22 | 19.59 | 0.92 | 20.68 | 0.83 |
| MSKIP_ON | ENSSSCG00000000152 | 5 | 11887743 | 11897527 | 20.19 | 1.49 | 21.61 | 0.24 | 29.15 | 1.63 | 22.90 | 1.31 | 19.59 | 0.58 | 20.68 | 1.03 |
| MSKIP_ON | ENSSSCG00000000152 | 5 | 11887743 | 11893041 | 20.19 | 1.49 | 21.61 | 0.24 | 29.15 | 1.63 | 22.90 | 1.31 | 19.59 | 0.58 | 20.68 | 1.03 |
| SKIP_ON | ENSSSCG00000000152 | 5 | 11893010 | 11893041 | 20.19 | 1.49 | 21.61 | 0.24 | 29.15 | 1.63 | 22.90 | 1.31 | 19.59 | 0.58 | 20.68 | 1.03 |
| MSKIP_OFF | ENSSSCG00000000152 | 5 | 11887743 | 11893041 | 20.19 | 0.42 | 21.61 | 1.95 | 29.15 | 2.35 | 22.90 | 0.46 | 19.59 | 1.05 | 20.68 | 0.80 |
| SKIP_OFF | ENSSSCG00000000152 | 5 | 11887743 | 11887782 | 20.19 | 0.42 | 21.61 | 1.95 | 29.15 | 2.35 | 22.90 | 0.46 | 19.59 | 1.05 | 20.68 | 0.80 |
| SKIP_ON | ENSSSCG00000000152 | 5 | 11897439 | 11897527 | 20.19 | 0.42 | 21.61 | 1.95 | 29.15 | 2.35 | 22.90 | 0.46 | 19.59 | 1.05 | 20.68 | 0.80 |
| AE | ENSSSCG00000000152 | 5 | 11883894 | 11884038 | 20.19 | 2.38 | 21.61 | 1.21 | 29.15 | 2.41 | 22.90 | 1.78 | 19.59 | 1.50 | 20.68 | 1.86 |
| TSS | ENSSSCG00000000152 | 5 | 11633546 | 11633824 | 20.19 | 1.76 | 21.61 | 2.45 | 29.15 | 3.13 | 22.90 | 2.19 | 19.59 | 1.96 | 20.68 | 1.63 |
| TSS | ENSSSCG00000000152 | 5 | 11696934 | 11697001 | 20.19 | 4.72 | 21.61 | 6.53 | 29.15 | 6.68 | 22.90 | 3.77 | 19.59 | 4.43 | 20.68 | 6.70 |
| TSS | ENSSSCG00000000152 | 5 | 11808730 | 11809415 | 20.19 | 13.56 | 21.61 | 12.36 | 29.15 | 18.34 | 22.90 | 12.52 | 19.59 | 13.04 | 20.68 | 11.32 |
| MSKIP_ON | ENSSSCG00000000152 | 5 | 11887743 | 11897527 | 20.19 | 16.26 | 21.61 | 16.70 | 29.15 | 20.14 | 22.90 | 14.51 | 19.59 | 15.85 | 20.68 | 16.13 |
| SKIP_OFF | ENSSSCG00000000152 | 5 | 11893010 | 11893041 | 20.19 | 16.26 | 21.61 | 16.70 | 29.15 | 20.14 | 22.90 | 14.51 | 19.59 | 15.85 | 20.68 | 16.13 |
| SKIP_ON | ENSSSCG00000000152 | 5 | 11887743 | 11887782 | 20.19 | 16.26 | 21.61 | 16.70 | 29.15 | 20.14 | 22.90 | 14.51 | 19.59 | 15.85 | 20.68 | 16.13 |
| AE | ENSSSCG00000000152 | 5 | 11883906 | 11884038 | 20.19 | 16.05 | 21.61 | 17.96 | 29.15 | 23.60 | 22.90 | 18.92 | 19.59 | 16.12 | 20.68 | 17.19 |
| TTS | ENSSSCG00000000152 | 5 | 11899706 | 11899864 | 20.19 | 18.42 | 21.61 | 19.17 | 29.15 | 25.24 | 22.90 | 20.71 | 19.59 | 17.62 | 20.68 | 19.05 |
| SKIP_ON | ENSSSCG00000000152 | 5 | 11875949 | 11876041 | 20.19 | 19.30 | 21.61 | 20.64 | 29.15 | 28.54 | 22.90 | 22.67 | 19.59 | 18.66 | 20.68 | 19.85 |
| TSS | ENSSSCG00000000156 | 5 | 12525806 | 12525927 | 18.56 | 1.26 | 10.51 | 0.78 | 14.99 | 0.92 | 19.29 | 1.12 | 19.20 | 1.07 | 18.92 | 0.96 |
| TSS | ENSSSCG00000000156 | 5 | 12525569 | 12525727 | 18.56 | 17.31 | 10.51 | 9.72 | 14.99 | 14.07 | 19.29 | 18.17 | 19.20 | 18.12 | 18.92 | 17.96 |
| TTS | ENSSSCG00000000156 | 5 | 12501275 | 12506332 | 18.56 | 18.56 | 10.51 | 10.51 | 14.99 | 14.99 | 19.29 | 19.29 | 19.20 | 19.20 | 18.92 | 18.92 |
| TTS | ENSSSCG00000000157 | 5 | 12578357 | 12579900 | 0.16 | 0.09 | 0.10 | 0.09 | 0.27 | 0.16 | 0.89 | 0.61 | 0.70 | 0.36 | 1.22 | 0.78 |
| TSS | ENSSSCG00000000157 | 5 | 12535679 | 12535770 | 0.16 | 0.16 | 0.10 | 0.10 | 0.27 | 0.27 | 0.89 | 0.89 | 0.70 | 0.70 | 1.22 | 1.22 |
| TSS | ENSSSCG00000000158 | 5 | 12579975 | 12580462 | 15.95 | 15.95 | 11.53 | 11.53 | 14.21 | 14.21 | 13.22 | 13.22 | 11.95 | 11.95 | 16.86 | 16.86 |
| TTS | ENSSSCG00000000158 | 5 | 12605489 | 12607578 | 15.95 | 15.95 | 11.53 | 11.53 | 14.21 | 14.21 | 13.22 | 13.22 | 11.95 | 11.95 | 16.86 | 16.86 |
| SKIP_OFF | ENSSSCG00000000160 | 5 | 12654723 | 12654872 | 6.06 | 0.13 | 4.16 | 0.00 | 5.61 | 0.00 | 5.47 | 0.00 | 5.07 | 0.00 | 4.90 | 0.12 |
| TSS | ENSSSCG00000000160 | 5 | 12638291 | 12639128 | 6.06 | 0.13 | 4.16 | 0.00 | 5.61 | 0.00 | 5.47 | 0.00 | 5.07 | 0.00 | 4.90 | 0.12 |
| TSS | ENSSSCG00000000160 | 5 | 12638389 | 12638508 | 6.06 | 2.91 | 4.16 | 2.26 | 5.61 | 2.32 | 5.47 | 2.68 | 5.07 | 3.58 | 4.90 | 1.62 |
| TSS | ENSSSCG00000000160 | 5 | 12638514 | 12638667 | 6.06 | 3.02 | 4.16 | 1.90 | 5.61 | 3.29 | 5.47 | 2.80 | 5.07 | 1.50 | 4.90 | 3.17 |
| SKIP_ON | ENSSSCG00000000160 | 5 | 12654723 | 12654872 | 6.06 | 5.93 | 4.16 | 4.16 | 5.61 | 5.61 | 5.47 | 5.47 | 5.07 | 5.07 | 4.90 | 4.79 |
| TTS | ENSSSCG00000000160 | 5 | 12667225 | 12667537 | 6.06 | 6.06 | 4.16 | 4.16 | 5.61 | 5.61 | 5.47 | 5.47 | 5.07 | 5.07 | 4.90 | 4.90 |
| XSKIP_OFF | ENSSSCG00000000161 | 5 | 12709306 | 12709367 | 19.17 | 0.00 | 14.04 | 0.00 | 15.01 | 0.00 | 14.77 | 0.00 | 13.50 | 0.00 | 13.95 | 0.00 |
| XSKIP_ON | ENSSSCG00000000161 | 5 | 12709306 | 12709367 | 19.17 | 19.17 | 14.04 | 14.04 | 15.01 | 15.01 | 14.77 | 14.77 | 13.50 | 13.50 | 13.95 | 13.95 |
| TSS | ENSSSCG00000000161 | 5 | 12730655 | 12730726 | 19.17 | 19.17 | 14.04 | 14.04 | 15.01 | 15.01 | 14.77 | 14.77 | 13.50 | 13.50 | 13.95 | 13.95 |
| TTS | ENSSSCG00000000161 | 5 | 12703304 | 12703413 | 19.17 | 19.17 | 14.04 | 14.04 | 15.01 | 15.01 | 14.77 | 14.77 | 13.50 | 13.50 | 13.95 | 13.95 |
| SKIP_OFF | ENSSSCG00000000162 | 5 | 12797065 | 12797166 | 0.19 | 0.00 | 0.58 | 0.03 | 0.58 | 0.00 | 1.42 | 0.06 | 0.57 | 0.00 | 0.56 | 0.00 |
| MSKIP_OFF | ENSSSCG00000000162 | 5 | 12793906 | 12794985 | 0.19 | 0.00 | 0.58 | 0.00 | 0.58 | 0.07 | 1.42 | 0.00 | 0.57 | 0.00 | 0.56 | 0.00 |
| MSKIP_ON | ENSSSCG00000000162 | 5 | 12793906 | 12794985 | 0.19 | 0.19 | 0.58 | 0.58 | 0.58 | 0.50 | 1.42 | 1.42 | 0.57 | 0.57 | 0.56 | 0.56 |
| SKIP_ON | ENSSSCG00000000162 | 5 | 12797065 | 12797166 | 0.19 | 0.19 | 0.58 | 0.54 | 0.58 | 0.58 | 1.42 | 1.36 | 0.57 | 0.57 | 0.56 | 0.56 |
| TSS | ENSSSCG00000000162 | 5 | 13067791 | 13069699 | 0.19 | 0.19 | 0.58 | 0.58 | 0.58 | 0.58 | 1.42 | 1.42 | 0.57 | 0.57 | 0.56 | 0.56 |
| TTS | ENSSSCG00000000162 | 5 | 12754539 | 12756736 | 0.19 | 0.19 | 0.58 | 0.58 | 0.58 | 0.58 | 1.42 | 1.42 | 0.57 | 0.57 | 0.56 | 0.56 |
| IR_ON | ENSSSCG00000000164 | 5 | 13362997 | 13363077 | 11.33 | 3.21 | 8.05 | 2.47 | 9.16 | 2.22 | 15.33 | 5.11 | 7.83 | 1.74 | 13.44 | 3.93 |
| IR_OFF | ENSSSCG00000000164 | 5 | 13356674 | 13356738 | 11.33 | 4.71 | 8.05 | 2.56 | 9.16 | 2.94 | 15.33 | 5.15 | 7.83 | 3.08 | 13.44 | 3.90 |
| IR_OFF | ENSSSCG00000000164 | 5 | 13362997 | 13363077 | 11.33 | 3.41 | 8.05 | 3.02 | 9.16 | 4.01 | 15.33 | 5.07 | 7.83 | 3.00 | 13.44 | 5.62 |

| | | | | | | | | | | | | | | | | |
|---|---|---|---|---|---|---|---|---|---|---|---|---|---|---|---|---|
| SKIP_OFF | ENSSSCG00000000164 | 5 | 13356245 | 13356385 | 11.33 | 3.41 | 8.05 | 3.02 | 9.16 | 4.01 | 15.33 | 5.07 | 7.83 | 3.00 | 13.44 | 5.62 |
| SKIP_ON | ENSSSCG00000000164 | 5 | 13356245 | 13356385 | 11.33 | 7.92 | 8.05 | 5.03 | 9.16 | 5.15 | 15.33 | 10.27 | 7.83 | 4.82 | 13.44 | 7.82 |
| IR_ON | ENSSSCG00000000164 | 5 | 13356674 | 13356738 | 11.33 | 6.62 | 8.05 | 5.49 | 9.16 | 6.22 | 15.33 | 10.18 | 7.83 | 4.74 | 13.44 | 9.54 |
| TTS | ENSSSCG00000000164 | 5 | 13363959 | 13364594 | 11.33 | 6.62 | 8.05 | 5.49 | 9.16 | 6.22 | 15.33 | 10.18 | 7.83 | 4.74 | 13.44 | 9.54 |
| TSS | ENSSSCG00000000164 | 5 | 13275541 | 13276334 | 11.33 | 11.33 | 8.05 | 8.05 | 9.16 | 9.16 | 15.33 | 15.33 | 7.83 | 7.83 | 13.44 | 13.44 |
| SKIP_OFF | ENSSSCG00000000167 | 5 | 13570590 | 13570637 | 8.60 | 0.00 | 7.24 | 0.00 | 8.26 | 0.00 | 8.08 | 0.00 | 7.38 | 0.00 | 7.01 | 0.00 |
| TSS | ENSSSCG00000000167 | 5 | 13585717 | 13585781 | 8.60 | 0.49 | 7.24 | 0.33 | 8.26 | 0.41 | 8.08 | 0.35 | 7.38 | 0.29 | 7.01 | 0.71 |
| SKIP_ON | ENSSSCG00000000167 | 5 | 13486401 | 13486520 | 8.60 | 0.49 | 7.24 | 0.33 | 8.26 | 0.41 | 8.08 | 0.35 | 7.38 | 0.29 | 7.01 | 0.71 |
| SKIP_OFF | ENSSSCG00000000167 | 5 | 13486401 | 13486520 | 8.60 | 6.21 | 7.24 | 5.18 | 8.26 | 5.94 | 8.08 | 5.48 | 7.38 | 5.02 | 7.01 | 4.69 |
| TTS | ENSSSCG00000000167 | 5 | 13468944 | 13472409 | 8.60 | 6.70 | 7.24 | 5.51 | 8.26 | 6.35 | 8.08 | 5.83 | 7.38 | 5.31 | 7.01 | 5.40 |
| SKIP_ON | ENSSSCG00000000167 | 5 | 13570590 | 13570637 | 8.60 | 8.10 | 7.24 | 6.91 | 8.26 | 7.85 | 8.08 | 7.73 | 7.38 | 7.09 | 7.01 | 6.30 |
| TSS | ENSSSCG00000000167 | 5 | 13579953 | 13580164 | 8.60 | 8.10 | 7.24 | 6.91 | 8.26 | 7.85 | 8.08 | 7.73 | 7.38 | 7.09 | 7.01 | 6.30 |
| TSS | ENSSSCG00000000169 | 5 | 13758098 | 13758140 | 5.23 | 0.02 | 4.37 | 0.01 | 12.99 | 0.03 | 7.43 | 0.02 | 5.35 | 0.02 | 6.74 | 0.01 |
| TTS | ENSSSCG00000000169 | 5 | 13804930 | 13805703 | 5.23 | 5.19 | 4.37 | 4.33 | 12.99 | 12.91 | 7.43 | 7.08 | 5.35 | 4.15 | 6.74 | 5.91 |
| TSS | ENSSSCG00000000169 | 5 | 13926468 | 13926539 | 5.23 | 5.19 | 4.37 | 4.34 | 12.99 | 12.91 | 7.43 | 7.35 | 5.35 | 5.31 | 6.74 | 6.67 |
| TSS | ENSSSCG00000000171 | 5 | 14021321 | 14021776 | 50.66 | 50.66 | 28.70 | 28.70 | 38.04 | 38.04 | 11.86 | 11.86 | 21.06 | 21.06 | 31.69 | 31.69 |
| TTS | ENSSSCG00000000171 | 5 | 14028718 | 14030039 | 50.66 | 50.66 | 28.70 | 28.70 | 38.04 | 38.04 | 11.86 | 11.86 | 21.06 | 21.06 | 31.69 | 31.69 |
| TSS | ENSSSCG00000000175 | 5 | 14850477 | 14850728 | 15.73 | 15.73 | 12.70 | 12.70 | 15.02 | 15.02 | 10.40 | 10.40 | 11.30 | 11.30 | 13.70 | 13.70 |
| TTS | ENSSSCG00000000175 | 5 | 14836798 | 14836923 | 15.73 | 15.73 | 12.70 | 12.70 | 15.02 | 15.02 | 10.40 | 10.40 | 11.30 | 11.30 | 13.70 | 13.70 |
| TSS | ENSSSCG00000000176 | 5 | 14909604 | 14909741 | 13.24 | 13.24 | 9.75 | 9.75 | 12.14 | 12.14 | 11.96 | 11.96 | 11.51 | 11.51 | 12.88 | 12.88 |
| TTS | ENSSSCG00000000176 | 5 | 14886694 | 14889870 | 13.24 | 13.24 | 9.75 | 9.75 | 12.14 | 12.14 | 11.96 | 11.96 | 11.51 | 11.51 | 12.88 | 12.88 |
| TSS | ENSSSCG00000000179 | 5 | 14950492 | 14950839 | 2.44 | 2.44 | 0.49 | 0.49 | 0.73 | 0.73 | 21.73 | 21.73 | 11.76 | 11.76 | 18.04 | 18.04 |
| TTS | ENSSSCG00000000179 | 5 | 14958565 | 14959327 | 2.44 | 2.44 | 0.49 | 0.49 | 0.73 | 0.73 | 21.73 | 21.73 | 11.76 | 11.76 | 18.04 | 18.04 |
| TSS | ENSSSCG00000000180 | 5 | 14970662 | 14970752 | 23.45 | 23.45 | 18.27 | 18.27 | 22.11 | 22.11 | 32.88 | 32.88 | 20.83 | 20.83 | 36.70 | 36.70 |
| TTS | ENSSSCG00000000180 | 5 | 14958782 | 14959544 | 23.45 | 23.45 | 18.27 | 18.27 | 22.11 | 22.11 | 32.88 | 32.88 | 20.83 | 20.83 | 36.70 | 36.70 |
| IR_OFF | ENSSSCG00000000182 | 5 | 14999806 | 14999949 | 0.70 | 0.26 | 1.31 | 0.68 | 0.76 | 0.23 | 0.24 | 0.15 | 0.37 | 0.21 | 0.67 | 0.31 |
| IR_ON | ENSSSCG00000000182 | 5 | 14999806 | 14999949 | 0.70 | 0.43 | 1.31 | 0.63 | 0.76 | 0.54 | 0.24 | 0.09 | 0.37 | 0.16 | 0.67 | 0.37 |
| TSS | ENSSSCG00000000182 | 5 | 15011691 | 15013613 | 0.70 | 0.43 | 1.31 | 0.63 | 0.76 | 0.54 | 0.24 | 0.09 | 0.37 | 0.16 | 0.67 | 0.37 |
| TTS | ENSSSCG00000000182 | 5 | 14995704 | 14998577 | 0.70 | 0.70 | 1.31 | 1.31 | 0.76 | 0.76 | 0.24 | 0.24 | 0.37 | 0.37 | 0.67 | 0.67 |
| SKIP_OFF | ENSSSCG00000000185 | 5 | 15035996 | 15036105 | 20.89 | 0.00 | 21.24 | 0.00 | 19.64 | 0.00 | 20.10 | 0.00 | 22.02 | 0.00 | 21.31 | 0.00 |
| TSS | ENSSSCG00000000185 | 5 | 15045088 | 15045102 | 20.89 | 0.00 | 21.24 | 0.00 | 19.64 | 0.00 | 20.10 | 0.00 | 22.02 | 0.00 | 21.31 | 0.00 |
| TTS | ENSSSCG00000000185 | 5 | 15033060 | 15033334 | 20.89 | 0.38 | 21.24 | 0.22 | 19.64 | 0.68 | 20.10 | 0.00 | 22.02 | 0.57 | 21.31 | 1.72 |
| TSS | ENSSSCG00000000185 | 5 | 15049313 | 15049439 | 20.89 | 0.93 | 21.24 | 0.93 | 19.64 | 1.07 | 20.10 | 1.56 | 22.02 | 0.73 | 21.31 | 0.42 |
| TSS | ENSSSCG00000000185 | 5 | 15035996 | 15036181 | 20.89 | 1.19 | 21.24 | 2.11 | 19.64 | 2.04 | 20.10 | 1.63 | 22.02 | 2.54 | 21.31 | 2.30 |
| SKIP_OFF | ENSSSCG00000000185 | 5 | 15034340 | 15034377 | 20.89 | 1.70 | 21.24 | 2.80 | 19.64 | 2.43 | 20.10 | 3.19 | 22.02 | 2.68 | 21.31 | 1.01 |
| TSS | ENSSSCG00000000185 | 5 | 15049571 | 15049656 | 20.89 | 18.77 | 21.24 | 18.20 | 19.64 | 16.53 | 20.10 | 16.91 | 22.02 | 18.75 | 21.31 | 18.58 |
| SKIP_ON | ENSSSCG00000000185 | 5 | 15034340 | 15034377 | 20.89 | 19.19 | 21.24 | 18.44 | 19.64 | 17.21 | 20.10 | 16.91 | 22.02 | 19.34 | 21.31 | 20.30 |
| SKIP_ON | ENSSSCG00000000185 | 5 | 15035996 | 15036105 | 20.89 | 19.70 | 21.24 | 19.13 | 19.64 | 17.60 | 20.10 | 18.47 | 22.02 | 19.48 | 21.31 | 19.01 |
| TSS | ENSSSCG00000000186 | 5 | 15097490 | 15097764 | 0.50 | 0.50 | 0.24 | 0.24 | 0.27 | 0.27 | 0.38 | 0.38 | 0.38 | 0.38 | 0.18 | 0.18 |
| TTS | ENSSSCG00000000186 | 5 | 15093609 | 15094920 | 0.50 | 0.50 | 0.24 | 0.24 | 0.27 | 0.27 | 0.38 | 0.38 | 0.38 | 0.38 | 0.18 | 0.18 |
| TSS | ENSSSCG00000000188 | 5 | 15118996 | 15119577 | 1.57 | 1.57 | 0.98 | 0.98 | 0.73 | 0.73 | 0.33 | 0.33 | 0.85 | 0.85 | 0.80 | 0.80 |
| TTS | ENSSSCG00000000188 | 5 | 15114527 | 15115537 | 1.57 | 1.57 | 0.98 | 0.98 | 0.73 | 0.73 | 0.33 | 0.33 | 0.85 | 0.85 | 0.80 | 0.80 |
| MSKIP_OFF | ENSSSCG00000000189 | 5 | 15130504 | 15131765 | 10.35 | 0.17 | 9.76 | 0.00 | 12.74 | 0.00 | 14.15 | 0.42 | 14.13 | 0.03 | 10.98 | 0.46 |
| SKIP_OFF | ENSSSCG00000000189 | 5 | 15124452 | 15124529 | 10.35 | 0.00 | 9.76 | 0.00 | 12.74 | 0.22 | 14.15 | 0.00 | 14.13 | 0.00 | 10.98 | 0.00 |
| TSS | ENSSSCG00000000189 | 5 | 15135226 | 15135252 | 10.35 | 9.87 | 9.76 | 9.40 | 12.74 | 12.19 | 14.15 | 13.55 | 14.13 | 13.40 | 10.98 | 10.30 |
| SKIP_ON | ENSSSCG00000000189 | 5 | 15124452 | 15124529 | 10.35 | 10.35 | 9.76 | 9.76 | 12.74 | 12.53 | 14.15 | 14.15 | 14.13 | 14.13 | 10.98 | 10.98 |
| MSKIP_ON | ENSSSCG00000000189 | 5 | 15130504 | 15131765 | 10.35 | 10.18 | 9.76 | 9.76 | 12.74 | 12.74 | 14.15 | 13.74 | 14.13 | 14.10 | 10.98 | 10.52 |
| TTS | ENSSSCG00000000189 | 5 | 15119638 | 15122154 | 10.35 | 10.35 | 9.76 | 9.76 | 12.74 | 12.74 | 14.15 | 14.15 | 14.13 | 14.13 | 10.98 | 10.98 |
| TSS | ENSSSCG00000000190 | 5 | 15190076 | 15190862 | 416.62 | 50.69 | 276.00 | 43.50 | 202.65 | 25.96 | 312.52 | 45.43 | 262.27 | 41.59 | 265.03 | 39.38 |
| TTS | ENSSSCG00000000190 | 5 | 15150240 | 15150804 | 416.62 | 145.17 | 276.00 | 42.40 | 202.65 | 17.06 | 312.52 | 19.01 | 262.27 | 31.79 | 265.03 | 39.95 |

| | | | | | | | | | | | | | | | |
|---|---|---|---|---|---|---|---|---|---|---|---|---|---|---|---|
| TSS | ENSSSCG00000000190 | 5 | 15189714 | 15190062 | 416.62 | 170.24 | 276.00 | 83.55 | 202.65 | 62.50 | 312.52 | 117.57 | 262.27 | 113.01 | 265.03 | 108.42 |
| TSS | ENSSSCG00000000190 | 5 | 15153516 | 15153620 | 416.62 | 195.69 | 276.00 | 148.95 | 202.65 | 114.19 | 312.52 | 149.53 | 262.27 | 107.67 | 265.03 | 117.22 |
| TSS | ENSSSCG00000000191 | 5 | 15083878 | 15084043 | 4.68 | 4.68 | 6.38 | 6.38 | 6.05 | 6.05 | 5.13 | 5.13 | 6.68 | 6.68 | 4.49 | 4.49 |
| TTS | ENSSSCG00000000191 | 5 | 15058089 | 15058595 | 4.68 | 4.68 | 6.38 | 6.38 | 6.05 | 6.05 | 5.13 | 5.13 | 6.68 | 6.68 | 4.49 | 4.49 |
| TSS | ENSSSCG00000000194 | 5 | 15153184 | 15153599 | 86.78 | 18.33 | 33.77 | 3.44 | 20.49 | 2.35 | 45.30 | 27.44 | 30.83 | 11.90 | 33.50 | 10.59 |
| TSS | ENSSSCG00000000194 | 5 | 15260842 | 15261045 | 86.78 | 68.46 | 33.77 | 30.34 | 20.49 | 18.13 | 45.30 | 17.86 | 30.83 | 18.93 | 33.50 | 22.91 |
| TTS | ENSSSCG00000000194 | 5 | 15268707 | 15269779 | 86.78 | 86.78 | 33.77 | 33.77 | 20.49 | 20.49 | 45.30 | 45.30 | 30.83 | 30.83 | 33.50 | 33.50 |
| TTS | ENSSSCG00000000195 | 5 | 15288398 | 15289524 | 0.29 | 0.29 | 0.34 | 0.24 | 0.26 | 0.26 | 0.10 | 0.10 | 0.19 | 0.19 | 0.12 | 0.12 |
| TSS | ENSSSCG00000000195 | 5 | 15285252 | 15285891 | 0.29 | 0.29 | 0.34 | 0.34 | 0.26 | 0.26 | 0.10 | 0.10 | 0.19 | 0.19 | 0.12 | 0.12 |
| TSS | ENSSSCG00000000197 | 5 | 15311180 | 15312520 | 3.65 | 0.00 | 1.49 | 0.00 | 2.66 | 0.00 | 8.24 | 0.00 | 2.70 | 0.00 | 0.77 | 0.00 |
| TTS | ENSSSCG00000000197 | 5 | 15308282 | 15309162 | 3.65 | 0.00 | 1.49 | 0.00 | 2.66 | 0.00 | 8.24 | 0.00 | 2.70 | 0.00 | 0.77 | 0.00 |
| TSS | ENSSSCG00000000198 | 5 | 15316383 | 15316951 | 0.23 | 0.00 | 0.00 | 0.00 | 0.37 | 0.00 | 0.96 | 0.00 | 0.52 | 0.00 | 1.05 | 0.00 |
| TSS | ENSSSCG00000000198 | 5 | 15322508 | 15322771 | 0.23 | 0.23 | 0.00 | 0.00 | 0.37 | 0.37 | 0.96 | 0.96 | 0.52 | 0.52 | 1.05 | 1.05 |
| TTS | ENSSSCG00000000198 | 5 | 15334737 | 15335026 | 0.23 | 0.23 | 0.00 | 0.00 | 0.37 | 0.37 | 0.96 | 0.96 | 0.52 | 0.52 | 1.05 | 1.05 |
| SKIP_ON | ENSSSCG00000000199 | 5 | 15471648 | 15471693 | 9.61 | 0.21 | 6.41 | 0.11 | 9.85 | 0.14 | 10.78 | 0.00 | 8.86 | 0.11 | 11.98 | 0.20 |
| SKIP_OFF | ENSSSCG00000000199 | 5 | 15471648 | 15471693 | 9.61 | 9.40 | 6.41 | 6.30 | 9.85 | 9.71 | 10.78 | 10.78 | 8.86 | 8.74 | 11.98 | 11.78 |
| TSS | ENSSSCG00000000199 | 5 | 15347633 | 15347928 | 9.61 | 9.61 | 6.41 | 6.41 | 9.85 | 9.85 | 10.78 | 10.78 | 8.86 | 8.86 | 11.98 | 11.98 |
| TTS | ENSSSCG00000000199 | 5 | 15483736 | 15484756 | 9.61 | 9.61 | 6.41 | 6.41 | 9.85 | 9.85 | 10.78 | 10.78 | 8.86 | 8.86 | 11.98 | 11.98 |
| IR_ON | ENSSSCG00000000202 | 5 | 15524605 | 15524794 | 14.90 | 2.06 | 11.62 | 1.57 | 13.84 | 4.12 | 11.71 | 2.93 | 11.93 | 2.46 | 13.20 | 1.26 |
| IR_OFF | ENSSSCG00000000202 | 5 | 15524605 | 15524794 | 14.90 | 12.84 | 11.62 | 10.05 | 13.84 | 9.72 | 11.71 | 8.78 | 11.93 | 9.47 | 13.20 | 11.93 |
| TSS | ENSSSCG00000000202 | 5 | 15526422 | 15526535 | 14.90 | 14.90 | 11.62 | 11.62 | 13.84 | 13.84 | 11.71 | 11.71 | 11.93 | 11.93 | 13.20 | 13.20 |
| TTS | ENSSSCG00000000202 | 5 | 15517033 | 15517464 | 14.90 | 14.90 | 11.62 | 11.62 | 13.84 | 13.84 | 11.71 | 11.71 | 11.93 | 11.93 | 13.20 | 13.20 |
| TSS | ENSSSCG00000000203 | 5 | 15495902 | 15496975 | 2.09 | 0.94 | 1.02 | 0.20 | 0.85 | 0.33 | 7.08 | 3.49 | 4.94 | 2.86 | 6.56 | 2.88 |
| TSS | ENSSSCG00000000203 | 5 | 15495251 | 15495326 | 2.09 | 1.16 | 1.02 | 0.82 | 0.85 | 0.52 | 7.08 | 3.59 | 4.94 | 2.08 | 6.56 | 3.68 |
| TTS | ENSSSCG00000000203 | 5 | 15516051 | 15517458 | 2.09 | 2.09 | 1.02 | 1.02 | 0.85 | 0.85 | 7.08 | 7.08 | 4.94 | 4.94 | 6.56 | 6.56 |
| SKIP_ON | ENSSSCG00000000204 | 5 | 15577357 | 15577437 | 8.38 | 0.51 | 6.73 | 1.25 | 7.21 | 0.34 | 11.03 | 0.31 | 8.87 | 0.39 | 8.46 | 0.18 |
| SKIP_OFF | ENSSSCG00000000204 | 5 | 15577357 | 15577437 | 8.38 | 7.87 | 6.73 | 5.48 | 7.21 | 6.86 | 11.03 | 10.73 | 8.87 | 8.48 | 8.46 | 8.28 |
| TTS | ENSSSCG00000000204 | 5 | 15591821 | 15596650 | 8.38 | 7.87 | 6.73 | 5.48 | 7.21 | 6.86 | 11.03 | 10.73 | 8.87 | 8.48 | 8.46 | 8.28 |
| TSS | ENSSSCG00000000204 | 5 | 15570454 | 15570659 | 8.38 | 8.38 | 6.73 | 6.73 | 7.21 | 7.21 | 11.03 | 11.03 | 8.87 | 8.87 | 8.46 | 8.46 |
| TSS | ENSSSCG00000000205 | 5 | 15559626 | 15560066 | 1.40 | 1.40 | 0.91 | 0.91 | 0.89 | 0.89 | 1.00 | 1.00 | 1.76 | 1.76 | 1.71 | 1.71 |
| TTS | ENSSSCG00000000205 | 5 | 15542246 | 15542345 | 1.40 | 1.40 | 0.91 | 0.91 | 0.89 | 0.89 | 1.00 | 1.00 | 1.76 | 1.76 | 1.71 | 1.71 |
| TSS | ENSSSCG00000000206 | 5 | 15812439 | 15813034 | 7.37 | 7.37 | 10.05 | 10.05 | 6.22 | 6.22 | 14.79 | 14.79 | 7.26 | 7.26 | 2.63 | 2.63 |
| TTS | ENSSSCG00000000206 | 5 | 15779334 | 15782890 | 7.37 | 7.37 | 10.05 | 10.05 | 6.22 | 6.22 | 14.79 | 14.79 | 7.26 | 7.26 | 2.63 | 2.63 |
| TSS | ENSSSCG00000000207 | 5 | 15774938 | 15775227 | 0.31 | 0.31 | 0.41 | 0.41 | 0.63 | 0.63 | 0.67 | 0.67 | 0.95 | 0.95 | 0.33 | 0.33 |
| TTS | ENSSSCG00000000207 | 5 | 15775251 | 15775449 | 0.31 | 0.31 | 0.41 | 0.41 | 0.63 | 0.63 | 0.67 | 0.67 | 0.95 | 0.95 | 0.33 | 0.33 |
| SKIP_OFF | ENSSSCG00000000209 | 5 | 15725036 | 15725097 | 4.60 | 0.73 | 2.92 | 0.00 | 5.36 | 0.00 | 3.26 | 0.17 | 4.48 | 0.00 | 3.98 | 0.00 |
| SKIP_ON | ENSSSCG00000000209 | 5 | 15725036 | 15725097 | 4.60 | 1.53 | 2.92 | 1.40 | 5.36 | 2.23 | 3.26 | 1.33 | 4.48 | 2.17 | 3.98 | 2.56 |
| TSS | ENSSSCG00000000209 | 5 | 15742295 | 15742388 | 4.60 | 2.27 | 2.92 | 1.40 | 5.36 | 2.24 | 3.26 | 1.50 | 4.48 | 2.17 | 3.98 | 2.56 |
| TTS | ENSSSCG00000000209 | 5 | 15711254 | 15712132 | 4.60 | 2.33 | 2.92 | 1.52 | 5.36 | 3.13 | 3.26 | 1.76 | 4.48 | 2.31 | 3.98 | 1.42 |
| TSS | ENSSSCG00000000209 | 5 | 15761292 | 15761537 | 4.60 | 5.74 | 2.92 | 4.70 | 5.36 | 6.39 | 3.26 | 5.46 | 4.48 | 5.91 | 3.98 | 4.45 |
| TSS | ENSSSCG00000000211 | 5 | 15876401 | 15876890 | 0.17 | 0.17 | 0.13 | 0.13 | 0.13 | 0.13 | 0.26 | 0.26 | 0.05 | 0.05 | 0.50 | 0.50 |
| TTS | ENSSSCG00000000211 | 5 | 15879345 | 15881332 | 0.17 | 0.17 | 0.13 | 0.13 | 0.13 | 0.13 | 0.26 | 0.26 | 0.05 | 0.05 | 0.50 | 0.50 |
| TSS | ENSSSCG00000000212 | 5 | 15887947 | 15888678 | 0.13 | 0.13 | 0.06 | 0.06 | 0.19 | 0.19 | 0.10 | 0.10 | 0.23 | 0.23 | 0.12 | 0.12 |
| TTS | ENSSSCG00000000212 | 5 | 15890590 | 15892148 | 0.13 | 0.13 | 0.06 | 0.06 | 0.19 | 0.19 | 0.10 | 0.10 | 0.23 | 0.23 | 0.12 | 0.12 |
| SKIP_OFF | ENSSSCG00000000214 | 5 | 16012723 | 16012829 | 5.19 | 0.22 | 2.76 | 0.00 | 4.05 | 0.00 | 6.75 | 0.00 | 4.87 | 0.00 | 6.68 | 0.34 |
| TSS | ENSSSCG00000000214 | 5 | 16008485 | 16008530 | 5.19 | 0.22 | 2.76 | 0.00 | 4.05 | 0.00 | 6.75 | 0.00 | 4.87 | 0.00 | 6.68 | 0.34 |
| TSS | ENSSSCG00000000214 | 5 | 16008533 | 16008824 | 5.19 | 2.01 | 2.76 | 0.53 | 4.05 | 0.83 | 6.75 | 3.73 | 4.87 | 2.24 | 6.68 | 2.19 |
| TSS | ENSSSCG00000000214 | 5 | 16007448 | 16007793 | 5.19 | 2.95 | 2.76 | 2.23 | 4.05 | 3.21 | 6.75 | 3.02 | 4.87 | 2.63 | 6.68 | 4.14 |
| TTS | ENSSSCG00000000214 | 5 | 16021020 | 16021674 | 5.19 | 2.95 | 2.76 | 2.23 | 4.05 | 3.21 | 6.75 | 3.02 | 4.87 | 2.63 | 6.68 | 4.14 |
| SKIP_ON | ENSSSCG00000000214 | 5 | 16012723 | 16012829 | 5.19 | 4.96 | 2.76 | 2.76 | 4.05 | 4.05 | 6.75 | 6.75 | 4.87 | 4.87 | 6.68 | 6.33 |

### Table S4 Differentially spliced events

| data...1. | event_type | gene_id | chr | event start | event end | XL13 | XL17 | XL18 | XS12 | XS15 | XS16 | XL AveFPKM | XS AveFPKM | FC | logFC | AveExpr | t | P.Value | adj.P.Val | B |
|---|---|---|---|---|---|---|---|---|---|---|---|---|---|---|---|---|---|---|---|---|
| 1058034 | TTS | ENSSSCG00000002768 | 6 | 28446687 | 28446702 | 1.42 | 0.93 | 1.36 | 0.00 | 0.00 | 0.00 | 1.24 | 0.00 | 695762.58 | 19.41 | 0.62 | -8.18 | 0.0007 | 0.038 | 0.22 |
| 1000765 | TSS | ENSSSCG00000004263 | 1 | 44940794 | 44940850 | 1.55 | 1.24 | 1.02 | 0.00 | 0.00 | 0.00 | 1.27 | 0.00 | 51279.67 | 15.65 | 0.63 | -8.40 | 0.0006 | 0.036 | 0.35 |
| 1019643 | TSS | ENSSSCG00000029811 | 13 | 3980464 | 3980557 | 0.71 | 0.92 | 0.84 | 0.00 | 0.00 | 0.00 | 0.83 | 0.00 | 5943.19 | 12.54 | 0.41 | -11.90 | 0.0001 | 0.020 | 1.96 |
| 1014231 | SKIP_OFF | ENSSSCG00000033228 | 12 | 27261499 | 27261513 | 2.15 | 1.93 | 1.54 | 0.00 | 0.00 | 0.00 | 1.87 | 0.00 | 4076.38 | 11.99 | 0.94 | -10.91 | 0.0002 | 0.023 | 1.57 |
| 1014220 | SKIP_ON | ENSSSCG00000033228 | 12 | 27262711 | 27262722 | 2.15 | 1.93 | 1.54 | 0.00 | 0.00 | 0.00 | 1.87 | 0.00 | 4076.38 | 11.99 | 0.94 | -10.91 | 0.0002 | 0.023 | 1.57 |
| 1049709 | TSS | ENSSSCG00000006191 | 4 | 64710542 | 64710633 | 0.90 | 0.76 | 0.79 | 0.00 | 0.00 | 0.00 | 0.82 | 0.00 | 3976.77 | 11.96 | 0.41 | -14.88 | 0.0001 | 0.015 | 2.93 |
| 1023585 | SKIP_OFF | ENSSSCG00000010281 | 14 | 74747072 | 74747197 | 4.37 | 5.26 | 5.11 | 0.00 | 0.00 | 0.02 | 4.91 | 0.01 | 761.07 | 9.57 | 2.46 | -18.82 | 0.0000 | 0.010 | 3.85 |
| 1069392 | SKIP_OFF | ENSSSCG00000028225 | 7 | 116374054 | 116374209 | 0.31 | 0.33 | 0.31 | 0.00 | 0.00 | 0.00 | 0.32 | 0.00 | 728.24 | 9.51 | 0.16 | -8.23 | 0.0007 | 0.037 | 0.25 |
| 1029818 | TTS | ENSSSCG00000034942 | 15 | 48383713 | 48388326 | 107.11 | 101.29 | 85.77 | 0.41 | 2.00 | 2.25 | 98.06 | 1.55 | 63.07 | 5.98 | 49.81 | -16.03 | 0.0000 | 0.013 | 3.23 |
| 1029817 | TSS | ENSSSCG00000034942 | 15 | 48377513 | 48377756 | 111.90 | 104.71 | 87.37 | 0.50 | 2.11 | 2.33 | 101.33 | 1.65 | 61.40 | 5.94 | 51.49 | -14.51 | 0.0001 | 0.015 | 2.82 |
| 1075974 | SKIP_OFF | ENSSSCG00000015537 | 9 | 122359024 | 122359125 | 0.51 | 0.53 | 0.42 | 0.00 | 0.03 | 0.00 | 0.49 | 0.01 | 41.96 | 5.39 | 0.25 | -9.42 | 0.0004 | 0.030 | 0.89 |
| 1067382 | MSKIP_OFF | ENSSSCG00000001825 | 7 | 53686932 | 53687906 | 1.26 | 1.12 | 1.14 | 0.00 | 0.08 | 0.00 | 1.18 | 0.03 | 41.89 | 5.39 | 0.60 | -18.80 | 0.0000 | 0.010 | 3.85 |
| 1024037 | TSS | ENSSSCG00000010455 | 14 | 101367855 | 101368158 | 0.98 | 0.61 | 0.86 | 0.08 | 0.00 | 0.00 | 0.81 | 0.03 | 31.72 | 4.99 | 0.42 | -7.09 | 0.0013 | 0.047 | -0.45 |
| 1016020 | TSS | ENSSSCG00000011423 | 13 | 33926804 | 33926915 | 0.34 | 0.27 | 0.28 | 0.01 | 0.03 | 0.02 | 0.30 | 0.02 | 16.50 | 4.04 | 0.16 | -6.47 | 0.0019 | 0.055 | -0.88 |
| 1032155 | MSKIP_OFF | ENSSSCG00000007133 | 17 | 30804841 | 30805130 | 0.70 | 1.03 | 0.76 | 0.17 | 0.00 | 0.00 | 0.83 | 0.06 | 14.43 | 3.85 | 0.44 | -6.61 | 0.0017 | 0.053 | -0.78 |
| 1032151 | XSKIP_OFF | ENSSSCG00000007133 | 17 | 30805003 | 30805130 | 0.70 | 1.03 | 0.76 | 0.17 | 0.00 | 0.00 | 0.83 | 0.06 | 14.43 | 3.85 | 0.44 | -6.61 | 0.0017 | 0.053 | -0.78 |
| 1031358 | TSS | ENSSSCG00000030343 | 16 | 34871780 | 34871822 | 0.60 | 0.77 | 0.62 | 0.00 | 0.00 | 0.19 | 0.66 | 0.06 | 10.55 | 3.40 | 0.36 | -6.99 | 0.0014 | 0.048 | -0.52 |
| 1031365 | XAE | ENSSSCG00000030343 | 16 | 34839254 | 34839386 | 0.60 | 0.77 | 0.62 | 0.00 | 0.00 | 0.19 | 0.66 | 0.06 | 10.55 | 3.40 | 0.36 | -6.99 | 0.0014 | 0.048 | -0.52 |
| 1000555 | TSS | ENSSSCG00000004195 | 1 | 32051209 | 32051376 | 1.39 | 1.37 | 1.53 | 0.16 | 0.26 | 0.00 | 1.43 | 0.14 | 10.28 | 3.36 | 0.79 | -13.70 | 0.0001 | 0.017 | 2.58 |
| 1008290 | TTS | ENSSSCG00000011147 | 10 | 65572686 | 65574571 | 119.94 | 95.45 | 81.49 | 1.17 | 16.69 | 16.71 | 98.96 | 11.52 | 8.59 | 3.10 | 55.24 | -7.51 | 0.0010 | 0.043 | -0.18 |
| 1019143 | MSKIP_OFF | ENSSSCG00000026746 | 13 | 79332564 | 79332882 | 1.09 | 1.00 | 1.10 | 0.13 | 0.26 | 0.00 | 1.06 | 0.13 | 8.10 | 3.02 | 0.60 | -10.75 | 0.0002 | 0.024 | 1.50 |
| 1019148 | SKIP_OFF | ENSSSCG00000026746 | 13 | 79330784 | 79330858 | 1.09 | 1.00 | 1.10 | 0.13 | 0.26 | 0.00 | 1.06 | 0.13 | 8.10 | 3.02 | 0.60 | -10.75 | 0.0002 | 0.024 | 1.50 |
| 1045599 | SKIP_OFF | ENSSSCG00000008421 | 3 | 92002381 | 92002550 | 23.42 | 15.59 | 20.60 | 0.12 | 5.72 | 2.14 | 19.87 | 2.66 | 7.47 | 2.90 | 11.26 | -6.50 | 0.0019 | 0.054 | -0.86 |
| 1062615 | TSS | ENSSSCG00000027684 | 6 | 83507215 | 83508007 | 0.61 | 0.50 | 0.57 | 0.01 | 0.16 | 0.06 | 0.56 | 0.08 | 7.34 | 2.87 | 0.32 | -7.53 | 0.0010 | 0.043 | -0.17 |
| 1062616 | TTS | ENSSSCG00000027684 | 6 | 83488973 | 83493215 | 0.61 | 0.50 | 0.57 | 0.01 | 0.16 | 0.06 | 0.56 | 0.08 | 7.34 | 2.87 | 0.32 | -7.53 | 0.0010 | 0.043 | -0.17 |
| 1045593 | TSS | ENSSSCG00000008421 | 3 | 91964415 | 91964639 | 24.13 | 16.40 | 21.30 | 0.12 | 6.08 | 2.36 | 20.61 | 2.85 | 7.22 | 2.85 | 11.73 | -6.62 | 0.0017 | 0.053 | -0.77 |
| 1045594 | TTS | ENSSSCG00000008421 | 3 | 92020227 | 92021398 | 21.40 | 15.59 | 20.17 | 0.12 | 6.08 | 2.22 | 19.05 | 2.81 | 6.79 | 2.76 | 10.93 | -6.95 | 0.0014 | 0.049 | -0.55 |
| 1045601 | XSKIP_OFF | ENSSSCG00000008421 | 3 | 92015950 | 92015950 | 21.40 | 15.59 | 20.17 | 0.12 | 6.08 | 2.22 | 19.05 | 2.81 | 6.79 | 2.76 | 10.93 | -6.95 | 0.0014 | 0.049 | -0.55 |
| 1045596 | SKIP_ON | ENSSSCG00000008421 | 3 | 91984228 | 91984299 | 21.40 | 15.59 | 20.17 | 0.12 | 6.08 | 2.22 | 19.05 | 2.81 | 6.79 | 2.76 | 10.93 | -6.95 | 0.0014 | 0.049 | -0.55 |
| 1010172 | TSS | ENSSSCG00000022159 | 11 | 18756040 | 18756356 | 5.62 | 6.01 | 5.44 | 0.17 | 2.05 | 0.74 | 5.69 | 0.98 | 5.78 | 2.53 | 3.34 | -8.59 | 0.0006 | 0.035 | 0.45 |
| 1000556 | TTS | ENSSSCG00000004195 | 1 | 32006042 | 32007153 | 1.72 | 1.37 | 1.97 | 0.26 | 0.48 | 0.19 | 1.68 | 0.31 | 5.44 | 2.44 | 1.00 | -7.36 | 0.0011 | 0.045 | -0.27 |
| 1079470 | TSS | ENSSSCG00000012638 | X | 100826849 | 100827728 | 13.78 | 17.69 | 16.20 | 0.83 | 5.48 | 2.63 | 15.89 | 2.98 | 5.33 | 2.41 | 9.44 | -7.76 | 0.0009 | 0.041 | -0.02 |
| 1023991 | SKIP_ON | ENSSSCG00000010437 | 14 | 99783084 | 99783098 | 45.82 | 54.54 | 58.86 | 1.47 | 18.39 | 14.04 | 53.07 | 11.30 | 4.70 | 2.23 | 32.19 | -6.98 | 0.0014 | 0.049 | -0.52 |
| 1018359 | TTS | ENSSSCG00000021966 | 13 | 74766604 | 74766639 | 0.92 | 1.04 | 0.80 | 0.19 | 0.24 | 0.19 | 0.92 | 0.21 | 4.46 | 2.16 | 0.56 | -9.21 | 0.0004 | 0.031 | 0.78 |
| 1065608 | TTS | ENSSSCG00000040607 | 6 | 8484601 | 8485030 | 2.36 | 2.82 | 2.31 | 0.16 | 1.07 | 0.53 | 2.49 | 0.59 | 4.24 | 2.08 | 1.54 | -6.49 | 0.0019 | 0.055 | -0.87 |
| 1080548 | TSS | ENSSSCG00000030241 | X | 88234558 | 88234608 | 38.74 | 32.90 | 32.73 | 3.64 | 8.80 | 12.52 | 34.79 | 8.32 | 4.18 | 2.06 | 21.56 | -8.67 | 0.0006 | 0.034 | 0.50 |
| 1039380 | SKIP_OFF | ENSSSCG00000014136 | 2 | 91768020 | 91773257 | 8.63 | 6.72 | 7.94 | 0.51 | 2.68 | 2.82 | 7.76 | 2.00 | 3.88 | 1.96 | 4.88 | -6.55 | 0.0018 | 0.054 | -0.82 |
| 1039385 | SKIP_ON | ENSSSCG00000014136 | 2 | 91746365 | 91749304 | 8.63 | 6.72 | 7.94 | 0.51 | 2.68 | 2.82 | 7.76 | 2.00 | 3.88 | 1.96 | 4.88 | -6.55 | 0.0018 | 0.054 | -0.82 |
| 1007330 | TSS | ENSSSCG00000010840 | 10 | 12322181 | 12322304 | 2.56 | 1.84 | 2.57 | 0.48 | 0.68 | 0.68 | 2.33 | 0.61 | 3.80 | 1.92 | 1.47 | -7.14 | 0.0013 | 0.047 | -0.42 |
| 1007336 | XSKIP_ON | ENSSSCG00000010840 | 10 | 12321629 | 12321642 | 2.56 | 1.84 | 2.57 | 0.48 | 0.68 | 0.68 | 2.33 | 0.61 | 3.80 | 1.92 | 1.47 | -7.14 | 0.0013 | 0.047 | -0.42 |
| 1048761 | TSS | ENSSSCG00000040786 | 3 | 86783366 | 86783498 | 11.93 | 10.98 | 11.60 | 2.06 | 3.13 | 3.95 | 11.50 | 3.05 | 3.78 | 1.92 | 7.27 | -14.66 | 0.0001 | 0.015 | 2.86 |
| 1004422 | SKIP_ON | ENSSSCG00000005585 | 1 | 264307939 | 264308067 | 1.51 | 1.32 | 1.07 | 0.36 | 0.44 | 0.30 | 1.30 | 0.37 | 3.54 | 1.82 | 0.83 | -6.97 | 0.0014 | 0.049 | -0.53 |
| 1078216 | TTS | ENSSSCG00000012112 | X | 7910855 | 7912885 | 15.47 | 21.90 | 20.62 | 6.49 | 5.77 | 6.20 | 19.33 | 6.15 | 3.14 | 1.65 | 12.74 | -7.09 | 0.0013 | 0.047 | -0.45 |
| 1057413 | SKIP_ON | ENSSSCG00000038732 | 5 | 23018912 | 23018971 | 1.48 | 1.61 | 1.18 | 0.48 | 0.46 | 0.47 | 1.42 | 0.47 | 3.03 | 1.60 | 0.95 | -7.50 | 0.0010 | 0.043 | -0.19 |
| 1067628 | SKIP_ON | ENSSSCG00000001914 | 7 | 59486916 | 59487053 | 28.94 | 28.08 | 25.52 | 3.98 | 10.76 | 12.66 | 27.52 | 9.13 | 3.01 | 1.59 | 18.32 | -6.91 | 0.0015 | 0.049 | -0.57 |
| 1067626 | TSS | ENSSSCG00000001914 | 7 | 59503667 | 59505269 | 29.42 | 28.08 | 25.63 | 4.25 | 10.86 | 12.76 | 27.71 | 9.29 | 2.98 | 1.58 | 18.50 | -6.97 | 0.0014 | 0.049 | -0.53 |
| 1067627 | TTS | ENSSSCG00000001914 | 7 | 59481407 | 59481749 | 29.42 | 28.08 | 25.63 | 4.25 | 10.86 | 12.76 | 27.71 | 9.29 | 2.98 | 1.58 | 18.50 | -6.97 | 0.0014 | 0.049 | -0.53 |
| 1080443 | TSS | ENSSSCG00000027465 | X | 57176103 | 57176254 | 3.55 | 3.43 | 3.92 | 1.25 | 1.47 | 1.01 | 3.63 | 1.24 | 2.92 | 1.54 | 2.44 | -12.57 | 0.0001 | 0.019 | 2.20 |
| 1080444 | TTS | ENSSSCG00000027465 | X | 57198622 | 57201459 | 3.55 | 3.43 | 3.92 | 1.25 | 1.47 | 1.01 | 3.63 | 1.24 | 2.92 | 1.54 | 2.44 | -12.57 | 0.0001 | 0.019 | 2.20 |

| | | | | | | | | | | | | | | | | | | | | |
|---|---|---|---|---|---|---|---|---|---|---|---|---|---|---|---|---|---|---|---|---|
| 1010176 | TSS | ENSSSCG00000022159 | 11 | 18757615 | 18757700 | 10.68 | 14.85 | 13.81 | 4.31 | 5.31 | 4.37 | 13.11 | 4.66 | 2.81 | 1.49 | 8.89 | -6.93 | 0.0014 | 0.049 | -0.56 |
| 1010181 | XSKIP_OFF | ENSSSCG00000022159 | 11 | 18756754 | 18756822 | 10.68 | 14.85 | 13.81 | 4.31 | 5.31 | 4.37 | 13.11 | 4.66 | 2.81 | 1.49 | 8.89 | -6.93 | 0.0014 | 0.049 | -0.56 |
| 1025836 | SKIP_OFF | ENSSSCG00000032622 | 14 | 6791993 | 6792022 | 4.08 | 3.47 | 3.51 | 0.85 | 1.50 | 1.81 | 3.69 | 1.39 | 2.66 | 1.41 | 2.54 | -7.01 | 0.0014 | 0.048 | -0.50 |
| 1042954 | SKIP_OFF | ENSSSCG00000039045 | 2 | 151018018 | 151018741 | 4.78 | 5.92 | 5.42 | 1.68 | 1.65 | 2.73 | 5.37 | 2.02 | 2.66 | 1.41 | 3.70 | -7.34 | 0.0011 | 0.045 | -0.29 |
| 1042957 | SKIP_ON | ENSSSCG00000039045 | 2 | 151012135 | 151012311 | 4.78 | 5.92 | 5.42 | 1.68 | 1.65 | 2.73 | 5.37 | 2.02 | 2.66 | 1.41 | 3.70 | -7.34 | 0.0011 | 0.045 | -0.29 |
| 1063447 | TSS | ENSSSCG00000032060 | 6 | 7255180 | 7255332 | 22.83 | 30.01 | 26.13 | 8.05 | 8.98 | 12.73 | 26.32 | 9.92 | 2.65 | 1.41 | 18.12 | -6.91 | 0.0015 | 0.049 | -0.57 |
| 1063449 | TTS | ENSSSCG00000032060 | 6 | 7278340 | 7278710 | 23.81 | 30.75 | 29.47 | 8.41 | 9.53 | 14.17 | 28.01 | 10.70 | 2.62 | 1.39 | 19.35 | -6.65 | 0.0017 | 0.053 | -0.76 |
| 1042059 | TSS | ENSSSCG00000033765 | 2 | 142983077 | 142985349 | 0.63 | 0.64 | 0.69 | 0.18 | 0.30 | 0.28 | 0.65 | 0.25 | 2.59 | 1.37 | 0.45 | -7.36 | 0.0011 | 0.045 | -0.28 |
| 1042060 | TTS | ENSSSCG00000033765 | 2 | 142985373 | 142985628 | 0.63 | 0.64 | 0.69 | 0.18 | 0.30 | 0.28 | 0.65 | 0.25 | 2.59 | 1.37 | 0.45 | -7.36 | 0.0011 | 0.045 | -0.28 |
| 1025041 | TSS | ENSSSCG00000010746 | 14 | 135672821 | 135672926 | 17.13 | 17.05 | 15.32 | 3.67 | 8.87 | 6.59 | 16.50 | 6.38 | 2.59 | 1.37 | 11.44 | -6.65 | 0.0017 | 0.053 | -0.75 |
| 1025043 | TTS | ENSSSCG00000010746 | 14 | 135302758 | 135307810 | 17.17 | 17.05 | 15.35 | 3.67 | 8.90 | 6.59 | 16.53 | 6.39 | 2.59 | 1.37 | 11.46 | -6.64 | 0.0017 | 0.053 | -0.76 |
| 1034342 | TSS | ENSSSCG00000016444 | 18 | 6342709 | 6343108 | 43.38 | 41.94 | 43.24 | 10.61 | 20.82 | 19.00 | 42.85 | 16.81 | 2.55 | 1.35 | 29.83 | -8.71 | 0.0005 | 0.034 | 0.52 |
| 1034343 | TTS | ENSSSCG00000016444 | 18 | 6348590 | 6348958 | 43.38 | 41.94 | 43.24 | 10.61 | 20.82 | 19.00 | 42.85 | 16.81 | 2.55 | 1.35 | 29.83 | -8.71 | 0.0005 | 0.034 | 0.52 |
| 1011118 | TSS | ENSSSCG00000017215 | 12 | 6197682 | 6197717 | 2.67 | 2.49 | 3.04 | 0.95 | 1.37 | 0.92 | 2.73 | 1.08 | 2.53 | 1.34 | 1.91 | -7.89 | 0.0008 | 0.040 | 0.05 |
| 1020290 | XMSKIP_ON | ENSSSCG00000034493 | 13 | 160191825 | 160204381 | 5.73 | 5.69 | 7.14 | 2.45 | 3.00 | 1.97 | 6.19 | 2.47 | 2.50 | 1.32 | 4.33 | -7.03 | 0.0014 | 0.048 | -0.49 |
| 1020271 | TSS | ENSSSCG00000034493 | 13 | 160224671 | 160224803 | 5.73 | 5.69 | 7.14 | 2.45 | 3.00 | 2.02 | 6.19 | 2.49 | 2.49 | 1.32 | 4.34 | -7.09 | 0.0013 | 0.047 | -0.45 |
| 1020293 | XSKIP_OFF | ENSSSCG00000034493 | 13 | 160204668 | 160204775 | 5.73 | 5.69 | 7.14 | 2.45 | 3.00 | 2.02 | 6.19 | 2.49 | 2.49 | 1.32 | 4.34 | -7.09 | 0.0013 | 0.047 | -0.45 |
| 1049789 | TTS | ENSSSCG00000006206 | 4 | 67899254 | 67901126 | 8.27 | 7.36 | 9.44 | 2.71 | 4.35 | 3.15 | 8.36 | 3.40 | 2.45 | 1.30 | 5.88 | -6.78 | 0.0016 | 0.051 | -0.66 |
| 1005309 | TSS | ENSSSCG00000026042 | 1 | 16119103 | 16119704 | 0.80 | 0.78 | 0.85 | 0.38 | 0.35 | 0.26 | 0.81 | 0.33 | 2.45 | 1.29 | 0.57 | -8.56 | 0.0006 | 0.035 | 0.44 |
| 1005310 | TTS | ENSSSCG00000026042 | 1 | 16112178 | 16114324 | 0.80 | 0.78 | 0.85 | 0.38 | 0.35 | 0.26 | 0.81 | 0.33 | 2.45 | 1.29 | 0.57 | -8.56 | 0.0006 | 0.035 | 0.44 |
| 1053460 | TSS | ENSSSCG00000000038 | 5 | 6152922 | 6152942 | 139.14 | 147.27 | 162.53 | 50.47 | 60.85 | 74.59 | 149.65 | 61.97 | 2.41 | 1.27 | 105.81 | -9.52 | 0.0004 | 0.029 | 0.94 |
| 1053461 | TTS | ENSSSCG00000000038 | 5 | 6169090 | 6169262 | 139.14 | 147.27 | 162.53 | 50.47 | 60.85 | 74.59 | 149.65 | 61.97 | 2.41 | 1.27 | 105.81 | -9.52 | 0.0004 | 0.029 | 0.94 |
| 1060176 | IR_ON | ENSSSCG00000003660 | 6 | 95386229 | 95386908 | 6.37 | 7.40 | 7.23 | 3.03 | 3.30 | 2.39 | 7.00 | 2.91 | 2.41 | 1.27 | 4.95 | -10.31 | 0.0003 | 0.026 | 1.31 |
| 1060174 | MIR_ON | ENSSSCG00000003660 | 6 | 95386229 | 95386908 | 6.37 | 7.40 | 7.23 | 3.03 | 3.30 | 2.39 | 7.00 | 2.91 | 2.41 | 1.27 | 4.95 | -10.31 | 0.0003 | 0.026 | 1.31 |
| 1060171 | SKIP_OFF | ENSSSCG00000003660 | 6 | 95391911 | 95392005 | 6.37 | 7.40 | 7.23 | 3.03 | 3.30 | 2.39 | 7.00 | 2.91 | 2.41 | 1.27 | 4.95 | -10.31 | 0.0003 | 0.026 | 1.31 |
| 1016507 | TSS | ENSSSCG00000011550 | 13 | 66055124 | 66055295 | 7.84 | 6.04 | 6.53 | 3.45 | 2.73 | 2.42 | 6.80 | 2.87 | 2.37 | 1.25 | 4.84 | -6.76 | 0.0016 | 0.051 | -0.67 |
| 1005180 | SKIP_OFF | ENSSSCG00000024088 | 1 | 109266431 | 109266550 | 5.39 | 4.25 | 4.49 | 2.31 | 1.74 | 2.11 | 4.71 | 2.05 | 2.29 | 1.20 | 3.38 | -7.29 | 0.0012 | 0.045 | -0.32 |
| 1005177 | TSS | ENSSSCG00000024088 | 1 | 109584637 | 109584733 | 5.39 | 4.25 | 4.49 | 2.31 | 1.74 | 2.11 | 4.71 | 2.05 | 2.29 | 1.20 | 3.38 | -7.29 | 0.0012 | 0.045 | -0.32 |
| 1020278 | MSKIP_OFF | ENSSSCG00000034493 | 13 | 160191825 | 160204775 | 1.63 | 1.68 | 1.85 | 0.55 | 0.91 | 0.81 | 1.72 | 0.76 | 2.27 | 1.18 | 1.24 | -7.82 | 0.0009 | 0.041 | 0.01 |
| 1020291 | XMSKIP_OFF | ENSSSCG00000034493 | 13 | 160191825 | 160204381 | 1.63 | 1.68 | 1.85 | 0.55 | 0.91 | 0.81 | 1.72 | 0.76 | 2.27 | 1.18 | 1.24 | -7.82 | 0.0009 | 0.041 | 0.01 |
| 1020298 | XSKIP_OFF | ENSSSCG00000034493 | 13 | 160204236 | 160204381 | 1.63 | 1.68 | 1.85 | 0.55 | 0.91 | 0.81 | 1.72 | 0.76 | 2.27 | 1.18 | 1.24 | -7.82 | 0.0009 | 0.041 | 0.01 |
| 1074531 | TSS | ENSSSCG00000015015 | 9 | 38582399 | 38583148 | 1.67 | 1.97 | 2.05 | 0.78 | 1.03 | 0.74 | 1.90 | 0.85 | 2.23 | 1.16 | 1.37 | -7.19 | 0.0012 | 0.046 | -0.38 |
| 1017953 | TSS | ENSSSCG00000011977 | 13 | 160064051 | 160064647 | 25.94 | 30.48 | 25.28 | 10.04 | 14.97 | 12.12 | 27.23 | 12.38 | 2.20 | 1.14 | 19.81 | -7.27 | 0.0012 | 0.046 | -0.34 |
| 1013963 | AE | ENSSSCG00000030167 | 12 | 18917114 | 18917247 | 24.16 | 24.48 | 23.67 | 9.38 | 9.55 | 14.50 | 24.11 | 11.14 | 2.16 | 1.11 | 17.62 | -8.12 | 0.0007 | 0.038 | 0.19 |
| 1013959 | TSS | ENSSSCG00000030167 | 12 | 18913190 | 18913407 | 24.16 | 24.48 | 23.67 | 9.38 | 9.55 | 14.50 | 24.11 | 11.14 | 2.16 | 1.11 | 17.62 | -8.12 | 0.0007 | 0.038 | 0.19 |
| 1049667 | TTS | ENSSSCG00000006177 | 4 | 62097975 | 62098400 | 3.25 | 2.70 | 3.54 | 1.39 | 1.42 | 1.58 | 3.16 | 1.46 | 2.16 | 1.11 | 2.31 | -7.06 | 0.0013 | 0.048 | -0.47 |
| 1038547 | TSS | ENSSSCG00000013849 | 2 | 61476477 | 61476814 | 20.77 | 20.19 | 22.38 | 8.67 | 8.89 | 11.91 | 21.11 | 9.82 | 2.15 | 1.10 | 15.47 | -9.71 | 0.0003 | 0.028 | 1.03 |
| 1038548 | TTS | ENSSSCG00000013849 | 2 | 61450736 | 61455161 | 20.77 | 20.19 | 22.38 | 8.67 | 8.89 | 11.91 | 21.11 | 9.82 | 2.15 | 1.10 | 15.47 | -9.71 | 0.0003 | 0.028 | 1.03 |
| 1010177 | TTS | ENSSSCG00000022159 | 11 | 18564277 | 18566808 | 42.31 | 45.92 | 44.72 | 17.67 | 24.64 | 19.75 | 44.32 | 20.69 | 2.14 | 1.10 | 32.50 | -10.80 | 0.0002 | 0.024 | 1.52 |
| 1078313 | SKIP_OFF | ENSSSCG00000012126 | X | 10463816 | 10463935 | 33.86 | 36.07 | 36.82 | 14.79 | 21.11 | 15.79 | 35.58 | 17.23 | 2.07 | 1.05 | 26.40 | -9.06 | 0.0005 | 0.032 | 0.70 |
| 1078308 | TSS | ENSSSCG00000012126 | X | 10582201 | 10582431 | 33.98 | 36.49 | 37.33 | 15.14 | 21.27 | 15.79 | 35.93 | 17.40 | 2.07 | 1.05 | 26.66 | -8.99 | 0.0005 | 0.032 | 0.67 |
| 1013962 | TTS | ENSSSCG00000030167 | 12 | 18919292 | 18919407 | 44.79 | 48.30 | 44.40 | 18.51 | 20.97 | 27.82 | 45.83 | 22.43 | 2.04 | 1.03 | 34.13 | -8.15 | 0.0007 | 0.038 | 0.21 |
| 1050981 | TSS | ENSSSCG00000006644 | 4 | 98250968 | 98251593 | 11.98 | 13.16 | 12.17 | 4.36 | 7.12 | 6.93 | 12.44 | 6.14 | 2.03 | 1.02 | 9.29 | -6.96 | 0.0014 | 0.049 | -0.54 |
| 1050986 | XSKIP_OFF | ENSSSCG00000006644 | 4 | 98254271 | 98254459 | 11.98 | 13.16 | 12.17 | 4.36 | 7.12 | 6.93 | 12.44 | 6.14 | 2.03 | 1.02 | 9.29 | -6.96 | 0.0014 | 0.049 | -0.54 |
| 1056491 | TTS | ENSSSCG00000027792 | 5 | 79545430 | 79545440 | 7.35 | 7.44 | 7.11 | 3.47 | 4.64 | 2.70 | 7.30 | 3.60 | 2.03 | 1.02 | 5.45 | -6.84 | 0.0015 | 0.050 | -0.62 |
| 1036268 | TSS | ENSSSCG00000012955 | 2 | 6126815 | 6126932 | 14.14 | 15.61 | 15.69 | 7.93 | 6.10 | 8.64 | 15.15 | 7.56 | 2.00 | 1.00 | 11.35 | -8.86 | 0.0005 | 0.033 | 0.60 |
| 1063581 | TSS | ENSSSCG00000032536 | 6 | 49402430 | 49402661 | 2.36 | 2.07 | 3.23 | 4.94 | 5.11 | 5.30 | 2.55 | 5.12 | 0.50 | -1.00 | 3.83 | 7.42 | 0.0011 | 0.044 | -0.24 |
| 1063582 | TTS | ENSSSCG00000032536 | 6 | 49399123 | 49400510 | 2.36 | 2.07 | 3.23 | 4.94 | 5.11 | 5.30 | 2.55 | 5.12 | 0.50 | -1.00 | 3.83 | 7.42 | 0.0011 | 0.044 | -0.24 |
| 1062300 | TSS | ENSSSCG00000026045 | 6 | 60759168 | 60759606 | 1.68 | 1.45 | 1.96 | 3.28 | 3.72 | 3.23 | 1.70 | 3.41 | 0.50 | -1.01 | 2.55 | 8.40 | 0.0006 | 0.036 | 0.35 |
| 1062301 | TTS | ENSSSCG00000026045 | 6 | 60768214 | 60773371 | 1.68 | 1.45 | 1.96 | 3.28 | 3.72 | 3.23 | 1.70 | 3.41 | 0.50 | -1.01 | 2.55 | 8.40 | 0.0006 | 0.036 | 0.35 |
| 1065387 | TSS | ENSSSCG00000039793 | 6 | 150605432 | 150605505 | 5.30 | 5.77 | 5.28 | 12.36 | 10.15 | 10.46 | 5.45 | 10.99 | 0.50 | -1.01 | 8.22 | 8.30 | 0.0007 | 0.037 | 0.29 |
| 1065394 | SKIP_OFF | ENSSSCG00000039793 | 6 | 150322722 | 150322811 | 5.30 | 5.77 | 5.28 | 12.36 | 10.15 | 10.46 | 5.45 | 10.99 | 0.50 | -1.01 | 8.22 | 8.30 | 0.0007 | 0.037 | 0.29 |
| 1037214 | TSS | ENSSSCG00000013360 | 2 | 40550623 | 40550843 | 7.63 | 7.90 | 10.29 | 17.43 | 19.29 | 15.38 | 8.61 | 17.36 | 0.50 | -1.01 | 12.98 | 6.60 | 0.0018 | 0.053 | -0.79 |
| 1037215 | TTS | ENSSSCG00000013360 | 2 | 40545774 | 40547345 | 7.63 | 7.90 | 10.29 | 17.43 | 19.29 | 15.38 | 8.61 | 17.36 | 0.50 | -1.01 | 12.98 | 6.60 | 0.0018 | 0.053 | -0.79 |

| | | | | | | | | | | | | | | | | | | | | |
|---|---|---|---|---|---|---|---|---|---|---|---|---|---|---|---|---|---|---|---|---|
| 1062338 | TSS | ENSSSCG00000026152 | 6 | 52598136 | 52598375 | 6.60 | 5.63 | 6.06 | 13.95 | 10.58 | 12.41 | 6.10 | 12.31 | 0.50 | -1.01 | 9.21 | 6.51 | 0.0019 | 0.054 | -0.85 |
| 1062339 | TTS | ENSSSCG00000026152 | 6 | 52559614 | 52559826 | 6.60 | 5.63 | 6.06 | 13.95 | 10.58 | 12.41 | 6.10 | 12.31 | 0.50 | -1.01 | 9.21 | 6.51 | 0.0019 | 0.054 | -0.85 |
| 1002950 | TSS | ENSSSCG00000004983 | 1 | 169801523 | 169802376 | 5.47 | 3.76 | 3.64 | 8.43 | 8.17 | 9.41 | 4.29 | 8.67 | 0.49 | -1.02 | 6.48 | 6.64 | 0.0017 | 0.053 | -0.76 |
| 1015161 | TSS | ENSSSCG00000011192 | 13 | 2840435 | 2840955 | 5.82 | 4.44 | 5.45 | 10.49 | 10.51 | 10.85 | 5.24 | 10.62 | 0.49 | -1.02 | 7.93 | 13.28 | 0.0001 | 0.017 | 2.44 |
| 1015162 | TTS | ENSSSCG00000011192 | 13 | 2786676 | 2786825 | 5.82 | 4.44 | 5.45 | 10.49 | 10.51 | 10.85 | 5.24 | 10.62 | 0.49 | -1.02 | 7.93 | 13.28 | 0.0001 | 0.017 | 2.44 |
| 1024532 | TSS | ENSSSCG00000010589 | 14 | 113707659 | 113707899 | 3.05 | 2.04 | 2.89 | 5.66 | 4.76 | 5.77 | 2.66 | 5.40 | 0.49 | -1.02 | 4.03 | 6.47 | 0.0019 | 0.055 | -0.88 |
| 1024533 | TTS | ENSSSCG00000010589 | 14 | 113732447 | 113733826 | 3.05 | 2.04 | 2.89 | 5.66 | 4.76 | 5.77 | 2.66 | 5.40 | 0.49 | -1.02 | 4.03 | 6.47 | 0.0019 | 0.055 | -0.88 |
| 1071761 | TSS | ENSSSCG00000009071 | 8 | 96269180 | 96269321 | 1.70 | 1.73 | 2.36 | 4.13 | 3.84 | 3.75 | 1.93 | 3.91 | 0.49 | -1.02 | 2.92 | 8.57 | 0.0006 | 0.035 | 0.44 |
| 1015163 | MSKIP_ON | ENSSSCG00000011192 | 13 | 2830603 | 2835308 | 5.82 | 4.44 | 5.14 | 10.26 | 10.30 | 10.85 | 5.13 | 10.47 | 0.49 | -1.03 | 7.80 | 12.78 | 0.0001 | 0.019 | 2.28 |
| 1015165 | MSKIP_ON | ENSSSCG00000011192 | 13 | 2833855 | 2836900 | 5.82 | 4.44 | 5.14 | 10.26 | 10.30 | 10.85 | 5.13 | 10.47 | 0.49 | -1.03 | 7.80 | 12.78 | 0.0001 | 0.019 | 2.28 |
| 1013209 | TSS | ENSSSCG00000017995 | 12 | 54584353 | 54585051 | 2.21 | 1.56 | 1.62 | 3.96 | 3.30 | 3.78 | 1.79 | 3.68 | 0.49 | -1.04 | 2.74 | 6.98 | 0.0014 | 0.049 | -0.52 |
| 1013210 | TTS | ENSSSCG00000017995 | 12 | 54640087 | 54642376 | 2.21 | 1.56 | 1.62 | 3.96 | 3.30 | 3.78 | 1.79 | 3.68 | 0.49 | -1.04 | 2.74 | 6.98 | 0.0014 | 0.049 | -0.52 |
| 1017551 | MSKIP_ON | ENSSSCG00000011849 | 13 | 134122648 | 134123063 | 4.04 | 4.15 | 5.96 | 9.45 | 8.95 | 10.71 | 4.72 | 9.70 | 0.49 | -1.04 | 7.21 | 6.51 | 0.0019 | 0.054 | -0.85 |
| 1017553 | SKIP_ON | ENSSSCG00000011849 | 13 | 134122913 | 134123063 | 4.04 | 4.15 | 5.96 | 9.45 | 8.95 | 10.71 | 4.72 | 9.70 | 0.49 | -1.04 | 7.21 | 6.51 | 0.0019 | 0.054 | -0.85 |
| 1076241 | TTS | ENSSSCG00000015645 | 9 | 67113960 | 67114343 | 4.25 | 4.59 | 4.93 | 9.67 | 8.80 | 9.96 | 4.59 | 9.48 | 0.48 | -1.05 | 7.03 | 12.90 | 0.0001 | 0.019 | 2.32 |
| 1065392 | SKIP_OFF | ENSSSCG00000039793 | 6 | 150342486 | 150342578 | 4.10 | 4.34 | 3.04 | 8.88 | 7.41 | 7.44 | 3.83 | 7.91 | 0.48 | -1.05 | 5.87 | 6.89 | 0.0015 | 0.050 | -0.58 |
| 1062303 | IR_OFF | ENSSSCG00000026045 | 6 | 60763804 | 60764065 | 1.36 | 1.35 | 1.68 | 3.14 | 3.22 | 2.72 | 1.46 | 3.03 | 0.48 | -1.05 | 2.25 | 8.50 | 0.0006 | 0.035 | 0.41 |
| 1065388 | TTS | ENSSSCG00000039793 | 6 | 150247296 | 150251485 | 5.01 | 5.08 | 4.47 | 11.55 | 9.29 | 9.40 | 4.85 | 10.08 | 0.48 | -1.05 | 7.47 | 7.30 | 0.0012 | 0.045 | -0.31 |
| 1033458 | SKIP_OFF | ENSSSCG00000028004 | 17 | 28029891 | 28031062 | 1.66 | 1.34 | 1.28 | 3.16 | 2.90 | 2.84 | 1.43 | 2.97 | 0.48 | -1.05 | 2.20 | 10.16 | 0.0003 | 0.026 | 1.24 |
| 1033454 | TSS | ENSSSCG00000028004 | 17 | 27855900 | 27856020 | 1.66 | 1.34 | 1.28 | 3.16 | 2.90 | 2.84 | 1.43 | 2.97 | 0.48 | -1.05 | 2.20 | 10.16 | 0.0003 | 0.026 | 1.24 |
| 1017891 | AE | ENSSSCG00000011949 | 13 | 157384808 | 157384870 | 2.56 | 1.83 | 3.03 | 5.53 | 5.15 | 4.74 | 2.47 | 5.14 | 0.48 | -1.06 | 3.81 | 6.80 | 0.0016 | 0.050 | -0.65 |
| 1044306 | TSS | ENSSSCG00000007958 | 3 | 38813966 | 38815170 | 3.18 | 2.58 | 3.22 | 5.87 | 6.46 | 6.35 | 2.99 | 6.23 | 0.48 | -1.06 | 4.61 | 12.33 | 0.0001 | 0.019 | 2.12 |
| 1044307 | TTS | ENSSSCG00000007958 | 3 | 38805243 | 38808516 | 3.18 | 2.58 | 3.22 | 5.87 | 6.46 | 6.35 | 2.99 | 6.23 | 0.48 | -1.06 | 4.61 | 12.33 | 0.0001 | 0.019 | 2.12 |
| 1075908 | TSS | ENSSSCG00000015509 | 9 | 118113600 | 118114655 | 12.70 | 11.48 | 13.42 | 27.51 | 23.72 | 27.16 | 12.53 | 26.13 | 0.48 | -1.06 | 19.33 | 10.81 | 0.0002 | 0.024 | 1.52 |
| 1075973 | SKIP_ON | ENSSSCG00000015537 | 9 | 122359024 | 122359125 | 3.66 | 3.19 | 2.96 | 7.40 | 6.24 | 6.87 | 3.27 | 6.84 | 0.48 | -1.06 | 5.05 | 9.57 | 0.0004 | 0.029 | 0.96 |
| 1075909 | TTS | ENSSSCG00000015509 | 9 | 117915443 | 117915775 | 12.65 | 11.37 | 13.37 | 27.34 | 23.67 | 27.16 | 12.46 | 26.06 | 0.48 | -1.06 | 19.26 | 10.84 | 0.0002 | 0.024 | 1.54 |
| 1065397 | SKIP_ON | ENSSSCG00000039793 | 6 | 150548748 | 150548891 | 5.10 | 5.47 | 4.64 | 12.19 | 9.74 | 9.93 | 5.07 | 10.62 | 0.48 | -1.07 | 7.85 | 7.15 | 0.0013 | 0.047 | -0.41 |
| 1017546 | SKIP_OFF | ENSSSCG00000011849 | 13 | 134120411 | 134120455 | 6.44 | 5.37 | 7.72 | 13.43 | 12.37 | 15.12 | 6.51 | 13.64 | 0.48 | -1.07 | 10.07 | 7.21 | 0.0012 | 0.046 | -0.37 |
| 1072749 | AE | ENSSSCG00000026655 | 8 | 132130280 | 132130358 | 2.63 | 2.51 | 2.68 | 5.54 | 5.23 | 5.61 | 2.60 | 5.46 | 0.48 | -1.07 | 4.03 | 22.63 | 0.0000 | 0.009 | 4.49 |
| 1072746 | IR_OFF | ENSSSCG00000026655 | 8 | 132144939 | 132144955 | 2.63 | 2.51 | 2.68 | 5.54 | 5.23 | 5.61 | 2.60 | 5.46 | 0.48 | -1.07 | 4.03 | 22.63 | 0.0000 | 0.009 | 4.49 |
| 1072747 | XAE | ENSSSCG00000026655 | 8 | 132074583 | 132074699 | 2.63 | 2.51 | 2.68 | 5.54 | 5.23 | 5.61 | 2.60 | 5.46 | 0.48 | -1.07 | 4.03 | 22.63 | 0.0000 | 0.009 | 4.49 |
| 1072744 | XMSKIP_OFF | ENSSSCG00000026655 | 8 | 132051031 | 132054454 | 2.63 | 2.51 | 2.68 | 5.54 | 5.23 | 5.61 | 2.60 | 5.46 | 0.48 | -1.07 | 4.03 | 22.63 | 0.0000 | 0.009 | 4.49 |
| 1079410 | TSS | ENSSSCG00000012619 | X | 98095128 | 98095236 | 1.23 | 1.31 | 1.29 | 2.75 | 2.54 | 2.79 | 1.28 | 2.69 | 0.48 | -1.07 | 1.99 | 16.63 | 0.0000 | 0.012 | 3.38 |
| 1073767 | SKIP_OFF | ENSSSCG00000014598 | 9 | 2008898 | 2008930 | 6.80 | 7.94 | 6.40 | 15.93 | 14.81 | 13.88 | 7.05 | 14.87 | 0.47 | -1.08 | 10.96 | 11.06 | 0.0002 | 0.023 | 1.63 |
| 1017544 | TTS | ENSSSCG00000011849 | 13 | 134124013 | 134124823 | 7.83 | 6.08 | 8.15 | 16.00 | 14.10 | 16.46 | 7.35 | 15.52 | 0.47 | -1.08 | 11.44 | 8.95 | 0.0005 | 0.032 | 0.65 |
| 1019259 | SKIP_ON | ENSSSCG00000027571 | 13 | 165841504 | 165841572 | 4.46 | 4.64 | 5.20 | 11.57 | 9.41 | 9.29 | 4.77 | 10.09 | 0.47 | -1.08 | 7.43 | 7.28 | 0.0012 | 0.045 | -0.33 |
| 1037489 | AE | ENSSSCG00000013427 | 2 | 77193065 | 77193206 | 7.89 | 7.63 | 9.98 | 18.85 | 18.94 | 16.26 | 8.50 | 18.01 | 0.47 | -1.08 | 13.26 | 8.77 | 0.0005 | 0.033 | 0.55 |
| 1037490 | XAE | ENSSSCG00000013427 | 2 | 77193065 | 77193206 | 7.89 | 7.63 | 9.98 | 18.85 | 18.94 | 16.26 | 8.50 | 18.01 | 0.47 | -1.08 | 13.26 | 8.77 | 0.0005 | 0.033 | 0.55 |
| 1013923 | TTS | ENSSSCG00000029553 | 12 | 917062 | 917264 | 1.47 | 1.32 | 1.87 | 3.50 | 3.05 | 3.33 | 1.55 | 3.29 | 0.47 | -1.08 | 2.42 | 8.60 | 0.0006 | 0.035 | 0.46 |
| 1032383 | SKIP_ON | ENSSSCG00000007247 | 17 | 35929730 | 35929802 | 3.31 | 3.03 | 2.41 | 6.54 | 6.02 | 6.08 | 2.92 | 6.21 | 0.47 | -1.09 | 4.56 | 11.13 | 0.0002 | 0.023 | 1.66 |
| 1047244 | AE | ENSSSCG00000029185 | 3 | 112087391 | 112087592 | 7.54 | 4.68 | 6.95 | 13.21 | 12.92 | 14.76 | 6.39 | 13.63 | 0.47 | -1.09 | 10.01 | 7.38 | 0.0011 | 0.045 | -0.26 |
| 1047241 | TSS | ENSSSCG00000029185 | 3 | 112089263 | 112089586 | 7.54 | 4.68 | 6.95 | 13.21 | 12.92 | 14.76 | 6.39 | 13.63 | 0.47 | -1.09 | 10.01 | 7.38 | 0.0011 | 0.045 | -0.26 |
| 1012554 | XMSKIP_OFF | ENSSSCG00000017754 | 12 | 44165136 | 44166843 | 1.14 | 1.33 | 1.80 | 3.28 | 3.08 | 2.78 | 1.43 | 3.04 | 0.47 | -1.09 | 2.24 | 6.91 | 0.0015 | 0.049 | -0.57 |
| 1012558 | XSKIP_OFF | ENSSSCG00000017754 | 12 | 44166808 | 44166843 | 1.14 | 1.33 | 1.80 | 3.28 | 3.08 | 2.78 | 1.43 | 3.04 | 0.47 | -1.09 | 2.24 | 6.91 | 0.0015 | 0.049 | -0.57 |
| 1060060 | TSS | ENSSSCG00000003619 | 6 | 89346932 | 89347036 | 1.16 | 1.30 | 1.47 | 2.62 | 2.72 | 3.07 | 1.31 | 2.80 | 0.47 | -1.10 | 2.06 | 9.36 | 0.0004 | 0.030 | 0.86 |
| 1060061 | TTS | ENSSSCG00000003619 | 6 | 89384841 | 89384979 | 1.16 | 1.30 | 1.47 | 2.62 | 2.72 | 3.07 | 1.31 | 2.80 | 0.47 | -1.10 | 2.06 | 9.36 | 0.0004 | 0.030 | 0.86 |
| 1057139 | AE | ENSSSCG00000035733 | 5 | 414555 | 414679 | 13.28 | 10.68 | 9.12 | 22.71 | 22.85 | 25.29 | 11.03 | 23.62 | 0.47 | -1.10 | 17.32 | 9.07 | 0.0005 | 0.032 | 0.71 |
| 1057132 | TSS | ENSSSCG00000035733 | 5 | 401269 | 401381 | 13.28 | 10.68 | 9.12 | 22.71 | 22.85 | 25.29 | 11.03 | 23.62 | 0.47 | -1.10 | 17.32 | 9.07 | 0.0005 | 0.032 | 0.71 |
| 1051165 | TSS | ENSSSCG00000006733 | 4 | 103676385 | 103676441 | 2.84 | 2.78 | 2.33 | 6.24 | 6.11 | 4.82 | 2.65 | 5.72 | 0.46 | -1.11 | 4.18 | 6.75 | 0.0016 | 0.051 | -0.68 |
| 1000061 | IR_OFF | ENSSSCG00000004023 | 1 | 2098700 | 2101548 | 6.10 | 4.77 | 4.48 | 11.79 | 9.79 | 11.73 | 5.12 | 11.10 | 0.46 | -1.12 | 8.11 | 7.69 | 0.0009 | 0.042 | -0.07 |
| 1000059 | MIR_OFF | ENSSSCG00000004023 | 1 | 2097678 | 2101548 | 6.10 | 4.77 | 4.48 | 11.79 | 9.79 | 11.73 | 5.12 | 11.10 | 0.46 | -1.12 | 8.11 | 7.69 | 0.0009 | 0.042 | -0.07 |
| 1013922 | TSS | ENSSSCG00000029553 | 12 | 837752 | 837887 | 1.37 | 1.32 | 1.75 | 3.45 | 2.92 | 3.29 | 1.48 | 3.22 | 0.46 | -1.12 | 2.35 | 8.78 | 0.0005 | 0.033 | 0.56 |
| 1013931 | XAE | ENSSSCG00000029553 | 12 | 864924 | 865100 | 1.37 | 1.32 | 1.75 | 3.45 | 2.92 | 3.29 | 1.48 | 3.22 | 0.46 | -1.12 | 2.35 | 8.78 | 0.0005 | 0.033 | 0.56 |
| 1013924 | XSKIP_ON | ENSSSCG00000029553 | 12 | 890862 | 890958 | 1.37 | 1.32 | 1.75 | 3.45 | 2.92 | 3.29 | 1.48 | 3.22 | 0.46 | -1.12 | 2.35 | 8.78 | 0.0005 | 0.033 | 0.56 |

| | | | | | | | | | | | | | | | | | | | | |
|---|---|---|---|---|---|---|---|---|---|---|---|---|---|---|---|---|---|---|---|---|
| 1038771 | SKIP_ON | ENSSSCG00000013895 | 2 | 59681239 | 59681259 | 1.32 | 1.26 | 1.49 | 3.31 | 2.75 | 2.80 | 1.36 | 2.95 | 0.46 | -1.12 | 2.15 | 8.62 | 0.0006 | 0.035 | 0.47 |
| 1013921 | TSS | ENSSSCG00000029553 | 12 | 833115 | 833527 | 1.09 | 1.00 | 1.21 | 2.09 | 2.39 | 2.71 | 1.10 | 2.39 | 0.46 | -1.12 | 1.75 | 7.07 | 0.0013 | 0.047 | -0.46 |
| 1055810 | SKIP_OFF | ENSSSCG00000000939 | 5 | 100357554 | 100357634 | 2.95 | 1.74 | 1.83 | 4.67 | 4.95 | 4.58 | 2.17 | 4.74 | 0.46 | -1.13 | 3.45 | 6.72 | 0.0016 | 0.052 | -0.71 |
| 1055813 | SKIP_ON | ENSSSCG00000000939 | 5 | 100357377 | 100357472 | 2.95 | 1.74 | 1.83 | 4.67 | 4.95 | 4.58 | 2.17 | 4.74 | 0.46 | -1.13 | 3.45 | 6.72 | 0.0016 | 0.052 | -0.71 |
| 1021979 | TSS | ENSSSCG00000009822 | 14 | 31849687 | 31850332 | 1.09 | 0.98 | 1.24 | 2.35 | 2.70 | 2.21 | 1.11 | 2.42 | 0.46 | -1.13 | 1.76 | 8.26 | 0.0007 | 0.037 | 0.27 |
| 1021980 | TTS | ENSSSCG00000009822 | 14 | 31859213 | 31862651 | 1.09 | 0.98 | 1.24 | 2.35 | 2.70 | 2.21 | 1.11 | 2.42 | 0.46 | -1.13 | 1.76 | 8.26 | 0.0007 | 0.037 | 0.27 |
| 1017887 | TTS | ENSSSCG00000011949 | 13 | 157356251 | 157361581 | 2.56 | 1.84 | 3.03 | 5.89 | 5.45 | 4.94 | 2.48 | 5.43 | 0.46 | -1.13 | 3.95 | 7.06 | 0.0013 | 0.048 | -0.47 |
| 1063595 | AE | ENSSSCG00000032585 | 6 | 58187493 | 58187630 | 2.07 | 2.37 | 2.88 | 5.88 | 5.63 | 4.54 | 2.44 | 5.35 | 0.46 | -1.13 | 3.89 | 6.50 | 0.0019 | 0.054 | -0.86 |
| 1063592 | TSS | ENSSSCG00000032585 | 6 | 58188909 | 58189093 | 2.07 | 2.37 | 2.88 | 5.88 | 5.63 | 4.54 | 2.44 | 5.35 | 0.46 | -1.13 | 3.89 | 6.50 | 0.0019 | 0.054 | -0.86 |
| 1022693 | TTS | ENSSSCG00000010031 | 14 | 48303322 | 48303506 | 2.74 | 2.21 | 3.03 | 6.70 | 5.20 | 5.63 | 2.66 | 5.84 | 0.45 | -1.14 | 4.25 | 6.62 | 0.0017 | 0.053 | -0.77 |
| 1038769 | SKIP_ON | ENSSSCG00000013895 | 2 | 59679199 | 59679274 | 1.40 | 1.56 | 2.05 | 4.18 | 3.38 | 3.49 | 1.67 | 3.68 | 0.45 | -1.14 | 2.68 | 6.66 | 0.0017 | 0.053 | -0.74 |
| 1075912 | SKIP_OFF | ENSSSCG00000015509 | 9 | 118058399 | 118058467 | 6.96 | 4.43 | 6.27 | 14.16 | 11.28 | 13.48 | 5.89 | 12.97 | 0.45 | -1.14 | 9.43 | 6.53 | 0.0018 | 0.054 | -0.84 |
| 1016834 | TTS | ENSSSCG00000011670 | 13 | 82042731 | 82045106 | 10.12 | 9.90 | 14.20 | 22.16 | 27.81 | 25.62 | 11.41 | 25.20 | 0.45 | -1.14 | 18.30 | 6.79 | 0.0016 | 0.051 | -0.65 |
| 1063252 | TSS | ENSSSCG00000031102 | 6 | 65333764 | 65333894 | 3.33 | 2.75 | 3.61 | 6.77 | 6.87 | 7.79 | 3.23 | 7.14 | 0.45 | -1.14 | 5.19 | 10.06 | 0.0003 | 0.027 | 1.19 |
| 1063253 | TTS | ENSSSCG00000031102 | 6 | 65323756 | 65327576 | 3.33 | 2.75 | 3.61 | 6.77 | 6.87 | 7.79 | 3.23 | 7.14 | 0.45 | -1.14 | 5.19 | 10.06 | 0.0003 | 0.027 | 1.19 |
| 1013926 | XAE | ENSSSCG00000029553 | 12 | 851814 | 851945 | 2.36 | 2.32 | 2.83 | 5.48 | 5.18 | 5.95 | 2.50 | 5.54 | 0.45 | -1.15 | 4.02 | 11.49 | 0.0002 | 0.022 | 1.80 |
| 1013929 | XAE | ENSSSCG00000029553 | 12 | 856994 | 857213 | 2.36 | 2.32 | 2.83 | 5.48 | 5.18 | 5.95 | 2.50 | 5.54 | 0.45 | -1.15 | 4.02 | 11.49 | 0.0002 | 0.022 | 1.80 |
| 1021823 | SKIP_ON | ENSSSCG00000009783 | 14 | 29894182 | 29894238 | 5.81 | 4.02 | 6.22 | 11.91 | 10.89 | 12.71 | 5.35 | 11.84 | 0.45 | -1.15 | 8.60 | 8.05 | 0.0008 | 0.039 | 0.15 |
| 1015649 | SKIP_ON | ENSSSCG00000011330 | 13 | 29857575 | 29857724 | 0.51 | 0.60 | 0.60 | 1.20 | 1.28 | 1.30 | 0.57 | 1.26 | 0.45 | -1.15 | 0.91 | 12.47 | 0.0001 | 0.019 | 2.17 |
| 1015643 | TSS | ENSSSCG00000011330 | 13 | 29855380 | 29855461 | 0.51 | 0.60 | 0.60 | 1.20 | 1.28 | 1.30 | 0.57 | 1.26 | 0.45 | -1.15 | 0.91 | 12.47 | 0.0001 | 0.019 | 2.17 |
| 1038765 | TSS | ENSSSCG00000013895 | 2 | 59684201 | 59684511 | 1.26 | 1.09 | 1.49 | 3.19 | 2.75 | 2.56 | 1.28 | 2.83 | 0.45 | -1.15 | 2.06 | 7.36 | 0.0011 | 0.045 | -0.28 |
| 1038778 | XMIR_OFF | ENSSSCG00000013895 | 2 | 59680706 | 59681442 | 1.26 | 1.09 | 1.49 | 3.19 | 2.75 | 2.56 | 1.28 | 2.83 | 0.45 | -1.15 | 2.06 | 7.36 | 0.0011 | 0.045 | -0.28 |
| 1009882 | TSS | ENSSSCG00000009501 | 11 | 65201457 | 65201471 | 10.16 | 8.87 | 10.31 | 24.06 | 19.13 | 22.15 | 9.78 | 21.78 | 0.45 | -1.15 | 15.78 | 8.47 | 0.0006 | 0.035 | 0.39 |
| 1009885 | XSKIP_OFF | ENSSSCG00000009501 | 11 | 65155199 | 65155225 | 10.16 | 8.87 | 10.31 | 24.06 | 19.13 | 22.15 | 9.78 | 21.78 | 0.45 | -1.15 | 15.78 | 8.47 | 0.0006 | 0.035 | 0.39 |
| 1077402 | SKIP_ON | ENSSSCG00000032721 | 9 | 133192899 | 133192958 | 1.59 | 1.72 | 1.79 | 4.04 | 3.21 | 4.12 | 1.70 | 3.79 | 0.45 | -1.16 | 2.74 | 7.39 | 0.0011 | 0.045 | -0.26 |
| 1021820 | TSS | ENSSSCG00000009783 | 14 | 29894683 | 29895548 | 5.81 | 4.02 | 6.22 | 12.08 | 11.06 | 12.71 | 5.35 | 11.95 | 0.45 | -1.16 | 8.65 | 8.45 | 0.0006 | 0.036 | 0.38 |
| 1074123 | SKIP_ON | ENSSSCG00000014880 | 9 | 12071027 | 12071143 | 0.62 | 1.02 | 0.88 | 1.86 | 1.78 | 2.00 | 0.84 | 1.88 | 0.45 | -1.16 | 1.36 | 8.00 | 0.0008 | 0.039 | 0.12 |
| 1003157 | TTS | ENSSSCG00000005055 | 1 | 184495850 | 184496155 | 110.23 | 115.17 | 100.51 | 249.15 | 213.19 | 268.93 | 108.63 | 243.76 | 0.45 | -1.17 | 176.20 | 8.51 | 0.0006 | 0.035 | 0.41 |
| 1003155 | TSS | ENSSSCG00000005055 | 1 | 184478643 | 184478771 | 102.18 | 115.07 | 96.89 | 238.87 | 205.65 | 263.04 | 104.72 | 235.85 | 0.44 | -1.17 | 170.28 | 7.97 | 0.0008 | 0.039 | 0.10 |
| 1013903 | IR_ON | ENSSSCG00000029102 | 12 | 22308513 | 22309044 | 1.06 | 0.97 | 1.43 | 2.45 | 2.74 | 2.61 | 1.15 | 2.60 | 0.44 | -1.17 | 1.88 | 9.05 | 0.0005 | 0.032 | 0.70 |
| 1013902 | SKIP_OFF | ENSSSCG00000029102 | 12 | 22318990 | 22319046 | 1.06 | 0.97 | 1.43 | 2.45 | 2.74 | 2.61 | 1.15 | 2.60 | 0.44 | -1.17 | 1.88 | 9.05 | 0.0005 | 0.032 | 0.70 |
| 1047242 | TTS | ENSSSCG00000029185 | 3 | 112083268 | 112085409 | 9.17 | 6.53 | 9.05 | 18.12 | 16.98 | 20.99 | 8.25 | 18.70 | 0.44 | -1.18 | 13.47 | 7.55 | 0.0010 | 0.043 | -0.15 |
| 1079419 | XAE | ENSSSCG00000012619 | X | 98104754 | 98104856 | 2.06 | 2.50 | 2.92 | 5.75 | 5.50 | 5.71 | 2.49 | 5.65 | 0.44 | -1.18 | 4.07 | 12.76 | 0.0001 | 0.019 | 2.27 |
| 1037865 | IR_ON | ENSSSCG00000013572 | 2 | 71577682 | 71577768 | 1.33 | 1.39 | 0.83 | 2.97 | 2.66 | 2.43 | 1.18 | 2.69 | 0.44 | -1.18 | 1.94 | 6.70 | 0.0017 | 0.052 | -0.72 |
| 1037862 | SKIP_OFF | ENSSSCG00000013572 | 2 | 71601762 | 71601874 | 1.33 | 1.39 | 0.83 | 2.97 | 2.66 | 2.43 | 1.18 | 2.69 | 0.44 | -1.18 | 1.94 | 6.70 | 0.0017 | 0.052 | -0.72 |
| 1017547 | SKIP_ON | ENSSSCG00000011849 | 13 | 134122648 | 134122743 | 6.59 | 5.30 | 7.58 | 14.77 | 13.60 | 15.83 | 6.49 | 14.73 | 0.44 | -1.18 | 10.61 | 9.49 | 0.0004 | 0.029 | 0.92 |
| 1002142 | SKIP_ON | ENSSSCG00000004704 | 1 | 127869779 | 127869923 | 0.83 | 1.23 | 1.49 | 2.45 | 2.93 | 2.68 | 1.18 | 2.69 | 0.44 | -1.18 | 1.93 | 6.65 | 0.0017 | 0.053 | -0.76 |
| 1065839 | TTS | ENSSSCG00000001011 | 7 | 1681355 | 1685563 | 12.73 | 16.36 | 11.79 | 32.59 | 29.43 | 31.14 | 13.63 | 31.05 | 0.44 | -1.19 | 22.34 | 11.11 | 0.0002 | 0.023 | 1.65 |
| 1065838 | TSS | ENSSSCG00000001011 | 7 | 1693212 | 1693281 | 12.70 | 16.35 | 11.78 | 32.53 | 29.39 | 31.14 | 13.61 | 31.02 | 0.44 | -1.19 | 22.32 | 11.11 | 0.0002 | 0.023 | 1.65 |
| 1012159 | TTS | ENSSSCG00000017601 | 12 | 31272730 | 31272764 | 0.81 | 0.59 | 0.75 | 1.43 | 1.66 | 1.84 | 0.72 | 1.64 | 0.44 | -1.19 | 1.18 | 7.00 | 0.0014 | 0.048 | -0.51 |
| 1076398 | TSS | ENSSSCG00000021702 | 9 | 9280491 | 9280627 | 1.01 | 0.91 | 0.86 | 1.88 | 2.46 | 2.02 | 0.93 | 2.12 | 0.44 | -1.20 | 1.52 | 6.85 | 0.0015 | 0.050 | -0.62 |
| 1076399 | TTS | ENSSSCG00000021702 | 9 | 9176032 | 9179459 | 1.01 | 0.91 | 0.86 | 1.88 | 2.46 | 2.02 | 0.93 | 2.12 | 0.44 | -1.20 | 1.52 | 6.85 | 0.0015 | 0.050 | -0.62 |
| 1066035 | IR_OFF | ENSSSCG00000001095 | 7 | 19588399 | 19591335 | 6.63 | 7.09 | 5.22 | 12.16 | 15.78 | 15.67 | 6.31 | 14.53 | 0.43 | -1.20 | 10.42 | 6.64 | 0.0017 | 0.053 | -0.76 |
| 1017564 | AE | ENSSSCG00000011849 | 13 | 134108471 | 134108692 | 4.60 | 3.79 | 5.17 | 11.54 | 9.53 | 10.20 | 4.52 | 10.42 | 0.43 | -1.21 | 7.47 | 8.80 | 0.0005 | 0.033 | 0.57 |
| 1069238 | TTS | ENSSSCG00000025768 | 7 | 112167899 | 112167927 | 132.41 | 115.25 | 176.24 | 347.68 | 286.09 | 344.34 | 141.30 | 326.04 | 0.43 | -1.21 | 233.67 | 7.27 | 0.0012 | 0.046 | -0.33 |
| 1027502 | TSS | ENSSSCG00000015931 | 15 | 75852198 | 75852290 | 2.62 | 2.22 | 3.29 | 6.81 | 6.46 | 5.52 | 2.71 | 6.26 | 0.43 | -1.21 | 4.49 | 7.56 | 0.0010 | 0.043 | -0.15 |
| 1027503 | TTS | ENSSSCG00000015931 | 15 | 75872968 | 75874947 | 2.62 | 2.22 | 3.29 | 6.81 | 6.46 | 5.52 | 2.71 | 6.26 | 0.43 | -1.21 | 4.49 | 7.56 | 0.0010 | 0.043 | -0.15 |
| 1068303 | SKIP_OFF | ENSSSCG00000002332 | 7 | 94942115 | 94942361 | 1.83 | 1.52 | 1.31 | 3.91 | 3.16 | 3.70 | 1.55 | 3.59 | 0.43 | -1.21 | 2.57 | 7.93 | 0.0008 | 0.040 | 0.08 |
| 1068310 | SKIP_ON | ENSSSCG00000002332 | 7 | 94879654 | 94879720 | 1.83 | 1.52 | 1.31 | 3.91 | 3.16 | 3.70 | 1.55 | 3.59 | 0.43 | -1.21 | 2.57 | 7.93 | 0.0008 | 0.040 | 0.08 |
| 1068308 | XSKIP_ON | ENSSSCG00000002332 | 7 | 94829992 | 94830087 | 1.83 | 1.52 | 1.31 | 3.91 | 3.16 | 3.70 | 1.55 | 3.59 | 0.43 | -1.21 | 2.57 | 7.93 | 0.0008 | 0.040 | 0.08 |
| 1017555 | SKIP_ON | ENSSSCG00000011849 | 13 | 134106510 | 134106580 | 4.60 | 3.70 | 5.17 | 11.54 | 9.44 | 10.20 | 4.49 | 10.39 | 0.43 | -1.21 | 7.44 | 8.38 | 0.0006 | 0.036 | 0.34 |
| 1045306 | TTS | ENSSSCG00000008336 | 3 | 72620712 | 72623370 | 16.85 | 23.47 | 20.13 | 50.44 | 39.63 | 50.40 | 20.15 | 46.82 | 0.43 | -1.22 | 33.49 | 6.96 | 0.0014 | 0.049 | -0.54 |
| 1045305 | TSS | ENSSSCG00000008336 | 3 | 72694121 | 72694173 | 16.85 | 23.47 | 20.13 | 50.44 | 39.63 | 50.40 | 20.15 | 46.82 | 0.43 | -1.22 | 33.49 | 6.96 | 0.0014 | 0.049 | -0.54 |
| 1070914 | TSS | ENSSSCG00000008761 | 8 | 20254332 | 20255039 | 2.93 | 2.14 | 3.17 | 6.10 | 6.79 | 6.29 | 2.75 | 6.39 | 0.43 | -1.22 | 4.57 | 10.32 | 0.0003 | 0.026 | 1.31 |

| | | | | | | | | | | | | | | | | | | | | |
|---|---|---|---|---|---|---|---|---|---|---|---|---|---|---|---|---|---|---|---|---|
| 1070915 | TTS | ENSSSCG00000008761 | 8 | 20368774 | 20368848 | 2.93 | 2.14 | 3.17 | 6.10 | 6.79 | 6.29 | 2.75 | 6.39 | 0.43 | -1.22 | 4.57 | 10.32 | 0.0003 | 0.026 | 1.31 |
| 1056727 | TTS | ENSSSCG00000031441 | 5 | 2873428 | 2875485 | 4.73 | 3.17 | 3.38 | 8.67 | 8.13 | 9.49 | 3.76 | 8.76 | 0.43 | -1.22 | 6.26 | 8.44 | 0.0006 | 0.036 | 0.37 |
| 1044426 | TSS | ENSSSCG00000008004 | 3 | 41060908 | 41062032 | 4.91 | 4.01 | 4.92 | 9.89 | 11.28 | 11.14 | 4.61 | 10.77 | 0.43 | -1.22 | 7.69 | 12.17 | 0.0001 | 0.019 | 2.06 |
| 1044427 | TTS | ENSSSCG00000008004 | 3 | 41053268 | 41057978 | 4.91 | 4.01 | 4.92 | 9.89 | 11.28 | 11.14 | 4.61 | 10.77 | 0.43 | -1.22 | 7.69 | 12.17 | 0.0001 | 0.019 | 2.06 |
| 1008516 | SKIP_OFF | ENSSSCG00000024127 | 10 | 49056944 | 49057030 | 4.77 | 3.88 | 5.23 | 10.95 | 9.89 | 11.74 | 4.63 | 10.86 | 0.43 | -1.23 | 7.74 | 9.90 | 0.0003 | 0.027 | 1.12 |
| 1008517 | SKIP_ON | ENSSSCG00000024127 | 10 | 49053729 | 49053905 | 4.77 | 3.88 | 5.23 | 10.95 | 9.89 | 11.74 | 4.63 | 10.86 | 0.43 | -1.23 | 7.74 | 9.90 | 0.0003 | 0.027 | 1.12 |
| 1002956 | SKIP_ON | ENSSSCG00000004983 | 1 | 169843986 | 169844114 | 5.57 | 3.82 | 3.64 | 10.74 | 9.25 | 10.79 | 4.34 | 10.26 | 0.42 | -1.24 | 7.30 | 7.90 | 0.0008 | 0.040 | 0.06 |
| 1024461 | MSKIP_OFF | ENSSSCG00000010563 | 14 | 112495835 | 112510827 | 2.53 | 2.28 | 1.58 | 5.05 | 5.11 | 4.95 | 2.13 | 5.04 | 0.42 | -1.24 | 3.58 | 10.54 | 0.0002 | 0.025 | 1.41 |
| 1024455 | SKIP_OFF | ENSSSCG00000010563 | 14 | 112495835 | 112495912 | 2.53 | 2.28 | 1.58 | 5.05 | 5.11 | 4.95 | 2.13 | 5.04 | 0.42 | -1.24 | 3.58 | 10.54 | 0.0002 | 0.025 | 1.41 |
| 1043988 | TSS | ENSSSCG00000007839 | 3 | 23979880 | 23980230 | 6.59 | 5.76 | 7.82 | 15.62 | 14.71 | 17.37 | 6.72 | 15.90 | 0.42 | -1.24 | 11.31 | 9.90 | 0.0003 | 0.027 | 1.12 |
| 1039468 | SKIP_OFF | ENSSSCG00000014170 | 2 | 103324699 | 103324764 | 12.24 | 11.19 | 12.24 | 28.59 | 26.64 | 29.31 | 11.89 | 28.18 | 0.42 | -1.24 | 20.04 | 19.86 | 0.0000 | 0.009 | 4.05 |
| 1005482 | MSKIP_ON | ENSSSCG00000027765 | 1 | 175234854 | 175244312 | 3.20 | 3.79 | 3.95 | 9.58 | 7.94 | 8.42 | 3.65 | 8.65 | 0.42 | -1.25 | 6.15 | 9.86 | 0.0003 | 0.028 | 1.10 |
| 1005474 | SKIP_ON | ENSSSCG00000027765 | 1 | 175234854 | 175235006 | 3.20 | 3.79 | 3.95 | 9.58 | 7.94 | 8.42 | 3.65 | 8.65 | 0.42 | -1.25 | 6.15 | 9.86 | 0.0003 | 0.028 | 1.10 |
| 1005478 | SKIP_ON | ENSSSCG00000027765 | 1 | 175244214 | 175244312 | 3.20 | 3.79 | 3.95 | 9.58 | 7.94 | 8.42 | 3.65 | 8.65 | 0.42 | -1.25 | 6.15 | 9.86 | 0.0003 | 0.028 | 1.10 |
| 1056726 | TSS | ENSSSCG00000031441 | 5 | 2947615 | 2947785 | 4.53 | 3.17 | 3.38 | 8.67 | 8.13 | 9.49 | 3.69 | 8.76 | 0.42 | -1.25 | 6.23 | 9.30 | 0.0004 | 0.030 | 0.83 |
| 1043990 | TTS | ENSSSCG00000007839 | 3 | 23887124 | 23891252 | 6.80 | 6.74 | 8.03 | 16.49 | 16.05 | 18.95 | 7.19 | 17.16 | 0.42 | -1.26 | 12.18 | 10.64 | 0.0002 | 0.025 | 1.45 |
| 1061958 | TSS | ENSSSCG00000024476 | 6 | 27627759 | 27627840 | 7.17 | 4.30 | 5.82 | 13.27 | 12.54 | 15.45 | 5.76 | 13.75 | 0.42 | -1.26 | 9.76 | 7.05 | 0.0013 | 0.048 | -0.48 |
| 1061959 | TTS | ENSSSCG00000024476 | 6 | 27637313 | 27637508 | 7.17 | 4.30 | 5.82 | 13.27 | 12.54 | 15.45 | 5.76 | 13.75 | 0.42 | -1.26 | 9.76 | 7.05 | 0.0013 | 0.048 | -0.48 |
| 1053150 | TSS | ENSSSCG00000038558 | 4 | 98470392 | 98470664 | 20.01 | 15.75 | 22.66 | 41.58 | 45.66 | 52.45 | 19.47 | 46.56 | 0.42 | -1.26 | 33.02 | 7.66 | 0.0009 | 0.042 | -0.08 |
| 1003727 | SKIP_ON | ENSSSCG00000005272 | 1 | 228123311 | 228123382 | 1.96 | 2.23 | 2.60 | 5.53 | 5.27 | 5.45 | 2.26 | 5.42 | 0.42 | -1.26 | 3.84 | 16.22 | 0.0000 | 0.013 | 3.28 |
| 1043422 | AE | ENSSSCG00000007568 | 3 | 1884375 | 1885672 | 1.06 | 0.66 | 0.98 | 2.22 | 2.08 | 2.21 | 0.90 | 2.17 | 0.41 | -1.27 | 1.54 | 10.01 | 0.0003 | 0.027 | 1.17 |
| 1004414 | TTS | ENSSSCG00000005582 | 1 | 264045670 | 264046096 | 1.48 | 1.78 | 1.94 | 4.23 | 3.98 | 4.34 | 1.73 | 4.19 | 0.41 | -1.27 | 2.96 | 14.72 | 0.0001 | 0.015 | 2.88 |
| 1043413 | SKIP_ON | ENSSSCG00000007568 | 3 | 1893261 | 1893395 | 1.13 | 0.80 | 1.13 | 2.51 | 2.26 | 2.68 | 1.02 | 2.48 | 0.41 | -1.28 | 1.75 | 9.10 | 0.0005 | 0.032 | 0.72 |
| 1003724 | TSS | ENSSSCG00000005272 | 1 | 228141654 | 228142091 | 1.33 | 1.71 | 1.68 | 4.03 | 3.34 | 4.15 | 1.57 | 3.84 | 0.41 | -1.29 | 2.71 | 8.54 | 0.0006 | 0.035 | 0.43 |
| 1041230 | TTS | ENSSSCG00000028144 | 2 | 62352434 | 62353106 | 1.91 | 1.61 | 2.59 | 5.18 | 4.57 | 5.18 | 2.04 | 4.98 | 0.41 | -1.29 | 3.51 | 8.80 | 0.0005 | 0.033 | 0.57 |
| 1006683 | TSS | ENSSSCG00000037508 | 1 | 261314310 | 261315093 | 1.85 | 2.31 | 2.05 | 5.06 | 5.10 | 5.03 | 2.07 | 5.07 | 0.41 | -1.29 | 3.57 | 22.55 | 0.0000 | 0.009 | 4.48 |
| 1074872 | TTS | ENSSSCG00000015120 | 9 | 46563482 | 46565228 | 5.46 | 2.65 | 4.13 | 10.71 | 9.69 | 9.62 | 4.08 | 10.01 | 0.41 | -1.29 | 7.04 | 7.11 | 0.0013 | 0.047 | -0.44 |
| 1053153 | SKIP_ON | ENSSSCG00000038558 | 4 | 98500781 | 98500912 | 17.35 | 14.26 | 19.19 | 36.04 | 40.70 | 47.85 | 16.93 | 41.53 | 0.41 | -1.29 | 29.23 | 7.02 | 0.0014 | 0.048 | -0.50 |
| 1004111 | TSS | ENSSSCG00000005425 | 1 | 246548367 | 246548794 | 20.67 | 19.37 | 22.65 | 50.18 | 48.37 | 55.47 | 20.90 | 51.34 | 0.41 | -1.30 | 36.12 | 13.85 | 0.0001 | 0.017 | 2.62 |
| 1016206 | SKIP_ON | ENSSSCG00000011482 | 13 | 40582144 | 40582518 | 1.10 | 1.20 | 1.29 | 2.97 | 2.49 | 3.35 | 1.19 | 2.94 | 0.41 | -1.30 | 2.07 | 7.20 | 0.0012 | 0.046 | -0.38 |
| 1061792 | TSS | ENSSSCG00000023276 | 6 | 119192757 | 119193356 | 2.61 | 1.36 | 1.99 | 4.90 | 4.55 | 5.23 | 1.99 | 4.89 | 0.41 | -1.30 | 3.44 | 7.49 | 0.0010 | 0.043 | -0.19 |
| 1002583 | TTS | ENSSSCG00000004856 | 1 | 145914838 | 145914887 | 2.35 | 2.39 | 2.12 | 5.97 | 6.16 | 4.88 | 2.29 | 5.67 | 0.40 | -1.31 | 3.98 | 8.78 | 0.0005 | 0.033 | 0.56 |
| 1004114 | IR_ON | ENSSSCG00000005425 | 1 | 246706001 | 246706092 | 20.54 | 19.10 | 21.87 | 49.43 | 47.84 | 55.27 | 20.50 | 50.85 | 0.40 | -1.31 | 35.68 | 13.45 | 0.0001 | 0.017 | 2.50 |
| 1004113 | TTS | ENSSSCG00000005425 | 1 | 246712492 | 246714861 | 20.54 | 19.00 | 21.87 | 49.43 | 47.84 | 55.27 | 20.47 | 50.85 | 0.40 | -1.31 | 35.66 | 13.41 | 0.0001 | 0.017 | 2.49 |
| 1043201 | TSS | ENSSSCG00000040416 | 2 | 5906314 | 5906388 | 6.60 | 5.61 | 7.62 | 17.85 | 14.07 | 17.34 | 6.61 | 16.42 | 0.40 | -1.31 | 11.51 | 7.91 | 0.0008 | 0.040 | 0.06 |
| 1043202 | TTS | ENSSSCG00000040416 | 2 | 5886993 | 5889017 | 6.60 | 5.61 | 7.62 | 17.85 | 14.07 | 17.34 | 6.61 | 16.42 | 0.40 | -1.31 | 11.51 | 7.91 | 0.0008 | 0.040 | 0.06 |
| 1045468 | MSKIP_ON | ENSSSCG00000008386 | 3 | 80725341 | 80728852 | 2.89 | 1.39 | 2.22 | 5.84 | 5.07 | 5.27 | 2.17 | 5.40 | 0.40 | -1.32 | 3.78 | 6.97 | 0.0014 | 0.049 | -0.53 |
| 1003725 | TTS | ENSSSCG00000005272 | 1 | 228110584 | 228113906 | 0.62 | 0.52 | 0.93 | 1.58 | 1.70 | 1.89 | 0.69 | 1.72 | 0.40 | -1.32 | 1.21 | 6.99 | 0.0014 | 0.048 | -0.52 |
| 1066031 | TTS | ENSSSCG00000001095 | 7 | 19594926 | 19595084 | 8.55 | 8.40 | 7.30 | 18.22 | 20.48 | 21.89 | 8.09 | 20.20 | 0.40 | -1.32 | 14.14 | 11.28 | 0.0002 | 0.022 | 1.72 |
| 1074876 | IR_OFF | ENSSSCG00000015120 | 9 | 46565827 | 46566044 | 5.25 | 2.57 | 4.08 | 10.61 | 9.51 | 9.62 | 3.97 | 9.92 | 0.40 | -1.32 | 6.94 | 7.43 | 0.0011 | 0.044 | -0.23 |
| 1021241 | SKIP_ON | ENSSSCG00000009578 | 14 | 20143 | 20331 | 0.46 | 0.71 | 0.46 | 1.38 | 1.50 | 1.18 | 0.54 | 1.35 | 0.40 | -1.32 | 0.95 | 6.47 | 0.0019 | 0.055 | -0.88 |
| 1063119 | TTS | ENSSSCG00000030018 | 6 | 119185481 | 119188299 | 2.72 | 1.36 | 1.99 | 5.18 | 4.55 | 5.46 | 2.02 | 5.06 | 0.40 | -1.32 | 3.54 | 6.77 | 0.0016 | 0.051 | -0.67 |
| 1013571 | TSS | ENSSSCG00000023423 | 12 | 5208271 | 5208679 | 18.61 | 14.57 | 16.37 | 39.83 | 38.40 | 45.95 | 16.51 | 41.39 | 0.40 | -1.33 | 28.95 | 10.19 | 0.0003 | 0.026 | 1.25 |
| 1013572 | TTS | ENSSSCG00000023423 | 12 | 5203195 | 5204286 | 18.61 | 14.57 | 16.37 | 39.83 | 38.40 | 45.95 | 16.51 | 41.39 | 0.40 | -1.33 | 28.95 | 10.19 | 0.0003 | 0.026 | 1.25 |
| 1074206 | SKIP_OFF | ENSSSCG00000014909 | 9 | 19639974 | 19640021 | 0.79 | 0.99 | 1.26 | 2.25 | 2.45 | 2.92 | 1.01 | 2.54 | 0.40 | -1.33 | 1.78 | 6.69 | 0.0017 | 0.052 | -0.72 |
| 1029414 | SKIP_OFF | ENSSSCG00000029002 | 15 | 120347643 | 120347759 | 3.67 | 4.07 | 3.73 | 10.08 | 9.23 | 9.50 | 3.82 | 9.60 | 0.40 | -1.33 | 6.71 | 21.66 | 0.0000 | 0.009 | 4.35 |
| 1029412 | TTS | ENSSSCG00000029002 | 15 | 120409735 | 120411641 | 3.67 | 4.07 | 3.73 | 10.08 | 9.23 | 9.50 | 3.82 | 9.60 | 0.40 | -1.33 | 6.71 | 21.66 | 0.0000 | 0.009 | 4.35 |
| 1073937 | TTS | ENSSSCG00000014818 | 9 | 7301154 | 7303228 | 2.51 | 2.98 | 3.87 | 8.53 | 7.05 | 7.95 | 3.12 | 7.84 | 0.40 | -1.33 | 5.48 | 8.52 | 0.0006 | 0.035 | 0.42 |
| 1069830 | TSS | ENSSSCG00000033093 | 7 | 95746786 | 95746887 | 0.76 | 1.06 | 1.41 | 2.51 | 2.77 | 2.85 | 1.08 | 2.71 | 0.40 | -1.33 | 1.89 | 8.03 | 0.0008 | 0.039 | 0.14 |
| 1069831 | TTS | ENSSSCG00000033093 | 7 | 95926094 | 95930034 | 0.76 | 1.06 | 1.41 | 2.51 | 2.77 | 2.85 | 1.08 | 2.71 | 0.40 | -1.33 | 1.89 | 8.03 | 0.0008 | 0.039 | 0.14 |
| 1062528 | TSS | ENSSSCG00000027169 | 6 | 105613258 | 105613386 | 2.84 | 3.24 | 3.50 | 7.58 | 7.96 | 8.59 | 3.19 | 8.04 | 0.40 | -1.33 | 5.62 | 14.57 | 0.0001 | 0.015 | 2.84 |
| 1062529 | TTS | ENSSSCG00000027169 | 6 | 105645096 | 105645113 | 2.84 | 3.24 | 3.50 | 7.58 | 7.96 | 8.59 | 3.19 | 8.04 | 0.40 | -1.33 | 5.62 | 14.57 | 0.0001 | 0.015 | 2.84 |
| 1010604 | TTS | ENSSSCG00000035318 | 11 | 3265765 | 3266161 | 0.81 | 0.94 | 0.77 | 2.25 | 1.98 | 2.12 | 0.84 | 2.12 | 0.40 | -1.33 | 1.48 | 13.13 | 0.0001 | 0.018 | 2.40 |
| 1063392 | IR_OFF | ENSSSCG00000031781 | 6 | 28545357 | 28546178 | 8.70 | 8.18 | 6.63 | 22.23 | 18.01 | 19.12 | 7.84 | 19.78 | 0.40 | -1.34 | 13.81 | 9.01 | 0.0005 | 0.032 | 0.68 |

| | | | | | | | | | | | | | | | | | | | | |
|---|---|---|---|---|---|---|---|---|---|---|---|---|---|---|---|---|---|---|---|---|
| 1048013 | TSS | ENSSSCG00000035256 | 3 | 17978746 | 17978861 | 0.57 | 0.52 | 0.72 | 1.57 | 1.69 | 1.32 | 0.60 | 1.53 | 0.40 | -1.34 | 1.07 | 7.42 | 0.0011 | 0.044 | -0.23 |
| 1048014 | TTS | ENSSSCG00000035256 | 3 | 17979260 | 17985101 | 0.57 | 0.52 | 0.72 | 1.57 | 1.69 | 1.32 | 0.60 | 1.53 | 0.40 | -1.34 | 1.07 | 7.42 | 0.0011 | 0.044 | -0.23 |
| 1002951 | TTS | ENSSSCG00000004983 | 1 | 169867741 | 169867907 | 8.62 | 6.82 | 7.02 | 21.19 | 15.51 | 20.49 | 7.49 | 19.06 | 0.39 | -1.35 | 13.27 | 6.55 | 0.0018 | 0.054 | -0.82 |
| 1004016 | TSS | ENSSSCG00000005376 | 1 | 240049360 | 240049734 | 0.96 | 1.06 | 1.09 | 3.05 | 2.58 | 2.29 | 1.04 | 2.64 | 0.39 | -1.35 | 1.84 | 7.51 | 0.0010 | 0.043 | -0.18 |
| 1004017 | TTS | ENSSSCG00000005376 | 1 | 240004944 | 240005157 | 0.96 | 1.06 | 1.09 | 3.05 | 2.58 | 2.29 | 1.04 | 2.64 | 0.39 | -1.35 | 1.84 | 7.51 | 0.0010 | 0.043 | -0.18 |
| 1017541 | TSS | ENSSSCG00000011849 | 13 | 134098408 | 134098661 | 1.62 | 1.06 | 1.60 | 3.31 | 3.42 | 4.16 | 1.42 | 3.63 | 0.39 | -1.35 | 2.53 | 7.17 | 0.0012 | 0.047 | -0.40 |
| 1062740 | SKIP_OFF | ENSSSCG00000028126 | 6 | 97325537 | 97325616 | 2.09 | 1.23 | 2.35 | 4.67 | 4.89 | 4.96 | 1.89 | 4.84 | 0.39 | -1.36 | 3.37 | 8.91 | 0.0005 | 0.033 | 0.62 |
| 1043924 | SKIP_OFF | ENSSSCG00000007814 | 3 | 19240628 | 19240687 | 0.62 | 0.61 | 0.94 | 1.87 | 1.61 | 2.08 | 0.72 | 1.85 | 0.39 | -1.36 | 1.29 | 6.76 | 0.0016 | 0.051 | -0.68 |
| 1049488 | TSS | ENSSSCG00000006108 | 4 | 42930834 | 42931137 | 6.61 | 7.21 | 9.60 | 23.19 | 18.29 | 18.83 | 7.81 | 20.10 | 0.39 | -1.36 | 13.96 | 7.25 | 0.0012 | 0.046 | -0.34 |
| 1001960 | IR_OFF | ENSSSCG00000004651 | 1 | 122426153 | 122426241 | 1.73 | 2.32 | 1.41 | 4.55 | 4.70 | 4.81 | 1.82 | 4.69 | 0.39 | -1.37 | 3.25 | 10.79 | 0.0002 | 0.024 | 1.52 |
| 1029968 | MSKIP_OFF | ENSSSCG00000036679 | 15 | 46633434 | 46635454 | 0.69 | 0.80 | 0.96 | 2.01 | 1.94 | 2.36 | 0.81 | 2.10 | 0.39 | -1.37 | 1.46 | 8.75 | 0.0005 | 0.034 | 0.54 |
| 1002141 | TTS | ENSSSCG00000004704 | 1 | 127913530 | 127915445 | 1.69 | 1.62 | 2.79 | 5.82 | 4.75 | 5.31 | 2.03 | 5.29 | 0.38 | -1.38 | 3.66 | 7.04 | 0.0013 | 0.048 | -0.48 |
| 1029965 | SKIP_ON | ENSSSCG00000036679 | 15 | 46621430 | 46622995 | 0.70 | 0.80 | 0.97 | 2.06 | 1.99 | 2.39 | 0.82 | 2.15 | 0.38 | -1.38 | 1.48 | 9.30 | 0.0004 | 0.030 | 0.83 |
| 1013448 | SKIP_ON | ENSSSCG00000022196 | 12 | 40212120 | 40212293 | 0.89 | 0.46 | 0.49 | 1.61 | 1.62 | 1.58 | 0.62 | 1.60 | 0.38 | -1.38 | 1.11 | 7.20 | 0.0012 | 0.046 | -0.38 |
| 1013447 | TTS | ENSSSCG00000022196 | 12 | 40229934 | 40230006 | 0.89 | 0.46 | 0.49 | 1.61 | 1.62 | 1.58 | 0.62 | 1.60 | 0.38 | -1.38 | 1.11 | 7.20 | 0.0012 | 0.046 | -0.38 |
| 1062737 | TSS | ENSSSCG00000028126 | 6 | 97324691 | 97324885 | 2.37 | 1.50 | 2.82 | 5.61 | 5.94 | 5.90 | 2.23 | 5.82 | 0.38 | -1.38 | 4.03 | 9.47 | 0.0004 | 0.029 | 0.91 |
| 1062738 | TTS | ENSSSCG00000028126 | 6 | 97343305 | 97345993 | 2.37 | 1.50 | 2.82 | 5.61 | 5.94 | 5.90 | 2.23 | 5.82 | 0.38 | -1.38 | 4.03 | 9.47 | 0.0004 | 0.029 | 0.91 |
| 1071350 | TTS | ENSSSCG00000008942 | 8 | 67695793 | 67702227 | 12.21 | 10.94 | 11.11 | 32.40 | 30.53 | 26.50 | 11.42 | 29.81 | 0.38 | -1.38 | 20.61 | 10.94 | 0.0002 | 0.023 | 1.58 |
| 1067624 | TSS | ENSSSCG00000001913 | 7 | 59452117 | 59452420 | 4.01 | 2.89 | 3.89 | 8.11 | 9.39 | 10.71 | 3.60 | 9.40 | 0.38 | -1.39 | 6.50 | 7.41 | 0.0011 | 0.044 | -0.24 |
| 1067625 | TTS | ENSSSCG00000001913 | 7 | 59460278 | 59461206 | 4.01 | 2.89 | 3.89 | 8.11 | 9.39 | 10.71 | 3.60 | 9.40 | 0.38 | -1.39 | 6.50 | 7.41 | 0.0011 | 0.044 | -0.24 |
| 1070064 | IR_ON | ENSSSCG00000035153 | 7 | 20684479 | 20684559 | 2.03 | 1.65 | 2.57 | 5.67 | 4.81 | 5.90 | 2.08 | 5.46 | 0.38 | -1.39 | 3.77 | 8.41 | 0.0006 | 0.036 | 0.36 |
| 1070062 | MIR_ON | ENSSSCG00000035153 | 7 | 20684479 | 20684750 | 2.03 | 1.65 | 2.57 | 5.67 | 4.81 | 5.90 | 2.08 | 5.46 | 0.38 | -1.39 | 3.77 | 8.41 | 0.0006 | 0.036 | 0.36 |
| 1079416 | SKIP_ON | ENSSSCG00000012619 | X | 98040878 | 98040922 | 1.23 | 2.14 | 1.74 | 4.19 | 4.54 | 4.73 | 1.71 | 4.48 | 0.38 | -1.39 | 3.10 | 9.54 | 0.0004 | 0.029 | 0.95 |
| 1075717 | TSS | ENSSSCG00000015434 | 9 | 105985831 | 105986060 | 33.61 | 31.34 | 33.17 | 90.19 | 70.87 | 96.79 | 32.71 | 85.95 | 0.38 | -1.39 | 59.33 | 7.25 | 0.0012 | 0.046 | -0.35 |
| 1075720 | XAE | ENSSSCG00000015434 | 9 | 105984989 | 105985060 | 33.61 | 31.34 | 33.17 | 90.19 | 70.87 | 96.79 | 32.71 | 85.95 | 0.38 | -1.39 | 59.33 | 7.25 | 0.0012 | 0.046 | -0.35 |
| 1001823 | TSS | ENSSSCG00000004617 | 1 | 119020990 | 119021978 | 5.38 | 4.61 | 7.10 | 16.18 | 14.53 | 14.30 | 5.70 | 15.00 | 0.38 | -1.40 | 10.35 | 10.45 | 0.0002 | 0.025 | 1.37 |
| 1002149 | SKIP_OFF | ENSSSCG00000004704 | 1 | 127910069 | 127910182 | 0.37 | 0.34 | 0.44 | 0.88 | 0.97 | 1.16 | 0.38 | 1.01 | 0.38 | -1.40 | 0.69 | 6.81 | 0.0016 | 0.050 | -0.64 |
| 1002140 | TSS | ENSSSCG00000004704 | 1 | 127869890 | 127869923 | 0.37 | 0.34 | 0.44 | 0.88 | 0.97 | 1.16 | 0.38 | 1.01 | 0.38 | -1.40 | 0.69 | 6.81 | 0.0016 | 0.050 | -0.64 |
| 1079562 | SKIP_OFF | ENSSSCG00000012660 | X | 106829230 | 106829476 | 1.99 | 1.37 | 2.26 | 5.24 | 4.39 | 5.21 | 1.87 | 4.94 | 0.38 | -1.40 | 3.41 | 8.51 | 0.0006 | 0.035 | 0.41 |
| 1013446 | TSS | ENSSSCG00000022196 | 12 | 40205096 | 40205358 | 0.89 | 0.46 | 0.49 | 1.61 | 1.62 | 1.65 | 0.62 | 1.63 | 0.38 | -1.40 | 1.12 | 7.36 | 0.0011 | 0.045 | -0.28 |
| 1002581 | TSS | ENSSSCG00000004856 | 1 | 146002702 | 146003308 | 2.05 | 2.05 | 2.24 | 6.04 | 5.25 | 5.61 | 2.12 | 5.63 | 0.38 | -1.41 | 3.87 | 15.60 | 0.0000 | 0.013 | 3.12 |
| 1075718 | TTS | ENSSSCG00000015434 | 9 | 105951383 | 105954349 | 37.64 | 36.04 | 37.10 | 102.56 | 82.22 | 110.32 | 36.93 | 98.37 | 0.38 | -1.41 | 67.65 | 7.78 | 0.0009 | 0.041 | -0.01 |
| 1049492 | SKIP_ON | ENSSSCG00000006108 | 4 | 42884699 | 42884803 | 6.75 | 7.13 | 9.43 | 24.60 | 18.42 | 19.19 | 7.77 | 20.74 | 0.37 | -1.42 | 14.25 | 6.51 | 0.0019 | 0.054 | -0.85 |
| 1076007 | XSKIP_ON | ENSSSCG00000015549 | 9 | 123909838 | 123910034 | 1.26 | 1.27 | 1.56 | 3.66 | 3.44 | 3.84 | 1.37 | 3.65 | 0.37 | -1.42 | 2.51 | 15.67 | 0.0000 | 0.013 | 3.14 |
| 1076338 | MSKIP_OFF | ENSSSCG00000015664 | 9 | 67954724 | 67958377 | 3.63 | 2.29 | 2.31 | 6.99 | 7.06 | 7.94 | 2.74 | 7.33 | 0.37 | -1.42 | 5.04 | 9.03 | 0.0005 | 0.032 | 0.69 |
| 1076341 | SKIP_OFF | ENSSSCG00000015664 | 9 | 67954724 | 67954744 | 3.63 | 2.29 | 2.31 | 6.99 | 7.06 | 7.94 | 2.74 | 7.33 | 0.37 | -1.42 | 5.04 | 9.03 | 0.0005 | 0.032 | 0.69 |
| 1076331 | SKIP_ON | ENSSSCG00000015664 | 9 | 67952156 | 67952176 | 3.63 | 2.29 | 2.31 | 6.99 | 7.06 | 7.94 | 2.74 | 7.33 | 0.37 | -1.42 | 5.04 | 9.03 | 0.0005 | 0.032 | 0.69 |
| 1076319 | XSKIP_ON | ENSSSCG00000015664 | 9 | 67947529 | 67947897 | 3.63 | 2.29 | 2.31 | 6.99 | 7.06 | 7.94 | 2.74 | 7.33 | 0.37 | -1.42 | 5.04 | 9.03 | 0.0005 | 0.032 | 0.69 |
| 1066029 | TSS | ENSSSCG00000001095 | 7 | 19586002 | 19586585 | 5.03 | 5.04 | 4.33 | 11.35 | 13.46 | 13.69 | 4.80 | 12.83 | 0.37 | -1.42 | 8.82 | 10.90 | 0.0002 | 0.023 | 1.56 |
| 1063389 | TSS | ENSSSCG00000031781 | 6 | 28548002 | 28548152 | 8.31 | 7.77 | 6.09 | 22.23 | 18.01 | 19.12 | 7.39 | 19.78 | 0.37 | -1.42 | 13.59 | 9.20 | 0.0004 | 0.031 | 0.78 |
| 1006689 | SKIP_OFF | ENSSSCG00000037508 | 1 | 261345165 | 261345226 | 1.48 | 2.06 | 1.69 | 4.70 | 4.78 | 4.58 | 1.75 | 4.69 | 0.37 | -1.43 | 3.22 | 16.97 | 0.0000 | 0.012 | 3.46 |
| 1006690 | SKIP_ON | ENSSSCG00000037508 | 1 | 261326633 | 261326716 | 1.48 | 2.06 | 1.69 | 4.70 | 4.78 | 4.58 | 1.75 | 4.69 | 0.37 | -1.43 | 3.22 | 16.97 | 0.0000 | 0.012 | 3.46 |
| 1069237 | TSS | ENSSSCG00000025768 | 7 | 112160789 | 112161017 | 83.68 | 73.90 | 101.76 | 238.13 | 209.30 | 252.83 | 86.44 | 233.42 | 0.37 | -1.43 | 159.93 | 10.30 | 0.0003 | 0.026 | 1.30 |
| 1017566 | AE | ENSSSCG00000011849 | 13 | 134122913 | 134123040 | 2.78 | 1.39 | 1.84 | 5.70 | 5.15 | 5.44 | 2.00 | 5.43 | 0.37 | -1.44 | 3.71 | 8.26 | 0.0007 | 0.037 | 0.27 |
| 1040768 | TSS | ENSSSCG00000024389 | 2 | 229252 | 229570 | 1.31 | 1.37 | 1.54 | 4.33 | 3.08 | 4.05 | 1.41 | 3.82 | 0.37 | -1.44 | 2.61 | 6.59 | 0.0018 | 0.053 | -0.80 |
| 1040769 | TTS | ENSSSCG00000024389 | 2 | 221989 | 223854 | 1.31 | 1.37 | 1.54 | 4.33 | 3.08 | 4.05 | 1.41 | 3.82 | 0.37 | -1.44 | 2.61 | 6.59 | 0.0018 | 0.053 | -0.80 |
| 1029989 | AE | ENSSSCG00000036679 | 15 | 46673660 | 46673715 | 3.51 | 3.90 | 5.22 | 12.20 | 10.17 | 11.89 | 4.21 | 11.42 | 0.37 | -1.44 | 7.82 | 9.35 | 0.0004 | 0.030 | 0.86 |
| 1070060 | SKIP_ON | ENSSSCG00000035153 | 7 | 20682902 | 20682924 | 1.75 | 1.52 | 2.42 | 5.32 | 4.58 | 5.61 | 1.90 | 5.17 | 0.37 | -1.45 | 3.53 | 8.45 | 0.0006 | 0.036 | 0.38 |
| 1050088 | XSKIP_ON | ENSSSCG00000006309 | 4 | 83579006 | 83579041 | 1.86 | 1.13 | 1.86 | 4.87 | 4.34 | 4.08 | 1.62 | 4.43 | 0.37 | -1.45 | 3.02 | 8.77 | 0.0005 | 0.033 | 0.55 |
| 1029983 | MSKIP_ON | ENSSSCG00000036679 | 15 | 46684528 | 46727479 | 2.81 | 3.10 | 4.26 | 10.15 | 8.18 | 9.50 | 3.39 | 9.28 | 0.37 | -1.45 | 6.33 | 8.58 | 0.0006 | 0.035 | 0.45 |
| 1029986 | SKIP_OFF | ENSSSCG00000036679 | 15 | 46636139 | 46636297 | 2.81 | 3.10 | 4.26 | 10.15 | 8.18 | 9.50 | 3.39 | 9.28 | 0.37 | -1.45 | 6.33 | 8.58 | 0.0006 | 0.035 | 0.45 |
| 1029969 | SKIP_ON | ENSSSCG00000036679 | 15 | 46727424 | 46727479 | 2.81 | 3.10 | 4.26 | 10.15 | 8.18 | 9.50 | 3.39 | 9.28 | 0.37 | -1.45 | 6.33 | 8.58 | 0.0006 | 0.035 | 0.45 |
| 1025401 | TTS | ENSSSCG00000025876 | 14 | 71442852 | 71443821 | 2.45 | 1.59 | 3.63 | 6.69 | 7.38 | 6.96 | 2.56 | 7.01 | 0.36 | -1.45 | 4.78 | 7.56 | 0.0010 | 0.043 | -0.15 |
| 1018959 | SKIP_ON | ENSSSCG00000025408 | 13 | 25726122 | 25726223 | 1.90 | 2.01 | 2.74 | 6.34 | 5.36 | 6.57 | 2.22 | 6.09 | 0.36 | -1.46 | 4.15 | 8.96 | 0.0005 | 0.032 | 0.65 |

| | | | | | | | | | | | | | | | | | | | | |
|---|---|---|---|---|---|---|---|---|---|---|---|---|---|---|---|---|---|---|---|---|
| 1029967 | MSKIP_ON | ENSSSCG00000036679 | 15 | 46633434 | 46635454 | 2.82 | 3.10 | 4.27 | 10.19 | 8.23 | 9.53 | 3.40 | 9.32 | 0.36 | -1.46 | 6.36 | 8.65 | 0.0006 | 0.034 | 0.49 |
| 1068589 | IR_OFF | ENSSSCG00000002428 | 7 | 110750762 | 110750877 | 5.80 | 5.65 | 7.01 | 17.61 | 13.94 | 19.11 | 6.15 | 16.89 | 0.36 | -1.46 | 11.52 | 7.15 | 0.0013 | 0.047 | -0.41 |
| 1068584 | TSS | ENSSSCG00000002428 | 7 | 110731668 | 110731940 | 5.80 | 5.65 | 7.01 | 17.61 | 13.94 | 19.11 | 6.15 | 16.89 | 0.36 | -1.46 | 11.52 | 7.15 | 0.0013 | 0.047 | -0.41 |
| 1021593 | TSS | ENSSSCG00000009714 | 14 | 20144601 | 20144723 | 1.07 | 1.16 | 0.59 | 2.24 | 2.59 | 2.90 | 0.94 | 2.58 | 0.36 | -1.46 | 1.76 | 6.64 | 0.0017 | 0.053 | -0.76 |
| 1068585 | TTS | ENSSSCG00000002428 | 7 | 110783385 | 110783501 | 5.80 | 5.65 | 7.01 | 17.61 | 14.06 | 19.11 | 6.15 | 16.93 | 0.36 | -1.46 | 11.54 | 7.35 | 0.0011 | 0.045 | -0.28 |
| 1022433 | SKIP_ON | ENSSSCG00000009956 | 14 | 43364583 | 43364656 | 2.91 | 3.28 | 2.66 | 8.03 | 7.50 | 8.82 | 2.95 | 8.12 | 0.36 | -1.46 | 5.53 | 12.88 | 0.0001 | 0.019 | 2.31 |
| 1018952 | TTS | ENSSSCG00000025408 | 13 | 25210498 | 25213056 | 1.90 | 1.98 | 2.70 | 6.28 | 5.36 | 6.50 | 2.19 | 6.04 | 0.36 | -1.46 | 4.12 | 9.43 | 0.0004 | 0.030 | 0.89 |
| 1070056 | TSS | ENSSSCG00000035153 | 7 | 20682082 | 20682755 | 1.98 | 1.91 | 2.87 | 6.08 | 5.41 | 7.28 | 2.26 | 6.26 | 0.36 | -1.47 | 4.26 | 6.78 | 0.0016 | 0.051 | -0.66 |
| 1043933 | SKIP_OFF | ENSSSCG00000007814 | 3 | 19274300 | 19274428 | 0.76 | 0.54 | 0.66 | 1.93 | 1.58 | 1.96 | 0.66 | 1.83 | 0.36 | -1.48 | 1.24 | 8.64 | 0.0006 | 0.034 | 0.49 |
| 1055225 | TSS | ENSSSCG00000000741 | 5 | 67280924 | 67281762 | 0.13 | 0.17 | 0.22 | 0.49 | 0.46 | 0.51 | 0.17 | 0.49 | 0.36 | -1.49 | 0.33 | 6.57 | 0.0018 | 0.054 | -0.81 |
| 1055226 | TTS | ENSSSCG00000000741 | 5 | 67290121 | 67292348 | 0.13 | 0.17 | 0.22 | 0.49 | 0.46 | 0.51 | 0.17 | 0.49 | 0.36 | -1.49 | 0.33 | 6.57 | 0.0018 | 0.054 | -0.81 |
| 1076005 | TSS | ENSSSCG00000015549 | 9 | 123910348 | 123910397 | 1.36 | 1.36 | 1.93 | 4.63 | 4.03 | 4.37 | 1.55 | 4.34 | 0.36 | -1.49 | 2.95 | 11.35 | 0.0002 | 0.022 | 1.75 |
| 1076006 | TTS | ENSSSCG00000015549 | 9 | 123892115 | 123895470 | 1.36 | 1.36 | 1.93 | 4.63 | 4.03 | 4.37 | 1.55 | 4.34 | 0.36 | -1.49 | 2.95 | 11.35 | 0.0002 | 0.022 | 1.75 |
| 1032862 | TSS | ENSSSCG00000007454 | 17 | 49337732 | 49337793 | 7.89 | 5.62 | 6.88 | 20.78 | 16.88 | 19.44 | 6.79 | 19.03 | 0.36 | -1.49 | 12.91 | 9.86 | 0.0003 | 0.028 | 1.10 |
| 1032863 | TTS | ENSSSCG00000007454 | 17 | 49258553 | 49260250 | 7.89 | 5.62 | 6.88 | 20.78 | 16.88 | 19.44 | 6.79 | 19.03 | 0.36 | -1.49 | 12.91 | 9.86 | 0.0003 | 0.028 | 1.10 |
| 1018951 | TSS | ENSSSCG00000025408 | 13 | 25751275 | 25751362 | 1.90 | 2.01 | 2.74 | 6.66 | 5.36 | 6.64 | 2.22 | 6.22 | 0.36 | -1.49 | 4.22 | 8.37 | 0.0006 | 0.036 | 0.33 |
| 1022694 | SKIP_ON | ENSSSCG00000010031 | 14 | 48298349 | 48298509 | 1.83 | 0.96 | 1.46 | 4.57 | 3.50 | 3.85 | 1.42 | 3.97 | 0.36 | -1.49 | 2.69 | 6.70 | 0.0017 | 0.052 | -0.72 |
| 1034429 | TSS | ENSSSCG00000016491 | 18 | 8205725 | 8205996 | 2.28 | 2.00 | 2.14 | 4.90 | 6.40 | 6.71 | 2.14 | 6.00 | 0.36 | -1.49 | 4.07 | 7.26 | 0.0012 | 0.046 | -0.34 |
| 1034430 | TTS | ENSSSCG00000016491 | 18 | 8256074 | 8256269 | 2.28 | 2.00 | 2.14 | 4.90 | 6.40 | 6.71 | 2.14 | 6.00 | 0.36 | -1.49 | 4.07 | 7.26 | 0.0012 | 0.046 | -0.34 |
| 1023179 | TSS | ENSSSCG00000010164 | 14 | 57480793 | 57481988 | 3.18 | 2.67 | 3.75 | 9.44 | 7.85 | 9.69 | 3.20 | 8.99 | 0.36 | -1.49 | 6.10 | 9.37 | 0.0004 | 0.030 | 0.86 |
| 1023180 | TTS | ENSSSCG00000010164 | 14 | 57424780 | 57428021 | 3.18 | 2.67 | 3.75 | 9.44 | 7.85 | 9.69 | 3.20 | 8.99 | 0.36 | -1.49 | 6.10 | 9.37 | 0.0004 | 0.030 | 0.86 |
| 1012768 | AE | ENSSSCG00000017829 | 12 | 48417303 | 48418064 | 0.43 | 0.25 | 0.55 | 1.12 | 1.24 | 1.10 | 0.41 | 1.15 | 0.36 | -1.49 | 0.78 | 7.49 | 0.0010 | 0.043 | -0.19 |
| 1012764 | IR_OFF | ENSSSCG00000017829 | 12 | 48456557 | 48456663 | 0.43 | 0.25 | 0.55 | 1.12 | 1.24 | 1.10 | 0.41 | 1.15 | 0.36 | -1.49 | 0.78 | 7.49 | 0.0010 | 0.043 | -0.19 |
| 1012754 | TSS | ENSSSCG00000017829 | 12 | 48417089 | 48417225 | 0.43 | 0.25 | 0.55 | 1.12 | 1.24 | 1.10 | 0.41 | 1.15 | 0.36 | -1.49 | 0.78 | 7.49 | 0.0010 | 0.043 | -0.19 |
| 1012762 | XIR_OFF | ENSSSCG00000017829 | 12 | 48417225 | 48417303 | 0.43 | 0.25 | 0.55 | 1.12 | 1.24 | 1.10 | 0.41 | 1.15 | 0.36 | -1.49 | 0.78 | 7.49 | 0.0010 | 0.043 | -0.19 |
| 1009292 | TSS | ENSSSCG00000009293 | 11 | 2829290 | 2829535 | 1.88 | 2.27 | 1.82 | 6.67 | 5.14 | 5.10 | 1.99 | 5.64 | 0.35 | -1.50 | 3.81 | 7.20 | 0.0012 | 0.046 | -0.38 |
| 1076252 | AE | ENSSSCG00000015645 | 9 | 66839798 | 66840598 | 1.79 | 1.43 | 1.47 | 4.94 | 4.23 | 4.12 | 1.56 | 4.43 | 0.35 | -1.50 | 3.00 | 10.75 | 0.0002 | 0.024 | 1.50 |
| 1015605 | TSS | ENSSSCG00000011314 | 13 | 28996446 | 28996615 | 17.74 | 9.82 | 16.65 | 46.62 | 35.74 | 43.09 | 14.74 | 41.82 | 0.35 | -1.50 | 28.28 | 7.10 | 0.0013 | 0.047 | -0.44 |
| 1015606 | TTS | ENSSSCG00000011314 | 13 | 28977503 | 28977963 | 17.74 | 9.82 | 16.65 | 46.62 | 35.74 | 43.09 | 14.74 | 41.82 | 0.35 | -1.50 | 28.28 | 7.10 | 0.0013 | 0.047 | -0.44 |
| 1017557 | MSKIP_ON | ENSSSCG00000011849 | 13 | 134122648 | 134123040 | 2.55 | 1.15 | 1.62 | 5.32 | 4.65 | 5.12 | 1.77 | 5.03 | 0.35 | -1.51 | 3.40 | 7.55 | 0.0010 | 0.043 | -0.16 |
| 1017558 | SKIP_ON | ENSSSCG00000011849 | 13 | 134122913 | 134123040 | 2.55 | 1.15 | 1.62 | 5.32 | 4.65 | 5.12 | 1.77 | 5.03 | 0.35 | -1.51 | 3.40 | 7.55 | 0.0010 | 0.043 | -0.16 |
| 1025402 | SKIP_ON | ENSSSCG00000025876 | 14 | 71456564 | 71456662 | 2.45 | 1.59 | 2.90 | 6.69 | 7.38 | 5.69 | 2.32 | 6.59 | 0.35 | -1.51 | 4.45 | 7.26 | 0.0012 | 0.046 | -0.34 |
| 1003920 | TSS | ENSSSCG00000005324 | 1 | 236550424 | 236551267 | 1.34 | 0.59 | 1.47 | 3.57 | 2.97 | 3.14 | 1.13 | 3.23 | 0.35 | -1.51 | 2.18 | 6.72 | 0.0016 | 0.052 | -0.70 |
| 1003921 | TTS | ENSSSCG00000005324 | 1 | 236539348 | 236539523 | 1.34 | 0.59 | 1.47 | 3.57 | 2.97 | 3.14 | 1.13 | 3.23 | 0.35 | -1.51 | 2.18 | 6.72 | 0.0016 | 0.052 | -0.70 |
| 1021957 | TSS | ENSSSCG00000009814 | 14 | 31259727 | 31260038 | 7.57 | 4.94 | 6.92 | 19.63 | 14.97 | 20.82 | 6.47 | 18.48 | 0.35 | -1.51 | 12.48 | 6.53 | 0.0018 | 0.054 | -0.84 |
| 1021958 | TTS | ENSSSCG00000009814 | 14 | 31306107 | 31306441 | 7.57 | 4.94 | 6.92 | 19.63 | 14.97 | 20.82 | 6.47 | 18.48 | 0.35 | -1.51 | 12.48 | 6.53 | 0.0018 | 0.054 | -0.84 |
| 1044489 | TTS | ENSSSCG00000008020 | 3 | 40371484 | 40372136 | 6.06 | 5.52 | 5.78 | 18.65 | 13.23 | 17.66 | 5.79 | 16.52 | 0.35 | -1.51 | 11.15 | 6.81 | 0.0016 | 0.050 | -0.64 |
| 1076248 | SKIP_OFF | ENSSSCG00000015645 | 9 | 67111018 | 67111056 | 1.83 | 1.43 | 1.47 | 4.94 | 4.35 | 4.24 | 1.58 | 4.51 | 0.35 | -1.52 | 3.04 | 12.20 | 0.0001 | 0.019 | 2.07 |
| 1076245 | SKIP_ON | ENSSSCG00000015645 | 9 | 67107456 | 67107602 | 1.83 | 1.43 | 1.47 | 4.94 | 4.35 | 4.24 | 1.58 | 4.51 | 0.35 | -1.52 | 3.04 | 12.20 | 0.0001 | 0.019 | 2.07 |
| 1044488 | TSS | ENSSSCG00000008020 | 3 | 40309635 | 40309750 | 6.06 | 5.45 | 5.74 | 18.62 | 13.16 | 17.63 | 5.75 | 16.47 | 0.35 | -1.52 | 11.11 | 6.74 | 0.0016 | 0.051 | -0.69 |
| 1052735 | TSS | ENSSSCG00000034398 | 4 | 41372785 | 41373922 | 3.46 | 3.17 | 3.55 | 9.94 | 10.04 | 9.26 | 3.39 | 9.75 | 0.35 | -1.52 | 6.57 | 24.56 | 0.0000 | 0.009 | 4.75 |
| 1052736 | TTS | ENSSSCG00000034398 | 4 | 41392294 | 41397071 | 3.46 | 3.17 | 3.55 | 9.94 | 10.04 | 9.26 | 3.39 | 9.75 | 0.35 | -1.52 | 6.57 | 24.56 | 0.0000 | 0.009 | 4.75 |
| 1047455 | TSS | ENSSSCG00000031876 | 3 | 24910674 | 24910849 | 4.86 | 5.23 | 7.67 | 15.49 | 18.83 | 16.86 | 5.92 | 17.06 | 0.35 | -1.53 | 11.49 | 9.04 | 0.0005 | 0.032 | 0.69 |
| 1062739 | SKIP_ON | ENSSSCG00000028126 | 6 | 97325537 | 97325616 | 0.27 | 0.27 | 0.47 | 0.94 | 1.05 | 0.94 | 0.34 | 0.97 | 0.35 | -1.53 | 0.66 | 7.91 | 0.0008 | 0.040 | 0.07 |
| 1037654 | SKIP_ON | ENSSSCG00000013494 | 2 | 74915417 | 74915482 | 0.48 | 0.61 | 0.51 | 1.48 | 1.41 | 1.76 | 0.54 | 1.55 | 0.35 | -1.53 | 1.04 | 8.80 | 0.0005 | 0.033 | 0.57 |
| 1067990 | SKIP_ON | ENSSSCG00000002041 | 7 | 76202082 | 76202212 | 0.48 | 0.66 | 0.81 | 2.10 | 1.63 | 1.89 | 0.65 | 1.87 | 0.35 | -1.53 | 1.26 | 7.58 | 0.0010 | 0.043 | -0.14 |
| 1067986 | TSS | ENSSSCG00000002041 | 7 | 76176590 | 76176688 | 0.48 | 0.66 | 0.81 | 2.10 | 1.63 | 1.89 | 0.65 | 1.87 | 0.35 | -1.53 | 1.26 | 7.58 | 0.0010 | 0.043 | -0.14 |
| 1017072 | MSKIP_ON | ENSSSCG00000011733 | 13 | 100063060 | 100066548 | 4.87 | 3.66 | 5.08 | 14.81 | 13.90 | 10.68 | 4.53 | 13.13 | 0.35 | -1.53 | 8.83 | 6.87 | 0.0015 | 0.050 | -0.60 |
| 1016624 | TSS | ENSSSCG00000011589 | 13 | 68828618 | 68828870 | 13.33 | 10.13 | 9.45 | 35.77 | 25.99 | 33.63 | 10.97 | 31.80 | 0.34 | -1.54 | 21.39 | 6.91 | 0.0015 | 0.049 | -0.57 |
| 1016626 | SKIP_ON | ENSSSCG00000011589 | 13 | 68877935 | 68877988 | 13.24 | 9.91 | 9.40 | 35.62 | 25.80 | 33.54 | 10.85 | 31.65 | 0.34 | -1.54 | 21.25 | 6.86 | 0.0015 | 0.050 | -0.60 |
| 1016625 | TTS | ENSSSCG00000011589 | 13 | 68901616 | 68902060 | 13.24 | 9.91 | 9.40 | 35.62 | 25.80 | 33.54 | 10.85 | 31.65 | 0.34 | -1.54 | 21.25 | 6.86 | 0.0015 | 0.050 | -0.60 |
| 1047458 | IR_ON | ENSSSCG00000031876 | 3 | 24908295 | 24908427 | 3.58 | 3.32 | 5.83 | 11.22 | 12.74 | 13.22 | 4.24 | 12.39 | 0.34 | -1.55 | 8.32 | 8.66 | 0.0006 | 0.034 | 0.49 |
| 1047456 | TTS | ENSSSCG00000031876 | 3 | 24876637 | 24877434 | 3.58 | 3.32 | 5.83 | 11.22 | 12.74 | 13.22 | 4.24 | 12.39 | 0.34 | -1.55 | 8.32 | 8.66 | 0.0006 | 0.034 | 0.49 |
| 1007767 | TSS | ENSSSCG00000011014 | 10 | 40055608 | 40057661 | 0.36 | 0.20 | 0.28 | 0.77 | 0.82 | 0.87 | 0.28 | 0.82 | 0.34 | -1.55 | 0.55 | 8.29 | 0.0007 | 0.037 | 0.29 |

| | | | | | | | | | | | | | | | | | | | | |
|---|---|---|---|---|---|---|---|---|---|---|---|---|---|---|---|---|---|---|---|---|
| 1075976 | SKIP_OFF | ENSSSCG00000015537 | 9 | 122374193 | 122374276 | 1.10 | 0.61 | 0.80 | 2.24 | 2.27 | 2.83 | 0.83 | 2.44 | 0.34 | -1.55 | 1.64 | 7.06 | 0.0013 | 0.048 | -0.47 |
| 1004665 | TSS | ENSSSCG00000005655 | 1 | 269030324 | 269030719 | 22.57 | 14.22 | 14.69 | 50.95 | 44.98 | 54.94 | 17.16 | 50.29 | 0.34 | -1.55 | 33.73 | 8.88 | 0.0005 | 0.033 | 0.61 |
| 1004666 | TTS | ENSSSCG00000005655 | 1 | 269002245 | 269002599 | 22.57 | 14.22 | 14.69 | 50.95 | 44.98 | 54.94 | 17.16 | 50.29 | 0.34 | -1.55 | 33.73 | 8.88 | 0.0005 | 0.033 | 0.61 |
| 1058127 | TSS | ENSSSCG00000002805 | 6 | 19969041 | 19969347 | 7.56 | 5.70 | 5.74 | 20.55 | 16.07 | 19.08 | 6.33 | 18.57 | 0.34 | -1.55 | 12.45 | 8.92 | 0.0005 | 0.032 | 0.63 |
| 1058128 | TTS | ENSSSCG00000002805 | 6 | 19948918 | 19950689 | 7.56 | 5.70 | 5.74 | 20.55 | 16.07 | 19.08 | 6.33 | 18.57 | 0.34 | -1.55 | 12.45 | 8.92 | 0.0005 | 0.032 | 0.63 |
| 1017062 | TSS | ENSSSCG00000011733 | 13 | 100077743 | 100077857 | 6.21 | 3.66 | 5.08 | 15.77 | 16.00 | 12.18 | 4.98 | 14.65 | 0.34 | -1.56 | 9.82 | 7.13 | 0.0013 | 0.047 | -0.43 |
| 1075716 | TSS | ENSSSCG00000015434 | 9 | 105985376 | 105985968 | 4.03 | 4.70 | 3.94 | 12.36 | 11.35 | 13.53 | 4.22 | 12.41 | 0.34 | -1.56 | 8.32 | 12.87 | 0.0001 | 0.019 | 2.31 |
| 1075719 | XAE | ENSSSCG00000015434 | 9 | 105984989 | 105985090 | 4.03 | 4.70 | 3.94 | 12.36 | 11.35 | 13.53 | 4.22 | 12.41 | 0.34 | -1.56 | 8.32 | 12.87 | 0.0001 | 0.019 | 2.31 |
| 1022928 | TTS | ENSSSCG00000010099 | 14 | 50593561 | 50594491 | 2.02 | 1.36 | 1.29 | 5.33 | 4.02 | 4.37 | 1.56 | 4.58 | 0.34 | -1.56 | 3.07 | 7.03 | 0.0014 | 0.048 | -0.49 |
| 1048105 | TSS | ENSSSCG00000036096 | 3 | 39263447 | 39264019 | 1.02 | 1.15 | 1.40 | 3.56 | 3.26 | 3.70 | 1.19 | 3.50 | 0.34 | -1.56 | 2.35 | 13.87 | 0.0001 | 0.017 | 2.63 |
| 1048106 | TTS | ENSSSCG00000036096 | 3 | 39261903 | 39262192 | 1.02 | 1.15 | 1.40 | 3.56 | 3.26 | 3.70 | 1.19 | 3.50 | 0.34 | -1.56 | 2.35 | 13.87 | 0.0001 | 0.017 | 2.63 |
| 1070554 | TSS | ENSSSCG00000039469 | 7 | 96999538 | 96999655 | 8.51 | 9.85 | 11.44 | 33.39 | 25.67 | 28.67 | 9.93 | 29.24 | 0.34 | -1.56 | 19.59 | 8.55 | 0.0006 | 0.035 | 0.43 |
| 1070555 | TTS | ENSSSCG00000039469 | 7 | 97041926 | 97047090 | 8.51 | 9.85 | 11.44 | 33.39 | 25.67 | 28.67 | 9.93 | 29.24 | 0.34 | -1.56 | 19.59 | 8.55 | 0.0006 | 0.035 | 0.43 |
| 1017692 | SKIP_ON | ENSSSCG00000011881 | 13 | 138736852 | 138736983 | 1.60 | 1.15 | 1.87 | 4.70 | 4.57 | 4.38 | 1.54 | 4.55 | 0.34 | -1.56 | 3.04 | 13.66 | 0.0001 | 0.017 | 2.56 |
| 1028643 | MSKIP_ON | ENSSSCG00000016263 | 15 | 131322801 | 131339081 | 2.29 | 1.82 | 2.71 | 8.01 | 6.40 | 5.73 | 2.27 | 6.72 | 0.34 | -1.56 | 4.49 | 6.52 | 0.0019 | 0.054 | -0.85 |
| 1028620 | TTS | ENSSSCG00000016263 | 15 | 131343089 | 131343135 | 2.29 | 1.82 | 2.71 | 8.01 | 6.40 | 5.73 | 2.27 | 6.72 | 0.34 | -1.56 | 4.49 | 6.52 | 0.0019 | 0.054 | -0.85 |
| 1037779 | TSS | ENSSSCG00000013546 | 2 | 72610577 | 72610691 | 1.05 | 1.64 | 1.79 | 4.67 | 4.01 | 4.59 | 1.49 | 4.42 | 0.34 | -1.57 | 2.96 | 9.98 | 0.0003 | 0.027 | 1.16 |
| 1037780 | TTS | ENSSSCG00000013546 | 2 | 72619546 | 72620207 | 1.05 | 1.64 | 1.79 | 4.67 | 4.01 | 4.59 | 1.49 | 4.42 | 0.34 | -1.57 | 2.96 | 9.98 | 0.0003 | 0.027 | 1.16 |
| 1025405 | IR_OFF | ENSSSCG00000025876 | 14 | 71445904 | 71448305 | 2.07 | 1.39 | 2.89 | 5.80 | 6.79 | 6.23 | 2.12 | 6.27 | 0.34 | -1.57 | 4.19 | 8.44 | 0.0006 | 0.036 | 0.37 |
| 1001798 | MSKIP_ON | ENSSSCG00000004612 | 1 | 116551464 | 116574940 | 0.82 | 0.96 | 0.51 | 2.57 | 2.30 | 1.94 | 0.76 | 2.27 | 0.34 | -1.57 | 1.52 | 7.02 | 0.0014 | 0.048 | -0.50 |
| 1001805 | SKIP_ON | ENSSSCG00000004612 | 1 | 116574759 | 116574940 | 0.82 | 0.96 | 0.51 | 2.57 | 2.30 | 1.94 | 0.76 | 2.27 | 0.34 | -1.57 | 1.52 | 7.02 | 0.0014 | 0.048 | -0.50 |
| 1001807 | SKIP_ON | ENSSSCG00000004612 | 1 | 116551464 | 116551581 | 0.82 | 0.96 | 0.51 | 2.57 | 2.30 | 1.94 | 0.76 | 2.27 | 0.34 | -1.57 | 1.52 | 7.02 | 0.0014 | 0.048 | -0.50 |
| 1058928 | AE | ENSSSCG00000003152 | 6 | 54253754 | 54254088 | 0.41 | 0.29 | 0.28 | 0.82 | 1.03 | 1.06 | 0.33 | 0.97 | 0.34 | -1.58 | 0.65 | 7.18 | 0.0012 | 0.046 | -0.39 |
| 1058925 | TSS | ENSSSCG00000003152 | 6 | 54250531 | 54250860 | 0.41 | 0.29 | 0.28 | 0.82 | 1.03 | 1.06 | 0.33 | 0.97 | 0.34 | -1.58 | 0.65 | 7.18 | 0.0012 | 0.046 | -0.39 |
| 1058926 | TTS | ENSSSCG00000003152 | 6 | 54266697 | 54266893 | 0.41 | 0.29 | 0.28 | 0.82 | 1.03 | 1.06 | 0.33 | 0.97 | 0.34 | -1.58 | 0.65 | 7.18 | 0.0012 | 0.046 | -0.39 |
| 1057699 | TSS | ENSSSCG00000002637 | 6 | 112234 | 112267 | 2.46 | 1.53 | 3.62 | 7.26 | 6.95 | 8.53 | 2.54 | 7.58 | 0.33 | -1.58 | 5.06 | 6.92 | 0.0014 | 0.049 | -0.57 |
| 1057700 | TTS | ENSSSCG00000002637 | 6 | 117929 | 118086 | 2.46 | 1.53 | 3.62 | 7.26 | 6.95 | 8.53 | 2.54 | 7.58 | 0.33 | -1.58 | 5.06 | 6.92 | 0.0014 | 0.049 | -0.57 |
| 1020171 | XSKIP_OFF | ENSSSCG00000033860 | 13 | 117146539 | 117146594 | 4.80 | 4.67 | 5.48 | 17.68 | 14.71 | 12.45 | 4.98 | 14.94 | 0.33 | -1.58 | 9.96 | 6.88 | 0.0015 | 0.050 | -0.59 |
| 1018955 | SKIP_ON | ENSSSCG00000025408 | 13 | 25634994 | 25635155 | 1.27 | 1.63 | 1.89 | 5.32 | 4.52 | 4.52 | 1.60 | 4.79 | 0.33 | -1.58 | 3.19 | 10.54 | 0.0002 | 0.025 | 1.41 |
| 1018957 | SKIP_ON | ENSSSCG00000025408 | 13 | 25646911 | 25646994 | 1.27 | 1.63 | 1.89 | 5.32 | 4.52 | 4.52 | 1.60 | 4.79 | 0.33 | -1.58 | 3.19 | 10.54 | 0.0002 | 0.025 | 1.41 |
| 1069540 | SKIP_ON | ENSSSCG00000030626 | 7 | 53785612 | 53785713 | 0.42 | 0.45 | 0.44 | 1.17 | 1.38 | 1.40 | 0.44 | 1.32 | 0.33 | -1.59 | 0.88 | 10.87 | 0.0002 | 0.024 | 1.55 |
| 1069537 | TSS | ENSSSCG00000030626 | 7 | 53794648 | 53794796 | 0.42 | 0.45 | 0.44 | 1.17 | 1.38 | 1.40 | 0.44 | 1.32 | 0.33 | -1.59 | 0.88 | 10.87 | 0.0002 | 0.024 | 1.55 |
| 1018456 | TSS | ENSSSCG00000022618 | 13 | 18243374 | 18243908 | 4.79 | 3.26 | 3.17 | 11.56 | 9.25 | 13.01 | 3.74 | 11.27 | 0.33 | -1.59 | 7.51 | 6.59 | 0.0018 | 0.053 | -0.80 |
| 1018457 | TTS | ENSSSCG00000022618 | 13 | 18349953 | 18350381 | 4.79 | 3.26 | 3.17 | 11.56 | 9.25 | 13.01 | 3.74 | 11.27 | 0.33 | -1.59 | 7.51 | 6.59 | 0.0018 | 0.053 | -0.80 |
| 1015607 | SKIP_ON | ENSSSCG00000011314 | 13 | 28990147 | 28990271 | 15.61 | 8.39 | 16.17 | 45.45 | 34.50 | 41.36 | 13.39 | 40.44 | 0.33 | -1.59 | 26.91 | 7.08 | 0.0013 | 0.047 | -0.46 |
| 1033030 | TSS | ENSSSCG00000007504 | 17 | 57783154 | 57783577 | 10.13 | 5.13 | 3.94 | 18.81 | 18.94 | 20.29 | 6.40 | 19.35 | 0.33 | -1.60 | 12.87 | 7.03 | 0.0014 | 0.048 | -0.49 |
| 1033031 | TTS | ENSSSCG00000007504 | 17 | 57796389 | 57797990 | 10.13 | 5.13 | 3.94 | 18.81 | 18.94 | 20.29 | 6.40 | 19.35 | 0.33 | -1.60 | 12.87 | 7.03 | 0.0014 | 0.048 | -0.49 |
| 1043084 | TTS | ENSSSCG00000039731 | 2 | 102427004 | 102427598 | 1.60 | 2.26 | 1.92 | 6.26 | 5.23 | 6.00 | 1.93 | 5.83 | 0.33 | -1.60 | 3.88 | 11.31 | 0.0002 | 0.022 | 1.73 |
| 1034185 | TSS | ENSSSCG00000040392 | 17 | 49380787 | 49380788 | 4.40 | 2.72 | 2.63 | 9.68 | 8.69 | 11.08 | 3.25 | 9.82 | 0.33 | -1.60 | 6.53 | 7.72 | 0.0009 | 0.041 | -0.05 |
| 1034186 | TTS | ENSSSCG00000040392 | 17 | 49320614 | 49320680 | 4.40 | 2.72 | 2.63 | 9.68 | 8.69 | 11.08 | 3.25 | 9.82 | 0.33 | -1.60 | 6.53 | 7.72 | 0.0009 | 0.041 | -0.05 |
| 1073921 | TSS | ENSSSCG00000014812 | 9 | 6785303 | 6785635 | 2.70 | 2.15 | 1.88 | 7.97 | 5.58 | 6.79 | 2.24 | 6.78 | 0.33 | -1.60 | 4.51 | 6.60 | 0.0018 | 0.053 | -0.79 |
| 1001955 | SKIP_ON | ENSSSCG00000004651 | 1 | 122514570 | 122514658 | 3.59 | 3.57 | 3.06 | 10.39 | 9.90 | 10.80 | 3.41 | 10.36 | 0.33 | -1.60 | 6.88 | 23.35 | 0.0000 | 0.009 | 4.59 |
| 1074212 | AE | ENSSSCG00000014909 | 9 | 19646385 | 19650290 | 0.51 | 0.27 | 0.35 | 1.06 | 1.29 | 1.05 | 0.37 | 1.13 | 0.33 | -1.61 | 0.75 | 7.10 | 0.0013 | 0.047 | -0.44 |
| 1074204 | TTS | ENSSSCG00000014909 | 9 | 19640768 | 19641300 | 0.51 | 0.27 | 0.35 | 1.06 | 1.29 | 1.05 | 0.37 | 1.13 | 0.33 | -1.61 | 0.75 | 7.10 | 0.0013 | 0.047 | -0.44 |
| 1068886 | TSS | ENSSSCG00000002520 | 7 | 121187349 | 121187737 | 4.72 | 2.48 | 3.16 | 12.03 | 9.88 | 9.67 | 3.45 | 10.53 | 0.33 | -1.61 | 6.99 | 7.49 | 0.0010 | 0.043 | -0.19 |
| 1028431 | TSS | ENSSSCG00000016214 | 15 | 121284920 | 121285147 | 3.86 | 5.32 | 3.00 | 13.98 | 10.22 | 13.12 | 4.06 | 12.44 | 0.33 | -1.62 | 8.25 | 6.72 | 0.0016 | 0.052 | -0.70 |
| 1017543 | TSS | ENSSSCG00000011849 | 13 | 134101345 | 134101771 | 1.71 | 1.60 | 1.60 | 5.22 | 4.71 | 5.14 | 1.64 | 5.02 | 0.33 | -1.62 | 3.33 | 21.69 | 0.0000 | 0.009 | 4.35 |
| 1020541 | SKIP_ON | ENSSSCG00000036033 | 13 | 11035201 | 11035264 | 1.34 | 1.80 | 2.03 | 5.85 | 5.20 | 4.83 | 1.72 | 5.29 | 0.33 | -1.62 | 3.51 | 10.41 | 0.0003 | 0.025 | 1.35 |
| 1008123 | MSKIP_OFF | ENSSSCG00000011090 | 10 | 53925155 | 53926881 | 4.67 | 4.38 | 3.18 | 13.69 | 10.85 | 13.28 | 4.08 | 12.61 | 0.32 | -1.63 | 8.34 | 9.06 | 0.0005 | 0.032 | 0.71 |
| 1050127 | TTS | ENSSSCG00000006325 | 4 | 85069603 | 85069852 | 35.85 | 22.64 | 32.54 | 93.16 | 77.86 | 111.26 | 30.34 | 94.09 | 0.32 | -1.63 | 62.22 | 6.49 | 0.0019 | 0.055 | -0.87 |
| 1009252 | TTS | ENSSSCG00000009270 | 11 | 1022644 | 1025613 | 9.36 | 5.90 | 6.56 | 25.66 | 19.22 | 22.90 | 7.27 | 22.59 | 0.32 | -1.64 | 14.93 | 7.58 | 0.0010 | 0.043 | -0.13 |
| 1000678 | TTS | ENSSSCG00000004222 | 1 | 37043805 | 37045906 | 1.45 | 1.00 | 1.02 | 3.52 | 3.07 | 4.21 | 1.16 | 3.60 | 0.32 | -1.64 | 2.38 | 7.13 | 0.0013 | 0.047 | -0.42 |
| 1000673 | TSS | ENSSSCG00000004222 | 1 | 37054910 | 37055099 | 1.45 | 1.00 | 1.02 | 3.52 | 3.07 | 4.21 | 1.16 | 3.60 | 0.32 | -1.64 | 2.38 | 7.13 | 0.0013 | 0.047 | -0.42 |
| 1017690 | TSS | ENSSSCG00000011881 | 13 | 138681330 | 138681831 | 1.48 | 1.12 | 1.51 | 4.59 | 4.23 | 4.00 | 1.37 | 4.27 | 0.32 | -1.64 | 2.82 | 14.10 | 0.0001 | 0.016 | 2.70 |

| | | | | | | | | | | | | | | | | | | | | |
|---|---|---|---|---|---|---|---|---|---|---|---|---|---|---|---|---|---|---|---|---|
| 1017691 | TTS | ENSSSCG00000011881 | 13 | 138740653 | 138743791 | 1.48 | 1.12 | 1.51 | 4.59 | 4.23 | 4.00 | 1.37 | 4.27 | 0.32 | -1.64 | 2.82 | 14.10 | 0.0001 | 0.016 | 2.70 |
| 1013375 | TTS | ENSSSCG00000021363 | 12 | 53113963 | 53117122 | 7.00 | 5.47 | 5.85 | 20.08 | 15.05 | 22.03 | 6.11 | 19.05 | 0.32 | -1.64 | 12.58 | 6.46 | 0.0019 | 0.055 | -0.89 |
| 1069832 | SKIP_ON | ENSSSCG00000033093 | 7 | 95904553 | 95904606 | 0.38 | 0.78 | 1.04 | 2.13 | 2.13 | 2.59 | 0.73 | 2.28 | 0.32 | -1.64 | 1.51 | 6.61 | 0.0017 | 0.053 | -0.78 |
| 1054358 | AE | ENSSSCG00000000449 | 5 | 22869184 | 22869861 | 1.22 | 2.13 | 3.43 | 6.69 | 7.34 | 7.15 | 2.26 | 7.06 | 0.32 | -1.64 | 4.66 | 7.60 | 0.0010 | 0.042 | -0.13 |
| 1054350 | TSS | ENSSSCG00000000449 | 5 | 22868430 | 22868453 | 1.22 | 2.13 | 3.43 | 6.69 | 7.34 | 7.15 | 2.26 | 7.06 | 0.32 | -1.64 | 4.66 | 7.60 | 0.0010 | 0.042 | -0.13 |
| 1054355 | XIR_ON | ENSSSCG00000000449 | 5 | 22869411 | 22869801 | 1.22 | 2.13 | 3.43 | 6.69 | 7.34 | 7.15 | 2.26 | 7.06 | 0.32 | -1.64 | 4.66 | 7.60 | 0.0010 | 0.042 | -0.13 |
| 1001083 | SKIP_ON | ENSSSCG00000004371 | 1 | 72747720 | 72747752 | 0.85 | 1.04 | 1.35 | 3.81 | 3.10 | 3.22 | 1.08 | 3.38 | 0.32 | -1.65 | 2.23 | 9.12 | 0.0004 | 0.031 | 0.74 |
| 1039952 | TSS | ENSSSCG00000014348 | 2 | 141293051 | 141293194 | 3.89 | 3.06 | 3.41 | 10.57 | 9.12 | 12.87 | 3.45 | 10.85 | 0.32 | -1.65 | 7.15 | 7.04 | 0.0013 | 0.048 | -0.48 |
| 1044200 | TSS | ENSSSCG00000007932 | 3 | 37591842 | 37593729 | 4.81 | 3.57 | 5.32 | 15.94 | 11.87 | 15.32 | 4.57 | 14.38 | 0.32 | -1.65 | 9.47 | 7.61 | 0.0010 | 0.042 | -0.12 |
| 1060782 | TSS | ENSSSCG00000003828 | 6 | 153197599 | 153197756 | 1.72 | 1.82 | 1.85 | 6.39 | 4.46 | 6.08 | 1.79 | 5.64 | 0.32 | -1.65 | 3.72 | 6.81 | 0.0015 | 0.050 | -0.64 |
| 1039141 | TSS | ENSSSCG00000014056 | 2 | 81372439 | 81372674 | 0.81 | 0.96 | 1.32 | 3.33 | 2.89 | 3.53 | 1.03 | 3.25 | 0.32 | -1.66 | 2.14 | 9.60 | 0.0004 | 0.029 | 0.98 |
| 1077091 | TSS | ENSSSCG00000029649 | 9 | 45861694 | 45861923 | 31.31 | 34.00 | 28.07 | 85.54 | 91.84 | 117.61 | 31.13 | 98.33 | 0.32 | -1.66 | 64.73 | 7.17 | 0.0012 | 0.047 | -0.40 |
| 1077092 | TTS | ENSSSCG00000029649 | 9 | 45844619 | 45845264 | 31.31 | 34.00 | 28.07 | 85.54 | 91.84 | 117.61 | 31.13 | 98.33 | 0.32 | -1.66 | 64.73 | 7.17 | 0.0012 | 0.047 | -0.40 |
| 1040317 | TSS | ENSSSCG00000021082 | 2 | 141823828 | 141824530 | 0.25 | 0.25 | 0.21 | 0.75 | 0.69 | 0.80 | 0.24 | 0.75 | 0.32 | -1.66 | 0.49 | 10.03 | 0.0003 | 0.027 | 1.18 |
| 1040318 | TTS | ENSSSCG00000021082 | 2 | 141662775 | 141663149 | 0.25 | 0.25 | 0.21 | 0.75 | 0.69 | 0.80 | 0.24 | 0.75 | 0.32 | -1.66 | 0.49 | 10.03 | 0.0003 | 0.027 | 1.18 |
| 1014970 | TSS | ENSSSCG00000039424 | 12 | 1141096 | 1142937 | 16.61 | 9.71 | 13.69 | 39.22 | 44.65 | 42.60 | 13.34 | 42.15 | 0.32 | -1.66 | 27.75 | 12.00 | 0.0001 | 0.020 | 2.00 |
| 1066098 | TSS | ENSSSCG00000001203 | 7 | 22082220 | 22082282 | 1.63 | 0.77 | 1.23 | 3.75 | 3.69 | 4.06 | 1.21 | 3.83 | 0.32 | -1.66 | 2.52 | 10.12 | 0.0003 | 0.026 | 1.22 |
| 1066099 | TTS | ENSSSCG00000001203 | 7 | 22087495 | 22088469 | 1.63 | 0.77 | 1.23 | 3.75 | 3.69 | 4.06 | 1.21 | 3.83 | 0.32 | -1.66 | 2.52 | 10.12 | 0.0003 | 0.026 | 1.22 |
| 1070467 | SKIP_OFF | ENSSSCG00000038886 | 7 | 4721548 | 4721658 | 0.56 | 0.73 | 0.76 | 2.48 | 2.20 | 1.82 | 0.68 | 2.17 | 0.32 | -1.67 | 1.43 | 7.62 | 0.0010 | 0.042 | -0.11 |
| 1070463 | TSS | ENSSSCG00000038886 | 7 | 4750286 | 4750752 | 0.56 | 0.73 | 0.76 | 2.48 | 2.20 | 1.82 | 0.68 | 2.17 | 0.32 | -1.67 | 1.43 | 7.62 | 0.0010 | 0.042 | -0.11 |
| 1070470 | XIR_ON | ENSSSCG00000038886 | 7 | 4746693 | 4746736 | 0.56 | 0.73 | 0.76 | 2.48 | 2.20 | 1.82 | 0.68 | 2.17 | 0.32 | -1.67 | 1.43 | 7.62 | 0.0010 | 0.042 | -0.11 |
| 1073503 | TSS | ENSSSCG00000038080 | 8 | 120145621 | 120145879 | 9.03 | 8.29 | 9.17 | 26.75 | 27.16 | 30.11 | 8.83 | 28.01 | 0.32 | -1.67 | 18.42 | 18.61 | 0.0000 | 0.010 | 3.81 |
| 1073504 | TTS | ENSSSCG00000038080 | 8 | 120226377 | 120229307 | 9.03 | 8.29 | 9.17 | 26.75 | 27.16 | 30.11 | 8.83 | 28.01 | 0.32 | -1.67 | 18.42 | 18.61 | 0.0000 | 0.010 | 3.81 |
| 1002726 | TSS | ENSSSCG00000004919 | 1 | 162512444 | 162513718 | 1.54 | 0.75 | 1.34 | 3.96 | 3.22 | 4.32 | 1.21 | 3.84 | 0.31 | -1.67 | 2.52 | 6.94 | 0.0014 | 0.049 | -0.55 |
| 1048426 | TSS | ENSSSCG00000038001 | 3 | 111327821 | 111328868 | 2.18 | 2.70 | 2.36 | 7.51 | 6.92 | 8.58 | 2.41 | 7.67 | 0.31 | -1.67 | 5.04 | 10.91 | 0.0002 | 0.023 | 1.56 |
| 1059697 | SKIP_ON | ENSSSCG00000003486 | 6 | 76150473 | 76150589 | 0.18 | 0.13 | 0.13 | 0.45 | 0.52 | 0.43 | 0.15 | 0.47 | 0.31 | -1.67 | 0.31 | 6.55 | 0.0018 | 0.054 | -0.83 |
| 1015910 | SKIP_ON | ENSSSCG00000011393 | 13 | 32334870 | 32335061 | 2.93 | 2.02 | 3.46 | 10.42 | 8.44 | 7.92 | 2.80 | 8.92 | 0.31 | -1.67 | 5.86 | 7.47 | 0.0010 | 0.043 | -0.20 |
| 1050025 | TTS | ENSSSCG00000006294 | 4 | 81911463 | 81911495 | 8.27 | 8.37 | 7.69 | 25.89 | 26.95 | 24.78 | 8.11 | 25.87 | 0.31 | -1.67 | 16.99 | 28.45 | 0.0000 | 0.009 | 5.17 |
| 1050087 | TTS | ENSSSCG00000006309 | 4 | 83583375 | 83586639 | 1.86 | 1.13 | 1.86 | 4.87 | 5.79 | 4.92 | 1.62 | 5.19 | 0.31 | -1.68 | 3.41 | 9.78 | 0.0003 | 0.028 | 1.06 |
| 1066434 | SKIP_OFF | ENSSSCG00000001476 | 7 | 25249107 | 25249149 | 0.64 | 1.04 | 0.59 | 2.53 | 2.20 | 2.57 | 0.76 | 2.43 | 0.31 | -1.69 | 1.59 | 9.40 | 0.0004 | 0.030 | 0.88 |
| 1000001 | TSS | ENSSSCG00000004008 | 1 | 578465 | 578588 | 1.05 | 0.53 | 0.96 | 2.81 | 2.83 | 2.55 | 0.85 | 2.73 | 0.31 | -1.69 | 1.79 | 10.49 | 0.0002 | 0.025 | 1.39 |
| 1000002 | TTS | ENSSSCG00000004008 | 1 | 719872 | 719928 | 1.05 | 0.53 | 0.96 | 2.81 | 2.83 | 2.55 | 0.85 | 2.73 | 0.31 | -1.69 | 1.79 | 10.49 | 0.0002 | 0.025 | 1.39 |
| 1022432 | SKIP_OFF | ENSSSCG00000009956 | 14 | 43436685 | 43436720 | 6.82 | 7.32 | 6.50 | 22.29 | 19.05 | 25.37 | 6.88 | 22.24 | 0.31 | -1.69 | 14.56 | 8.87 | 0.0005 | 0.033 | 0.61 |
| 1077969 | TTS | ENSSSCG00000038710 | 9 | 45236741 | 45240812 | 0.14 | 0.22 | 0.18 | 0.63 | 0.51 | 0.59 | 0.18 | 0.58 | 0.31 | -1.69 | 0.38 | 7.39 | 0.0011 | 0.045 | -0.26 |
| 1076592 | TTS | ENSSSCG00000023451 | 9 | 128429092 | 128432727 | 0.98 | 1.19 | 0.74 | 2.67 | 3.63 | 3.15 | 0.97 | 3.15 | 0.31 | -1.70 | 2.06 | 7.50 | 0.0010 | 0.043 | -0.18 |
| 1022430 | TTS | ENSSSCG00000009956 | 14 | 43438709 | 43438870 | 6.82 | 7.32 | 6.50 | 22.39 | 19.21 | 25.49 | 6.88 | 22.36 | 0.31 | -1.70 | 14.62 | 9.00 | 0.0005 | 0.032 | 0.67 |
| 1022428 | TSS | ENSSSCG00000009956 | 14 | 43304626 | 43305485 | 6.82 | 7.32 | 6.50 | 22.39 | 19.21 | 25.49 | 6.88 | 22.36 | 0.31 | -1.70 | 14.62 | 9.00 | 0.0005 | 0.032 | 0.67 |
| 1015904 | XMSKIP_ON | ENSSSCG00000011393 | 13 | 32340609 | 32341006 | 3.06 | 1.72 | 3.57 | 10.72 | 8.51 | 7.96 | 2.78 | 9.06 | 0.31 | -1.70 | 5.92 | 6.61 | 0.0018 | 0.053 | -0.78 |
| 1003706 | TSS | ENSSSCG00000005266 | 1 | 225951643 | 225951810 | 0.55 | 0.43 | 0.56 | 1.67 | 1.96 | 1.37 | 0.51 | 1.67 | 0.31 | -1.70 | 1.09 | 6.79 | 0.0016 | 0.051 | -0.65 |
| 1003707 | TTS | ENSSSCG00000005266 | 1 | 226109327 | 226111095 | 0.55 | 0.43 | 0.56 | 1.67 | 1.96 | 1.37 | 0.51 | 1.67 | 0.31 | -1.70 | 1.09 | 6.79 | 0.0016 | 0.051 | -0.65 |
| 1034993 | TSS | ENSSSCG00000016690 | 18 | 43975913 | 43976006 | 0.51 | 0.89 | 0.80 | 1.92 | 2.71 | 2.54 | 0.73 | 2.39 | 0.31 | -1.70 | 1.56 | 6.50 | 0.0019 | 0.054 | -0.86 |
| 1044298 | SKIP_ON | ENSSSCG00000007955 | 3 | 38750963 | 38751074 | 1.90 | 1.21 | 1.32 | 5.22 | 4.23 | 4.98 | 1.48 | 4.81 | 0.31 | -1.70 | 3.14 | 9.61 | 0.0004 | 0.029 | 0.98 |
| 1009251 | TSS | ENSSSCG00000009270 | 11 | 943456 | 943592 | 11.05 | 7.26 | 8.54 | 34.29 | 24.31 | 28.88 | 8.95 | 29.16 | 0.31 | -1.70 | 19.05 | 6.95 | 0.0014 | 0.049 | -0.55 |
| 1039443 | TSS | ENSSSCG00000014161 | 2 | 101291853 | 101292482 | 1.35 | 1.09 | 1.34 | 4.56 | 4.58 | 3.20 | 1.26 | 4.11 | 0.31 | -1.71 | 2.69 | 6.49 | 0.0019 | 0.055 | -0.87 |
| 1039444 | TTS | ENSSSCG00000014161 | 2 | 100937214 | 100937379 | 1.35 | 1.09 | 1.34 | 4.56 | 4.58 | 3.20 | 1.26 | 4.11 | 0.31 | -1.71 | 2.69 | 6.49 | 0.0019 | 0.055 | -0.87 |
| 1032057 | MSKIP_OFF | ENSSSCG00000007094 | 17 | 26652875 | 26654116 | 2.03 | 0.81 | 1.87 | 5.75 | 5.14 | 4.50 | 1.57 | 5.13 | 0.31 | -1.71 | 3.35 | 7.15 | 0.0013 | 0.047 | -0.41 |
| 1032046 | SKIP_OFF | ENSSSCG00000007094 | 17 | 26652875 | 26654116 | 2.03 | 0.81 | 1.87 | 5.75 | 5.14 | 4.50 | 1.57 | 5.13 | 0.31 | -1.71 | 3.35 | 7.15 | 0.0013 | 0.047 | -0.41 |
| 1032054 | XSKIP_OFF | ENSSSCG00000007094 | 17 | 26652875 | 26654816 | 2.03 | 0.81 | 1.87 | 5.75 | 5.14 | 4.50 | 1.57 | 5.13 | 0.31 | -1.71 | 3.35 | 7.15 | 0.0013 | 0.047 | -0.41 |
| 1043415 | XSKIP_ON | ENSSSCG00000007568 | 3 | 1867257 | 1867357 | 4.25 | 2.43 | 2.78 | 11.24 | 8.54 | 11.15 | 3.15 | 10.31 | 0.31 | -1.71 | 6.73 | 7.25 | 0.0012 | 0.046 | -0.35 |
| 1043417 | XSKIP_ON | ENSSSCG00000007568 | 3 | 1875787 | 1875877 | 4.25 | 2.43 | 2.78 | 11.24 | 8.54 | 11.15 | 3.15 | 10.31 | 0.31 | -1.71 | 6.73 | 7.25 | 0.0012 | 0.046 | -0.35 |
| 1043419 | XSKIP_ON | ENSSSCG00000007568 | 3 | 1884171 | 1884220 | 4.25 | 2.43 | 2.78 | 11.24 | 8.54 | 11.15 | 3.15 | 10.31 | 0.31 | -1.71 | 6.73 | 7.25 | 0.0012 | 0.046 | -0.35 |
| 1001081 | TSS | ENSSSCG00000004371 | 1 | 72683320 | 72683322 | 0.81 | 1.04 | 1.25 | 3.81 | 3.10 | 3.22 | 1.03 | 3.38 | 0.31 | -1.71 | 2.20 | 9.70 | 0.0003 | 0.028 | 1.02 |
| 1001957 | SKIP_ON | ENSSSCG00000004651 | 1 | 122444378 | 122444476 | 7.51 | 5.99 | 5.66 | 18.19 | 20.28 | 24.49 | 6.39 | 20.99 | 0.30 | -1.72 | 13.69 | 8.00 | 0.0008 | 0.039 | 0.12 |
| 1078719 | TSS | ENSSSCG00000012295 | X | 43207054 | 43207146 | 2.15 | 1.59 | 2.14 | 6.93 | 5.46 | 6.95 | 1.96 | 6.45 | 0.30 | -1.72 | 4.20 | 9.00 | 0.0005 | 0.032 | 0.68 |

| | | | | | | | | | | | | | | | | | | | | |
|---|---|---|---|---|---|---|---|---|---|---|---|---|---|---|---|---|---|---|---|---|
| 1078720 | TTS | ENSSSCG00000012295 | X | 43211102 | 43211626 | 2.15 | 1.59 | 2.14 | 6.93 | 5.46 | 6.95 | 1.96 | 6.45 | 0.30 | -1.72 | 4.20 | 9.00 | 0.0005 | 0.032 | 0.68 |
| 1001952 | TSS | ENSSSCG00000004651 | 1 | 122526534 | 122526981 | 7.51 | 5.99 | 5.66 | 18.30 | 20.28 | 24.49 | 6.39 | 21.02 | 0.30 | -1.72 | 13.70 | 8.13 | 0.0007 | 0.038 | 0.20 |
| 1049633 | TSS | ENSSSCG00000006161 | 4 | 57684704 | 57685354 | 0.28 | 0.65 | 0.39 | 1.43 | 1.30 | 1.60 | 0.44 | 1.45 | 0.30 | -1.72 | 0.94 | 7.29 | 0.0012 | 0.045 | -0.32 |
| 1049634 | TTS | ENSSSCG00000006161 | 4 | 57743673 | 57744709 | 0.28 | 0.65 | 0.39 | 1.43 | 1.30 | 1.60 | 0.44 | 1.45 | 0.30 | -1.72 | 0.94 | 7.29 | 0.0012 | 0.045 | -0.32 |
| 1063362 | TSS | ENSSSCG00000031652 | 6 | 31564986 | 31565042 | 2.64 | 1.92 | 2.45 | 8.52 | 7.97 | 6.57 | 2.33 | 7.69 | 0.30 | -1.72 | 5.01 | 9.15 | 0.0004 | 0.031 | 0.75 |
| 1063363 | TTS | ENSSSCG00000031652 | 6 | 31658230 | 31661190 | 2.64 | 1.92 | 2.45 | 8.52 | 7.97 | 6.57 | 2.33 | 7.69 | 0.30 | -1.72 | 5.01 | 9.15 | 0.0004 | 0.031 | 0.75 |
| 1063364 | SKIP_ON | ENSSSCG00000031652 | 6 | 31622310 | 31622390 | 2.64 | 1.92 | 2.45 | 8.52 | 7.97 | 6.57 | 2.33 | 7.69 | 0.30 | -1.72 | 5.01 | 9.15 | 0.0004 | 0.031 | 0.75 |
| 1044682 | TSS | ENSSSCG00000008089 | 3 | 43750687 | 43750795 | 1.05 | 0.59 | 1.33 | 2.79 | 3.25 | 3.73 | 0.99 | 3.26 | 0.30 | -1.72 | 2.12 | 6.88 | 0.0015 | 0.050 | -0.59 |
| 1044683 | TTS | ENSSSCG00000008089 | 3 | 43771701 | 43771926 | 1.05 | 0.59 | 1.33 | 2.79 | 3.25 | 3.73 | 0.99 | 3.26 | 0.30 | -1.72 | 2.12 | 6.88 | 0.0015 | 0.050 | -0.59 |
| 1002144 | SKIP_ON | ENSSSCG00000004704 | 1 | 127886234 | 127886383 | 1.01 | 1.30 | 1.88 | 5.12 | 4.43 | 4.23 | 1.39 | 4.59 | 0.30 | -1.72 | 2.99 | 9.10 | 0.0004 | 0.032 | 0.73 |
| 1066398 | TTS | ENSSSCG00000001459 | 7 | 25029182 | 25033067 | 0.24 | 0.28 | 0.33 | 1.03 | 0.88 | 0.90 | 0.28 | 0.94 | 0.30 | -1.72 | 0.61 | 10.20 | 0.0003 | 0.026 | 1.26 |
| 1002151 | SKIP_OFF | ENSSSCG00000004704 | 1 | 127894725 | 127894907 | 1.01 | 1.30 | 1.88 | 5.12 | 4.48 | 4.23 | 1.39 | 4.61 | 0.30 | -1.73 | 3.00 | 9.24 | 0.0004 | 0.031 | 0.80 |
| 1002146 | SKIP_ON | ENSSSCG00000004704 | 1 | 127890879 | 127890941 | 1.01 | 1.30 | 1.88 | 5.12 | 4.48 | 4.23 | 1.39 | 4.61 | 0.30 | -1.73 | 3.00 | 9.24 | 0.0004 | 0.031 | 0.80 |
| 1004123 | SKIP_ON | ENSSSCG00000005426 | 1 | 246833278 | 246833310 | 2.00 | 1.77 | 1.50 | 6.63 | 5.52 | 5.32 | 1.76 | 5.83 | 0.30 | -1.73 | 3.79 | 9.94 | 0.0003 | 0.027 | 1.14 |
| 1004125 | XSKIP_ON | ENSSSCG00000005426 | 1 | 246833278 | 246833310 | 2.00 | 1.77 | 1.50 | 6.63 | 5.52 | 5.32 | 1.76 | 5.83 | 0.30 | -1.73 | 3.79 | 9.94 | 0.0003 | 0.027 | 1.14 |
| 1056118 | TSS | ENSSSCG00000023788 | 5 | 29205704 | 29206656 | 0.37 | 0.48 | 0.70 | 1.61 | 1.63 | 1.90 | 0.52 | 1.71 | 0.30 | -1.73 | 1.12 | 8.88 | 0.0005 | 0.033 | 0.61 |
| 1033550 | TTS | ENSSSCG00000031228 | 17 | 62911874 | 62913752 | 0.36 | 0.21 | 0.21 | 0.85 | 0.88 | 0.86 | 0.26 | 0.86 | 0.30 | -1.73 | 0.56 | 10.05 | 0.0003 | 0.027 | 1.19 |
| 1021559 | TSS | ENSSSCG00000009699 | 14 | 15718522 | 15718821 | 0.54 | 0.40 | 0.63 | 1.42 | 1.90 | 1.89 | 0.52 | 1.74 | 0.30 | -1.73 | 1.13 | 7.38 | 0.0011 | 0.045 | -0.26 |
| 1021560 | TTS | ENSSSCG00000009699 | 14 | 15752844 | 15755457 | 0.54 | 0.40 | 0.63 | 1.42 | 1.90 | 1.89 | 0.52 | 1.74 | 0.30 | -1.73 | 1.13 | 7.38 | 0.0011 | 0.045 | -0.26 |
| 1039473 | SKIP_ON | ENSSSCG00000014170 | 2 | 103270394 | 103270456 | 1.05 | 1.34 | 1.39 | 3.71 | 4.51 | 4.34 | 1.26 | 4.19 | 0.30 | -1.73 | 2.72 | 11.54 | 0.0002 | 0.022 | 1.82 |
| 1050889 | SKIP_ON | ENSSSCG00000006620 | 4 | 97676065 | 97676139 | 1.18 | 0.59 | 1.32 | 3.49 | 2.84 | 3.92 | 1.03 | 3.42 | 0.30 | -1.73 | 2.22 | 6.57 | 0.0018 | 0.054 | -0.81 |
| 1021822 | TTS | ENSSSCG00000009783 | 14 | 29890066 | 29890329 | 1.93 | 1.49 | 2.52 | 6.71 | 5.81 | 7.25 | 1.98 | 6.59 | 0.30 | -1.74 | 4.28 | 9.48 | 0.0004 | 0.029 | 0.92 |
| 1021828 | XIR_OFF | ENSSSCG00000009783 | 14 | 29890329 | 29890404 | 1.93 | 1.49 | 2.52 | 6.71 | 5.81 | 7.25 | 1.98 | 6.59 | 0.30 | -1.74 | 4.28 | 9.48 | 0.0004 | 0.029 | 0.92 |
| 1050026 | SKIP_ON | ENSSSCG00000006294 | 4 | 81769768 | 81769869 | 8.10 | 7.30 | 6.65 | 24.91 | 24.86 | 23.85 | 7.35 | 24.54 | 0.30 | -1.74 | 15.94 | 33.36 | 0.0000 | 0.009 | 5.55 |
| 1050028 | SKIP_ON | ENSSSCG00000006294 | 4 | 81756380 | 81756448 | 8.10 | 7.30 | 6.65 | 24.91 | 24.86 | 23.85 | 7.35 | 24.54 | 0.30 | -1.74 | 15.94 | 33.36 | 0.0000 | 0.009 | 5.55 |
| 1064453 | SKIP_ON | ENSSSCG00000035849 | 6 | 167680823 | 167681128 | 19.25 | 16.77 | 12.45 | 55.75 | 44.43 | 61.87 | 16.16 | 54.02 | 0.30 | -1.74 | 35.09 | 7.34 | 0.0011 | 0.045 | -0.29 |
| 1067233 | SKIP_OFF | ENSSSCG00000001779 | 7 | 49010454 | 49010575 | 0.75 | 0.55 | 0.57 | 1.78 | 2.43 | 2.07 | 0.63 | 2.09 | 0.30 | -1.74 | 1.36 | 7.58 | 0.0010 | 0.043 | -0.13 |
| 1067235 | XSKIP_OFF | ENSSSCG00000001779 | 7 | 49010454 | 49010575 | 0.75 | 0.55 | 0.57 | 1.78 | 2.43 | 2.07 | 0.63 | 2.09 | 0.30 | -1.74 | 1.36 | 7.58 | 0.0010 | 0.043 | -0.13 |
| 1063366 | SKIP_ON | ENSSSCG00000031652 | 6 | 31569558 | 31569702 | 2.64 | 1.81 | 2.44 | 8.52 | 7.97 | 6.57 | 2.29 | 7.69 | 0.30 | -1.74 | 4.99 | 9.03 | 0.0005 | 0.032 | 0.69 |
| 1012505 | TTS | ENSSSCG00000017747 | 12 | 43362342 | 43367139 | 0.47 | 0.36 | 0.26 | 1.05 | 1.16 | 1.45 | 0.36 | 1.22 | 0.30 | -1.74 | 0.79 | 6.54 | 0.0018 | 0.054 | -0.83 |
| 1067425 | SKIP_ON | ENSSSCG00000001837 | 7 | 54885999 | 54886070 | 3.48 | 1.69 | 2.05 | 7.22 | 7.42 | 9.56 | 2.41 | 8.06 | 0.30 | -1.75 | 5.24 | 6.47 | 0.0019 | 0.055 | -0.88 |
| 1060733 | IR_ON | ENSSSCG00000003814 | 6 | 148940163 | 148942244 | 1.52 | 1.16 | 1.96 | 6.05 | 5.19 | 4.34 | 1.55 | 5.19 | 0.30 | -1.75 | 3.37 | 7.08 | 0.0013 | 0.047 | -0.46 |
| 1060645 | TSS | ENSSSCG00000003784 | 6 | 138950089 | 138950599 | 0.33 | 0.28 | 0.44 | 0.95 | 1.35 | 1.20 | 0.35 | 1.17 | 0.30 | -1.75 | 0.76 | 6.59 | 0.0018 | 0.053 | -0.80 |
| 1066395 | TSS | ENSSSCG00000001459 | 7 | 25037763 | 25037892 | 1.86 | 1.21 | 1.38 | 5.47 | 4.73 | 4.78 | 1.48 | 4.99 | 0.30 | -1.75 | 3.24 | 11.97 | 0.0001 | 0.020 | 1.99 |
| 1029785 | TSS | ENSSSCG00000034567 | 15 | 68107090 | 68107339 | 0.86 | 1.42 | 1.02 | 3.14 | 3.64 | 4.32 | 1.10 | 3.70 | 0.30 | -1.75 | 2.40 | 7.23 | 0.0012 | 0.046 | -0.36 |
| 1029786 | TTS | ENSSSCG00000034567 | 15 | 68057757 | 68059443 | 0.86 | 1.42 | 1.02 | 3.14 | 3.64 | 4.32 | 1.10 | 3.70 | 0.30 | -1.75 | 2.40 | 7.23 | 0.0012 | 0.046 | -0.36 |
| 1000420 | TTS | ENSSSCG00000004149 | 1 | 25985796 | 25986510 | 2.67 | 3.31 | 5.17 | 12.32 | 10.97 | 14.30 | 3.72 | 12.53 | 0.30 | -1.75 | 8.12 | 7.65 | 0.0009 | 0.042 | -0.09 |
| 1028292 | AE | ENSSSCG00000016164 | 15 | 115572770 | 115573036 | 0.79 | 0.53 | 0.79 | 2.80 | 2.40 | 1.94 | 0.71 | 2.38 | 0.30 | -1.75 | 1.54 | 6.72 | 0.0016 | 0.052 | -0.71 |
| 1063287 | TSS | ENSSSCG00000031300 | 6 | 163066161 | 163066663 | 0.38 | 0.27 | 0.24 | 1.01 | 1.02 | 0.98 | 0.30 | 1.00 | 0.30 | -1.75 | 0.65 | 12.79 | 0.0001 | 0.019 | 2.28 |
| 1064254 | TSS | ENSSSCG00000035104 | 6 | 27489130 | 27489238 | 0.56 | 0.41 | 0.34 | 1.43 | 1.42 | 1.59 | 0.44 | 1.48 | 0.30 | -1.76 | 0.96 | 11.63 | 0.0002 | 0.021 | 1.86 |
| 1064255 | TTS | ENSSSCG00000035104 | 6 | 27451465 | 27451652 | 0.56 | 0.41 | 0.34 | 1.43 | 1.42 | 1.59 | 0.44 | 1.48 | 0.30 | -1.76 | 0.96 | 11.63 | 0.0002 | 0.021 | 1.86 |
| 1001953 | TTS | ENSSSCG00000004651 | 1 | 122370623 | 122374061 | 7.15 | 5.99 | 5.45 | 18.30 | 20.28 | 24.24 | 6.20 | 20.94 | 0.30 | -1.76 | 13.57 | 8.61 | 0.0006 | 0.035 | 0.47 |
| 1027848 | TSS | ENSSSCG00000016052 | 15 | 95015284 | 95015635 | 1.69 | 0.59 | 1.02 | 4.18 | 3.70 | 3.30 | 1.10 | 3.73 | 0.30 | -1.76 | 2.41 | 6.78 | 0.0016 | 0.051 | -0.66 |
| 1050024 | TSS | ENSSSCG00000006294 | 4 | 81640178 | 81640307 | 7.37 | 6.75 | 5.98 | 22.67 | 23.36 | 21.92 | 6.70 | 22.65 | 0.30 | -1.76 | 14.67 | 29.16 | 0.0000 | 0.009 | 5.23 |
| 1060785 | TTS | ENSSSCG00000003828 | 6 | 152754730 | 152755149 | 2.81 | 3.12 | 2.32 | 10.18 | 7.57 | 10.27 | 2.75 | 9.34 | 0.29 | -1.76 | 6.05 | 7.65 | 0.0009 | 0.042 | -0.09 |
| 1043412 | TTS | ENSSSCG00000007568 | 3 | 1861492 | 1864621 | 3.20 | 2.43 | 2.78 | 11.08 | 7.70 | 9.92 | 2.80 | 9.57 | 0.29 | -1.77 | 6.18 | 7.06 | 0.0013 | 0.047 | -0.47 |
| 1039447 | SKIP_ON | ENSSSCG00000014161 | 2 | 101194422 | 101194583 | 1.17 | 1.09 | 1.34 | 4.56 | 4.58 | 3.20 | 1.20 | 4.11 | 0.29 | -1.78 | 2.66 | 6.65 | 0.0017 | 0.053 | -0.75 |
| 1030340 | TSS | ENSSSCG00000016782 | 16 | 3969915 | 3970084 | 3.82 | 3.73 | 5.73 | 16.09 | 13.98 | 15.42 | 4.42 | 15.16 | 0.29 | -1.78 | 9.79 | 12.65 | 0.0001 | 0.019 | 2.23 |
| 1033548 | TSS | ENSSSCG00000031228 | 17 | 62905780 | 62906298 | 0.33 | 0.21 | 0.21 | 0.85 | 0.88 | 0.86 | 0.25 | 0.86 | 0.29 | -1.78 | 0.56 | 11.42 | 0.0002 | 0.022 | 1.78 |
| 1020056 | TSS | ENSSSCG00000033000 | 13 | 68190712 | 68190982 | 1.55 | 2.31 | 1.80 | 7.42 | 6.07 | 6.02 | 1.89 | 6.51 | 0.29 | -1.79 | 4.20 | 9.59 | 0.0004 | 0.029 | 0.97 |
| 1020058 | TTS | ENSSSCG00000033000 | 13 | 68179806 | 68184769 | 1.55 | 2.31 | 1.80 | 7.42 | 6.07 | 6.02 | 1.89 | 6.51 | 0.29 | -1.79 | 4.20 | 9.59 | 0.0004 | 0.029 | 0.97 |
| 1070110 | TSS | ENSSSCG00000035596 | 7 | 23852615 | 23852971 | 1.11 | 0.67 | 0.89 | 3.09 | 3.27 | 2.87 | 0.89 | 3.08 | 0.29 | -1.79 | 1.98 | 13.19 | 0.0001 | 0.018 | 2.42 |
| 1070111 | TTS | ENSSSCG00000035596 | 7 | 23872509 | 23872620 | 1.11 | 0.67 | 0.89 | 3.09 | 3.27 | 2.87 | 0.89 | 3.08 | 0.29 | -1.79 | 1.98 | 13.19 | 0.0001 | 0.018 | 2.42 |
| 1064443 | TSS | ENSSSCG00000035849 | 6 | 167743530 | 167743708 | 19.25 | 16.77 | 12.81 | 60.54 | 45.26 | 62.77 | 16.28 | 56.19 | 0.29 | -1.79 | 36.23 | 7.29 | 0.0012 | 0.045 | -0.32 |

### Table S5 The most significantly upregulated and downregulated AS events

| data...1. | event_type | gene_id | chrom | event_start | event_end | XL13 | XL17 | XL18 | XS12 | XS15 | XS16 | XL AveFPKM | XS AveFPKM | FC | logFC | AveExpr | t | P.Value | adj.P.Val | B |
|---|---|---|---|---|---|---|---|---|---|---|---|---|---|---|---|---|---|---|---|---|
| 1037279 | SKIP_OFF | ENSSSCG00000013382 | 2 | 42257144 | 42257614 | 0.00 | 0.00 | 0.00 | 0.47 | 0.78 | 0.59 | 0.00 | 0.61 | 0.00 | -22.93 | 0.31 | 6.57 | 0.002 | 0.05 | -0.81 |
| 1062231 | TTS | ENSSSCG00000025652 | 6 | 18075478 | 18075965 | 0.00 | 0.00 | 0.00 | 4.78 | 3.45 | 5.89 | 0.00 | 4.71 | 0.00 | -22.12 | 2.35 | 7.09 | 0.001 | 0.05 | -0.45 |
| 1062235 | XIR_OFF | ENSSSCG00000025652 | 6 | 18075965 | 18076256 | 0.00 | 0.00 | 0.00 | 4.78 | 3.45 | 5.89 | 0.00 | 4.71 | 0.00 | -22.12 | 2.35 | 7.09 | 0.001 | 0.05 | -0.45 |
| 1044201 | TSS | ENSSSCG00000007932 | 3 | 37594724 | 37594788 | 0.00 | 0.00 | 0.00 | 11.42 | 6.45 | 9.66 | 0.00 | 9.18 | 0.00 | -20.06 | 4.59 | 6.70 | 0.002 | 0.05 | -0.72 |
| 1031206 | SKIP_OFF | ENSSSCG00000021527 | 16 | 62909457 | 62909644 | 0.00 | 0.00 | 0.00 | 0.86 | 0.78 | 1.02 | 0.00 | 0.89 | 0.00 | -18.57 | 0.44 | 11.42 | 0.000 | 0.02 | 1.77 |
| 1069930 | TSS | ENSSSCG00000033721 | 7 | 76489466 | 76489520 | 0.00 | 0.00 | 0.00 | 5.13 | 4.04 | 5.46 | 0.00 | 4.87 | 0.00 | -17.76 | 2.44 | 12.01 | 0.000 | 0.02 | 2.00 |
| 1037695 | SKIP_OFF | ENSSSCG00000013506 | 2 | 74469449 | 74469532 | 0.00 | 0.00 | 0.00 | 0.42 | 0.52 | 0.63 | 0.00 | 0.52 | 0.00 | -17.63 | 0.26 | 7.52 | 0.001 | 0.04 | -0.17 |
| 1037692 | SKIP_ON | ENSSSCG00000013506 | 2 | 74465870 | 74465947 | 0.00 | 0.00 | 0.00 | 0.42 | 0.52 | 0.63 | 0.00 | 0.52 | 0.00 | -17.47 | 0.26 | 7.52 | 0.001 | 0.04 | -0.17 |
| 1052716 | TSS | ENSSSCG00000034242 | 4 | 60637049 | 60637238 | 0.00 | 0.00 | 0.00 | 1.74 | 1.16 | 1.43 | 0.00 | 1.44 | 0.00 | -17.45 | 0.72 | 8.87 | 0.001 | 0.03 | 0.61 |
| 1074745 | TSS | ENSSSCG00000015089 | 9 | 45535018 | 45535259 | 0.00 | 0.00 | 0.00 | 1.38 | 0.90 | 1.38 | 0.00 | 1.22 | 0.00 | -16.61 | 0.61 | 7.80 | 0.001 | 0.04 | 0.00 |
| 1074753 | XSKIP_ON | ENSSSCG00000015089 | 9 | 45534751 | 45534780 | 0.00 | 0.00 | 0.00 | 1.38 | 0.90 | 1.38 | 0.00 | 1.22 | 0.00 | -16.61 | 0.61 | 7.80 | 0.001 | 0.04 | 0.00 |
| 1065341 | TSS | ENSSSCG00000039678 | 6 | 168651154 | 168651178 | 0.00 | 0.00 | 0.00 | 0.69 | 1.01 | 0.66 | 0.00 | 0.79 | 0.00 | -16.39 | 0.39 | 7.09 | 0.001 | 0.05 | -0.45 |
| 1072030 | TSS | ENSSSCG00000009169 | 8 | 118605293 | 118605600 | 0.00 | 0.00 | 0.00 | 0.97 | 0.77 | 0.64 | 0.00 | 0.79 | 0.00 | -15.52 | 0.40 | 8.08 | 0.001 | 0.04 | 0.17 |
| 1071119 | SKIP_OFF | ENSSSCG00000008838 | 8 | 40250019 | 40250153 | 0.00 | 0.00 | 0.00 | 1.77 | 1.00 | 1.65 | 0.00 | 1.47 | 0.00 | -14.71 | 0.74 | 6.49 | 0.002 | 0.05 | -0.87 |
| 1071114 | TSS | ENSSSCG00000008838 | 8 | 40369803 | 40369921 | 0.00 | 0.00 | 0.00 | 1.77 | 1.00 | 1.65 | 0.00 | 1.47 | 0.00 | -14.71 | 0.74 | 6.49 | 0.002 | 0.05 | -0.87 |
| 1056121 | SKIP_ON | ENSSSCG00000023788 | 5 | 29291627 | 29291686 | 0.00 | 0.00 | 0.00 | 0.81 | 0.70 | 0.90 | 0.00 | 0.80 | 0.00 | -14.43 | 0.40 | 12.32 | 0.000 | 0.02 | 2.11 |
| 1075337 | TSS | ENSSSCG00000015313 | 9 | 72185595 | 72185727 | 0.00 | 0.00 | 0.00 | 0.75 | 0.81 | 0.61 | 0.00 | 0.72 | 0.00 | -14.01 | 0.36 | 10.70 | 0.000 | 0.02 | 1.48 |
| 1075343 | XSKIP_ON | ENSSSCG00000015313 | 9 | 72184300 | 72184518 | 0.00 | 0.00 | 0.00 | 0.75 | 0.81 | 0.61 | 0.00 | 0.72 | 0.00 | -14.01 | 0.36 | 10.70 | 0.000 | 0.02 | 1.48 |
| 1037352 | SKIP_OFF | ENSSSCG00000013397 | 2 | 45928980 | 45929072 | 0.00 | 0.00 | 0.00 | 0.38 | 0.39 | 0.32 | 0.00 | 0.36 | 0.00 | -12.93 | 0.18 | 8.47 | 0.001 | 0.04 | 0.39 |
| 1072238 | MSKIP_OFF | ENSSSCG00000009237 | 8 | 135107334 | 135108713 | 0.00 | 0.00 | 0.00 | 0.25 | 0.29 | 0.25 | 0.00 | 0.27 | 0.00 | -12.67 | 0.13 | 6.60 | 0.002 | 0.05 | -0.79 |
| 1072235 | SKIP_ON | ENSSSCG00000009237 | 8 | 135091354 | 135091709 | 0.00 | 0.00 | 0.00 | 0.25 | 0.29 | 0.25 | 0.00 | 0.27 | 0.00 | -12.67 | 0.13 | 6.60 | 0.002 | 0.05 | -0.79 |
| 1051835 | MSKIP_OFF | ENSSSCG00000006950 | 4 | 130780652 | 130796097 | 0.01 | 0.00 | 0.00 | 10.84 | 7.64 | 13.67 | 0.00 | 10.72 | 0.00 | -12.51 | 5.36 | 6.53 | 0.002 | 0.05 | -0.84 |
| 1051830 | TTS | ENSSSCG00000006950 | 4 | 130775388 | 130775402 | 0.01 | 0.00 | 0.00 | 10.84 | 7.64 | 13.67 | 0.00 | 10.72 | 0.00 | -12.51 | 5.36 | 6.53 | 0.002 | 0.05 | -0.84 |
| 1045404 | TSS | ENSSSCG00000008368 | 3 | 78032364 | 78032476 | 0.00 | 0.00 | 0.00 | 0.45 | 0.45 | 0.46 | 0.00 | 0.46 | 0.00 | -12.19 | 0.23 | 11.83 | 0.000 | 0.02 | 1.93 |
| 1004079 | TSS | ENSSSCG00000005398 | 1 | 243039252 | 243039581 | 0.00 | 0.00 | 0.00 | 3.97 | 4.94 | 5.25 | 0.00 | 4.72 | 0.00 | -12.08 | 2.36 | 13.01 | 0.000 | 0.02 | 2.35 |
| 1055931 | AE | ENSSSCG00000000982 | 5 | 606408 | 606543 | 0.00 | 0.00 | 0.00 | 1.05 | 1.03 | 0.97 | 0.00 | 1.01 | 0.00 | -11.96 | 0.51 | 22.53 | 0.000 | 0.01 | 4.48 |
| 1055921 | TSS | ENSSSCG00000000982 | 5 | 599863 | 599923 | 0.00 | 0.00 | 0.00 | 1.05 | 1.03 | 0.97 | 0.00 | 1.01 | 0.00 | -11.96 | 0.51 | 22.53 | 0.000 | 0.01 | 4.48 |
| 1055928 | XIR_ON | ENSSSCG00000000982 | 5 | 629518 | 629546 | 0.00 | 0.00 | 0.00 | 1.05 | 1.03 | 0.97 | 0.00 | 1.01 | 0.00 | -11.96 | 0.51 | 22.53 | 0.000 | 0.01 | 4.48 |
| 1055926 | XSKIP_ON | ENSSSCG00000000982 | 5 | 632575 | 632605 | 0.00 | 0.00 | 0.00 | 1.05 | 1.03 | 0.97 | 0.00 | 1.01 | 0.00 | -11.96 | 0.51 | 22.53 | 0.000 | 0.01 | 4.48 |
| 1042711 | TSS | ENSSSCG00000037835 | 2 | 57310171 | 57311418 | 0.00 | 0.00 | 0.00 | 0.90 | 0.64 | 0.83 | 0.00 | 0.79 | 0.00 | -11.55 | 0.39 | 9.48 | 0.000 | 0.03 | 0.92 |

| | | | | | | | | | | | | | | | | | | | | |
|---|---|---|---|---|---|---|---|---|---|---|---|---|---|---|---|---|---|---|---|---|
| 1069242 | SKIP_ON | ENSSSCG00000025788 | 7 | 40906172 | 40906181 | 0.00 | 0.00 | 0.00 | 0.45 | 0.44 | 0.40 | 0.00 | 0.43 | 0.00 | -11.52 | 0.22 | 10.38 | 0.000 | 0.03 | 1.34 |
| 1011794 | XSKIP_ON | ENSSSCG00000017444 | 12 | 21129339 | 21129364 | 0.00 | 0.00 | 0.00 | 0.62 | 0.73 | 0.84 | 0.00 | 0.73 | 0.00 | -9.31 | 0.36 | 10.17 | 0.000 | 0.03 | 1.24 |
| 1023980 | TSS | ENSSSCG00000010433 | 14 | 99216806 | 99216885 | 0.00 | 0.00 | 0.00 | 1.41 | 1.16 | 1.83 | 0.00 | 1.47 | 0.00 | -8.95 | 0.74 | 7.75 | 0.001 | 0.04 | -0.03 |
| 1046634 | TTS | ENSSSCG00000023273 | 3 | 132730306 | 132730998 | 0.05 | 0.00 | 0.00 | 5.21 | 3.96 | 5.89 | 0.02 | 5.02 | 0.00 | -8.25 | 2.52 | 9.40 | 0.000 | 0.03 | 0.88 |
| 1023649 | TSS | ENSSSCG00000010302 | 14 | 76393149 | 76393174 | 0.01 | 0.01 | 0.01 | 2.68 | 2.24 | 2.53 | 0.01 | 2.48 | 0.00 | -8.03 | 1.25 | 19.18 | 0.000 | 0.01 | 3.92 |
| 1061055 | SKIP_OFF | ENSSSCG00000003908 | 6 | 165225268 | 165225351 | 0.11 | 0.12 | 0.00 | 16.83 | 13.30 | 16.59 | 0.08 | 15.57 | 0.00 | -7.69 | 7.82 | 14.43 | 0.000 | 0.02 | 2.80 |
| 1061052 | TSS | ENSSSCG00000003908 | 6 | 165229255 | 165229516 | 0.11 | 0.12 | 0.00 | 16.83 | 13.30 | 16.59 | 0.08 | 15.57 | 0.00 | -7.69 | 7.82 | 14.43 | 0.000 | 0.02 | 2.80 |
| 1009660 | SKIP_ON | ENSSSCG00000009428 | 11 | 24320403 | 24320462 | 0.06 | 0.36 | 0.34 | 28.09 | 37.33 | 27.05 | 0.25 | 30.82 | 0.01 | -6.93 | 15.54 | 9.94 | 0.000 | 0.03 | 1.14 |
| 1024132 | SKIP_ON | ENSSSCG00000010478 | 14 | 105021390 | 105021518 | 0.20 | 0.00 | 0.00 | 6.93 | 4.33 | 5.97 | 0.07 | 5.75 | 0.01 | -6.41 | 2.91 | 7.90 | 0.001 | 0.04 | 0.06 |
| 1052719 | SKIP_ON | ENSSSCG00000034242 | 4 | 60521119 | 60521274 | 0.02 | 0.00 | 0.04 | 1.86 | 1.24 | 1.71 | 0.02 | 1.61 | 0.01 | -6.39 | 0.81 | 8.86 | 0.001 | 0.03 | 0.60 |
| 1055923 | TSS | ENSSSCG00000000982 | 5 | 613477 | 613850 | 0.04 | 0.00 | 0.00 | 0.93 | 1.11 | 1.22 | 0.01 | 1.09 | 0.01 | -6.26 | 0.55 | 12.22 | 0.000 | 0.02 | 2.08 |
| 1027743 | TSS | ENSSSCG00000016030 | 15 | 91863966 | 91864130 | 0.23 | 0.00 | 0.07 | 5.40 | 6.73 | 6.71 | 0.10 | 6.28 | 0.02 | -5.96 | 3.19 | 14.70 | 0.000 | 0.02 | 2.88 |
| 1027744 | TTS | ENSSSCG00000016030 | 15 | 91841945 | 91842751 | 0.23 | 0.00 | 0.07 | 5.40 | 6.73 | 6.71 | 0.10 | 6.28 | 0.02 | -5.96 | 3.19 | 14.70 | 0.000 | 0.02 | 2.88 |
| 1031202 | TTS | ENSSSCG00000021527 | 16 | 62917712 | 62918473 | 0.00 | 0.00 | 0.04 | 0.82 | 0.51 | 0.82 | 0.01 | 0.72 | 0.02 | -5.60 | 0.37 | 6.62 | 0.002 | 0.05 | -0.78 |
| 1031209 | XIR_ON | ENSSSCG00000021527 | 16 | 62915281 | 62916956 | 0.00 | 0.00 | 0.04 | 0.82 | 0.51 | 0.82 | 0.01 | 0.72 | 0.02 | -5.60 | 0.37 | 6.62 | 0.002 | 0.05 | -0.78 |
| 1068439 | TSS | ENSSSCG00000002380 | 7 | 98265232 | 98265373 | 0.34 | 0.06 | 0.03 | 6.75 | 5.85 | 8.37 | 0.15 | 6.99 | 0.02 | -5.59 | 3.57 | 9.78 | 0.000 | 0.03 | 1.06 |
| 1060754 | TSS | ENSSSCG00000003819 | 6 | 149845338 | 149845854 | 0.22 | 0.03 | 0.00 | 5.00 | 3.50 | 3.12 | 0.08 | 3.87 | 0.02 | -5.57 | 1.98 | 6.94 | 0.001 | 0.05 | -0.55 |
| 1060755 | TTS | ENSSSCG00000003819 | 6 | 149836418 | 149836641 | 0.22 | 0.03 | 0.00 | 5.00 | 3.50 | 3.12 | 0.08 | 3.87 | 0.02 | -5.57 | 1.98 | 6.94 | 0.001 | 0.05 | -0.55 |
| 1061053 | TTS | ENSSSCG00000003908 | 6 | 165224620 | 165225107 | 14.06 | 0.82 | 1.05 | 267.71 | 179.95 | 308.57 | 5.31 | 252.08 | 0.02 | -5.57 | 128.69 | 6.86 | 0.001 | 0.05 | -0.60 |
| 1002948 | TSS | ENSSSCG00000004983 | 1 | 169777107 | 169777927 | 0.10 | 0.06 | 0.00 | 3.04 | 1.84 | 2.50 | 0.05 | 2.46 | 0.02 | -5.57 | 1.26 | 7.29 | 0.001 | 0.05 | -0.32 |
| 1031201 | TSS | ENSSSCG00000021527 | 16 | 62786414 | 62787095 | 0.00 | 0.06 | 0.05 | 1.68 | 1.34 | 1.87 | 0.04 | 1.63 | 0.02 | -5.51 | 0.83 | 10.52 | 0.000 | 0.02 | 1.40 |
| 1061050 | TSS | ENSSSCG00000003908 | 6 | 165229389 | 165229512 | 13.70 | 0.70 | 1.05 | 246.37 | 164.19 | 289.80 | 5.15 | 233.45 | 0.02 | -5.50 | 119.30 | 6.54 | 0.002 | 0.05 | -0.83 |
| 1061054 | SKIP_ON | ENSSSCG00000003908 | 6 | 165225268 | 165225351 | 13.95 | 0.70 | 1.05 | 250.88 | 166.65 | 291.99 | 5.23 | 236.51 | 0.02 | -5.50 | 120.87 | 6.62 | 0.002 | 0.05 | -0.78 |
| 1032964 | SKIP_OFF | ENSSSCG00000007485 | 17 | 54951144 | 54951185 | 0.11 | 0.00 | 0.00 | 1.94 | 1.21 | 1.73 | 0.04 | 1.63 | 0.02 | -5.49 | 0.83 | 7.59 | 0.001 | 0.04 | -0.13 |
| 1032971 | SKIP_ON | ENSSSCG00000007485 | 17 | 54942992 | 54943162 | 0.11 | 0.00 | 0.00 | 1.94 | 1.21 | 1.73 | 0.04 | 1.63 | 0.02 | -5.49 | 0.83 | 7.59 | 0.001 | 0.04 | -0.13 |
| 1004397 | TSS | ENSSSCG00000005529 | 1 | 262258298 | 262258395 | 1.22 | 0.05 | 0.03 | 17.12 | 23.30 | 17.68 | 0.43 | 19.37 | 0.02 | -5.48 | 9.90 | 10.00 | 0.000 | 0.03 | 1.16 |
| 1080598 | TSS | ENSSSCG00000030800 | X | 32018003 | 32018231 | 0.24 | 0.00 | 0.05 | 4.70 | 4.64 | 3.41 | 0.10 | 4.25 | 0.02 | -5.44 | 2.18 | 10.31 | 0.000 | 0.03 | 1.31 |
| 1080599 | TTS | ENSSSCG00000030800 | X | 32067493 | 32067687 | 0.24 | 0.00 | 0.05 | 4.70 | 4.64 | 3.41 | 0.10 | 4.25 | 0.02 | -5.44 | 2.18 | 10.31 | 0.000 | 0.03 | 1.31 |
| 1015890 | TSS | ENSSSCG00000011391 | 13 | 32329090 | 32329138 | 1.47 | 0.03 | 0.06 | 17.95 | 22.56 | 27.38 | 0.52 | 22.63 | 0.02 | -5.44 | 11.58 | 8.50 | 0.001 | 0.04 | 0.40 |
| 1015894 | XSKIP_OFF | ENSSSCG00000011391 | 13 | 32322687 | 32322717 | 1.47 | 0.03 | 0.06 | 17.95 | 22.56 | 27.38 | 0.52 | 22.63 | 0.02 | -5.44 | 11.58 | 8.50 | 0.001 | 0.04 | 0.40 |
| 1073936 | TSS | ENSSSCG00000014818 | 9 | 7326351 | 7326438 | 0.00 | 0.14 | 0.12 | 4.59 | 2.72 | 4.15 | 0.09 | 3.82 | 0.02 | -5.41 | 1.96 | 6.96 | 0.001 | 0.05 | -0.54 |
| 1058161 | TSS | ENSSSCG00000002817 | 6 | 19603448 | 19603471 | 1.44 | 0.11 | 0.18 | 24.21 | 22.04 | 26.64 | 0.58 | 24.30 | 0.02 | -5.39 | 12.44 | 18.01 | 0.000 | 0.01 | 3.69 |
| 1058162 | TTS | ENSSSCG00000002817 | 6 | 19628861 | 19629173 | 1.44 | 0.11 | 0.18 | 24.21 | 22.04 | 26.64 | 0.58 | 24.30 | 0.02 | -5.39 | 12.44 | 18.01 | 0.000 | 0.01 | 3.69 |
| 1008077 | TTS | ENSSSCG00000011078 | 10 | 51955000 | 51960197 | 1.39 | 0.07 | 0.20 | 17.32 | 25.28 | 27.29 | 0.56 | 23.30 | 0.02 | -5.39 | 11.93 | 7.87 | 0.001 | 0.04 | 0.04 |
| 1028387 | SKIP_OFF | ENSSSCG00000016203 | 15 | 121030260 | 121030304 | 0.18 | 0.00 | 0.02 | 3.35 | 2.92 | 2.13 | 0.07 | 2.80 | 0.02 | -5.37 | 1.43 | 7.98 | 0.001 | 0.04 | 0.10 |

| | | | | | | | | | | | | | | | | | | | | |
|---|---|---|---|---|---|---|---|---|---|---|---|---|---|---|---|---|---|---|---|---|
| 1032352 | SKIP_OFF | ENSSSCG00000007236 | 17 | 35585086 | 35585154 | 0.32 | 0.00 | 0.07 | 5.41 | 4.10 | 6.76 | 0.13 | 5.42 | 0.02 | -5.37 | 2.78 | 7.26 | 0.001 | 0.05 | -0.34 |
| 1069809 | TSS | ENSSSCG00000032691 | 7 | 41546878 | 41547100 | 1.31 | 0.15 | 0.18 | 18.13 | 20.05 | 29.44 | 0.55 | 22.54 | 0.02 | -5.37 | 11.54 | 6.65 | 0.002 | 0.05 | -0.75 |
| 1069810 | TTS | ENSSSCG00000032691 | 7 | 41558356 | 41562366 | 1.31 | 0.15 | 0.18 | 18.13 | 20.05 | 29.44 | 0.55 | 22.54 | 0.02 | -5.37 | 11.54 | 6.65 | 0.002 | 0.05 | -0.75 |
| 1008076 | TSS | ENSSSCG00000011078 | 10 | 52049317 | 52050127 | 1.39 | 0.07 | 0.20 | 16.99 | 24.93 | 26.90 | 0.56 | 22.94 | 0.02 | -5.37 | 11.75 | 7.78 | 0.001 | 0.04 | -0.01 |
| 1047690 | SKIP_ON | ENSSSCG00000033394 | 3 | 56061027 | 56061125 | 0.26 | 0.00 | 0.02 | 4.03 | 3.30 | 4.44 | 0.10 | 3.93 | 0.02 | -5.37 | 2.01 | 11.73 | 0.000 | 0.02 | 1.90 |
| 1060768 | MSKIP_ON | ENSSSCG00000003823 | 6 | 152205755 | 152212627 | 1.55 | 0.05 | 0.06 | 20.61 | 19.53 | 26.66 | 0.55 | 22.27 | 0.02 | -5.34 | 11.41 | 10.14 | 0.000 | 0.03 | 1.23 |
| 1018502 | TSS | ENSSSCG00000023084 | 13 | 66664365 | 66664474 | 0.31 | 0.00 | 0.00 | 4.96 | 3.60 | 3.86 | 0.10 | 4.14 | 0.02 | -5.33 | 2.12 | 9.96 | 0.000 | 0.03 | 1.15 |
| 1018507 | XMSKIP_ON | ENSSSCG00000023084 | 13 | 66534662 | 66538763 | 0.31 | 0.00 | 0.00 | 4.96 | 3.60 | 3.86 | 0.10 | 4.14 | 0.02 | -5.33 | 2.12 | 9.96 | 0.000 | 0.03 | 1.15 |
| 1018522 | XSKIP_ON | ENSSSCG00000023084 | 13 | 66534662 | 66534717 | 0.31 | 0.00 | 0.00 | 4.96 | 3.60 | 3.86 | 0.10 | 4.14 | 0.02 | -5.33 | 2.12 | 9.96 | 0.000 | 0.03 | 1.15 |
| 1066808 | TSS | ENSSSCG00000001594 | 7 | 35018707 | 35018856 | 0.13 | 0.03 | 0.03 | 2.70 | 2.25 | 3.14 | 0.07 | 2.70 | 0.02 | -5.32 | 1.38 | 10.70 | 0.000 | 0.02 | 1.48 |
| 1066809 | TTS | ENSSSCG00000001594 | 7 | 34918893 | 34919090 | 0.13 | 0.03 | 0.03 | 2.70 | 2.25 | 3.14 | 0.07 | 2.70 | 0.02 | -5.32 | 1.38 | 10.70 | 0.000 | 0.02 | 1.48 |
| 1059585 | TTS | ENSSSCG00000003455 | 6 | 74629758 | 74629778 | 0.00 | 0.13 | 0.16 | 3.68 | 3.38 | 4.91 | 0.10 | 3.99 | 0.03 | -5.32 | 2.05 | 8.75 | 0.001 | 0.03 | 0.54 |
| 1043107 | SKIP_ON | ENSSSCG00000039815 | 2 | 78003931 | 78003993 | 0.04 | 0.00 | 0.06 | 1.15 | 1.20 | 1.55 | 0.03 | 1.30 | 0.03 | -5.31 | 0.67 | 10.13 | 0.000 | 0.03 | 1.23 |
| 1043104 | TSS | ENSSSCG00000039815 | 2 | 77977459 | 77977954 | 0.04 | 0.00 | 0.06 | 1.15 | 1.20 | 1.55 | 0.03 | 1.30 | 0.03 | -5.31 | 0.67 | 10.13 | 0.000 | 0.03 | 1.23 |
| 1001220 | TSS | ENSSSCG00000004434 | 1 | 81713433 | 81714250 | 0.27 | 0.01 | 0.02 | 4.70 | 2.70 | 4.26 | 0.10 | 3.89 | 0.03 | -5.30 | 1.99 | 6.56 | 0.002 | 0.05 | -0.82 |
| 1001221 | TTS | ENSSSCG00000004434 | 1 | 81603273 | 81603701 | 0.27 | 0.01 | 0.02 | 4.70 | 2.70 | 4.26 | 0.10 | 3.89 | 0.03 | -5.30 | 1.99 | 6.56 | 0.002 | 0.05 | -0.82 |
| 1021955 | TSS | ENSSSCG00000009809 | 14 | 30991763 | 30992099 | 1.06 | 0.10 | 0.05 | 13.32 | 14.33 | 19.93 | 0.40 | 15.86 | 0.03 | -5.30 | 8.13 | 7.89 | 0.001 | 0.04 | 0.05 |
| 1021956 | TTS | ENSSSCG00000009809 | 14 | 31006309 | 31006620 | 1.06 | 0.10 | 0.05 | 13.32 | 14.33 | 19.93 | 0.40 | 15.86 | 0.03 | -5.30 | 8.13 | 7.89 | 0.001 | 0.04 | 0.05 |
| 1067076 | TTS | ENSSSCG00000001705 | 7 | 39281997 | 39282593 | 0.59 | 0.01 | 0.00 | 7.67 | 6.29 | 9.40 | 0.20 | 7.78 | 0.03 | -5.29 | 3.99 | 8.74 | 0.001 | 0.03 | 0.54 |
| 1012897 | TSS | ENSSSCG00000017884 | 12 | 50549637 | 50549758 | 2.68 | 0.16 | 0.28 | 41.60 | 34.24 | 44.99 | 1.04 | 40.28 | 0.03 | -5.28 | 20.66 | 12.72 | 0.000 | 0.02 | 2.26 |
| 1060766 | TSS | ENSSSCG00000003823 | 6 | 152175539 | 152175645 | 1.55 | 0.05 | 0.13 | 20.61 | 19.53 | 26.66 | 0.57 | 22.27 | 0.03 | -5.28 | 11.42 | 10.14 | 0.000 | 0.03 | 1.23 |
| 1060767 | TTS | ENSSSCG00000003823 | 6 | 152255060 | 152255220 | 1.55 | 0.05 | 0.13 | 20.61 | 19.53 | 26.66 | 0.57 | 22.27 | 0.03 | -5.28 | 11.42 | 10.14 | 0.000 | 0.03 | 1.23 |
| 1071357 | TSS | ENSSSCG00000008943 | 8 | 67872872 | 67872966 | 0.13 | 0.00 | 0.00 | 1.75 | 1.52 | 1.89 | 0.04 | 1.72 | 0.03 | -5.26 | 0.88 | 14.40 | 0.000 | 0.02 | 2.79 |
| 1052721 | SKIP_ON | ENSSSCG00000034242 | 4 | 60526376 | 60526470 | 0.07 | 0.01 | 0.04 | 1.86 | 1.34 | 1.72 | 0.04 | 1.64 | 0.03 | -5.25 | 0.84 | 10.48 | 0.000 | 0.02 | 1.38 |
| 1012898 | TTS | ENSSSCG00000017884 | 12 | 50524200 | 50527330 | 3.18 | 0.19 | 0.33 | 48.05 | 39.11 | 51.60 | 1.23 | 46.25 | 0.03 | -5.23 | 23.74 | 12.45 | 0.000 | 0.02 | 2.16 |
| 1020124 | TSS | ENSSSCG00000033462 | 13 | 146480431 | 146480525 | 1.55 | 0.22 | 0.46 | 25.41 | 36.21 | 22.11 | 0.74 | 27.91 | 0.03 | -5.23 | 14.33 | 6.75 | 0.002 | 0.05 | -0.68 |
| 1020126 | TTS | ENSSSCG00000033462 | 13 | 146488120 | 146488338 | 1.55 | 0.22 | 0.46 | 25.41 | 36.21 | 22.11 | 0.74 | 27.91 | 0.03 | -5.23 | 14.33 | 6.75 | 0.002 | 0.05 | -0.68 |
| 1055933 | TSS | ENSSSCG00000000983 | 5 | 646947 | 647067 | 0.21 | 0.03 | 0.06 | 2.87 | 4.44 | 3.68 | 0.10 | 3.66 | 0.03 | -5.23 | 1.88 | 8.25 | 0.001 | 0.04 | 0.27 |
| 1055934 | TTS | ENSSSCG00000000983 | 5 | 659313 | 662557 | 0.21 | 0.03 | 0.06 | 2.87 | 4.44 | 3.68 | 0.10 | 3.66 | 0.03 | -5.23 | 1.88 | 8.25 | 0.001 | 0.04 | 0.27 |
| 1072833 | TSS | ENSSSCG00000028471 | 8 | 27998300 | 27998796 | 1.78 | 0.11 | 0.23 | 31.79 | 22.86 | 24.39 | 0.71 | 26.35 | 0.03 | -5.22 | 13.53 | 9.70 | 0.000 | 0.03 | 1.03 |
| 1072834 | TTS | ENSSSCG00000028471 | 8 | 28071843 | 28072201 | 1.78 | 0.11 | 0.23 | 31.79 | 22.86 | 24.39 | 0.71 | 26.35 | 0.03 | -5.22 | 13.53 | 9.70 | 0.000 | 0.03 | 1.03 |
| 1056947 | TSS | ENSSSCG00000033574 | 5 | 87089867 | 87090254 | 0.14 | 0.00 | 0.00 | 1.86 | 1.83 | 1.70 | 0.05 | 1.80 | 0.03 | -5.21 | 0.92 | 23.26 | 0.000 | 0.01 | 4.58 |
| 1056950 | XAE | ENSSSCG00000033574 | 5 | 87058804 | 87058889 | 0.14 | 0.00 | 0.00 | 1.86 | 1.83 | 1.70 | 0.05 | 1.80 | 0.03 | -5.21 | 0.92 | 23.26 | 0.000 | 0.01 | 4.58 |
| 1047684 | TSS | ENSSSCG00000033394 | 3 | 56171560 | 56171826 | 0.31 | 0.02 | 0.02 | 4.42 | 4.11 | 4.80 | 0.12 | 4.45 | 0.03 | -5.20 | 2.28 | 20.37 | 0.000 | 0.01 | 4.14 |
| 1047685 | TTS | ENSSSCG00000033394 | 3 | 55950591 | 55951982 | 0.31 | 0.02 | 0.02 | 4.42 | 4.11 | 4.80 | 0.12 | 4.45 | 0.03 | -5.20 | 2.28 | 20.37 | 0.000 | 0.01 | 4.14 |

| | | | | | | | | | | | | | | | | | | | | |
|---|---|---|---|---|---|---|---|---|---|---|---|---|---|---|---|---|---|---|---|---|
| 1026732 | SKIP_ON | ENSSSCG00000015696 | 15 | 16916357 | 16916552 | 0.70 | 0.00 | 0.00 | 10.47 | 7.47 | 7.77 | 0.23 | 8.57 | 0.03 | -5.20 | 4.40 | 9.03 | 0.000 | 0.03 | 0.69 |
| 1011903 | TTS | ENSSSCG00000017495 | 12 | 22573320 | 22575118 | 0.33 | 0.06 | 0.10 | 7.30 | 4.24 | 6.32 | 0.17 | 5.95 | 0.03 | -5.17 | 3.06 | 6.77 | 0.002 | 0.05 | -0.67 |
| 1008021 | SKIP_OFF | ENSSSCG00000011069 | 10 | 39423888 | 39423986 | 0.97 | 0.00 | 0.13 | 15.58 | 11.85 | 11.95 | 0.37 | 13.12 | 0.03 | -5.16 | 6.75 | 10.71 | 0.000 | 0.02 | 1.48 |
| 1071282 | TSS | ENSSSCG00000008908 | 8 | 54945992 | 54946000 | 0.42 | 0.00 | 0.00 | 4.25 | 4.79 | 6.05 | 0.14 | 5.03 | 0.03 | -5.16 | 2.59 | 9.39 | 0.000 | 0.03 | 0.87 |
| 1071283 | TTS | ENSSSCG00000008908 | 8 | 54917973 | 54918127 | 0.42 | 0.00 | 0.00 | 4.25 | 4.79 | 6.05 | 0.14 | 5.03 | 0.03 | -5.16 | 2.59 | 9.39 | 0.000 | 0.03 | 0.87 |
| 1071107 | TSS | ENSSSCG00000008834 | 8 | 39162404 | 39163361 | 5.66 | 0.15 | 0.16 | 76.01 | 53.82 | 83.40 | 1.99 | 71.08 | 0.03 | -5.16 | 36.54 | 8.09 | 0.001 | 0.04 | 0.17 |
| 1018509 | MSKIP_ON | ENSSSCG00000023084 | 13 | 66583381 | 66589681 | 0.64 | 0.02 | 0.00 | 7.55 | 6.83 | 9.00 | 0.22 | 7.79 | 0.03 | -5.15 | 4.01 | 11.93 | 0.000 | 0.02 | 1.97 |
| 1018516 | MSKIP_ON | ENSSSCG00000023084 | 13 | 66588449 | 66589681 | 0.64 | 0.02 | 0.00 | 7.55 | 6.83 | 9.00 | 0.22 | 7.79 | 0.03 | -5.15 | 4.01 | 11.93 | 0.000 | 0.02 | 1.97 |
| 1018520 | SKIP_ON | ENSSSCG00000023084 | 13 | 66588449 | 66588508 | 0.64 | 0.02 | 0.00 | 7.55 | 6.83 | 9.00 | 0.22 | 7.79 | 0.03 | -5.15 | 4.01 | 11.93 | 0.000 | 0.02 | 1.97 |
| 1047687 | SKIP_OFF | ENSSSCG00000033394 | 3 | 56169058 | 56169214 | 0.27 | 0.02 | 0.00 | 3.28 | 3.73 | 3.39 | 0.10 | 3.47 | 0.03 | -5.14 | 1.78 | 21.72 | 0.000 | 0.01 | 4.36 |
| 1047692 | SKIP_ON | ENSSSCG00000033394 | 3 | 56166194 | 56166421 | 0.27 | 0.02 | 0.00 | 3.28 | 3.73 | 3.39 | 0.10 | 3.47 | 0.03 | -5.14 | 1.78 | 21.72 | 0.000 | 0.01 | 4.36 |
| 1000411 | TSS | ENSSSCG00000004147 | 1 | 25644193 | 25645094 | 1.27 | 0.09 | 0.12 | 18.35 | 16.93 | 16.46 | 0.49 | 17.25 | 0.03 | -5.13 | 8.87 | 25.78 | 0.000 | 0.01 | 4.89 |
| 1000412 | TTS | ENSSSCG00000004147 | 1 | 25566903 | 25567630 | 1.27 | 0.09 | 0.12 | 18.35 | 16.93 | 16.46 | 0.49 | 17.25 | 0.03 | -5.13 | 8.87 | 25.78 | 0.000 | 0.01 | 4.89 |
| 1023808 | IR_ON | ENSSSCG00000010339 | 14 | 82202414 | 82202538 | 0.00 | 0.00 | 0.04 | 0.43 | 0.49 | 0.33 | 0.01 | 0.42 | 0.03 | -5.12 | 0.21 | 6.76 | 0.002 | 0.05 | -0.67 |
| 1065726 | TSS | ENSSSCG00000040973 | 6 | 14233928 | 14233941 | 1.85 | 0.25 | 0.21 | 26.08 | 25.32 | 28.61 | 0.77 | 26.67 | 0.03 | -5.11 | 13.72 | 24.32 | 0.000 | 0.01 | 4.72 |
| 1065727 | TTS | ENSSSCG00000040973 | 6 | 13895493 | 13895975 | 1.85 | 0.25 | 0.21 | 26.08 | 25.32 | 28.61 | 0.77 | 26.67 | 0.03 | -5.11 | 13.72 | 24.32 | 0.000 | 0.01 | 4.72 |
| 1024130 | TSS | ENSSSCG00000010478 | 14 | 105011408 | 105012556 | 0.85 | 0.03 | 0.00 | 12.00 | 7.38 | 11.05 | 0.29 | 10.14 | 0.03 | -5.11 | 5.22 | 7.27 | 0.001 | 0.05 | -0.33 |
| 1024131 | TTS | ENSSSCG00000010478 | 14 | 105033136 | 105033525 | 0.85 | 0.03 | 0.00 | 12.00 | 7.38 | 11.05 | 0.29 | 10.14 | 0.03 | -5.11 | 5.22 | 7.27 | 0.001 | 0.05 | -0.33 |
| 1018229 | TSS | ENSSSCG00000020990 | 13 | 39318872 | 39319018 | 0.79 | 0.03 | 0.11 | 11.27 | 12.59 | 8.34 | 0.31 | 10.74 | 0.03 | -5.10 | 5.52 | 8.66 | 0.001 | 0.03 | 0.50 |
| 1018230 | TTS | ENSSSCG00000020990 | 13 | 39148454 | 39148644 | 0.79 | 0.03 | 0.11 | 11.27 | 12.59 | 8.34 | 0.31 | 10.74 | 0.03 | -5.10 | 5.52 | 8.66 | 0.001 | 0.03 | 0.50 |
| 1032349 | TSS | ENSSSCG00000007236 | 17 | 35558301 | 35558596 | 0.91 | 0.03 | 0.12 | 13.24 | 9.15 | 13.57 | 0.35 | 11.99 | 0.03 | -5.09 | 6.17 | 8.53 | 0.001 | 0.04 | 0.42 |
| 1032350 | TTS | ENSSSCG00000007236 | 17 | 35607702 | 35607778 | 0.91 | 0.03 | 0.12 | 13.24 | 9.15 | 13.57 | 0.35 | 11.99 | 0.03 | -5.09 | 6.17 | 8.53 | 0.001 | 0.04 | 0.42 |
| 1026730 | TSS | ENSSSCG00000015696 | 15 | 16911090 | 16911378 | 0.70 | 0.01 | 0.05 | 10.47 | 7.47 | 7.77 | 0.25 | 8.57 | 0.03 | -5.09 | 4.41 | 9.03 | 0.000 | 0.03 | 0.69 |
| 1026731 | TTS | ENSSSCG00000015696 | 15 | 16970116 | 16970182 | 0.70 | 0.01 | 0.05 | 10.47 | 7.47 | 7.77 | 0.25 | 8.57 | 0.03 | -5.09 | 4.41 | 9.03 | 0.000 | 0.03 | 0.69 |
| 1068842 | TSS | ENSSSCG00000002504 | 7 | 117609840 | 117609972 | 3.62 | 0.31 | 0.78 | 54.86 | 46.89 | 56.34 | 1.57 | 52.69 | 0.03 | -5.07 | 27.13 | 17.47 | 0.000 | 0.01 | 3.57 |
| 1068843 | TTS | ENSSSCG00000002504 | 7 | 117666773 | 117667455 | 3.62 | 0.31 | 0.78 | 54.86 | 46.89 | 56.34 | 1.57 | 52.69 | 0.03 | -5.07 | 27.13 | 17.47 | 0.000 | 0.01 | 3.57 |
| 1028177 | TTS | ENSSSCG00000016133 | 15 | 109989062 | 109992312 | 0.10 | 0.03 | 0.01 | 1.92 | 1.55 | 1.33 | 0.05 | 1.60 | 0.03 | -5.07 | 0.82 | 9.30 | 0.000 | 0.03 | 0.83 |
| 1000305 | TSS | ENSSSCG00000004114 | 1 | 19029280 | 19029353 | 0.54 | 0.05 | 0.19 | 9.74 | 9.17 | 6.99 | 0.26 | 8.63 | 0.03 | -5.06 | 4.45 | 10.47 | 0.000 | 0.03 | 1.38 |
| 1022929 | TSS | ENSSSCG00000010100 | 14 | 50622937 | 50623069 | 0.36 | 0.10 | 0.03 | 5.93 | 4.70 | 5.42 | 0.16 | 5.35 | 0.03 | -5.06 | 2.76 | 14.88 | 0.000 | 0.01 | 2.93 |
| 1022930 | TTS | ENSSSCG00000010100 | 14 | 50603323 | 50603460 | 0.36 | 0.10 | 0.03 | 5.93 | 4.70 | 5.42 | 0.16 | 5.35 | 0.03 | -5.06 | 2.76 | 14.88 | 0.000 | 0.01 | 2.93 |
| 1028390 | IR_ON | ENSSSCG00000016203 | 15 | 121035540 | 121035685 | 0.34 | 0.00 | 0.02 | 5.11 | 3.50 | 3.16 | 0.12 | 3.93 | 0.03 | -5.05 | 2.02 | 6.61 | 0.002 | 0.05 | -0.78 |
| 1028384 | TTS | ENSSSCG00000016203 | 15 | 121027040 | 121028735 | 0.34 | 0.00 | 0.02 | 5.11 | 3.50 | 3.16 | 0.12 | 3.93 | 0.03 | -5.05 | 2.02 | 6.61 | 0.002 | 0.05 | -0.78 |
| 1028388 | XIR_ON | ENSSSCG00000016203 | 15 | 121035540 | 121035685 | 0.34 | 0.00 | 0.02 | 5.11 | 3.50 | 3.16 | 0.12 | 3.93 | 0.03 | -5.05 | 2.02 | 6.61 | 0.002 | 0.05 | -0.78 |
| 1011539 | TSS | ENSSSCG00000017344 | 12 | 18471963 | 18472114 | 0.23 | 0.06 | 0.06 | 3.17 | 3.86 | 4.42 | 0.12 | 3.82 | 0.03 | -5.02 | 1.97 | 10.70 | 0.000 | 0.02 | 1.48 |
| 1070596 | TSS | ENSSSCG00000039903 | 7 | 39366987 | 39367116 | 0.08 | 0.24 | 0.42 | 9.54 | 6.22 | 8.35 | 0.25 | 8.04 | 0.03 | -5.02 | 4.14 | 8.49 | 0.001 | 0.04 | 0.40 |

| | | | | | | | | | | | | | | | | | | | | |
|---|---|---|---|---|---|---|---|---|---|---|---|---|---|---|---|---|---|---|---|---|
| 1070597 | TTS | ENSSSCG00000039903 | 7 | 39381748 | 39381790 | 0.08 | 0.24 | 0.42 | 9.54 | 6.22 | 8.35 | 0.25 | 8.04 | 0.03 | -5.02 | 4.14 | 8.49 | 0.001 | 0.04 | 0.40 |
| 1009658 | TSS | ENSSSCG00000009428 | 11 | 24321827 | 24321920 | 4.09 | 0.36 | 0.34 | 43.78 | 53.23 | 57.73 | 1.60 | 51.58 | 0.03 | -5.01 | 26.59 | 12.37 | 0.000 | 0.02 | 2.13 |
| 1009659 | TTS | ENSSSCG00000009428 | 11 | 24312150 | 24314072 | 4.09 | 0.36 | 0.34 | 43.78 | 53.23 | 57.73 | 1.60 | 51.58 | 0.03 | -5.01 | 26.59 | 12.37 | 0.000 | 0.02 | 2.13 |
| 1051377 | TSS | ENSSSCG00000006792 | 4 | 108951206 | 108951820 | 0.04 | 0.00 | 0.01 | 0.72 | 0.56 | 0.61 | 0.02 | 0.63 | 0.03 | -5.01 | 0.32 | 10.41 | 0.000 | 0.03 | 1.35 |
| 1051378 | TTS | ENSSSCG00000006792 | 4 | 108961813 | 108963665 | 0.04 | 0.00 | 0.01 | 0.72 | 0.56 | 0.61 | 0.02 | 0.63 | 0.03 | -5.01 | 0.32 | 10.41 | 0.000 | 0.03 | 1.35 |
| 1026841 | TSS | ENSSSCG00000015741 | 15 | 31685571 | 31686108 | 0.70 | 0.08 | 0.04 | 9.59 | 6.63 | 10.26 | 0.27 | 8.83 | 0.03 | -5.01 | 4.55 | 8.00 | 0.001 | 0.04 | 0.12 |
| 1026842 | TTS | ENSSSCG00000015741 | 15 | 31599673 | 31600331 | 0.70 | 0.08 | 0.04 | 9.59 | 6.63 | 10.26 | 0.27 | 8.83 | 0.03 | -5.01 | 4.55 | 8.00 | 0.001 | 0.04 | 0.12 |
| 1052718 | TTS | ENSSSCG00000034242 | 4 | 60505066 | 60507900 | 0.09 | 0.01 | 0.05 | 1.86 | 1.34 | 1.72 | 0.05 | 1.64 | 0.03 | -5.01 | 0.84 | 10.37 | 0.000 | 0.03 | 1.33 |
| 1058034 | TTS | ENSSSCG00000002768 | 6 | 28446687 | 28446702 | 1.42 | 0.93 | 1.36 | 0.00 | 0.00 | 0.00 | 1.24 | 0.00 | 695762.58 | 19.41 | 0.62 | -8.18 | 0.001 | 0.04 | 0.22 |
| 1000765 | TSS | ENSSSCG00000004263 | 1 | 44940794 | 44940850 | 1.55 | 1.24 | 1.02 | 0.00 | 0.00 | 0.00 | 1.27 | 0.00 | 51279.67 | 15.65 | 0.63 | -8.40 | 0.001 | 0.04 | 0.35 |
| 1019643 | TSS | ENSSSCG00000029811 | 13 | 3980464 | 3980557 | 0.71 | 0.92 | 0.84 | 0.00 | 0.00 | 0.00 | 0.83 | 0.00 | 5943.19 | 12.54 | 0.41 | -11.90 | 0.000 | 0.02 | 1.96 |
| 1014231 | SKIP_OFF | ENSSSCG00000033228 | 12 | 27261499 | 27261513 | 2.15 | 1.93 | 1.54 | 0.00 | 0.00 | 0.00 | 1.87 | 0.00 | 4076.38 | 11.99 | 0.94 | -10.91 | 0.000 | 0.02 | 1.57 |
| 1014220 | SKIP_ON | ENSSSCG00000033228 | 12 | 27262711 | 27262722 | 2.15 | 1.93 | 1.54 | 0.00 | 0.00 | 0.00 | 1.87 | 0.00 | 4076.38 | 11.99 | 0.94 | -10.91 | 0.000 | 0.02 | 1.57 |
| 1049709 | TSS | ENSSSCG00000006191 | 4 | 64710542 | 64710633 | 0.90 | 0.76 | 0.79 | 0.00 | 0.00 | 0.00 | 0.82 | 0.00 | 3976.77 | 11.96 | 0.41 | -14.88 | 0.000 | 0.01 | 2.93 |
| 1023585 | SKIP_OFF | ENSSSCG00000010281 | 14 | 74747072 | 74747197 | 4.37 | 5.26 | 5.11 | 0.00 | 0.00 | 0.02 | 4.91 | 0.01 | 761.07 | 9.57 | 2.46 | -18.82 | 0.000 | 0.01 | 3.85 |
| 1069392 | SKIP_OFF | ENSSSCG00000028225 | 7 | 116374054 | 116374209 | 0.31 | 0.33 | 0.31 | 0.00 | 0.00 | 0.00 | 0.32 | 0.00 | 728.24 | 9.51 | 0.16 | -8.23 | 0.001 | 0.04 | 0.25 |
| 1029818 | TTS | ENSSSCG00000034942 | 15 | 48383713 | 48388326 | 107.11 | 101.29 | 85.77 | 0.41 | 2.00 | 2.25 | 98.06 | 1.55 | 63.07 | 5.98 | 49.81 | -16.03 | 0.000 | 0.01 | 3.23 |
| 1029817 | TSS | ENSSSCG00000034942 | 15 | 48377513 | 48377756 | 111.90 | 104.71 | 87.37 | 0.50 | 2.11 | 2.33 | 101.33 | 1.65 | 61.40 | 5.94 | 51.49 | -14.51 | 0.000 | 0.02 | 2.82 |
| 1075974 | SKIP_OFF | ENSSSCG00000015537 | 9 | 122359024 | 122359125 | 0.51 | 0.53 | 0.42 | 0.00 | 0.03 | 0.00 | 0.49 | 0.01 | 41.96 | 5.39 | 0.25 | -9.42 | 0.000 | 0.03 | 0.89 |
| 1067382 | MSKIP_OFF | ENSSSCG00000001825 | 7 | 53686932 | 53687906 | 1.26 | 1.12 | 1.14 | 0.00 | 0.08 | 0.00 | 1.18 | 0.03 | 41.89 | 5.39 | 0.60 | -18.80 | 0.000 | 0.01 | 3.85 |
| 1024037 | TSS | ENSSSCG00000010455 | 14 | 101367855 | 101368158 | 0.98 | 0.61 | 0.86 | 0.08 | 0.00 | 0.00 | 0.81 | 0.03 | 31.72 | 4.99 | 0.42 | -7.09 | 0.001 | 0.05 | -0.45 |
| 1016020 | TSS | ENSSSCG00000011423 | 13 | 33926804 | 33926915 | 0.34 | 0.27 | 0.28 | 0.01 | 0.03 | 0.02 | 0.30 | 0.02 | 16.50 | 4.04 | 0.16 | -6.47 | 0.002 | 0.05 | -0.88 |
| 1032155 | MSKIP_OFF | ENSSSCG00000007133 | 17 | 30804841 | 30805130 | 0.70 | 1.03 | 0.76 | 0.17 | 0.00 | 0.00 | 0.83 | 0.06 | 14.43 | 3.85 | 0.44 | -6.61 | 0.002 | 0.05 | -0.78 |
| 1032151 | XSKIP_OFF | ENSSSCG00000007133 | 17 | 30805003 | 30805130 | 0.70 | 1.03 | 0.76 | 0.17 | 0.00 | 0.00 | 0.83 | 0.06 | 14.43 | 3.85 | 0.44 | -6.61 | 0.002 | 0.05 | -0.78 |
| 1031358 | TSS | ENSSSCG00000030343 | 16 | 34871780 | 34871822 | 0.60 | 0.77 | 0.62 | 0.00 | 0.00 | 0.19 | 0.66 | 0.06 | 10.55 | 3.40 | 0.36 | -6.99 | 0.001 | 0.05 | -0.52 |
| 1031365 | XAE | ENSSSCG00000030343 | 16 | 34839254 | 34839386 | 0.60 | 0.77 | 0.62 | 0.00 | 0.00 | 0.19 | 0.66 | 0.06 | 10.55 | 3.40 | 0.36 | -6.99 | 0.001 | 0.05 | -0.52 |
| 1000555 | TSS | ENSSSCG00000004195 | 1 | 32051209 | 32051376 | 1.39 | 1.37 | 1.53 | 0.16 | 0.26 | 0.00 | 1.43 | 0.14 | 10.28 | 3.36 | 0.79 | -13.70 | 0.000 | 0.02 | 2.58 |
| 1008290 | TTS | ENSSSCG00000011147 | 10 | 65572686 | 65574571 | 119.94 | 95.45 | 81.49 | 1.17 | 16.69 | 16.71 | 98.96 | 11.52 | 8.59 | 3.10 | 55.24 | -7.51 | 0.001 | 0.04 | -0.18 |
| 1019143 | MSKIP_OFF | ENSSSCG00000026746 | 13 | 79332564 | 79332882 | 1.09 | 1.00 | 1.10 | 0.13 | 0.26 | 0.00 | 1.06 | 0.13 | 8.10 | 3.02 | 0.60 | -10.75 | 0.000 | 0.02 | 1.50 |
| 1019148 | SKIP_OFF | ENSSSCG00000026746 | 13 | 79330784 | 79330858 | 1.09 | 1.00 | 1.10 | 0.13 | 0.26 | 0.00 | 1.06 | 0.13 | 8.10 | 3.02 | 0.60 | -10.75 | 0.000 | 0.02 | 1.50 |
| 1045599 | SKIP_OFF | ENSSSCG00000008421 | 3 | 92002381 | 92002550 | 23.42 | 15.59 | 20.60 | 0.12 | 5.72 | 2.14 | 19.87 | 2.66 | 7.47 | 2.90 | 11.26 | -6.50 | 0.002 | 0.05 | -0.86 |
| 1062615 | TSS | ENSSSCG00000027684 | 6 | 83507215 | 83508007 | 0.61 | 0.50 | 0.57 | 0.01 | 0.16 | 0.06 | 0.56 | 0.08 | 7.34 | 2.87 | 0.32 | -7.53 | 0.001 | 0.04 | -0.17 |
| 1062616 | TTS | ENSSSCG00000027684 | 6 | 83488973 | 83493215 | 0.61 | 0.50 | 0.57 | 0.01 | 0.16 | 0.06 | 0.56 | 0.08 | 7.34 | 2.87 | 0.32 | -7.53 | 0.001 | 0.04 | -0.17 |
| 1045593 | TSS | ENSSSCG00000008421 | 3 | 91964415 | 91964639 | 24.13 | 16.40 | 21.30 | 0.12 | 6.08 | 2.36 | 20.61 | 2.85 | 7.22 | 2.85 | 11.73 | -6.62 | 0.002 | 0.05 | -0.77 |
| 1045594 | TTS | ENSSSCG00000008421 | 3 | 92020227 | 92021398 | 21.40 | 15.59 | 20.17 | 0.12 | 6.08 | 2.22 | 19.05 | 2.81 | 6.79 | 2.76 | 10.93 | -6.95 | 0.001 | 0.05 | -0.55 |

| | | | | | | | | | | | | | | | | | | | | |
|---|---|---|---|---|---|---|---|---|---|---|---|---|---|---|---|---|---|---|---|---|
| 1045601 | XSKIP_OFF | ENSSSCG00000008421 | 3 | 92015950 | 92015950 | 21.40 | 15.59 | 20.17 | 0.12 | 6.08 | 2.22 | 19.05 | 2.81 | 6.79 | 2.76 | 10.93 | -6.95 | 0.001 | 0.05 | -0.55 |
| 1045596 | SKIP_ON | ENSSSCG00000008421 | 3 | 91984228 | 91984299 | 21.40 | 15.59 | 20.17 | 0.12 | 6.08 | 2.22 | 19.05 | 2.81 | 6.79 | 2.76 | 10.93 | -6.95 | 0.001 | 0.05 | -0.55 |
| 1010172 | TSS | ENSSSCG00000022159 | 11 | 18756040 | 18756356 | 5.62 | 6.01 | 5.44 | 0.17 | 2.05 | 0.74 | 5.69 | 0.98 | 5.78 | 2.53 | 3.34 | -8.59 | 0.001 | 0.03 | 0.45 |
| 1000556 | TTS | ENSSSCG00000004195 | 1 | 32006042 | 32007153 | 1.72 | 1.37 | 1.97 | 0.26 | 0.48 | 0.19 | 1.68 | 0.31 | 5.44 | 2.44 | 1.00 | -7.36 | 0.001 | 0.04 | -0.27 |
| 1079470 | TSS | ENSSSCG00000012638 | X | 100826849 | 100827728 | 13.78 | 17.69 | 16.20 | 0.83 | 5.48 | 2.63 | 15.89 | 2.98 | 5.33 | 2.41 | 9.44 | -7.76 | 0.001 | 0.04 | -0.02 |
| 1023991 | SKIP_ON | ENSSSCG00000010437 | 14 | 99783084 | 99783098 | 45.82 | 54.54 | 58.86 | 1.47 | 18.39 | 14.04 | 53.07 | 11.30 | 4.70 | 2.23 | 32.19 | -6.98 | 0.001 | 0.05 | -0.52 |
| 1018359 | TTS | ENSSSCG00000021966 | 13 | 74766604 | 74766639 | 0.92 | 1.04 | 0.80 | 0.19 | 0.24 | 0.19 | 0.92 | 0.21 | 4.46 | 2.16 | 0.56 | -9.21 | 0.000 | 0.03 | 0.78 |
| 1065608 | TTS | ENSSSCG00000040607 | 6 | 8484601 | 8485030 | 2.36 | 2.82 | 2.31 | 0.16 | 1.07 | 0.53 | 2.49 | 0.59 | 4.24 | 2.08 | 1.54 | -6.49 | 0.002 | 0.05 | -0.87 |
| 1080548 | TSS | ENSSSCG00000030241 | X | 88234558 | 88234608 | 38.74 | 32.90 | 32.73 | 3.64 | 8.80 | 12.52 | 34.79 | 8.32 | 4.18 | 2.06 | 21.56 | -8.67 | 0.001 | 0.03 | 0.50 |
| 1031575 | TSS | ENSSSCG00000036438 | 16 | 71988875 | 71989011 | 1478.32 | 1110.57 | 1600.09 | 104.47 | 472.37 | 403.96 | 1396.32 | 326.93 | 4.27 | 2.09 | 861.63 | -5.20 | 0.004 | 0.06 | -1.89 |
| 1031576 | TTS | ENSSSCG00000036438 | 16 | 71980468 | 71981522 | 1478.32 | 1110.57 | 1600.09 | 104.47 | 472.37 | 403.96 | 1396.32 | 326.93 | 4.27 | 2.09 | 861.63 | -5.20 | 0.004 | 0.06 | -1.89 |

### Table S6 Differentially spliced Genes

| Gene stable ID | Gene description | Gene name |
|---|---|---|
| ENSSSCG00000009755 | acetoacetyl-CoA synthetase [Source:HGNC Symbol;Acc:HGNC:21298] | AACS |
| ENSSSCG00000026473 | ATP binding cassette subfamily A member 8 [Source:HGNC Symbol;Acc:HGNC:38] | ABCA8 |
| ENSSSCG00000024127 | abl interactor 1 [Source:HGNC Symbol;Acc:HGNC:11320] | ABI1 |
| ENSSSCG00000010737 | abraxas 2, BRISC complex subunit [Source:HGNC Symbol;Acc:HGNC:28975] | ABRAXAS2 |
| ENSSSCG00000007857 | acyl-CoA synthetase medium chain family member 3 [Source:HGNC Symbol;Acc:HGNC:10522] | ACSM3 |
| ENSSSCG00000007133 | acyl-CoA synthetase short chain family member 1 [Source:HGNC Symbol;Acc:HGNC:16091] | ACSS1 |
| ENSSSCG00000000939 | acyl-CoA synthetase short chain family member 3 [Source:HGNC Symbol;Acc:HGNC:24723] | ACSS3 |
| ENSSSCG00000010746 | ADAM metallopeptidase domain 12 [Source:HGNC Symbol;Acc:HGNC:190] | ADAM12 |
| ENSSSCG00000004270 | adhesion G protein-coupled receptor B3 [Source:HGNC Symbol;Acc:HGNC:945] | ADGRB3 |
| ENSSSCG00000038732 | ArfGAP with GTPase domain, ankyrin repeat and PH domain 2 | AGAP2 |
| ENSSSCG00000008558 | ATP/GTP binding protein like 5 [Source:HGNC Symbol;Acc:HGNC:26147] | AGBL5 |
| ENSSSCG00000002504 | adenylate kinase 7 [Source:HGNC Symbol;Acc:HGNC:20091] | AK7 |
| ENSSSCG00000023894 | AT-hook transcription factor [Source:HGNC Symbol;Acc:HGNC:24108] | AKNA |
| ENSSSCG00000028996 | aldehyde dehydrogenase 1 family member A1 [Source:HGNC Symbol;Acc:HGNC:402] | ALDH1A1 |
| ENSSSCG00000030626 | aldehyde dehydrogenase 1 family member L1 [Source:HGNC Symbol;Acc:HGNC:3978] | ALDH1L1 |
| ENSSSCG00000028501 | aldehyde dehydrogenase 3 family member B1 [Source:HGNC Symbol;Acc:HGNC:410] | ALDH3B1 |
| ENSSSCG00000021997 | ALS2 C-terminal like [Source:HGNC Symbol;Acc:HGNC:20605] | ALS2CL |
| ENSSSCG00000016105 | amyotrophic lateral sclerosis 2 chromosome region 12 [Source:HGNC Symbol;Acc:HGNC:14439] | ALS2CR12 |
| ENSSSCG00000009814 | anaphase promoting complex subunit 5 [Source:HGNC Symbol;Acc:HGNC:15713] | ANAPC5 |
| ENSSSCG00000003819 | Sus scrofa angiopoietin like 3 (ANGPTL3), mRNA. [Source:RefSeq mRNA;Acc:NM_001003926] | ANGPTL3 |
| ENSSSCG00000017609 | ankyrin repeat and fibronectin type III domain containing 1 | ANKFN1 |
| ENSSSCG00000016365 | ankyrin repeat and MYND domain containing 1 [Source:HGNC Symbol;Acc:HGNC:20987] | ANKMY1 |
| ENSSSCG00000013497 | ankyrin repeat domain 24 [Source:HGNC Symbol;Acc:HGNC:29424] | ANKRD24 |
| ENSSSCG00000015491 | ankyrin repeat domain 45 [Source:HGNC Symbol;Acc:HGNC:24786] | ANKRD45 |
| ENSSSCG00000032691 | ankyrin repeat domain 66 [Source:HGNC Symbol;Acc:HGNC:44669] | ANKRD66 |
| ENSSSCG00000024158 | anoctamin 1 [Source:HGNC Symbol;Acc:HGNC:21625] | ANO1 |
| ENSSSCG00000008336 | annexin A4 [Source:RefSeq peptide;Acc:NP_001161111] | ANXA4 |
| ENSSSCG00000017712 | adaptor related protein complex 2 beta 1 subunit [Source:HGNC Symbol;Acc:HGNC:563] | AP2B1 |
| ENSSSCG00000000668 | apolipoprotein B mRNA editing enzyme catalytic subunit 1 | APOBEC1 |
| ENSSSCG00000014880 | aquaporin-11 [Source:RefSeq peptide;Acc:NP_001106152] | AQP11 |
| ENSSSCG00000004195 | arginase 1 [Source:HGNC Symbol;Acc:HGNC:663] | ARG1 |
| ENSSSCG00000015015 | Rho GTPase activating protein 20 [Source:HGNC Symbol;Acc:HGNC:18357] | ARHGAP20 |
| ENSSSCG00000012112 | Rho GTPase activating protein 6 [Source:HGNC Symbol;Acc:HGNC:676] | ARHGAP6 |
| ENSSSCG00000003486 | Rho guanine nucleotide exchange factor 10 like [Source:HGNC Symbol;Acc:HGNC:25540] | ARHGEF10L |
| ENSSSCG00000006502 | Rho/Rac guanine nucleotide exchange factor 2 [Source:HGNC Symbol;Acc:HGNC:682] | ARHGEF2 |
| ENSSSCG00000030359 | Rho guanine nucleotide exchange factor 3 [Source:HGNC Symbol;Acc:HGNC:683] | ARHGEF3 |
| ENSSSCG00000021764 | Rho guanine nucleotide exchange factor 38 [Source:HGNC Symbol;Acc:HGNC:25968] | ARHGEF38 |
| ENSSSCG00000027278 | ADP ribosylation factor like GTPase 6 [Source:HGNC Symbol;Acc:HGNC:13210] | ARL6 |
| ENSSSCG00000011078 | armadillo repeat containing 3 [Source:HGNC Symbol;Acc:HGNC:30964] | ARMC3 |
| ENSSSCG00000011069 | armadillo repeat containing 4 [Source:HGNC Symbol;Acc:HGNC:25583] | ARMC4 |
| ENSSSCG00000013397 | aryl hydrocarbon receptor nuclear translocator like [Source:HGNC Symbol;Acc:HGNC:701] | ARNTL |
| ENSSSCG00000023130 | activating signal cointegrator 1 complex subunit 1 [Source:HGNC Symbol;Acc:HGNC:24268] | ASCC1 |
| ENSSSCG00000017932 | asialoglycoprotein receptor 2 [Source:HGNC Symbol;Acc:HGNC:743] | ASGR2 |
| ENSSSCG00000008072 | asporin [Source:HGNC Symbol;Acc:HGNC:14872] | ASPN |
| ENSSSCG00000021527 | ATPase phospholipid transporting 10B (putative) [Source:HGNC Symbol;Acc:HGNC:13543] | ATP10B |
| ENSSSCG00000023084 | ATPase plasma membrane Ca2+ transporting 2 [Source:HGNC Symbol;Acc:HGNC:815] | ATP2B2 |
| ENSSSCG00000002671 | ATPase secretory pathway Ca2+ transporting 2 [Source:HGNC Symbol;Acc:HGNC:29103] | ATP2C2 |
| ENSSSCG00000015528 | axonemal dynein light chain domain containing 1 [Source:HGNC Symbol;Acc:HGNC:26564] | AXDND1 |
| ENSSSCG00000003619 | antizyme inhibitor 2 [Source:RefSeq peptide;Acc:NP_001116665] | AZIN2 |
| ENSSSCG00000032536 | UDP-GlcNAc:betaGal beta-1,3-N-acetylglucosaminyltransferase 8 | B3GNT8 |
| ENSSSCG00000011014 | BMP and activin membrane-bound inhibitor homolog precursor | BAMBI |
| ENSSSCG00000002356 | basal body orientation factor 1 [Source:HGNC Symbol;Acc:HGNC:19855] | BBOF1 |
| ENSSSCG00000025417 | Bardet-Biedl syndrome 2 [Source:HGNC Symbol;Acc:HGNC:967] | BBS2 |
| ENSSSCG00000015931 | Bardet-Biedl syndrome 5 [Source:HGNC Symbol;Acc:HGNC:970] | BBS5 |
| ENSSSCG00000007485 | breast carcinoma amplified sequence 1 [Source:HGNC Symbol;Acc:HGNC:974] | BCAS1 |
| ENSSSCG00000024109 | 3-hydroxybutyrate dehydrogenase 1 [Source:HGNC Symbol;Acc:HGNC:1027] | BDH1 |
| ENSSSCG00000012137 | BMX non-receptor tyrosine kinase [Source:HGNC Symbol;Acc:HGNC:1079] | BMX |
| ENSSSCG00000010563 | beta-transducin repeat containing E3 ubiquitin protein ligase | BTRC |
| ENSSSCG00000015035 | chromosome 11 open reading frame 52 [Source:HGNC Symbol;Acc:HGNC:30531] | C11orf52 |

| | | |
|---|---|---|
| ENSSSCG00000015021 | chromosome 11 open reading frame 88 [Source:HGNC Symbol;Acc:HGNC:25061] | C11orf88 |
| ENSSSCG00000035388 | chromosome 16 open reading frame 46 [Source:HGNC Symbol;Acc:HGNC:26525] | C16orf46 |
| ENSSSCG00000013482 | chromosome 19 open reading frame 71 [Source:HGNC Symbol;Acc:HGNC:34496] | C19orf71 |
| ENSSSCG00000031102 | chromosome 1 open reading frame 174 [Source:HGNC Symbol;Acc:HGNC:27915] | C1orf174 |
| ENSSSCG00000003951 | chromosome 1 open reading frame 210 [Source:HGNC Symbol;Acc:HGNC:28755] | C1orf210 |
| ENSSSCG00000033259 | chromosome 1 open reading frame 228 [Source:HGNC Symbol;Acc:HGNC:34345] | C1orf228 |
| ENSSSCG00000003823 | chromosome 1 open reading frame 87 [Source:HGNC Symbol;Acc:HGNC:28547] | C1orf87 |
| ENSSSCG00000000117 | chromosome 22 open reading frame 23 [Source:HGNC Symbol;Acc:HGNC:18589] | C22orf23 |
| ENSSSCG00000008410 | chromosome 2 open reading frame 73 [Source:HGNC Symbol;Acc:HGNC:26861] | C2orf73 |
| ENSSSCG00000011482 | chromosome 3 open reading frame 67 [Source:HGNC Symbol;Acc:HGNC:24763] | C3orf67 |
| ENSSSCG00000011380 | chromosome 3 open reading frame 84 [Source:HGNC Symbol;Acc:HGNC:44666] | C3orf84 |
| ENSSSCG00000015662 | Sus scrofa complement component 4 binding protein, alpha (C4BPA), mRNA. | C4BPA |
| ENSSSCG00000015794 | chromosome 4 open reading frame 47 [Source:HGNC Symbol;Acc:HGNC:34346] | C4orf47 |
| ENSSSCG00000017098 | chromosome 5 open reading frame 49 [Source:HGNC Symbol;Acc:HGNC:27028] | C5orf49 |
| ENSSSCG00000004028 | chromosome 6 open reading frame 118 [Source:HGNC Symbol;Acc:HGNC:21233] | C6orf118 |
| ENSSSCG00000034398 | chromosome 8 open reading frame 37 [Source:HGNC Symbol;Acc:HGNC:27232] | C8orf37 |
| ENSSSCG00000032228 | chromosome 8 open reading frame 89 [Source:HGNC Symbol;Acc:HGNC:51258] | C8orf89 |
| ENSSSCG00000035668 | chromosome 9 open reading frame 135 [Source:HGNC Symbol;Acc:HGNC:31422] | C9orf135 |
| ENSSSCG00000033268 | ciliary associated calcium binding coiled-coil 1 [Source:HGNC Symbol;Acc:HGNC:28678] | CABCOCO1 |
| ENSSSCG00000003257 | calcium voltage-gated channel auxiliary subunit gamma 6 | CACNG6 |
| ENSSSCG00000038886 | cancer antigen 1 [Source:HGNC Symbol;Acc:HGNC:21622] | CAGE1 |
| ENSSSCG00000025768 | Sus scrofa calmodulin 1 (CALM1), mRNA. [Source:RefSeq mRNA;Acc:NM_001244210] | CALM3 |
| ENSSSCG00000013858 | calreticulin 3 [Source:HGNC Symbol;Acc:HGNC:20407] | CALR3 |
| ENSSSCG00000011550 | calcium/calmodulin dependent protein kinase I [Source:HGNC Symbol;Acc:HGNC:1459] | CAMK1 |
| ENSSSCG00000000517 | calcyphosine 2 [Source:HGNC Symbol;Acc:HGNC:16471] | CAPS2 |
| ENSSSCG00000031959 | calcyphosine like [Source:HGNC Symbol;Acc:HGNC:28375] | CAPSL |
| ENSSSCG00000040550 | cancer susceptibility 1 [Source:HGNC Symbol;Acc:HGNC:29599] | CASC1 |
| ENSSSCG00000028157 | caspase 8 [Source:HGNC Symbol;Acc:HGNC:1509] | CASP8 |
| ENSSSCG00000014170 | calpastatin [Source:HGNC Symbol;Acc:HGNC:1515] | CAST |
| ENSSSCG00000003407 | castor zinc finger 1 [Source:HGNC Symbol;Acc:HGNC:26002] | CASZ1 |
| ENSSSCG00000003083 | Cbl proto-oncogene C [Source:HGNC Symbol;Acc:HGNC:15961] | CBLC |
| ENSSSCG00000040334 | chromobox 6 [Source:HGNC Symbol;Acc:HGNC:1556] | CBX6 |
| ENSSSCG00000015795 | coiled-coil domain containing 110 [Source:HGNC Symbol;Acc:HGNC:28504] | CCDC110 |
| ENSSSCG00000002803 | coiled-coil domain containing 113 [Source:HGNC Symbol;Acc:HGNC:25002] | CCDC113 |
| ENSSSCG00000026689 | coiled-coil domain containing 114 [Source:HGNC Symbol;Acc:HGNC:26560] | CCDC114 |
| ENSSSCG00000015109 | coiled-coil domain containing 153 [Source:HGNC Symbol;Acc:HGNC:27446] | CCDC153 |
| ENSSSCG00000010003 | coiled-coil domain containing 157 [Source:HGNC Symbol;Acc:HGNC:33854] | CCDC157 |
| ENSSSCG00000021448 | coiled-coil domain containing 17 [Source:HGNC Symbol;Acc:HGNC:26574] | CCDC17 |
| ENSSSCG00000004087 | coiled-coil domain containing 170 [Source:HGNC Symbol;Acc:HGNC:21177] | CCDC170 |
| ENSSSCG00000027678 | coiled-coil domain containing 175 [Source:HGNC Symbol;Acc:HGNC:19847] | CCDC175 |
| ENSSSCG00000001908 | coiled-coil domain containing 33 [Source:HGNC Symbol;Acc:HGNC:26552] | CCDC33 |
| ENSSSCG00000017158 | coiled-coil domain containing 40 [Source:HGNC Symbol;Acc:HGNC:26090] | CCDC40 |
| ENSSSCG00000014917 | coiled-coil domain containing 81 [Source:HGNC Symbol;Acc:HGNC:26281] | CCDC81 |
| ENSSSCG00000024232 | coiled-coil domain containing 91 [Source:HGNC Symbol;Acc:HGNC:24855] | CCDC91 |
| ENSSSCG00000022196 | chaperonin containing TCP1 subunit 6B [Source:HGNC Symbol;Acc:HGNC:1621] | CCT6B |
| ENSSSCG00000006309 | CD247 molecule [Source:HGNC Symbol;Acc:HGNC:1677] | CD247 |
| ENSSSCG00000023993 | CD96 molecule [Source:HGNC Symbol;Acc:HGNC:16892] | CD96 |
| ENSSSCG00000013018 | CDC42 binding protein kinase gamma [Source:HGNC Symbol;Acc:HGNC:29829] | CDC42BPG |
| ENSSSCG00000011298 | CUB domain containing protein 1 [Source:HGNC Symbol;Acc:HGNC:24357] | CDCP1 |
| ENSSSCG00000025652 | Sus scrofa cadherin 1 (CDH1), mRNA. [Source:RefSeq mRNA;Acc:NM_001163060] | CDH1 |
| ENSSSCG00000011391 | cadherin related family member 4 [Source:HGNC Symbol;Acc:HGNC:34527] | CDHR4 |
| ENSSSCG00000008970 | cyclin dependent kinase like 2 [Source:HGNC Symbol;Acc:HGNC:1782] | CDKL2 |
| ENSSSCG00000013476 | CUGBP Elav-like family member 5 [Source:HGNC Symbol;Acc:HGNC:14058] | CELF5 |
| ENSSSCG00000031361 | cadherin EGF LAG seven-pass G-type receptor 1 [Source:HGNC Symbol;Acc:HGNC:1850] | CELSR1 |
| ENSSSCG00000002768 | centromere protein T [Source:HGNC Symbol;Acc:HGNC:25787] | CENPT |
| ENSSSCG00000011949 | centrosomal protein 97 [Source:HGNC Symbol;Acc:HGNC:26244] | CEP97 |
| ENSSSCG00000016010 | ceramide kinase like [Source:HGNC Symbol;Acc:HGNC:21699] | CERKL |
| ENSSSCG00000006644 | ceramide synthase 2 [Source:HGNC Symbol;Acc:HGNC:14076] | CERS2 |
| ENSSSCG00000024476 | carboxylesterase 3 precursor [Source:RefSeq peptide;Acc:NP_001230554] | CES3 |
| ENSSSCG00000023957 | cilia and flagella associated protein 126 [Source:HGNC Symbol;Acc:HGNC:32325] | CFAP126 |
| ENSSSCG00000005618 | cilia and flagella associated protein 157 [Source:HGNC Symbol;Acc:HGNC:27843] | CFAP157 |
| ENSSSCG00000001786 | cilia and flagella associated protein 161 [Source:HGNC Symbol;Acc:HGNC:26782] | CFAP161 |
| ENSSSCG00000002805 | cilia- and flagella-associated protein 20 [Source:RefSeq peptide;Acc:NP_001231715] | CFAP20 |

| | | |
|---|---|---|
| ENSSSCG00000015741 | cilia and flagella associated protein 221 [Source:HGNC Symbol;Acc:HGNC:33720] | CFAP221 |
| ENSSSCG00000010768 | cilia and flagella associated protein 46 [Source:HGNC Symbol;Acc:HGNC:25247] | CFAP46 |
| ENSSSCG00000017994 | cilia and flagella associated protein 52 [Source:HGNC Symbol;Acc:HGNC:16053] | CFAP52 |
| ENSSSCG00000033574 | cilia and flagella associated protein 54 [Source:HGNC Symbol;Acc:HGNC:26456] | CFAP54 |
| ENSSSCG00000023904 | cilia and flagella associated protein 57 [Source:HGNC Symbol;Acc:HGNC:26485] | CFAP57 |
| ENSSSCG00000010614 | cilia and flagella associated protein 58 [Source:HGNC Symbol;Acc:HGNC:26676] | CFAP58 |
| ENSSSCG00000016203 | cilia and flagella associated protein 65 [Source:HGNC Symbol;Acc:HGNC:25325] | CFAP65 |
| ENSSSCG00000015303 | cilia and flagella associated protein 69 [Source:HGNC Symbol;Acc:HGNC:26107] | CFAP69 |
| ENSSSCG00000010298 | cilia and flagella associated protein 70 [Source:HGNC Symbol;Acc:HGNC:30726] | CFAP70 |
| ENSSSCG00000030513 | cilia and flagella associated protein 74 [Source:HGNC Symbol;Acc:HGNC:29368] | CFAP74 |
| ENSSSCG00000005727 | cilia and flagella associated protein 77 [Source:HGNC Symbol;Acc:HGNC:33776] | CFAP77 |
| ENSSSCG00000034633 | cilia and flagella associated protein 99 [Source:HGNC Symbol;Acc:HGNC:51180] | CFAP99 |
| ENSSSCG00000016626 | cystic fibrosis transmembrane conductance regulator [Source:HGNC Symbol;Acc:HGNC:1884] | CFTR |
| ENSSSCG00000021363 | chromodomain helicase DNA binding protein 3 [Source:HGNC Symbol;Acc:HGNC:1918] | CHD3 |
| ENSSSCG00000025980 | charged multivesicular body protein 4C [Source:HGNC Symbol;Acc:HGNC:30599] | CHMP4C |
| ENSSSCG00000001989 | cell death-inducing DFFA-like effector b [Source:HGNC Symbol;Acc:HGNC:1977] | CIDEB |
| ENSSSCG00000013427 | cold inducible RNA binding protein [Source:HGNC Symbol;Acc:HGNC:1982] | CIRBP |
| ENSSSCG00000008089 | cytoskeleton associated protein 2 like [Source:HGNC Symbol;Acc:HGNC:26877] | CKAP2L |
| ENSSSCG00000008402 | clathrin heavy chain linker domain containing 1 [Source:HGNC Symbol;Acc:HGNC:26453] | CLHC1 |
| ENSSSCG00000007955 | clusterin associated protein 1 [Source:HGNC Symbol;Acc:HGNC:19009] | CLUAP1 |
| ENSSSCG00000032060 | C-X9-C motif containing 2 [Source:HGNC Symbol;Acc:HGNC:24447] | CMC2 |
| ENSSSCG00000022618 | CKLF like MARVEL transmembrane domain containing 8 | CMTM8 |
| ENSSSCG00000012164 | connector enhancer of kinase suppressor of Ras 2 [Source:HGNC Symbol;Acc:HGNC:19701] | CNKSR2 |
| ENSSSCG00000013547 | crumbs 3, cell polarity complex component [Source:HGNC Symbol;Acc:HGNC:20237] | CRB3 |
| ENSSSCG00000016690 | cAMP responsive element binding protein 5 [Source:HGNC Symbol;Acc:HGNC:16844] | CREB5 |
| ENSSSCG00000036261 | ciliary rootlet coiled-coil, rootletin family member 2 [Source:HGNC Symbol;Acc:HGNC:51677] | CROCC2 |
| ENSSSCG00000004371 | crystallin beta-gamma domain containing 1 [Source:HGNC Symbol;Acc:HGNC:356] | CRYBG1 |
| ENSSSCG00000007774 | cardiotrophin 1 [Source:HGNC Symbol;Acc:HGNC:2499] | CTF1 |
| ENSSSCG00000012970 | cathepsin W [Source:HGNC Symbol;Acc:HGNC:2546] | CTSW |
| ENSSSCG00000012249 | chromosome X open reading frame 38 [Source:HGNC Symbol;Acc:HGNC:28589] | CXorf38 |
| ENSSSCG00000012549 | chromosome X open reading frame 57 [Source:HGNC Symbol;Acc:HGNC:25486] | CXorf57 |
| ENSSSCG00000000038 | cytochrome b5 reductase 3 [Source:HGNC Symbol;Acc:HGNC:2873] | CYB5R3 |
| ENSSSCG00000003825 | Sus scrofa cytochrome P450, family 2, subfamily J, polypeptide 34 (CYP2J34), mRNA. | CYP2J34 |
| ENSSSCG00000016255 | dynein assembly factor with WD repeats 1 [Source:HGNC Symbol;Acc:HGNC:26383] | DAW1 |
| ENSSSCG00000011977 | discoidin, CUB and LCCL domain containing 2 [Source:HGNC Symbol;Acc:HGNC:24627] | DCBLD2 |
| ENSSSCG00000013326 | doublecortin domain containing 1 [Source:HGNC Symbol;Acc:HGNC:20625] | DCDC1 |
| ENSSSCG00000008942 | deoxycytidine kinase [Source:HGNC Symbol;Acc:HGNC:2704] | DCK |
| ENSSSCG00000006783 | DEAD-box helicase 20 [Source:HGNC Symbol;Acc:HGNC:2743] | DDX20 |
| ENSSSCG00000002640 | differentially expressed in FDCP 8 homolog [Source:HGNC Symbol;Acc:HGNC:25969] | DEF8 |
| ENSSSCG00000005585 | DENN domain containing 1A [Source:HGNC Symbol;Acc:HGNC:29324] | DENND1A |
| ENSSSCG00000013546 | DENN domain containing 1C [Source:HGNC Symbol;Acc:HGNC:26225] | DENND1C |
| ENSSSCG00000025704 | DENN domain containing 2D [Source:HGNC Symbol;Acc:HGNC:26192] | DENND2D |
| ENSSSCG00000014943 | deuterosome assembly protein 1 [Source:HGNC Symbol;Acc:HGNC:26344] | DEUP1 |
| ENSSSCG00000016521 | diacylglycerol kinase iota [Source:HGNC Symbol;Acc:HGNC:2855] | DGKI |
| ENSSSCG00000028225 | dicer 1, ribonuclease III [Source:HGNC Symbol;Acc:HGNC:17098] | DICER1 |
| ENSSSCG00000011249 | deleted in lung and esophageal cancer 1 [Source:HGNC Symbol;Acc:HGNC:2899] | DLEC1 |
| ENSSSCG00000004608 | dynein axonemal assembly factor 4 [Source:HGNC Symbol;Acc:HGNC:21493] | DNAAF4 |
| ENSSSCG00000028184 | dynein axonemal heavy chain 1 [Source:HGNC Symbol;Acc:HGNC:2940] | DNAH1 |
| ENSSSCG00000009765 | dynein axonemal heavy chain 10 [Source:HGNC Symbol;Acc:HGNC:2941] | DNAH10 |
| ENSSSCG00000031798 | dynein axonemal heavy chain 3 [Source:HGNC Symbol;Acc:HGNC:2949] | DNAH3 |
| ENSSSCG00000032814 | dynein axonemal heavy chain 7 [Source:HGNC Symbol;Acc:HGNC:18661] | DNAH7 |
| ENSSSCG00000018015 | dynein axonemal heavy chain 9 [Source:HGNC Symbol;Acc:HGNC:2953] | DNAH9 |
| ENSSSCG00000024357 | dynein axonemal intermediate chain 2 [Source:HGNC Symbol;Acc:HGNC:18744] | DNAI2 |
| ENSSSCG00000039469 | dynein axonemal light chain 1 [Source:HGNC Symbol;Acc:HGNC:23247] | DNAL1 |
| ENSSSCG00000000093 | dynein axonemal light chain 4 [Source:HGNC Symbol;Acc:HGNC:2955] | DNAL4 |
| ENSSSCG00000039034 | dynein axonemal light intermediate chain 1 [Source:HGNC Symbol;Acc:HGNC:14353] | DNALI1 |
| ENSSSCG00000007488 | docking protein 5 [Source:RefSeq peptide;Acc:NP_001116579] | DOK5 |
| ENSSSCG00000015711 | dipeptidyl peptidase like 10 [Source:HGNC Symbol;Acc:HGNC:20823] | DPP10 |
| ENSSSCG00000003423 | dorsal inhibitory axon guidance protein [Source:HGNC Symbol;Acc:HGNC:25054] | DRAXIN |
| ENSSSCG00000008568 | dynein regulatory complex subunit 1 [Source:HGNC Symbol;Acc:HGNC:24245] | DRC1 |
| ENSSSCG00000037557 | dynein regulatory complex subunit 3 [Source:HGNC Symbol;Acc:HGNC:25384] | DRC3 |
| ENSSSCG00000002817 | dynein regulatory complex subunit 7 [Source:HGNC Symbol;Acc:HGNC:25289] | DRC7 |
| ENSSSCG00000015048 | dopamine receptor D2 [Source:HGNC Symbol;Acc:HGNC:3023] | DRD2 |

| | | |
|---|---|---|
| ENSSSCG00000024800 | desmocollin 3 [Source:HGNC Symbol;Acc:HGNC:3037] | DSC3 |
| ENSSSCG00000015082 | DS cell adhesion molecule like 1 [Source:HGNC Symbol;Acc:HGNC:14656] | DSCAML1 |
| ENSSSCG00000022739 | desmoglein 2 [Source:HGNC Symbol;Acc:HGNC:3049] | DSG2 |
| ENSSSCG00000001025 | desmoplakin [Source:HGNC Symbol;Acc:HGNC:3052] | DSP |
| ENSSSCG00000028471 | death domain containing 1 [Source:HGNC Symbol;Acc:HGNC:37261] | DTHD1 |
| ENSSSCG00000000449 | deltex E3 ubiquitin ligase 3 [Source:HGNC Symbol;Acc:HGNC:24457] | DTX3 |
| ENSSSCG00000010339 | DPY30 domain containing 2 [Source:HGNC Symbol;Acc:HGNC:23468] | DYDC2 |
| ENSSSCG00000007094 | double zinc ribbon and ankyrin repeat domains 1 [Source:HGNC Symbol;Acc:HGNC:15858] | DZANK1 |
| ENSSSCG00000009501 | DAZ interacting zinc finger protein 1 [Source:HGNC Symbol;Acc:HGNC:20908] | DZIP1 |
| ENSSSCG00000027467 | DAZ interacting zinc finger protein 1 like [Source:HGNC Symbol;Acc:HGNC:26551] | DZIP1L |
| ENSSSCG00000004147 | epithelial cell transforming 2 like [Source:HGNC Symbol;Acc:HGNC:21118] | ECT2L |
| ENSSSCG00000007839 | eukaryotic elongation factor 2 kinase [Source:HGNC Symbol;Acc:HGNC:24615] | EEF2K |
| ENSSSCG00000032887 | EF-hand calcium binding domain 1 [Source:HGNC Symbol;Acc:HGNC:25678] | EFCAB1 |
| ENSSSCG00000002430 | EF-hand calcium binding domain 11 [Source:HGNC Symbol;Acc:HGNC:20357] | EFCAB11 |
| ENSSSCG00000003814 | EF-hand calcium binding domain 7 [Source:HGNC Symbol;Acc:HGNC:29379] | EFCAB7 |
| ENSSSCG00000008397 | EGF containing fibulin like extracellular matrix protein 1 [Source:HGNC Symbol;Acc:HGNC:3218] | EFEMP1 |
| ENSSSCG00000011205 | EF-hand domain family member B [Source:HGNC Symbol;Acc:HGNC:26330] | EFHB |
| ENSSSCG00000002620 | EF-hand domain containing 1 [Source:HGNC Symbol;Acc:HGNC:16406] | EFHC1 |
| ENSSSCG00000021846 | EF-hand domain containing 2 [Source:HGNC Symbol;Acc:HGNC:26233] | EFHC2 |
| ENSSSCG00000029484 | embigin [Source:HGNC Symbol;Acc:HGNC:30465] | EMB |
| ENSSSCG00000038080 | endomucin [Source:HGNC Symbol;Acc:HGNC:16041] | EMCN |
| ENSSSCG00000032929 | enkurin, TRPC channel interacting protein [Source:HGNC Symbol;Acc:HGNC:28388] | ENKUR |
| ENSSSCG00000010664 | enolase family member 4 [Source:HGNC Symbol;Acc:HGNC:31670] | ENO4 |
| ENSSSCG00000027169 | enolase superfamily member 1 [Source:HGNC Symbol;Acc:HGNC:30365] | ENOSF1 |
| ENSSSCG00000012663 | ecto-NOX disulfide-thiol exchanger 2 [Source:HGNC Symbol;Acc:HGNC:2259] | ENOX2 |
| ENSSSCG00000006001 | ectonucleotide pyrophosphatase/phosphodiesterase 2 [Source:HGNC Symbol;Acc:HGNC:3357] | ENPP2 |
| ENSSSCG00000025788 | ectonucleotide pyrophosphatase/phosphodiesterase 4 [Source:HGNC Symbol;Acc:HGNC:3359] | ENPP4 |
| ENSSSCG00000001715 | ectonucleotide pyrophosphatase/phosphodiesterase 5 (putative) | ENPP5 |
| ENSSSCG00000003586 | erythrocyte membrane protein band 4.1 [Source:HGNC Symbol;Acc:HGNC:3377] | EPB41 |
| ENSSSCG00000005446 | erythrocyte membrane protein band 4.1 like 4B [Source:HGNC Symbol;Acc:HGNC:19818] | EPB41L4B |
| ENSSSCG00000028144 | epoxide hydrolase 3 [Source:HGNC Symbol;Acc:HGNC:23760] | EPHX3 |
| ENSSSCG00000032203 | epiplakin 1 [Source:HGNC Symbol;Acc:HGNC:15577] | EPPK1 |
| ENSSSCG00000022706 | glutamate rich 6 [Source:HGNC Symbol;Acc:HGNC:28602] | ERICH6 |
| ENSSSCG00000016782 | family with sequence similarity 105 member A [Source:HGNC Symbol;Acc:HGNC:25629] | FAM105A |
| ENSSSCG00000007200 | family with sequence similarity 110 member A [Source:HGNC Symbol;Acc:HGNC:16188] | FAM110A |
| ENSSSCG00000017551 | family with sequence similarity 117 member A [Source:HGNC Symbol;Acc:HGNC:24179] | FAM117A |
| ENSSSCG00000008378 | family with sequence similarity 161 member A [Source:HGNC Symbol;Acc:HGNC:25808] | FAM161A |
| ENSSSCG00000002353 | family with sequence similarity 161 member B [Source:HGNC Symbol;Acc:HGNC:19854] | FAM161B |
| ENSSSCG00000032852 | family with sequence similarity 167 member A [Source:HGNC Symbol;Acc:HGNC:15549] | FAM167A |
| ENSSSCG00000008004 | family with sequence similarity 173 member A [Source:HGNC Symbol;Acc:HGNC:14152] | FAM173A |
| ENSSSCG00000004617 | family with sequence similarity 214 member A [Source:HGNC Symbol;Acc:HGNC:25609] | FAM214A |
| ENSSSCG00000009822 | protein FAM216A [Source:RefSeq peptide;Acc:NP_001231370] | FAM216A |
| ENSSSCG00000009428 | family with sequence similarity 216 member B [Source:HGNC Symbol;Acc:HGNC:26883] | FAM216B |
| ENSSSCG00000005324 | family with sequence similarity 221 member B [Source:HGNC Symbol;Acc:HGNC:30762] | FAM221B |
| ENSSSCG00000036621 | family with sequence similarity 229 member B [Source:HGNC Symbol;Acc:HGNC:33858] | FAM229B |
| ENSSSCG00000026175 | family with sequence similarity 81 member B [Source:HGNC Symbol;Acc:HGNC:26335] | FAM81B |
| ENSSSCG00000036964 | family with sequence similarity 83 member H [Source:HGNC Symbol;Acc:HGNC:24797] | FAM83H |
| ENSSSCG00000005967 | family with sequence similarity 84 member B [Source:HGNC Symbol;Acc:HGNC:24166] | FAM84B |
| ENSSSCG00000002668 | family with sequence similarity 92 member B [Source:HGNC Symbol;Acc:HGNC:24781] | FAM92B |
| ENSSSCG00000001837 | Fanconi anemia complementation group I [Source:HGNC Symbol;Acc:HGNC:25568] | FANCI |
| ENSSSCG00000010745 | fibronectin type III and ankyrin repeat domains 1 [Source:HGNC Symbol;Acc:HGNC:23527] | FANK1 |
| ENSSSCG00000039802 | F-box and leucine rich repeat protein 2 [Source:HGNC Symbol;Acc:HGNC:13598] | FBXL2 |
| ENSSSCG00000004873 | F-box protein 15 [Source:HGNC Symbol;Acc:HGNC:13617] | FBXO15 |
| ENSSSCG00000009678 | F-box protein 16 [Source:HGNC Symbol;Acc:HGNC:13618] | FBXO16 |
| ENSSSCG00000013725 | F-box and WD repeat domain containing 9 [Source:HGNC Symbol;Acc:HGNC:28136] | FBXW9 |
| ENSSSCG00000007052 | fermitin family member 1 [Source:HGNC Symbol;Acc:HGNC:15889] | FERMT1 |
| ENSSSCG00000010478 | free fatty acid receptor 4 [Source:HGNC Symbol;Acc:HGNC:19061] | FFAR4 |
| ENSSSCG00000004023 | FGFR1 oncogene partner [Source:HGNC Symbol;Acc:HGNC:17012] | FGFR1OP |
| ENSSSCG00000003828 | FGGY carbohydrate kinase domain containing [Source:HGNC Symbol;Acc:HGNC:25610] | FGGY |
| ENSSSCG00000003455 | forkhead associated phosphopeptide binding domain 1 | FHAD1 |
| ENSSSCG00000006702 | flavin containing monooxygenase 5 [Source:HGNC Symbol;Acc:HGNC:3773] | FMO5 |
| ENSSSCG00000022159 | fibronectin type III domain containing 3A [Source:HGNC Symbol;Acc:HGNC:20296] | FNDC3A |
| ENSSSCG00000014812 | folate receptor beta [Source:HGNC Symbol;Acc:HGNC:3793] | FOLR2 |

| | | |
|---|---|---|
| ENSSSCG00000017187 | forkhead box J1 [Source:HGNC Symbol;Acc:HGNC:3816] | FOXJ1 |
| ENSSSCG00000004434 | fyn related Src family tyrosine kinase [Source:HGNC Symbol;Acc:HGNC:3955] | FRK |
| ENSSSCG00000010379 | FERM and PDZ domain containing 2 [Source:HGNC Symbol;Acc:HGNC:28572] | FRMPD2 |
| ENSSSCG00000005426 | fibronectin type III and SPRY domain containing 1 like [Source:HGNC Symbol;Acc:HGNC:13753] | FSD1L |
| ENSSSCG00000040418 | fibrous sheath interacting protein 1 [Source:HGNC Symbol;Acc:HGNC:21674] | FSIP1 |
| ENSSSCG00000004651 | galactokinase 2 [Source:HGNC Symbol;Acc:HGNC:4119] | GALK2 |
| ENSSSCG00000015907 | polypeptide N-acetylgalactosaminyltransferase 3 [Source:HGNC Symbol;Acc:HGNC:4125] | GALNT3 |
| ENSSSCG00000004729 | glucosidase alpha, neutral C [Source:HGNC Symbol;Acc:HGNC:4139] | GANC |
| ENSSSCG00000039395 | growth arrest specific 2 like 2 [Source:HGNC Symbol;Acc:HGNC:24846] | GAS2L2 |
| ENSSSCG00000002633 | growth arrest specific 8 [Source:HGNC Symbol;Acc:HGNC:4166] | GAS8 |
| ENSSSCG00000016751 | glucokinase [Source:HGNC Symbol;Acc:HGNC:4195] | GCK |
| ENSSSCG00000035582 | GIPC PDZ domain containing family member 2 [Source:HGNC Symbol;Acc:HGNC:18177] | GIPC2 |
| ENSSSCG00000015258 | galactosidase beta 1 like 2 [Source:HGNC Symbol;Acc:HGNC:25129] | GLB1L2 |
| ENSSSCG00000007931 | glyoxylate reductase 1 homolog [Source:HGNC Symbol;Acc:HGNC:24434] | GLYR1 |
| ENSSSCG00000001095 | geminin, DNA replication inhibitor [Source:HGNC Symbol;Acc:HGNC:17493] | GMNN |
| ENSSSCG00000038558 | golgi phosphoprotein 3 like [Source:HGNC Symbol;Acc:HGNC:24882] | GOLPH3L |
| ENSSSCG00000040989 | G protein-coupled receptor class C group 5 member C | GPRC5C |
| ENSSSCG00000011915 | GRAM domain containing 1C [Source:HGNC Symbol;Acc:HGNC:25252] | GRAMD1C |
| ENSSSCG00000031441 | GRAM domain containing 4 [Source:HGNC Symbol;Acc:HGNC:29113] | GRAMD4 |
| ENSSSCG00000017495 | growth factor receptor bound protein 7 [Source:HGNC Symbol;Acc:HGNC:4567] | GRB7 |
| ENSSSCG00000012638 | glutamate ionotropic receptor AMPA type subunit 3 [Source:HGNC Symbol;Acc:HGNC:4573] | GRIA3 |
| ENSSSCG00000009956 | G protein-coupled receptor kinase 3 [Source:HGNC Symbol;Acc:HGNC:290] | GRK3 |
| ENSSSCG00000011423 | glutamate metabotropic receptor 2 [Source:HGNC Symbol;Acc:HGNC:4594] | GRM2 |
| ENSSSCG00000037508 | gelsolin [Source:HGNC Symbol;Acc:HGNC:4620] | GSN |
| ENSSSCG00000001978 | granzyme B precursor [Source:RefSeq peptide;Acc:NP_001137182] | GZMB |
| ENSSSCG00000011192 | 2-hydroxyacyl-CoA lyase 1 [Source:HGNC Symbol;Acc:HGNC:17856] | HACL1 |
| ENSSSCG00000008003 | hydroxyacylglutathione hydrolase like [Source:HGNC Symbol;Acc:HGNC:14177] | HAGHL |
| ENSSSCG00000028031 | histone deacetylase 11 [Source:HGNC Symbol;Acc:HGNC:19086] | HDAC11 |
| ENSSSCG00000030548 | HECT and RLD domain containing E3 ubiquitin protein ligase 5 | HERC5 |
| ENSSSCG00000006760 | homeodomain interacting protein kinase 1 [Source:HGNC Symbol;Acc:HGNC:19006] | HIPK1 |
| ENSSSCG00000001459 | major histocompatibility complex, class II, DO beta [Source:HGNC Symbol;Acc:HGNC:4937] | HLA-DOB |
| ENSSSCG00000034242 | hepatocyte nuclear factor 4 gamma [Source:HGNC Symbol;Acc:HGNC:5026] | HNF4G |
| ENSSSCG00000017534 | homeobox B3 [Source:HGNC Symbol;Acc:HGNC:5114] | HOXB3 |
| ENSSSCG00000038993 | homeobox C4 [Source:HGNC Symbol;Acc:HGNC:5126] | HOXC4 |
| ENSSSCG00000032664 | hippocalcin [Source:HGNC Symbol;Acc:HGNC:5144] | HPCA |
| ENSSSCG00000009699 | 15-hydroxyprostaglandin dehydrogenase [Source:HGNC Symbol;Acc:HGNC:5154] | HPGD |
| ENSSSCG00000009237 | heparanase precursor [Source:RefSeq peptide;Acc:NP_001139602] | HPSE |
| ENSSSCG00000001476 | hydroxysteroid 17-beta dehydrogenase 8 [Source:HGNC Symbol;Acc:HGNC:3554] | HSD17B8 |
| ENSSSCG00000040973 | HYDIN, axonemal central pair apparatus protein [Source:HGNC Symbol;Acc:HGNC:19368] | HYDIN |
| ENSSSCG00000039909 | intercellular adhesion molecule 2 precursor [Source:RefSeq peptide;Acc:NP_001001631] | ICAM2 |
| ENSSSCG00000011589 | intraflagellar transport 122 [Source:HGNC Symbol;Acc:HGNC:13556] | IFT122 |
| ENSSSCG00000008020 | intraflagellar transport 140 [Source:HGNC Symbol;Acc:HGNC:29077] | IFT140 |
| ENSSSCG00000026367 | intraflagellar transport 172 [Source:HGNC Symbol;Acc:HGNC:30391] | IFT172 |
| ENSSSCG00000029649 | intraflagellar transport 46 [Source:HGNC Symbol;Acc:HGNC:26146] | IFT46 |
| ENSSSCG00000009270 | intraflagellar transport 88 [Source:HGNC Symbol;Acc:HGNC:20606] | IFT88 |
| ENSSSCG00000016164 | IKAROS family zinc finger 2 [Source:HGNC Symbol;Acc:HGNC:13177] | IKZF2 |
| ENSSSCG00000038848 | interleukin 22 receptor subunit alpha 1 [Source:HGNC Symbol;Acc:HGNC:13700] | IL22RA1 |
| ENSSSCG00000006161 | interleukin 7 [Source:HGNC Symbol;Acc:HGNC:6023] | IL7 |
| ENSSSCG00000034987 | IQ motif and ankyrin repeat containing 1 [Source:HGNC Symbol;Acc:HGNC:49576] | IQANK1 |
| ENSSSCG00000016321 | IQ motif containing with AAA domain 1 [Source:HGNC Symbol;Acc:HGNC:26195] | IQCA1 |
| ENSSSCG00000011881 | IQ motif containing B1 [Source:HGNC Symbol;Acc:HGNC:28949] | IQCB1 |
| ENSSSCG00000009877 | IQ motif containing D [Source:HGNC Symbol;Acc:HGNC:25168] | IQCD |
| ENSSSCG00000007568 | IQ motif containing E [Source:HGNC Symbol;Acc:HGNC:29171] | IQCE |
| ENSSSCG00000011855 | IQ motif containing G [Source:HGNC Symbol;Acc:HGNC:25251] | IQCG |
| ENSSSCG00000034963 | IQ motif containing H [Source:HGNC Symbol;Acc:HGNC:25721] | IQCH |
| ENSSSCG00000033381 | interferon regulatory factor 4 [Source:HGNC Symbol;Acc:HGNC:6119] | IRF4 |
| ENSSSCG00000015612 | interferon regulatory factor 6 [Source:HGNC Symbol;Acc:HGNC:6121] | IRF6 |
| ENSSSCG00000009071 | jade family PHD finger 1 [Source:HGNC Symbol;Acc:HGNC:30027] | JADE1 |
| ENSSSCG00000015089 | junction adhesion molecule like [Source:HGNC Symbol;Acc:HGNC:19084] | JAML |
| ENSSSCG00000015138 | junctional cadherin complex regulator [Source:HGNC Symbol;Acc:HGNC:26288] | JHY |
| ENSSSCG00000022643 | katanin catalytic subunit A1 like 2 [Source:HGNC Symbol;Acc:HGNC:25387] | KATNAL2 |
| ENSSSCG00000000203 | potassium voltage-gated channel subfamily H member 3 | KCNH3 |
| ENSSSCG00000023451 | potassium two pore domain channel subfamily K member 2 | KCNK2 |

| | | |
|---|---|---|
| ENSSSCG00000007814 | KIAA0556 [Source:HGNC Symbol;Acc:HGNC:29068] | KIAA0556 |
| ENSSSCG00000016491 | KIAA1147 [Source:HGNC Symbol;Acc:HGNC:29472] | KIAA1147 |
| ENSSSCG00000011075 | KIAA1217 [Source:HGNC Symbol;Acc:HGNC:25428] | KIAA1217 |
| ENSSSCG00000006969 | KIAA1456 [Source:HGNC Symbol;Acc:HGNC:26725] | KIAA1456 |
| ENSSSCG00000004899 | KIAA1468 [Source:HGNC Symbol;Acc:HGNC:29289] | KIAA1468 |
| ENSSSCG00000013311 | KIAA1549 like [Source:HGNC Symbol;Acc:HGNC:24836] | KIAA1549L |
| ENSSSCG00000008386 | KIAA1841 [Source:HGNC Symbol;Acc:HGNC:29387] | KIAA1841 |
| ENSSSCG00000016110 | KIAA2012 [Source:HGNC Symbol;Acc:HGNC:51250] | KIAA2012 |
| ENSSSCG00000003508 | kinesin family member 17 [Source:HGNC Symbol;Acc:HGNC:19167] | KIF17 |
| ENSSSCG00000010984 | kinesin family member 24 [Source:HGNC Symbol;Acc:HGNC:19916] | KIF24 |
| ENSSSCG00000021571 | kinesin family member 27 [Source:HGNC Symbol;Acc:HGNC:18632] | KIF27 |
| ENSSSCG00000007247 | kinesin family member 3B [Source:HGNC Symbol;Acc:HGNC:6320] | KIF3B |
| ENSSSCG00000040510 | kinesin family member 9 [Source:HGNC Symbol;Acc:HGNC:16666] | KIF9 |
| ENSSSCG00000012955 | kinesin light chain 2 [Source:HGNC Symbol;Acc:HGNC:20716] | KLC2 |
| ENSSSCG00000023732 | kelch domain containing 7A [Source:HGNC Symbol;Acc:HGNC:26791] | KLHDC7A |
| ENSSSCG00000038597 | kelch domain containing 9 [Source:HGNC Symbol;Acc:HGNC:28489] | KLHDC9 |
| ENSSSCG00000012637 | kelch like family member 13 [Source:HGNC Symbol;Acc:HGNC:22931] | KLHL13 |
| ENSSSCG00000010770 | kinase non-catalytic C-lobe domain containing 1 [Source:HGNC Symbol;Acc:HGNC:29374] | KNDC1 |
| ENSSSCG00000015313 | KRIT1, ankyrin repeat containing [Source:HGNC Symbol;Acc:HGNC:1573] | KRIT1 |
| ENSSSCG00000017444 | keratin 15 [Source:HGNC Symbol;Acc:HGNC:6421] | KRT15 |
| ENSSSCG00000037854 | trans-L-3-hydroxyproline dehydratase [Source:HGNC Symbol;Acc:HGNC:20488] | L3HYPDH |
| ENSSSCG00000004469 | LCA5, lebercilin [Source:HGNC Symbol;Acc:HGNC:31923] | LCA5 |
| ENSSSCG00000028122 | LCA5L, lebercilin like [Source:HGNC Symbol;Acc:HGNC:1255] | LCA5L |
| ENSSSCG00000005055 | galectin 3 [Source:HGNC Symbol;Acc:HGNC:6563] | LGALS3 |
| ENSSSCG00000027144 | lamin tail domain containing 1 [Source:HGNC Symbol;Acc:HGNC:26683] | LMNTD1 |
| ENSSSCG00000023057 | LIM domain only 1 [Source:HGNC Symbol;Acc:HGNC:6641] | LMO1 |
| ENSSSCG00000040184 | LIM domain 7 [Source:HGNC Symbol;Acc:HGNC:6646] | LMO7 |
| ENSSSCG00000008838 | ligand of numb-protein X 1 [Source:HGNC Symbol;Acc:HGNC:6657] | LNX1 |
| ENSSSCG00000001914 | lysyl oxidase like 1 [Source:HGNC Symbol;Acc:HGNC:6665] | LOXL1 |
| ENSSSCG00000009021 | LPS responsive beige-like anchor protein [Source:HGNC Symbol;Acc:HGNC:1742] | LRBA |
| ENSSSCG00000016542 | leucine rich repeats and guanylate kinase domain containing | LRGUK |
| ENSSSCG00000015792 | LRP2 binding protein [Source:HGNC Symbol;Acc:HGNC:25434] | LRP2BP |
| ENSSSCG00000000679 | leucine rich repeat containing 23 [Source:HGNC Symbol;Acc:HGNC:19138] | LRRC23 |
| ENSSSCG00000024714 | leucine rich repeat containing 34 [Source:HGNC Symbol;Acc:HGNC:28408] | LRRC34 |
| ENSSSCG00000009796 | leucine rich repeat containing 43 [Source:HGNC Symbol;Acc:HGNC:28562] | LRRC43 |
| ENSSSCG00000004976 | leucine rich repeat containing 49 [Source:HGNC Symbol;Acc:HGNC:25965] | LRRC49 |
| ENSSSCG00000005952 | leucine rich repeat containing 6 [Source:HGNC Symbol;Acc:HGNC:16725] | LRRC6 |
| ENSSSCG00000006462 | leucine rich repeat containing 71 [Source:HGNC Symbol;Acc:HGNC:26556] | LRRC71 |
| ENSSSCG00000010100 | leucine rich repeat containing 74B [Source:HGNC Symbol;Acc:HGNC:34301] | LRRC74B |
| ENSSSCG00000005081 | leucine rich repeat containing 9 [Source:HGNC Symbol;Acc:HGNC:19848] | LRRC9 |
| ENSSSCG00000003784 | leucine rich repeats and IQ motif containing 3 [Source:HGNC Symbol;Acc:HGNC:28318] | LRRIQ3 |
| ENSSSCG00000038916 | LSM10, U7 small nuclear RNA associated [Source:HGNC Symbol;Acc:HGNC:17562] | LSM10 |
| ENSSSCG00000002368 | latent transforming growth factor beta binding protein 2 [Source:HGNC Symbol;Acc:HGNC:6715] | LTBP2 |
| ENSSSCG00000011314 | leucine zipper transcription factor like 1 [Source:HGNC Symbol;Acc:HGNC:6741] | LZTFL1 |
| ENSSSCG00000011892 | MYCBP associated and testis expressed 1 [Source:HGNC Symbol;Acc:HGNC:24010] | MAATS1 |
| ENSSSCG00000040607 | MAF bZIP transcription factor [Source:HGNC Symbol;Acc:HGNC:6776] | MAF |
| ENSSSCG00000012295 | MAGI family member, X-linked [Source:HGNC Symbol;Acc:HGNC:30006] | MAGIX |
| ENSSSCG00000001042 | male germ cell associated kinase [Source:HGNC Symbol;Acc:HGNC:6816] | MAK |
| ENSSSCG00000011791 | mitogen-activated protein kinase kinase kinase 13 [Source:HGNC Symbol;Acc:HGNC:6852] | MAP3K13 |
| ENSSSCG00000015696 | mitogen-activated protein kinase kinase kinase 19 [Source:HGNC Symbol;Acc:HGNC:26249] | MAP3K19 |
| ENSSSCG00000035933 | mitogen-activated protein kinase 15 [Source:HGNC Symbol;Acc:HGNC:24667] | MAPK15 |
| ENSSSCG00000029185 | microtubule associated protein RP/EB family member 3 [Source:HGNC Symbol;Acc:HGNC:6892] | MAPRE3 |
| ENSSSCG00000013895 | microtubule associated serine/threonine kinase 3 [Source:HGNC Symbol;Acc:HGNC:19036] | MAST3 |
| ENSSSCG00000013494 | megakaryocyte-associated tyrosine kinase [Source:HGNC Symbol;Acc:HGNC:6906] | MATK |
| ENSSSCG00000003755 | mucolipin 2 [Source:HGNC Symbol;Acc:HGNC:13357] | MCOLN2 |
| ENSSSCG00000039424 | MAPK regulated corepressor interacting protein 1 [Source:HGNC Symbol;Acc:HGNC:28007] | MCRIP1 |
| ENSSSCG00000016133 | malate dehydrogenase 1B [Source:HGNC Symbol;Acc:HGNC:17836] | MDH1B |
| ENSSSCG00000029102 | mediator complex subunit 24 [Source:HGNC Symbol;Acc:HGNC:22963] | MED24 |
| ENSSSCG00000016052 | major facilitator superfamily domain containing 6 [Source:HGNC Symbol;Acc:HGNC:24711] | MFSD6 |
| ENSSSCG00000038236 | MGAT4 family member D [Source:HGNC Symbol;Acc:HGNC:43619] | MGAT4D |
| ENSSSCG00000006325 | microsomal glutathione S-transferase 3 [Source:HGNC Symbol;Acc:HGNC:7064] | MGST3 |
| ENSSSCG00000016674 | MINDY lysine 48 deubiquitinase 4 [Source:HGNC Symbol;Acc:HGNC:21916] | MINDY4 |
| ENSSSCG00000037461 | myeloid leukemia factor 1 [Source:HGNC Symbol;Acc:HGNC:7125] | MLF1 |

| | | |
|---|---|---|
| ENSSSCG00000008480 | MORN repeat containing 2 [Source:HGNC Symbol;Acc:HGNC:30166] | MORN2 |
| ENSSSCG00000009809 | MORN repeat-containing protein 3 [Source:RefSeq peptide;Acc:NP_001230638] | MORN3 |
| ENSSSCG00000005529 | MORN repeat containing 5 [Source:HGNC Symbol;Acc:HGNC:17841] | MORN5 |
| ENSSSCG00000011070 | membrane palmitoylated protein 7 [Source:HGNC Symbol;Acc:HGNC:26542] | MPP7 |
| ENSSSCG00000028126 | metallophosphoesterase 1 [Source:HGNC Symbol;Acc:HGNC:15988] | MPPE1 |
| ENSSSCG00000027992 | macrophage stimulating 1 receptor [Source:HGNC Symbol;Acc:HGNC:7381] | MST1R |
| ENSSSCG00000004167 | MYB proto-oncogene, transcription factor [Source:HGNC Symbol;Acc:HGNC:7545] | MYB |
| ENSSSCG00000017563 | MYCBP associated protein [Source:HGNC Symbol;Acc:HGNC:19677] | MYCBPAP |
| ENSSSCG00000027407 | myosin heavy chain 14 [Source:HGNC Symbol;Acc:HGNC:23212] | MYH14 |
| ENSSSCG00000033196 | myogenesis regulating glycosidase (putative) [Source:HGNC Symbol;Acc:HGNC:19918] | MYORG |
| ENSSSCG00000011330 | neurobeachin like 2 [Source:HGNC Symbol;Acc:HGNC:31928] | NBEAL2 |
| ENSSSCG00000004222 | nuclear receptor coactivator 7 [Source:HGNC Symbol;Acc:HGNC:21081] | NCOA7 |
| ENSSSCG00000011090 | nebulette [Source:HGNC Symbol;Acc:HGNC:16932] | NEBL |
| ENSSSCG00000004919 | neural precursor cell expressed, developmentally down-regulated 4-like, E3 ubiquitin protein ligase | NEDD4L |
| ENSSSCG00000009714 | NIMA related kinase 1 [Source:HGNC Symbol;Acc:HGNC:7744] | NEK1 |
| ENSSSCG00000011217 | NIMA related kinase 10 [Source:HGNC Symbol;Acc:HGNC:18592] | NEK10 |
| ENSSSCG00000011181 | NIMA related kinase 11 [Source:HGNC Symbol;Acc:HGNC:18593] | NEK11 |
| ENSSSCG00000009380 | NIMA related kinase 5 [Source:HGNC Symbol;Acc:HGNC:7748] | NEK5 |
| ENSSSCG00000004856 | nuclear factor of activated T-cells, cytoplasmic 1 [Source:RefSeq peptide;Acc:NP_999326] | NFATC1 |
| ENSSSCG00000005190 | nuclear factor I B [Source:HGNC Symbol;Acc:HGNC:7785] | NFIB |
| ENSSSCG00000004149 | NHS like 1 [Source:HGNC Symbol;Acc:HGNC:21021] | NHSL1 |
| ENSSSCG00000037950 | NIPA like domain containing 1 [Source:HGNC Symbol;Acc:HGNC:27194] | NIPAL1 |
| ENSSSCG00000027465 | neuroligin 3 [Source:HGNC Symbol;Acc:HGNC:14289] | NLGN3 |
| ENSSSCG00000006294 | NME/NM23 family member 7 [Source:HGNC Symbol;Acc:HGNC:20461] | NME7 |
| ENSSSCG00000037044 | NME/NM23 family member 9 [Source:HGNC Symbol;Acc:HGNC:21343] | NME9 |
| ENSSSCG00000015557 | nicotinamide nucleotide adenylyltransferase 2 [Source:HGNC Symbol;Acc:HGNC:16789] | NMNAT2 |
| ENSSSCG00000005272 | nicotinamide riboside kinase 1 [Source:HGNC Symbol;Acc:HGNC:26057] | NMRK1 |
| ENSSSCG00000007336 | neuronatin [Source:HGNC Symbol;Acc:HGNC:7860] | NNAT |
| ENSSSCG00000010266 | Sus scrofa neuropeptide FF receptor 1 (NPFFR1), mRNA. | NPFFR1 |
| ENSSSCG00000008111 | nephrocystin 1 [Source:HGNC Symbol;Acc:HGNC:7905] | NPHP1 |
| ENSSSCG00000021082 | neuregulin 2 [Source:HGNC Symbol;Acc:HGNC:7998] | NRG2 |
| ENSSSCG00000000741 | nuclear receptor interacting protein 2 [Source:HGNC Symbol;Acc:HGNC:23078] | NRIP2 |
| ENSSSCG00000024496 | NOP2/Sun RNA methyltransferase family member 7 [Source:HGNC Symbol;Acc:HGNC:25857] | NSUN7 |
| ENSSSCG00000035318 | nucleoporin 58 [Source:HGNC Symbol;Acc:HGNC:20261] | NUP58 |
| ENSSSCG00000021514 | occludin [Source:HGNC Symbol;Acc:HGNC:8104] | OCLN |
| ENSSSCG00000006936 | outer dense fiber of sperm tails 2 like [Source:HGNC Symbol;Acc:HGNC:29225] | ODF2L |
| ENSSSCG00000000967 | outer dense fiber of sperm tails 3B [Source:HGNC Symbol;Acc:HGNC:34388] | ODF3B |
| ENSSSCG00000009783 | 2-oxoglutarate and iron dependent oxygenase domain containing 2 | OGFOD2 |
| ENSSSCG00000031228 | nociceptin receptor [Source:RefSeq peptide;Acc:NP_999341] | OPRL1 |
| ENSSSCG00000036709 | organic solute carrier partner 1 [Source:HGNC Symbol;Acc:HGNC:29971] | OSCP1 |
| ENSSSCG00000038153 | parkin coregulated [Source:HGNC Symbol;Acc:HGNC:19152] | PACRG |
| ENSSSCG00000010437 | 3'-phosphoadenosine 5'-phosphosulfate synthase 2 [Source:HGNC Symbol;Acc:HGNC:8604] | PAPSS2 |
| ENSSSCG00000025876 | phenazine biosynthesis like protein domain containing [Source:HGNC Symbol;Acc:HGNC:23301] | PBLD |
| ENSSSCG00000033765 | protocadherin gamma subfamily A, 3 [Source:HGNC Symbol;Acc:HGNC:8701] | PCDHGA3 |
| ENSSSCG00000005082 | pecanex homolog 4 [Source:HGNC Symbol;Acc:HGNC:20349] | PCNX4 |
| ENSSSCG00000005785 | proprotein convertase subtilisin/kexin type 6 [Source:HGNC Symbol;Acc:HGNC:8569] | PCSK6 |
| ENSSSCG00000008908 | phosducin like 2 [Source:HGNC Symbol;Acc:HGNC:29524] | PDCL2 |
| ENSSSCG00000027710 | PDZ domain containing 3 [Source:HGNC Symbol;Acc:HGNC:19891] | PDZD3 |
| ENSSSCG00000033735 | PPARGC1 and ESRR induced regulator, muscle 1 [Source:HGNC Symbol;Acc:HGNC:28208] | PERM1 |
| ENSSSCG00000011439 | PHD finger protein 7 [Source:HGNC Symbol;Acc:HGNC:18458] | PHF7 |
| ENSSSCG00000015031 | PIH1 domain containing 2 [Source:HGNC Symbol;Acc:HGNC:25210] | PIH1D2 |
| ENSSSCG00000012558 | PIH1 domain containing 3 [Source:HGNC Symbol;Acc:HGNC:28570] | PIH1D3 |
| ENSSSCG00000000528 | plakophilin 2 [Source:HGNC Symbol;Acc:HGNC:9024] | PKP2 |
| ENSSSCG00000029811 | phospholipase C like 2 [Source:HGNC Symbol;Acc:HGNC:9064] | PLCL2 |
| ENSSSCG00000015281 | pleckstrin homology domain containing A6 [Source:HGNC Symbol;Acc:HGNC:17053] | PLEKHA6 |
| ENSSSCG00000013382 | pleckstrin homology domain containing A7 [Source:HGNC Symbol;Acc:HGNC:27049] | PLEKHA7 |
| ENSSSCG00000035733 | plexin B2 [Source:HGNC Symbol;Acc:HGNC:9104] | PLXNB2 |
| ENSSSCG00000029002 | paroxysmal nonkinesigenic dyskinesia [Source:HGNC Symbol;Acc:HGNC:9153] | PNKD |
| ENSSSCG00000013572 | patatin like phospholipase domain containing 6 [Source:HGNC Symbol;Acc:HGNC:16268] | PNPLA6 |
| ENSSSCG00000014598 | PPFIA binding protein 2 [Source:HGNC Symbol;Acc:HGNC:9250] | PPFIBP2 |
| ENSSSCG00000004413 | peptidylprolyl isomerase like 6 [Source:HGNC Symbol;Acc:HGNC:21557] | PPIL6 |
| ENSSSCG00000004704 | diphosphoinositol pentakisphosphate kinase 1 [Source:HGNC Symbol;Acc:HGNC:29023] | PPIP5K1 |
| ENSSSCG00000028976 | protein phosphatase 2 regulatory subunit Bbeta [Source:HGNC Symbol;Acc:HGNC:9305] | PPP2R2B |

| | | |
|---|---|---|
| ENSSSCG00000032622 | protein phosphatase 3 catalytic subunit gamma [Source:HGNC Symbol;Acc:HGNC:9316] | PPP3CC |
| ENSSSCG00000003256 | protein kinase C gamma [Source:HGNC Symbol;Acc:HGNC:9402] | PRKCG |
| ENSSSCG00000026152 | protein kinase D2 [Source:HGNC Symbol;Acc:HGNC:17293] | PRKD2 |
| ENSSSCG00000013584 | proline rich 36 [Source:HGNC Symbol;Acc:HGNC:26172] | PRR36 |
| ENSSSCG00000040698 | proline rich 7, synaptic [Source:HGNC Symbol;Acc:HGNC:28130] | PRR7 |
| ENSSSCG00000013313 | proline rich and Gla domain 4 [Source:HGNC Symbol;Acc:HGNC:30799] | PRRG4 |
| ENSSSCG00000028587 | protease, serine 54 [Source:HGNC Symbol;Acc:HGNC:26336] | PRSS54 |
| ENSSSCG00000010281 | prosaposin [Source:HGNC Symbol;Acc:HGNC:9498] | PSAP |
| ENSSSCG00000031781 | proteasome subunit beta type-10 [Source:RefSeq peptide;Acc:NP_001038030] | PSMB10 |
| ENSSSCG00000009664 | protein tyrosine kinase 2 beta [Source:HGNC Symbol;Acc:HGNC:9612] | PTK2B |
| ENSSSCG00000026655 | protein tyrosine phosphatase, non-receptor type 13 [Source:HGNC Symbol;Acc:HGNC:9646] | PTPN13 |
| ENSSSCG00000025537 | protein tyrosine phosphatase, non-receptor type 6 [Source:HGNC Symbol;Acc:HGNC:9658] | PTPN6 |
| ENSSSCG00000035849 | protein tyrosine phosphatase, receptor type F [Source:HGNC Symbol;Acc:HGNC:9670] | PTPRF |
| ENSSSCG00000031716 | protein tyrosine phosphatase, receptor type Q [Source:HGNC Symbol;Acc:HGNC:9679] | PTPRQ |
| ENSSSCG00000015428 | pseudouridylate synthase 7 (putative) [Source:HGNC Symbol;Acc:HGNC:26033] | PUS7 |
| ENSSSCG00000011670 | 2-phosphoxylose phosphatase 1 [Source:HGNC Symbol;Acc:HGNC:26303] | PXYLP1 |
| ENSSSCG00000040825 | PYM homolog 1, exon junction complex associated factor | PYM1 |
| ENSSSCG00000008491 | glutaminyl-peptide cyclotransferase [Source:HGNC Symbol;Acc:HGNC:9753] | QPCT |
| ENSSSCG00000017747 | RAB11 family interacting protein 4 [Source:HGNC Symbol;Acc:HGNC:30267] | RAB11FIP4 |
| ENSSSCG00000038693 | RAB19, member RAS oncogene family [Source:HGNC Symbol;Acc:HGNC:19982] | RAB19 |
| ENSSSCG00000004612 | RAB27A, member RAS oncogene family [Source:HGNC Symbol;Acc:HGNC:9766] | RAB27A |
| ENSSSCG00000004114 | ras-related protein Rab-32 [Source:RefSeq peptide;Acc:NP_001116648] | RAB32 |
| ENSSSCG00000010048 | RAB36, member RAS oncogene family [Source:HGNC Symbol;Acc:HGNC:9775] | RAB36 |
| ENSSSCG00000000500 | RAB3A interacting protein [Source:HGNC Symbol;Acc:HGNC:16508] | RAB3IP |
| ENSSSCG00000015499 | RAB GTPase activating protein 1 like [Source:HGNC Symbol;Acc:HGNC:24663] | RABGAP1L |
| ENSSSCG00000009824 | RAD9 checkpoint clamp component B [Source:HGNC Symbol;Acc:HGNC:21700] | RAD9B |
| ENSSSCG00000008952 | Ras association domain family member 6 [Source:HGNC Symbol;Acc:HGNC:20796] | RASSF6 |
| ENSSSCG00000038001 | ribokinase [Source:HGNC Symbol;Acc:HGNC:30325] | RBKS |
| ENSSSCG00000007504 | RNA binding motif protein 38 [Source:HGNC Symbol;Acc:HGNC:15818] | RBM38 |
| ENSSSCG00000015509 | ring finger and WD repeat domain 2 [Source:HGNC Symbol;Acc:HGNC:17440] | RFWD2 |
| ENSSSCG00000035403 | regulatory factor X2 [Source:HGNC Symbol;Acc:HGNC:9983] | RFX2 |
| ENSSSCG00000006069 | regulator of G protein signaling 22 [Source:HGNC Symbol;Acc:HGNC:24499] | RGS22 |
| ENSSSCG00000033093 | regulator of G protein signaling 6 [Source:HGNC Symbol;Acc:HGNC:10002] | RGS6 |
| ENSSSCG00000010840 | regulator of G protein signaling 7 [Source:HGNC Symbol;Acc:HGNC:10003] | RGS7 |
| ENSSSCG00000012326 | RIB43A domain with coiled-coils 1 [Source:HGNC Symbol;Acc:HGNC:26537] | RIBC1 |
| ENSSSCG00000024718 | RIB43A domain with coiled-coils 2 [Source:HGNC Symbol;Acc:HGNC:13241] | RIBC2 |
| ENSSSCG00000028004 | Ras and Rab interactor 2 [Source:HGNC Symbol;Acc:HGNC:18750] | RIN2 |
| ENSSSCG00000033276 | ripply transcriptional repressor 1 [Source:HGNC Symbol;Acc:HGNC:25117] | RIPPLY1 |
| ENSSSCG00000013765 | relaxin 3 [Source:HGNC Symbol;Acc:HGNC:17135] | RLN3 |
| ENSSSCG00000015549 | 2-5A-dependent ribonuclease [Source:RefSeq peptide;Acc:NP_001090981] | RNASEL |
| ENSSSCG00000016414 | ring finger protein 32 [Source:HGNC Symbol;Acc:HGNC:17118] | RNF32 |
| ENSSSCG00000014056 | ring finger protein 44 [Source:HGNC Symbol;Acc:HGNC:19180] | RNF44 |
| ENSSSCG00000007932 | rogdi homolog [Source:HGNC Symbol;Acc:HGNC:29478] | ROGDI |
| ENSSSCG00000022306 | rhophilin associated tail protein 1 like [Source:HGNC Symbol;Acc:HGNC:24060] | ROPN1L |
| ENSSSCG00000026427 | RAR related orphan receptor C [Source:HGNC Symbol;Acc:HGNC:10260] | RORC |
| ENSSSCG00000031652 | RPGRIP1 like [Source:HGNC Symbol;Acc:HGNC:29168] | RPGRIP1L |
| ENSSSCG00000003648 | Ras related GTP binding C [Source:HGNC Symbol;Acc:HGNC:19902] | RRAGC |
| ENSSSCG00000040224 | radial spoke head 1 homolog [Source:HGNC Symbol;Acc:HGNC:12371] | RSPH1 |
| ENSSSCG00000010047 | radial spoke head 14 homolog [Source:HGNC Symbol;Acc:HGNC:13437] | RSPH14 |
| ENSSSCG00000040714 | radial spoke head 6 homolog A [Source:HGNC Symbol;Acc:HGNC:14241] | RSPH6A |
| ENSSSCG00000032007 | reticulon 4 [Source:HGNC Symbol;Acc:HGNC:14085] | RTN4 |
| ENSSSCG00000003152 | RuvB like AAA ATPase 2 [Source:HGNC Symbol;Acc:HGNC:10475] | RUVBL2 |
| ENSSSCG00000002400 | sterile alpha motif domain containing 15 [Source:HGNC Symbol;Acc:HGNC:18631] | SAMD15 |
| ENSSSCG00000005740 | sarcosine dehydrogenase [Source:HGNC Symbol;Acc:HGNC:10536] | SARDH |
| ENSSSCG00000031963 | stabilizer of axonemal microtubules 2 [Source:HGNC Symbol;Acc:HGNC:33727] | SAXO2 |
| ENSSSCG00000007842 | short chain dehydrogenase/reductase family 42E, member 2 | SDR42E2 |
| ENSSSCG00000010008 | SEC14 like lipid binding 3 [Source:HGNC Symbol;Acc:HGNC:18655] | SEC14L3 |
| ENSSSCG00000006495 | semaphorin 4A [Source:HGNC Symbol;Acc:HGNC:10729] | SEMA4A |
| ENSSSCG00000012619 | septin 6 [Source:HGNC Symbol;Acc:HGNC:15848] | SEPT6 |
| ENSSSCG00000001011 | serpin family B member 1 [Source:HGNC Symbol;Acc:HGNC:3311] | SERPINB1 |
| ENSSSCG00000011737 | serpin family I member 2 [Source:HGNC Symbol;Acc:HGNC:8945] | SERPINI2 |
| ENSSSCG00000010589 | sideroflexin 2 [Source:HGNC Symbol;Acc:HGNC:16086] | SFXN2 |
| ENSSSCG00000010433 | sphingomyelin synthase 1 [Source:HGNC Symbol;Acc:HGNC:29799] | SGMS1 |

| | | |
|---|---|---|
| ENSSSCG00000017829 | small G protein signaling modulator 2 [Source:HGNC Symbol;Acc:HGNC:29026] | SGSM2 |
| ENSSSCG00000022380 | SH2 domain containing 3A [Source:HGNC Symbol;Acc:HGNC:16885] | SH2D3A |
| ENSSSCG00000023273 | SH3 and SYLF domain containing 1 [Source:HGNC Symbol;Acc:HGNC:29546] | SH3YL1 |
| ENSSSCG00000033043 | SH3 and multiple ankyrin repeat domains 2 [Source:HGNC Symbol;Acc:HGNC:14295] | SHANK2 |
| ENSSSCG00000024389 | single Ig IL-1-related receptor [Source:RefSeq peptide;Acc:NP_001302618] | SIGIRR |
| ENSSSCG00000002332 | signal induced proliferation associated 1 like 1 [Source:HGNC Symbol;Acc:HGNC:20284] | SIPA1L1 |
| ENSSSCG00000016708 | src kinase associated phosphoprotein 2 [Source:HGNC Symbol;Acc:HGNC:15687] | SKAP2 |
| ENSSSCG00000007437 | solute carrier family 12 member 5 [Source:HGNC Symbol;Acc:HGNC:13818] | SLC12A5 |
| ENSSSCG00000017925 | solute carrier family 16 member 11 [Source:HGNC Symbol;Acc:HGNC:23093] | SLC16A11 |
| ENSSSCG00000010455 | solute carrier family 16 member 12 [Source:HGNC Symbol;Acc:HGNC:23094] | SLC16A12 |
| ENSSSCG00000017926 | solute carrier family 16 member 13 [Source:HGNC Symbol;Acc:HGNC:31037] | SLC16A13 |
| ENSSSCG00000029458 | solute carrier family 16 member 2 [Source:HGNC Symbol;Acc:HGNC:10923] | SLC16A2 |
| ENSSSCG00000017215 | solute carrier family 16 member 5 [Source:HGNC Symbol;Acc:HGNC:10926] | SLC16A5 |
| ENSSSCG00000021968 | solute carrier family 19 member 3 [Source:HGNC Symbol;Acc:HGNC:16266] | SLC19A3 |
| ENSSSCG00000012660 | solute carrier family 25 member 14 [Source:HGNC Symbol;Acc:HGNC:10984] | SLC25A14 |
| ENSSSCG00000002520 | solute carrier family 25 member 29 [Source:HGNC Symbol;Acc:HGNC:20116] | SLC25A29 |
| ENSSSCG00000030167 | solute carrier family 25 member 39 [Source:HGNC Symbol;Acc:HGNC:24279] | SLC25A39 |
| ENSSSCG00000039045 | solute carrier family 26 member 2 [Source:HGNC Symbol;Acc:HGNC:10994] | SLC26A2 |
| ENSSSCG00000004172 | solute carrier family 2 member 12 [Source:HGNC Symbol;Acc:HGNC:18067] | SLC2A12 |
| ENSSSCG00000030343 | solute carrier family 38 member 9 [Source:HGNC Symbol;Acc:HGNC:26907] | SLC38A9 |
| ENSSSCG00000009169 | solute carrier family 39 member 8 [Source:HGNC Symbol;Acc:HGNC:20862] | SLC39A8 |
| ENSSSCG00000001825 | solute carrier family 41 member 3 [Source:HGNC Symbol;Acc:HGNC:31046] | SLC41A3 |
| ENSSSCG00000005425 | solute carrier family 44 member 1 [Source:HGNC Symbol;Acc:HGNC:18798] | SLC44A1 |
| ENSSSCG00000001419 | solute carrier family 44 member 4 [Source:HGNC Symbol;Acc:HGNC:13941] | SLC44A4 |
| ENSSSCG00000008943 | solute carrier family 4 member 4 [Source:HGNC Symbol;Acc:HGNC:11030] | SLC4A4 |
| ENSSSCG00000028060 | solute carrier family 4 member 8 [Source:HGNC Symbol;Acc:HGNC:11034] | SLC4A8 |
| ENSSSCG00000022649 | solute carrier family 7 member 11 [Source:HGNC Symbol;Acc:HGNC:11059] | SLC7A11 |
| ENSSSCG00000002041 | solute carrier family 7 member 7 [Source:HGNC Symbol;Acc:HGNC:11065] | SLC7A7 |
| ENSSSCG00000012266 | solute carrier family 9 member A7 [Source:HGNC Symbol;Acc:HGNC:17123] | SLC9A7 |
| ENSSSCG00000038844 | small integral membrane protein 5 [Source:HGNC Symbol;Acc:HGNC:40030] | SMIM5 |
| ENSSSCG00000002764 | sphingomyelin phosphodiesterase 3 [Source:HGNC Symbol;Acc:HGNC:14240] | SMPD3 |
| ENSSSCG00000037598 | sorting nexin 10 [Source:HGNC Symbol;Acc:HGNC:14974] | SNX10 |
| ENSSSCG00000036679 | sorbin and SH3 domain containing 2 [Source:HGNC Symbol;Acc:HGNC:24098] | SORBS2 |
| ENSSSCG00000008478 | SOS Ras/Rac guanine nucleotide exchange factor 1 [Source:HGNC Symbol;Acc:HGNC:11187] | SOS1 |
| ENSSSCG00000034191 | SRY-box 6 [Source:HGNC Symbol;Acc:HGNC:16421] | SOX6 |
| ENSSSCG00000015197 | sperm autoantigenic protein 17 [Source:HGNC Symbol;Acc:HGNC:11210] | SPA17 |
| ENSSSCG00000022250 | sperm acrosome associated 9 [Source:HGNC Symbol;Acc:HGNC:1367] | SPACA9 |
| ENSSSCG00000027587 | sperm associated antigen 1 [Source:HGNC Symbol;Acc:HGNC:11212] | SPAG1 |
| ENSSSCG00000006726 | sperm associated antigen 17 [Source:HGNC Symbol;Acc:HGNC:26620] | SPAG17 |
| ENSSSCG00000011080 | Sus scrofa sperm associated antigen 6 (SPAG6), mRNA. | SPAG6 |
| ENSSSCG00000033228 | sperm associated antigen 9 [Source:HGNC Symbol;Acc:HGNC:14524] | SPAG9 |
| ENSSSCG00000029778 | spermatogenesis associated 17 [Source:HGNC Symbol;Acc:HGNC:25184] | SPATA17 |
| ENSSSCG00000008834 | spermatogenesis associated 18 [Source:HGNC Symbol;Acc:HGNC:29579] | SPATA18 |
| ENSSSCG00000014348 | spermatogenesis associated 24 [Source:HGNC Symbol;Acc:HGNC:27322] | SPATA24 |
| ENSSSCG00000015767 | spermatogenesis associated 4 [Source:HGNC Symbol;Acc:HGNC:17333] | SPATA4 |
| ENSSSCG00000005221 | spermatogenesis associated 6 like [Source:HGNC Symbol;Acc:HGNC:25472] | SPATA6L |
| ENSSSCG00000039903 | spermatogenesis associated serine rich 1 [Source:HGNC Symbol;Acc:HGNC:22957] | SPATS1 |
| ENSSSCG00000007149 | Sus scrofa sperm flagellar 1 (SPEF1), mRNA. [Source:RefSeq mRNA;Acc:NM_001195369] | SPEF1 |
| ENSSSCG00000040786 | spectrin beta, non-erythrocytic 1 [Source:HGNC Symbol;Acc:HGNC:11275] | SPTBN1 |
| ENSSSCG00000007072 | serine palmitoyltransferase long chain base subunit 3 [Source:HGNC Symbol;Acc:HGNC:16253] | SPTLC3 |
| ENSSSCG00000033624 | SLIT-ROBO Rho GTPase activating protein 3 [Source:HGNC Symbol;Acc:HGNC:19744] | SRGAP3 |
| ENSSSCG00000038730 | sperm specific antigen 2 [Source:HGNC Symbol;Acc:HGNC:11319] | SSFA2 |
| ENSSSCG00000034493 | ST3 beta-galactoside alpha-2,3-sialyltransferase 6 [Source:HGNC Symbol;Acc:HGNC:18080] | ST3GAL6 |
| ENSSSCG00000013506 | signal transducing adaptor family member 2 [Source:HGNC Symbol;Acc:HGNC:30430] | STAP2 |
| ENSSSCG00000034942 | steroidogenic acute regulatory protein, mitochondrial precursor | STAR |
| ENSSSCG00000014818 | StAR related lipid transfer domain containing 10 [Source:HGNC Symbol;Acc:HGNC:10666] | STARD10 |
| ENSSSCG00000016214 | serine/threonine kinase 16 [Source:HGNC Symbol;Acc:HGNC:11394] | STK16 |
| ENSSSCG00000001913 | stomatin like 1 [Source:HGNC Symbol;Acc:HGNC:14560] | STOML1 |
| ENSSSCG00000009366 | stomatin like 3 [Source:HGNC Symbol;Acc:HGNC:19420] | STOML3 |
| ENSSSCG00000002412 | stonin 2 [Source:HGNC Symbol;Acc:HGNC:30652] | STON2 |
| ENSSSCG00000005582 | spermatid perinuclear RNA binding protein [Source:HGNC Symbol;Acc:HGNC:16462] | STRBP |
| ENSSSCG00000023280 | Sus scrofa sulfotransferase family 2B member 1 (SULT2B1), mRNA. | SULT2B1 |
| ENSSSCG00000005110 | spectrin repeat containing nuclear envelope protein 2 [Source:HGNC Symbol;Acc:HGNC:17084] | SYNE2 |

| | | |
|---|---|---|
| ENSSSCG00000003222 | synaptotagmin 3 [Source:HGNC Symbol;Acc:HGNC:11511] | SYT3 |
| ENSSSCG00000022224 | TATA-box binding protein associated factor 8 [Source:HGNC Symbol;Acc:HGNC:17300] | TAF8 |
| ENSSSCG00000008761 | TBC1 domain family member 19 [Source:HGNC Symbol;Acc:HGNC:25624] | TBC1D19 |
| ENSSSCG00000005376 | TBC1 domain family member 2 [Source:HGNC Symbol;Acc:HGNC:18026] | TBC1D2 |
| ENSSSCG00000023788 | TBC1 domain family member 30 [Source:HGNC Symbol;Acc:HGNC:29164] | TBC1D30 |
| ENSSSCG00000001705 | t-complex-associated-testis-expressed 1 [Source:HGNC Symbol;Acc:HGNC:11693] | TCTE1 |
| ENSSSCG00000031352 | Tctex1 domain containing 1 [Source:HGNC Symbol;Acc:HGNC:26882] | TCTEX1D1 |
| ENSSSCG00000008820 | tec protein tyrosine kinase [Source:HGNC Symbol;Acc:HGNC:11719] | TEC |
| ENSSSCG00000017884 | tektin 1 [Source:HGNC Symbol;Acc:HGNC:15534] | TEKT1 |
| ENSSSCG00000003633 | tektin 2 [Source:HGNC Symbol;Acc:HGNC:11725] | TEKT2 |
| ENSSSCG00000018028 | tektin 3 [Source:HGNC Symbol;Acc:HGNC:14293] | TEKT3 |
| ENSSSCG00000035104 | telomere repeat binding bouquet formation protein 1 [Source:HGNC Symbol;Acc:HGNC:26675] | TERB1 |
| ENSSSCG00000017118 | telomerase reverse transcriptase [Source:RefSeq peptide;Acc:NP_001231229] | TERT |
| ENSSSCG00000009332 | testis expressed 26 [Source:HGNC Symbol;Acc:HGNC:28622] | TEX26 |
| ENSSSCG00000004602 | testis expressed 9 [Source:HGNC Symbol;Acc:HGNC:29585] | TEX9 |
| ENSSSCG00000015732 | transcription factor CP2 like 1 [Source:HGNC Symbol;Acc:HGNC:17925] | TFCP2L1 |
| ENSSSCG00000036033 | thyroid hormone receptor beta [Source:HGNC Symbol;Acc:HGNC:11799] | THRB |
| ENSSSCG00000038186 | tigger transposable element derived 4 [Source:HGNC Symbol;Acc:HGNC:18335] | TIGD4 |
| ENSSSCG00000033000 | TIMP metallopeptidase inhibitor 4 [Source:HGNC Symbol;Acc:HGNC:11823] | TIMP4 |
| ENSSSCG00000024088 | talin 2 [Source:HGNC Symbol;Acc:HGNC:15447] | TLN2 |
| ENSSSCG00000031112 | transmembrane 4 L six family member 5 [Source:HGNC Symbol;Acc:HGNC:11857] | TM4SF5 |
| ENSSSCG00000005266 | transmembrane channel like 1 [Source:HGNC Symbol;Acc:HGNC:16513] | TMC1 |
| ENSSSCG00000027528 | transmembrane protein 107 [Source:HGNC Symbol;Acc:HGNC:28128] | TMEM107 |
| ENSSSCG00000031876 | transmembrane protein 159 [Source:HGNC Symbol;Acc:HGNC:30136] | TMEM159 |
| ENSSSCG00000016444 | transmembrane protein 176B [Source:HGNC Symbol;Acc:HGNC:29596] | TMEM176B |
| ENSSSCG00000011751 | transmembrane protein 212 [Source:HGNC Symbol;Acc:HGNC:34295] | TMEM212 |
| ENSSSCG00000014196 | transmembrane protein 232 [Source:HGNC Symbol;Acc:HGNC:37270] | TMEM232 |
| ENSSSCG00000039678 | transmembrane protein 269 [Source:HGNC Symbol;Acc:HGNC:52381] | TMEM269 |
| ENSSSCG00000006108 | transmembrane protein 67 [Source:HGNC Symbol;Acc:HGNC:28396] | TMEM67 |
| ENSSSCG00000013360 | transmembrane protein 86A [Source:HGNC Symbol;Acc:HGNC:26890] | TMEM86A |
| ENSSSCG00000034013 | transmembrane protein 88B [Source:HGNC Symbol;Acc:HGNC:37099] | TMEM88B |
| ENSSSCG00000038710 | transmembrane protease, serine 13 [Source:HGNC Symbol;Acc:HGNC:29808] | TMPRSS13 |
| ENSSSCG00000011849 | tyrosine kinase non receptor 2 [Source:HGNC Symbol;Acc:HGNC:19297] | TNK2 |
| ENSSSCG00000024505 | troponin I3, cardiac type [Source:HGNC Symbol;Acc:HGNC:11947] | TNNI3 |
| ENSSSCG00000027765 | TOG array regulator of axonemal microtubules 1 [Source:HGNC Symbol;Acc:HGNC:19959] | TOGARAM1 |
| ENSSSCG00000017601 | target of myb1 like 1 membrane trafficking protein [Source:HGNC Symbol;Acc:HGNC:11983] | TOM1L1 |
| ENSSSCG00000021966 | DNA topoisomerase II binding protein 1 [Source:HGNC Symbol;Acc:HGNC:17008] | TOPBP1 |
| ENSSSCG00000025685 | tumor protein p73 [Source:HGNC Symbol;Acc:HGNC:12003] | TP73 |
| ENSSSCG00000013849 | tropomyosin 4 [Source:HGNC Symbol;Acc:HGNC:12013] | TPM4 |
| ENSSSCG00000032698 | tubulin polymerization promoting protein [Source:HGNC Symbol;Acc:HGNC:24164] | TPPP |
| ENSSSCG00000035983 | tubulin polymerization promoting protein family member 3 | TPPP3 |
| ENSSSCG00000032721 | TRAF3 interacting protein 3 [Source:HGNC Symbol;Acc:HGNC:30766] | TRAF3IP3 |
| ENSSSCG00000022512 | T-cell receptor delta constant [Source:HGNC Symbol;Acc:HGNC:12253] | TRDC |
| ENSSSCG00000035153 | tripartite motif containing 38 [Source:HGNC Symbol;Acc:HGNC:10059] | TRIM38 |
| ENSSSCG00000027684 | E3 ubiquitin-protein ligase TRIM63 [Source:RefSeq peptide;Acc:NP_001171685] | TRIM63 |
| ENSSSCG00000037835 | tripartite motif containing 7 [Source:HGNC Symbol;Acc:HGNC:16278] | TRIM7 |
| ENSSSCG00000030241 | TSC22 domain family member 3 [Source:HGNC Symbol;Acc:HGNC:3051] | TSC22D3 |
| ENSSSCG00000008186 | testis specific 10 [Source:HGNC Symbol;Acc:HGNC:14927] | TSGA10 |
| ENSSSCG00000002767 | translin associated factor X interacting protein 1 [Source:HGNC Symbol;Acc:HGNC:18586] | TSNAXIP1 |
| ENSSSCG00000003908 | tetraspanin 1 [Source:HGNC Symbol;Acc:HGNC:20657] | TSPAN1 |
| ENSSSCG00000015047 | tetratricopeptide repeat domain 12 [Source:HGNC Symbol;Acc:HGNC:23700] | TTC12 |
| ENSSSCG00000011263 | tetratricopeptide repeat domain 21A [Source:HGNC Symbol;Acc:HGNC:30761] | TTC21A |
| ENSSSCG00000016825 | tetratricopeptide repeat domain 23 like [Source:HGNC Symbol;Acc:HGNC:26355] | TTC23L |
| ENSSSCG00000016511 | tetratricopeptide repeat domain 26 [Source:HGNC Symbol;Acc:HGNC:21882] | TTC26 |
| ENSSSCG00000002428 | tetratricopeptide repeat domain 8 [Source:HGNC Symbol;Acc:HGNC:20087] | TTC8 |
| ENSSSCG00000006733 | transcription termination factor 2 [Source:HGNC Symbol;Acc:HGNC:12398] | TTF2 |
| ENSSSCG00000003336 | tubulin tyrosine ligase like 10 [Source:HGNC Symbol;Acc:HGNC:26693] | TTLL10 |
| ENSSSCG00000017542 | tubulin tyrosine ligase like 6 [Source:HGNC Symbol;Acc:HGNC:26664] | TTLL6 |
| ENSSSCG00000000982 | tubulin tyrosine ligase like 8 [Source:HGNC Symbol;Acc:HGNC:34000] | TTLL8 |
| ENSSSCG00000007236 | tubulin tyrosine ligase like 9 [Source:HGNC Symbol;Acc:HGNC:16118] | TTLL9 |
| ENSSSCG00000006620 | tuftelin 1 [Source:HGNC Symbol;Acc:HGNC:12422] | TUFT1 |
| ENSSSCG00000000737 | tubby like protein 3 [Source:HGNC Symbol;Acc:HGNC:12425] | TULP3 |
| ENSSSCG00000011393 | ubiquitin like modifier activating enzyme 7 [Source:HGNC Symbol;Acc:HGNC:12471] | UBA7 |

| | | |
|---|---|---|
| ENSSSCG00000023423 | UBA like domain containing 2 [Source:HGNC Symbol;Acc:HGNC:28438] | UBALD2 |
| ENSSSCG00000003810 | ubiquitin conjugating enzyme E2 U (putative) [Source:HGNC Symbol;Acc:HGNC:28559] | UBE2U |
| ENSSSCG00000021005 | ubiquilin 1 [Source:HGNC Symbol;Acc:HGNC:12508] | UBQLN1 |
| ENSSSCG00000003498 | UBX domain protein 10 [Source:HGNC Symbol;Acc:HGNC:26354] | UBXN10 |
| ENSSSCG00000033546 | UBX domain protein 11 [Source:HGNC Symbol;Acc:HGNC:30600] | UBXN11 |
| ENSSSCG00000008368 | UDP-glucose pyrophosphorylase 2 [Source:HGNC Symbol;Acc:HGNC:12527] | UGP2 |
| ENSSSCG00000025408 | unc-51 like kinase 4 [Source:HGNC Symbol;Acc:HGNC:15784] | ULK4 |
| ENSSSCG00000015100 | uroplakin 2 [Source:HGNC Symbol;Acc:HGNC:12579] | UPK2 |
| ENSSSCG00000001823 | urocanate hydratase 1 [Source:HGNC Symbol;Acc:HGNC:26444] | UROC1 |
| ENSSSCG00000015120 | ubiquitin specific peptidase 2 [Source:HGNC Symbol;Acc:HGNC:12618] | USP2 |
| ENSSSCG00000017995 | ubiquitin specific peptidase 43 [Source:HGNC Symbol;Acc:HGNC:20072] | USP43 |
| ENSSSCG00000010302 | ubiquitin specific peptidase 54 [Source:HGNC Symbol;Acc:HGNC:23513] | USP54 |
| ENSSSCG00000000703 | vesicle associated membrane protein 1 [Source:HGNC Symbol;Acc:HGNC:12642] | VAMP1 |
| ENSSSCG00000014136 | versican [Source:HGNC Symbol;Acc:HGNC:2464] | VCAN |
| ENSSSCG00000017915 | vitelline membrane outer layer protein 1 homolog precursor | VMO1 |
| ENSSSCG00000025266 | von Willebrand factor A domain containing 3A [Source:HGNC Symbol;Acc:HGNC:27088] | VWA3A |
| ENSSSCG00000033394 | von Willebrand factor A domain containing 3B [Source:HGNC Symbol;Acc:HGNC:28385] | VWA3B |
| ENSSSCG00000004008 | WD repeat domain 27 [Source:HGNC Symbol;Acc:HGNC:21248] | WDR27 |
| ENSSSCG00000020987 | WD repeat domain 31 [Source:HGNC Symbol;Acc:HGNC:21421] | WDR31 |
| ENSSSCG00000005655 | WD repeat domain 34 [Source:HGNC Symbol;Acc:HGNC:28296] | WDR34 |
| ENSSSCG00000011738 | WD repeat domain 49 [Source:HGNC Symbol;Acc:HGNC:26587] | WDR49 |
| ENSSSCG00000038486 | WD repeat domain 60 [Source:HGNC Symbol;Acc:HGNC:21862] | WDR60 |
| ENSSSCG00000006950 | WD repeat domain 63 [Source:HGNC Symbol;Acc:HGNC:30711] | WDR63 |
| ENSSSCG00000009802 | WD repeat domain 66 [Source:HGNC Symbol;Acc:HGNC:28506] | WDR66 |
| ENSSSCG00000022305 | WD repeat domain 78 [Source:HGNC Symbol;Acc:HGNC:26252] | WDR78 |
| ENSSSCG00000007991 | WD repeat domain 90 [Source:HGNC Symbol;Acc:HGNC:26960] | WDR90 |
| ENSSSCG00000005490 | whirlin [Source:HGNC Symbol;Acc:HGNC:16361] | WHRN |
| ENSSSCG00000006298 | lymphotactin precursor [Source:RefSeq peptide;Acc:NP_001127817] | XCL1 |
| ENSSSCG00000006191 | XK related 9 [Source:HGNC Symbol;Acc:HGNC:20937] | XKR9 |
| ENSSSCG00000012487 | XK related, X-linked [Source:HGNC Symbol;Acc:HGNC:29845] | XKRX |
| ENSSSCG00000015537 | xenotropic and polytropic retrovirus receptor 1 [Source:HGNC Symbol;Acc:HGNC:12827] | XPR1 |
| ENSSSCG00000021702 | X-ray radiation resistance associated 1 [Source:HGNC Symbol;Acc:HGNC:18868] | XRRA1 |
| ENSSSCG00000023393 | zinc finger B-box domain containing [Source:HGNC Symbol;Acc:HGNC:26245] | ZBBX |
| ENSSSCG00000002380 | zinc finger C2HC-type containing 1C [Source:HGNC Symbol;Acc:HGNC:20354] | ZC2HC1C |
| ENSSSCG00000002778 | zinc finger DHHC-type containing 1 [Source:HGNC Symbol;Acc:HGNC:17916] | ZDHHC1 |
| ENSSSCG00000010402 | zinc finger AN1-type containing 4 [Source:HGNC Symbol;Acc:HGNC:23504] | ZFAND4 |
| ENSSSCG00000033860 | zinc finger matrin-type 3 [Source:HGNC Symbol;Acc:HGNC:29983] | ZMAT3 |
| ENSSSCG00000011408 | zinc finger MYND-type containing 10 [Source:HGNC Symbol;Acc:HGNC:19412] | ZMYND10 |
| ENSSSCG00000003967 | zinc finger MYND-type containing 12 [Source:HGNC Symbol;Acc:HGNC:21192] | ZMYND12 |
| ENSSSCG00000007958 | zinc finger protein 174 [Source:HGNC Symbol;Acc:HGNC:12963] | ZNF174 |
| ENSSSCG00000023276 | zinc finger protein 24 [Source:HGNC Symbol;Acc:HGNC:13032] | ZNF24 |
| ENSSSCG00000030018 | zinc finger protein 396 [Source:HGNC Symbol;Acc:HGNC:18824] | ZNF396 |
| ENSSSCG00000040579 | zinc finger protein 683 [Source:HGNC Symbol;Acc:HGNC:28495] | ZNF683 |
| ENSSSCG00000001203 | zinc finger and SCAN domain containing 9 [Source:HGNC Symbol;Acc:HGNC:12984] | ZSCAN9 |
| ENSSSCG00000016030 | zinc finger SWIM-type containing 2 [Source:HGNC Symbol;Acc:HGNC:30990] | ZSWIM2 |
| ENSSSCG00000004263 | zinc finger with UFM1 specific peptidase domain [Source:HGNC Symbol;Acc:HGNC:21224] | ZUFSP |
| ENSSSCG00000011147 | aldo-keto reductase family 1, member C-like 1 [Source:RefSeq peptide;Acc:NP_001033715] |  |
| ENSSSCG00000038825 | cationic amino acid transporter 3-like [Source:RefSeq peptide;Acc:NP_001231094] |  |
| ENSSSCG00000015664 | complement decay-accelerating factor precursor [Source:RefSeq peptide;Acc:NP_998980] |  |
| ENSSSCG00000009578 | cyclin-dependent kinase 20 [Source:RefSeq peptide;Acc:NP_001182258] |  |
| ENSSSCG00000017754 | galectin-9 isoform 1 [Source:RefSeq peptide;Acc:NP_999097] |  |
| ENSSSCG00000039731 | glutaredoxin-1 [Source:RefSeq peptide;Acc:NP_999398] |  |
| ENSSSCG00000035293 | Insulin-like growth factor II Insulin-like growth factor II Preptin | |
| ENSSSCG00000031086 | Lutropin subunit beta [Source:UniProtKB/Swiss-Prot;Acc:P01232] |  |
| ENSSSCG00000035596 | mutS protein homolog 5 [Source:RefSeq peptide;Acc:NP_001182287] |  |
| ENSSSCG00000001229 | patr class I histocompatibility antigen, A-126 alpha chain-like precursor | |
| ENSSSCG00000010006 | SEC14-like protein 2 [Source:RefSeq peptide;Acc:NP_001185847] |  |
| ENSSSCG00000008421 | Sus scrofa luteinizing hormone/choriogonadotropin receptor (LHCGR), mRNA. | |
| ENSSSCG00000014004 | Sus scrofa phospholipid phosphatase 2 (PLPP2), mRNA. | |
| ENSSSCG00000027792 | uncharacterized protein LOC100623156 [Source:RefSeq peptide;Acc:NP_001231588] |  |

### Table S7. Gene Ontology on the DEGs and DSGs

#### (1) GO analysis on DEGs

| | |
|---|---|
| Analysis Type: | PANTHER Overrepresentation Test (Released 20171205) |
| Annotation Version and Release Date: | GO Ontology database Released 2018-02-02 |
| Analyzed List: | upload_1 (Sus scrofa) |
| Reference List: | Sus scrofa (all genes in database) |
| Test Type: | FISHER |

| **GO biological process complete** | REFLIST (22191) | upload_1 (599) | upload_1 (expected) | upload_1 (over/under) | upload_1 (fold Enrichment) | upload_1 (raw P-value) | upload_1 (FDR) |
|---|---|---|---|---|---|---|---|
| inner dynein arm assembly (GO:0036159) | 10 | 8 | 0.27 | + | 29.64 | 7.54E-09 | 3.56E-06 |
| regulation of cilium movement (GO:0003352) | 8 | 6 | 0.22 | + | 27.79 | 8.07E-07 | 2.42E-04 |
| cilium or flagellum-dependent cell motility (GO:0001539) | 12 | 9 | 0.32 | + | 27.79 | 1.25E-09 | 8.54E-07 |
| cilium-dependent cell motility (GO:0060285) | 11 | 8 | 0.3 | + | 26.94 | 1.27E-08 | 5.59E-06 |
| cilium movement involved in cell motility (GO:0060294) | 7 | 5 | 0.19 | + | 26.46 | 8.38E-06 | 2.19E-03 |
| regulation of cilium beat frequency (GO:0003356) | 6 | 4 | 0.16 | + | 24.7 | 8.75E-05 | 1.85E-02 |
| axonemal dynein complex assembly (GO:0070286) | 20 | 13 | 0.54 | + | 24.08 | 8.89E-13 | 7.80E-10 |
| cilium movement (GO:0003341) | 34 | 22 | 0.92 | + | 23.97 | 1.09E-20 | 1.67E-17 |
| epithelial cilium movement (GO:0003351) | 10 | 6 | 0.27 | + | 22.23 | 2.06E-06 | 6.02E-04 |
| regulation of microtubule-based movement (GO:0060632) | 10 | 6 | 0.27 | + | 22.23 | 2.06E-06 | 5.88E-04 |
| axoneme assembly (GO:0035082) | 34 | 20 | 0.92 | + | 21.79 | 2.53E-18 | 2.83E-15 |
| outer dynein arm assembly (GO:0036158) | 12 | 7 | 0.32 | + | 21.61 | 3.20E-07 | 1.12E-04 |
| sperm axoneme assembly (GO:0007288) | 8 | 4 | 0.22 | + | 18.52 | 1.98E-04 | 3.92E-02 |
| motile cilium assembly (GO:0044458) | 14 | 6 | 0.38 | + | 15.88 | 9.10E-06 | 2.33E-03 |
| microtubule bundle formation (GO:0001578) | 50 | 21 | 1.35 | + | 15.56 | 7.40E-17 | 7.57E-14 |
| intraciliary transport (GO:0042073) | 20 | 8 | 0.54 | + | 14.82 | 4.25E-07 | 1.41E-04 |
| protein transport along microtubule (GO:0098840) | 21 | 8 | 0.57 | + | 14.11 | 5.73E-07 | 1.85E-04 |
| microtubule-based protein transport (GO:0099118) | 21 | 8 | 0.57 | + | 14.11 | 5.73E-07 | 1.81E-04 |
| cilium organization (GO:0044782) | 159 | 55 | 4.29 | + | 12.81 | 1.30E-38 | 1.60E-34 |
| cilium assembly (GO:0060271) | 148 | 51 | 3.99 | + | 12.77 | 8.57E-36 | 5.26E-32 |
| flagellated sperm motility (GO:0030317) | 38 | 12 | 1.03 | + | 11.7 | 4.75E-09 | 2.54E-06 |
| sperm motility (GO:0097722) | 39 | 12 | 1.05 | + | 11.4 | 6.07E-09 | 3.11E-06 |
| microtubule-based movement (GO:0007018) | 132 | 38 | 3.56 | + | 10.66 | 1.37E-24 | 2.81E-21 |
| plasma membrane bounded cell projection assembly (GO:0120031) | 184 | 51 | 4.97 | + | 10.27 | 5.59E-32 | 2.29E-28 |
| cell projection assembly (GO:0030031) | 185 | 51 | 4.99 | + | 10.21 | 6.96E-32 | 2.14E-28 |
| determination of left/right symmetry (GO:0007368) | 54 | 14 | 1.46 | + | 9.6 | 2.16E-09 | 1.39E-06 |
| determination of bilateral symmetry (GO:0009855) | 57 | 14 | 1.54 | + | 9.1 | 3.92E-09 | 2.29E-06 |
| specification of symmetry (GO:0009799) | 57 | 14 | 1.54 | + | 9.1 | 3.92E-09 | 2.19E-06 |
| protein complex localization (GO:0031503) | 39 | 9 | 1.05 | + | 8.55 | 3.79E-06 | 1.06E-03 |
| microtubule-based transport (GO:0099111) | 50 | 9 | 1.35 | + | 6.67 | 2.19E-05 | 5.50E-03 |
| transport along microtubule (GO:0010970) | 50 | 9 | 1.35 | + | 6.67 | 2.19E-05 | 5.39E-03 |
| microtubule-based process (GO:0007017) | 339 | 56 | 9.15 | + | 6.12 | 4.37E-25 | 1.07E-21 |
| organelle assembly (GO:0070925) | 352 | 56 | 9.5 | + | 5.89 | 2.25E-24 | 3.95E-21 |
| cytoskeleton-dependent intracellular transport (GO:0030705) | 66 | 10 | 1.78 | + | 5.61 | 2.93E-05 | 7.06E-03 |
| microtubule cytoskeleton organization (GO:0000226) | 228 | 29 | 6.15 | + | 4.71 | 4.77E-11 | 3.45E-08 |
| cell projection organization (GO:0030030) | 450 | 57 | 12.15 | + | 4.69 | 1.75E-20 | 2.39E-17 |
| plasma membrane bounded cell projection organization (GO:0120036 | 445 | 56 | 12.01 | + | 4.66 | 5.08E-20 | 6.24E-17 |
| pattern specification process (GO:0007389) | 199 | 19 | 5.37 | + | 3.54 | 5.45E-06 | 1.49E-03 |

| | | | | | | | |
|---|---|---|---|---|---|---|---|
| movement of cell or subcellular component (GO:0006928) | 583 | 54 | 15.74 | + | 3.43 | 3.86E-14 | 3.64E-11 |
| cellular protein complex assembly (GO:0043623) | 194 | 16 | 5.24 | + | 3.06 | 1.47E-04 | 3.06E-02 |
| cytoskeleton organization (GO:0007010) | 548 | 41 | 14.79 | + | 2.77 | 1.73E-08 | 7.32E-06 |
| cellular component assembly (GO:0022607) | 1036 | 69 | 27.96 | + | 2.47 | 2.63E-11 | 2.16E-08 |
| cellular component biogenesis (GO:0044085) | 1185 | 70 | 31.99 | + | 2.19 | 2.91E-09 | 1.79E-06 |
| organelle organization (GO:0006996) | 1639 | 84 | 44.24 | + | 1.9 | 4.29E-08 | 1.65E-05 |
| cellular component organization (GO:0016043) | 2537 | 103 | 68.48 | + | 1.5 | 4.18E-05 | 9.34E-03 |
| regulation of cellular biosynthetic process (GO:0031326) | 2132 | 32 | 57.55 | - | 0.56 | 2.25E-04 | 4.31E-02 |
| regulation of macromolecule biosynthetic process (GO:0010556) | 2039 | 30 | 55.04 | - | 0.55 | 2.21E-04 | 4.31E-02 |
| regulation of cellular macromolecule biosynthetic process (GO:20001 | 1978 | 29 | 53.39 | - | 0.54 | 2.45E-04 | 4.55E-02 |
| macromolecule biosynthetic process (GO:0009059) | 1283 | 15 | 34.63 | - | 0.43 | 2.28E-04 | 4.31E-02 |
| nucleic acid metabolic process (GO:0090304) | 1511 | 17 | 40.79 | - | 0.42 | 3.04E-05 | 7.05E-03 |
| gene expression (GO:0010467) | 1448 | 16 | 39.09 | - | 0.41 | 4.30E-05 | 9.44E-03 |
| response to organic substance (GO:0010033) | 1076 | 11 | 29.04 | - | 0.38 | 1.98E-04 | 3.98E-02 |
| cellular response to chemical stimulus (GO:0070887) | 1087 | 11 | 29.34 | - | 0.37 | 1.48E-04 | 3.03E-02 |
| RNA metabolic process (GO:0016070) | 1198 | 12 | 32.34 | - | 0.37 | 6.16E-05 | 1.33E-02 |
| system process (GO:0003008) | 1252 | 11 | 33.8 | - | 0.33 | 7.43E-06 | 1.99E-03 |
| nervous system process (GO:0050877) | 1001 | 8 | 27.02 | - | 0.3 | 3.00E-05 | 7.09E-03 |
| response to chemical (GO:0042221) | 2204 | 17 | 59.49 | - | 0.29 | 4.56E-11 | 3.50E-08 |
| sensory perception (GO:0007600) | 876 | 6 | 23.65 | - | 0.25 | 3.46E-05 | 7.89E-03 |
| G-protein coupled receptor signaling pathway (GO:0007186) | 1196 | 8 | 32.28 | - | 0.25 | 6.26E-07 | 1.92E-04 |
| sensory perception of smell (GO:0007608) | 692 | 1 | 18.68 | - | 0.05 | 3.47E-07 | 1.18E-04 |
| detection of stimulus involved in sensory perception (GO:0050906) | 722 | 1 | 19.49 | - | 0.05 | 1.61E-07 | 6.01E-05 |
| sensory perception of chemical stimulus (GO:0007606) | 722 | 1 | 19.49 | - | 0.05 | 1.61E-07 | 5.83E-05 |
| detection of stimulus (GO:0051606) | 769 | 1 | 20.76 | - | 0.05 | 3.19E-08 | 1.26E-05 |

| **GO molecular function complete** | REFLIST (22191) | upload_1 (599) | upload_1 (expected) | upload_1 (over/under) | upload_1 (fold Enrichment) | upload_1 (raw P-value) | upload_1 (FDR) |
|---|---|---|---|---|---|---|---|
| dynein light chain binding (GO:0045503) | 12 | 10 | 0.32 | + | 30.87 | 7.11E-11 | 1.27E-07 |
| ATP-dependent microtubule motor activity, minus-end-directed (GO:0 | 8 | 6 | 0.22 | + | 27.79 | 8.07E-07 | 3.20E-04 |
| dynein heavy chain binding (GO:0045504) | 6 | 4 | 0.16 | + | 24.7 | 8.75E-05 | 1.83E-02 |
| dynein intermediate chain binding (GO:0045505) | 14 | 7 | 0.38 | + | 18.52 | 7.06E-07 | 3.15E-04 |
| ATP-dependent microtubule motor activity (GO:1990939) | 28 | 13 | 0.76 | + | 17.2 | 2.25E-11 | 8.03E-08 |
| dynein light intermediate chain binding (GO:0051959) | 18 | 8 | 0.49 | + | 16.47 | 2.24E-07 | 1.33E-04 |
| ATP-dependent microtubule motor activity, plus-end-directed (GO:000 | 20 | 7 | 0.54 | + | 12.97 | 4.70E-06 | 1.12E-03 |
| microtubule motor activity (GO:0003777) | 61 | 16 | 1.65 | + | 9.72 | 1.32E-10 | 1.57E-07 |
| motor activity (GO:0003774) | 99 | 19 | 2.67 | + | 7.11 | 2.60E-10 | 2.32E-07 |
| microtubule binding (GO:0008017) | 144 | 13 | 3.89 | + | 3.34 | 2.69E-04 | 4.80E-02 |
| tubulin binding (GO:0015631) | 181 | 15 | 4.89 | + | 3.07 | 2.24E-04 | 4.20E-02 |
| nucleic acid binding (GO:0003676) | 2480 | 33 | 66.94 | - | 0.49 | 3.18E-06 | 8.71E-04 |
| signal transducer activity (GO:0004871) | 1510 | 18 | 40.76 | - | 0.44 | 8.77E-05 | 1.74E-02 |
| molecular transducer activity (GO:0060089) | 1460 | 14 | 39.41 | - | 0.36 | 4.34E-06 | 1.11E-03 |
| receptor activity (GO:0004872) | 1427 | 12 | 38.52 | - | 0.31 | 8.92E-07 | 3.18E-04 |
| transmembrane signaling receptor activity (GO:0004888) | 1238 | 10 | 33.42 | - | 0.3 | 2.75E-06 | 8.92E-04 |
| transmembrane receptor activity (GO:0099600) | 1238 | 10 | 33.42 | - | 0.3 | 2.75E-06 | 8.18E-04 |
| signaling receptor activity (GO:0038023) | 1331 | 10 | 35.93 | - | 0.28 | 5.76E-07 | 2.94E-04 |
| G-protein coupled receptor activity (GO:0004930) | 1019 | 7 | 27.51 | - | 0.25 | 5.38E-06 | 1.20E-03 |

| GO cellular component complete | REFLIST (22191) | upload_1 (599) | upload_1 (expected) | upload_1 (over/under) | upload_1 (fold Enrichment) | upload_1 (raw P-value) | upload_1 (FDR) |
|---|---|---|---|---|---|---|---|
| axoneme part (GO:0044447) | 8 | 7 | 0.22 | + | 32.42 | 4.48E-08 | 3.33E-06 |
| axonemal dynein complex (GO:0005858) | 7 | 6 | 0.19 | + | 31.75 | 4.72E-07 | 2.94E-05 |
| dynein complex (GO:0030286) | 23 | 13 | 0.62 | + | 20.94 | 3.33E-12 | 3.71E-10 |
| intraciliary transport particle A (GO:0030991) | 8 | 4 | 0.22 | + | 18.52 | 1.98E-04 | 9.62E-03 |
| ciliary plasm (GO:0097014) | 47 | 23 | 1.27 | + | 18.13 | 1.68E-19 | 3.73E-17 |
| axoneme (GO:0005930) | 46 | 22 | 1.24 | + | 17.72 | 1.52E-18 | 2.95E-16 |
| intraciliary transport particle (GO:0030990) | 24 | 9 | 0.65 | + | 13.89 | 1.24E-07 | 8.03E-06 |
| plasma membrane bounded cell projection cytoplasm (GO:0032838) | 65 | 23 | 1.75 | + | 13.11 | 5.23E-17 | 6.79E-15 |
| intraciliary transport particle B (GO:0030992) | 15 | 5 | 0.4 | + | 12.35 | 1.38E-04 | 6.93E-03 |
| photoreceptor connecting cilium (GO:0032391) | 18 | 6 | 0.49 | + | 12.35 | 2.89E-05 | 1.61E-03 |
| ciliary transition zone (GO:0035869) | 32 | 10 | 0.86 | + | 11.58 | 1.01E-07 | 7.13E-06 |
| motile cilium (GO:0031514) | 73 | 22 | 1.97 | + | 11.16 | 4.01E-15 | 4.81E-13 |
| ciliary basal body (GO:0036064) | 54 | 15 | 1.46 | + | 10.29 | 2.54E-10 | 2.47E-08 |
| sperm flagellum (GO:0036126) | 47 | 13 | 1.27 | + | 10.25 | 4.18E-09 | 3.43E-07 |
| 9+2 motile cilium (GO:0097729) | 47 | 13 | 1.27 | + | 10.25 | 4.18E-09 | 3.26E-07 |
| cilium (GO:0005929) | 263 | 62 | 7.1 | + | 8.73 | 2.96E-35 | 4.61E-32 |
| ciliary part (GO:0044441) | 196 | 44 | 5.29 | + | 8.32 | 1.58E-24 | 1.23E-21 |
| microtubule associated complex (GO:0005875) | 88 | 17 | 2.38 | + | 7.16 | 2.09E-09 | 1.92E-07 |
| photoreceptor cell cilium (GO:0097733) | 47 | 8 | 1.27 | + | 6.31 | 8.91E-05 | 4.79E-03 |
| 9+0 non-motile cilium (GO:0097731) | 47 | 8 | 1.27 | + | 6.31 | 8.91E-05 | 4.63E-03 |
| sperm part (GO:0097223) | 90 | 15 | 2.43 | + | 6.17 | 1.04E-07 | 7.06E-06 |
| cytoplasmic region (GO:0099568) | 156 | 24 | 4.21 | + | 5.7 | 7.06E-11 | 7.34E-09 |
| non-motile cilium (GO:0097730) | 63 | 8 | 1.7 | + | 4.7 | 5.39E-04 | 2.27E-02 |
| cell projection part (GO:0044463) | 404 | 50 | 10.91 | + | 4.58 | 1.04E-17 | 1.80E-15 |
| plasma membrane bounded cell projection part (GO:0120038) | 404 | 50 | 10.91 | + | 4.58 | 1.04E-17 | 1.62E-15 |
| plasma membrane bounded cell projection (GO:0120025) | 652 | 71 | 17.6 | + | 4.03 | 7.00E-22 | 3.63E-19 |
| cell projection (GO:0042995) | 662 | 71 | 17.87 | + | 3.97 | 1.54E-21 | 5.99E-19 |
| microtubule cytoskeleton (GO:0015630) | 525 | 56 | 14.17 | + | 3.95 | 4.65E-17 | 6.58E-15 |
| microtubule organizing center (GO:0005815) | 313 | 31 | 8.45 | + | 3.67 | 2.85E-09 | 2.47E-07 |
| cytoskeletal part (GO:0044430) | 750 | 74 | 20.24 | + | 3.66 | 1.70E-20 | 5.30E-18 |
| microtubule (GO:0005874) | 135 | 12 | 3.64 | + | 3.29 | 5.21E-04 | 2.26E-02 |
| cytoskeleton (GO:0005856) | 955 | 83 | 25.78 | + | 3.22 | 9.82E-20 | 2.55E-17 |
| polymeric cytoskeletal fiber (GO:0099513) | 264 | 18 | 7.13 | + | 2.53 | 5.18E-04 | 2.31E-02 |
| intracellular non-membrane-bounded organelle (GO:0043232) | 2036 | 93 | 54.96 | + | 1.69 | 1.06E-06 | 6.36E-05 |
| non-membrane-bounded organelle (GO:0043228) | 2036 | 93 | 54.96 | + | 1.69 | 1.06E-06 | 6.12E-05 |
| intracellular membrane-bounded organelle (GO:0043231) | 5322 | 109 | 143.66 | - | 0.76 | 9.31E-04 | 3.82E-02 |
| intracellular ribonucleoprotein complex (GO:0030529) | 575 | 3 | 15.52 | - | 0.19 | 2.96E-04 | 1.40E-02 |
| ribonucleoprotein complex (GO:1990904) | 578 | 3 | 15.6 | - | 0.19 | 3.02E-04 | 1.38E-02 |

#### (2) GO analysis on DSGs

| | |
|---|---|
| Analysis Type: | PANTHER Overrepresentation Test (Released 20171205) |
| Annotation Version and Release Date: | GO Ontology database Released 2018-02-02 |
| Analyzed List: | upload_1 (Sus scrofa) |
| Reference List: | Sus scrofa (all genes in database) |
| Test Type: | FISHER |

| GO biological process complete | REFLIST (22191) | upload_1 (648) | upload_1 (expected) | upload_1 (over/under) | upload_1 (fold Enrichment) | upload_1 (raw P-value) | upload_1 (FDR) |
|---|---|---|---|---|---|---|---|
| cilium movement involved in cell motility (GO:0060294) | 7 | 5 | 0.2 | + | 24.46 | 1.22E-05 | 4.27E-03 |
| inner dynein arm assembly (GO:0036159) | 10 | 6 | 0.29 | + | 20.55 | 3.20E-06 | 1.23E-03 |
| cilium or flagellum-dependent cell motility (GO:0001539) | 12 | 7 | 0.35 | + | 19.98 | 5.36E-07 | 2.35E-04 |
| cilium-dependent cell motility (GO:0060285) | 11 | 6 | 0.32 | + | 18.68 | 4.83E-06 | 1.80E-03 |
| axonemal dynein complex assembly (GO:0070286) | 20 | 10 | 0.58 | + | 17.12 | 5.68E-09 | 4.10E-06 |
| cilium movement (GO:0003341) | 34 | 17 | 0.99 | + | 17.12 | 2.45E-14 | 3.76E-11 |
| axoneme assembly (GO:0035082) | 34 | 16 | 0.99 | + | 16.12 | 2.95E-13 | 4.03E-10 |
| outer dynein arm assembly (GO:0036158) | 12 | 5 | 0.35 | + | 14.27 | 8.44E-05 | 2.47E-02 |
| microtubule bundle formation (GO:0001578) | 50 | 17 | 1.46 | + | 11.64 | 3.66E-12 | 4.09E-09 |
| intraciliary transport (GO:0042073) | 20 | 6 | 0.58 | + | 10.27 | 7.23E-05 | 2.22E-02 |
| protein transport along microtubule (GO:0098840) | 21 | 6 | 0.61 | + | 9.78 | 9.07E-05 | 2.59E-02 |
| microtubule-based protein transport (GO:0099118) | 21 | 6 | 0.61 | + | 9.78 | 9.07E-05 | 2.53E-02 |
| cilium assembly (GO:0060271) | 148 | 40 | 4.32 | + | 9.26 | 1.03E-23 | 6.30E-20 |
| cilium organization (GO:0044782) | 159 | 42 | 4.64 | + | 9.05 | 1.62E-24 | 2.00E-20 |
| flagellated sperm motility (GO:0030317) | 38 | 10 | 1.11 | + | 9.01 | 7.80E-07 | 3.30E-04 |
| sperm motility (GO:0097722) | 39 | 10 | 1.14 | + | 8.78 | 9.55E-07 | 3.91E-04 |
| microtubule-based movement (GO:0007018) | 132 | 32 | 3.85 | + | 8.3 | 4.89E-18 | 1.00E-14 |
| plasma membrane bounded cell projection assembly (GO:0120031) | 184 | 40 | 5.37 | + | 7.44 | 9.41E-21 | 3.86E-17 |
| cell projection assembly (GO:0030031) | 185 | 40 | 5.4 | + | 7.4 | 1.12E-20 | 3.43E-17 |
| determination of left/right symmetry (GO:0007368) | 54 | 10 | 1.58 | + | 6.34 | 1.20E-05 | 4.34E-03 |
| determination of bilateral symmetry (GO:0009855) | 57 | 10 | 1.66 | + | 6.01 | 1.82E-05 | 6.22E-03 |
| specification of symmetry (GO:0009799) | 57 | 10 | 1.66 | + | 6.01 | 1.82E-05 | 6.05E-03 |
| microtubule-based process (GO:0007017) | 339 | 49 | 9.9 | + | 4.95 | 1.73E-18 | 4.24E-15 |
| organelle assembly (GO:0070925) | 352 | 46 | 10.28 | + | 4.48 | 6.27E-16 | 1.10E-12 |
| microtubule cytoskeleton organization (GO:0000226) | 228 | 25 | 6.66 | + | 3.75 | 7.02E-08 | 3.92E-05 |
| cell projection organization (GO:0030030) | 450 | 46 | 13.14 | + | 3.5 | 2.03E-12 | 2.49E-09 |
| plasma membrane bounded cell projection organization (GO:0120036 | 445 | 45 | 12.99 | + | 3.46 | 4.97E-12 | 5.09E-09 |
| movement of cell or subcellular component (GO:0006928) | 583 | 50 | 17.02 | + | 2.94 | 7.78E-11 | 7.36E-08 |
| cytoskeleton organization (GO:0007010) | 548 | 40 | 16 | + | 2.5 | 3.98E-07 | 1.81E-04 |
| cellular component assembly (GO:0022607) | 1036 | 54 | 30.25 | + | 1.78 | 7.36E-05 | 2.21E-02 |
| organelle organization (GO:0006996) | 1639 | 78 | 47.86 | + | 1.63 | 3.94E-05 | 1.24E-02 |
| cellular macromolecule biosynthetic process (GO:0034645) | 1263 | 16 | 36.88 | - | 0.43 | 1.72E-04 | 4.58E-02 |
| macromolecule biosynthetic process (GO:0009059) | 1283 | 16 | 37.46 | - | 0.43 | 9.79E-05 | 2.67E-02 |
| gene expression (GO:0010467) | 1448 | 18 | 42.28 | - | 0.43 | 2.99E-05 | 9.66E-03 |
| response to chemical (GO:0042221) | 2204 | 27 | 64.36 | - | 0.42 | 9.10E-08 | 4.66E-05 |
| system process (GO:0003008) | 1252 | 9 | 36.56 | - | 0.25 | 6.97E-08 | 4.08E-05 |
| nervous system process (GO:0050877) | 1001 | 6 | 29.23 | - | 0.21 | 3.07E-07 | 1.45E-04 |
| G-protein coupled receptor signaling pathway (GO:0007186) | 1196 | 7 | 34.92 | - | 0.2 | 1.76E-08 | 1.20E-05 |
| sensory perception (GO:0007600) | 876 | 5 | 25.58 | - | 0.2 | 1.36E-06 | 5.39E-04 |
| detection of stimulus (GO:0051606) | 769 | 2 | 22.46 | - | 0.09 | 1.01E-07 | 4.97E-05 |
| sensory perception of smell (GO:0007608) | 692 | 1 | 20.21 | - | 0.05 | 7.17E-08 | 3.83E-05 |
| detection of stimulus involved in sensory perception (GO:0050906) | 722 | 1 | 21.08 | - | 0.05 | 3.40E-08 | 2.20E-05 |
| sensory perception of chemical stimulus (GO:0007606) | 722 | 1 | 21.08 | - | 0.05 | 3.40E-08 | 2.09E-05 |

| GO molecular function complete | REFLIST (22191) | upload_1 (648) | upload_1 (expected) | upload_1 (over/under) | upload_1 (fold Enrichment) | upload_1 (raw P-value) | upload_1 (FDR) |
|---|---|---|---|---|---|---|---|
| dynein light chain binding (GO:0045503) | 12 | 10 | 0.35 | + | 28.54 | 1.50E-10 | 1.78E-07 |
| ATP-dependent microtubule motor activity, minus-end-directed (GO:0 | 8 | 6 | 0.23 | + | 25.68 | 1.26E-06 | 3.46E-04 |
| dynein heavy chain binding (GO:0045504) | 6 | 4 | 0.18 | + | 22.83 | 1.18E-04 | 1.61E-02 |
| dynein intermediate chain binding (GO:0045505) | 14 | 7 | 0.41 | + | 17.12 | 1.18E-06 | 3.50E-04 |
| ATP-dependent microtubule motor activity (GO:1990939) | 28 | 13 | 0.82 | + | 15.9 | 5.82E-11 | 1.04E-07 |
| dynein light intermediate chain binding (GO:0051959) | 18 | 8 | 0.53 | + | 15.22 | 4.00E-07 | 1.30E-04 |
| ATP-dependent microtubule motor activity, plus-end-directed (GO:000 | 20 | 7 | 0.58 | + | 11.99 | 7.76E-06 | 1.73E-03 |
| microtubule motor activity (GO:0003777) | 61 | 18 | 1.78 | + | 10.11 | 6.17E-12 | 2.20E-08 |
| motor activity (GO:0003774) | 99 | 19 | 2.89 | + | 6.57 | 9.31E-10 | 8.30E-07 |
| ATPase activity (GO:0016887) | 321 | 23 | 9.37 | + | 2.45 | 1.89E-04 | 2.41E-02 |
| pyrophosphatase activity (GO:0016462) | 601 | 35 | 17.55 | + | 1.99 | 2.21E-04 | 2.72E-02 |
| hydrolase activity, acting on acid anhydrides, in phosphorus-containin | 607 | 35 | 17.73 | + | 1.97 | 2.45E-04 | 2.91E-02 |
| hydrolase activity, acting on acid anhydrides (GO:0016817) | 609 | 35 | 17.78 | + | 1.97 | 2.54E-04 | 2.93E-02 |
| nucleoside-triphosphatase activity (GO:0017111) | 580 | 33 | 16.94 | + | 1.95 | 4.67E-04 | 4.76E-02 |
| ATP binding (GO:0005524) | 1055 | 55 | 30.81 | + | 1.79 | 5.87E-05 | 1.10E-02 |
| adenyl ribonucleotide binding (GO:0032559) | 1071 | 55 | 31.27 | + | 1.76 | 9.50E-05 | 1.47E-02 |
| adenyl nucleotide binding (GO:0030554) | 1075 | 55 | 31.39 | + | 1.75 | 9.92E-05 | 1.47E-02 |
| purine ribonucleoside triphosphate binding (GO:0035639) | 1321 | 66 | 38.57 | + | 1.71 | 3.78E-05 | 7.94E-03 |
| purine ribonucleotide binding (GO:0032555) | 1341 | 66 | 39.16 | + | 1.69 | 6.09E-05 | 1.09E-02 |
| purine nucleotide binding (GO:0017076) | 1346 | 66 | 39.3 | + | 1.68 | 6.39E-05 | 1.09E-02 |
| drug binding (GO:0008144) | 1204 | 59 | 35.16 | + | 1.68 | 1.63E-04 | 2.15E-02 |
| ribonucleotide binding (GO:0032553) | 1354 | 66 | 39.54 | + | 1.67 | 9.20E-05 | 1.49E-02 |
| nucleotide binding (GO:0000166) | 1519 | 70 | 44.36 | + | 1.58 | 2.88E-04 | 3.21E-02 |
| nucleoside phosphate binding (GO:1901265) | 1519 | 70 | 44.36 | + | 1.58 | 2.88E-04 | 3.11E-02 |
| small molecule binding (GO:0036094) | 1716 | 79 | 50.11 | + | 1.58 | 1.06E-04 | 1.52E-02 |
| anion binding (GO:0043168) | 1847 | 85 | 53.93 | + | 1.58 | 5.44E-05 | 1.08E-02 |
| carbohydrate derivative binding (GO:0097367) | 1504 | 69 | 43.92 | + | 1.57 | 3.66E-04 | 3.84E-02 |
| signal transducer activity (GO:0004871) | 1510 | 17 | 44.09 | - | 0.39 | 4.39E-06 | 1.04E-03 |
| molecular transducer activity (GO:0060089) | 1460 | 15 | 42.63 | - | 0.35 | 1.35E-06 | 3.43E-04 |
| receptor activity (GO:0004872) | 1427 | 13 | 41.67 | - | 0.31 | 2.72E-07 | 9.69E-05 |
| signaling receptor activity (GO:0038023) | 1331 | 10 | 38.87 | - | 0.26 | 5.88E-08 | 2.99E-05 |
| transmembrane signaling receptor activity (GO:0004888) | 1238 | 9 | 36.15 | - | 0.25 | 9.74E-08 | 4.34E-05 |
| transmembrane receptor activity (GO:0099600) | 1238 | 9 | 36.15 | - | 0.25 | 9.74E-08 | 3.86E-05 |
| G-protein coupled receptor activity (GO:0004930) | 1019 | 5 | 29.76 | - | 0.17 | 4.02E-08 | 2.39E-05 |

| **GO cellular component complete** | REFLIST (22191) | upload_1 (648) | upload_1 (expected) | upload_1 (over/under) | upload_1 (fold Enrichment) | upload_1 (raw P-value) | upload_1 (FDR) |
|---|---|---|---|---|---|---|---|
| axoneme part (GO:0044447) | 8 | 7 | 0.23 | + | 29.96 | 7.57E-08 | 6.55E-06 |
| axonemal dynein complex (GO:0005858) | 7 | 6 | 0.2 | + | 29.35 | 7.38E-07 | 5.23E-05 |
| dynein complex (GO:0030286) | 23 | 13 | 0.67 | + | 19.36 | 8.69E-12 | 1.23E-09 |
| intraciliary transport particle A (GO:0030991) | 8 | 4 | 0.23 | + | 17.12 | 2.65E-04 | 1.38E-02 |
| intraciliary transport particle (GO:0030990) | 24 | 9 | 0.7 | + | 12.84 | 2.36E-07 | 1.94E-05 |
| ciliary plasm (GO:0097014) | 47 | 17 | 1.37 | + | 12.39 | 1.62E-12 | 2.53E-10 |
| axoneme (GO:0005930) | 46 | 16 | 1.34 | + | 11.91 | 1.20E-11 | 1.55E-09 |
| intraciliary transport particle B (GO:0030992) | 15 | 5 | 0.44 | + | 11.42 | 1.97E-04 | 1.10E-02 |
| ciliary basal body (GO:0036064) | 54 | 16 | 1.58 | + | 10.15 | 8.78E-11 | 9.77E-09 |
| plasma membrane bounded cell projection cytoplasm (GO:0032838) | 65 | 17 | 1.9 | + | 8.96 | 1.19E-10 | 1.23E-08 |
| motile cilium (GO:0031514) | 73 | 18 | 2.13 | + | 8.44 | 7.71E-11 | 9.24E-09 |
| sperm flagellum (GO:0036126) | 47 | 11 | 1.37 | + | 8.01 | 5.97E-07 | 4.65E-05 |
| 9+2 motile cilium (GO:0097729) | 47 | 11 | 1.37 | + | 8.01 | 5.97E-07 | 4.43E-05 |
| microtubule associated complex (GO:0005875) | 88 | 19 | 2.57 | + | 7.39 | 1.64E-10 | 1.60E-08 |
| cilium (GO:0005929) | 263 | 55 | 7.68 | + | 7.16 | 2.07E-27 | 3.23E-24 |
| ciliary part (GO:0044441) | 196 | 38 | 5.72 | + | 6.64 | 2.63E-18 | 2.05E-15 |
| ciliary transition zone (GO:0035869) | 32 | 6 | 0.93 | + | 6.42 | 6.49E-04 | 3.16E-02 |
| sperm part (GO:0097223) | 90 | 12 | 2.63 | + | 4.57 | 3.24E-05 | 1.87E-03 |
| cytoplasmic region (GO:0099568) | 156 | 18 | 4.56 | + | 3.95 | 2.46E-06 | 1.67E-04 |
| cell projection part (GO:0044463) | 404 | 45 | 11.8 | + | 3.81 | 2.37E-13 | 4.62E-11 |
| plasma membrane bounded cell projection part (GO:0120038) | 404 | 45 | 11.8 | + | 3.81 | 2.37E-13 | 4.11E-11 |
| microtubule cytoskeleton (GO:0015630) | 525 | 58 | 15.33 | + | 3.78 | 9.70E-17 | 3.78E-14 |
| microtubule organizing center (GO:0005815) | 313 | 34 | 9.14 | + | 3.72 | 3.89E-10 | 3.56E-08 |
| plasma membrane bounded cell projection (GO:0120025) | 652 | 65 | 19.04 | + | 3.41 | 1.29E-16 | 4.02E-14 |
| cell projection (GO:0042995) | 662 | 65 | 19.33 | + | 3.36 | 2.55E-16 | 6.62E-14 |
| microtubule (GO:0005874) | 135 | 13 | 3.94 | + | 3.3 | 3.15E-04 | 1.58E-02 |
| cytoskeletal part (GO:0044430) | 750 | 71 | 21.9 | + | 3.24 | 5.58E-17 | 2.90E-14 |
| cytoskeleton (GO:0005856) | 955 | 80 | 27.89 | + | 2.87 | 3.26E-16 | 7.25E-14 |
| polymeric cytoskeletal fiber (GO:0099513) | 264 | 19 | 7.71 | + | 2.46 | 7.75E-04 | 3.66E-02 |
| intracellular non-membrane-bounded organelle (GO:0043232) | 2036 | 93 | 59.45 | + | 1.56 | 2.63E-05 | 1.64E-03 |
| non-membrane-bounded organelle (GO:0043228) | 2036 | 93 | 59.45 | + | 1.56 | 2.63E-05 | 1.57E-03 |
| membrane-bounded organelle (GO:0043227) | 6158 | 138 | 179.82 | - | 0.77 | 2.49E-04 | 1.34E-02 |
| intracellular membrane-bounded organelle (GO:0043231) | 5322 | 109 | 155.41 | - | 0.7 | 1.57E-05 | 1.02E-03 |
